# Formalized scientific methodology enables rigorous AI-conducted research across domains

**DOI:** 10.64898/2026.03.02.709102

**Authors:** Yanlin Zhang, Jing Zhao

## Abstract

We formalize scientific methodology—the end-to-end process from question formulation to evidence-grounded writing—as a phase-gated research protocol with explicit return paths and persistent constraints, and instantiate it for general-purpose language models as executable protocol specifications. The formalization decomposes methodology into three complementary layers: a procedural workflow, an integrity discipline, and project governance. Encoded as protocol specifications^$^ and activated across the lifecycle, these constraints externalize planning and verification artifacts and make integrity-relevant interventions auditable. We validate the approach in six end-to-end projects, including a matched controlled study, where the same agent produced two complete papers with and without the protocol. Across domains, the protocol-constrained agent produced evidence-backed, auditable research outputs—including closed-form derivations, quantitative ablations that resolve modeling design choices, and algorithmic refactors that preserve the objective while changing the computational primitive. In population-genomic applications, it also recovered well-studied biological signals as validity checks, including known admixture targets in the 1000 Genomes Project and Neanderthal-introgressed immune loci on chromosome 21 consistent with prior catalogs. In the controlled study, the protocol-free baseline could still produce a complete manuscript, but integrity-relevant risks were easier to introduce and harder to detect when constraints and artifacts were absent.

## Introduction

Doing good science requires two kinds of knowledge. The first—*what* is known in a field—is encoded in textbooks, papers, and databases. The second—*how* to generate reliable new knowledge—is the methodology that governs research practice: formulate research questions that matter, lock evaluation criteria before running experiments, report all results including failures, exclude alternative explanations before claiming mechanisms, ensure every assertion rests on specific evidence. This procedural and normative knowledge is what separates a publishable study from an exploratory exercise. Yet while much of it can in principle be articulated—and fragments have been codified in pre-registration protocols [1], registered reports [2], and reporting checklists [3, 4]—it is most commonly learned and refined through apprenticeship: PhD training, mentorship, lab culture, and the accumulated experience of reviewer feedback [5–7].

For agentic research, this apprenticeship-based transmission creates a more immediate problem: key methodological steps are often implicit, context-dependent, and enforced socially (through advisor feedback, lab norms, and peer review) rather than encoded as explicit, checkable rules. When these expectations are not externalized, an agent can complete a project while silently skipping steps that humans would normally insist on (e.g., freezing evaluation criteria before experiments, reporting negative results with equal prominence, or mapping claims to concrete evidence). The result is not an inability to produce manuscripts, but a gap in auditability and process discipline that makes failures harder to detect, comparisons harder to run fairly, and iteration harder to govern.

Scientific discovery follows a generative cycle: a researcher formulates a question, designs experiments to address it, collects observations, and then integrates those observations with existing knowledge to produce new understanding. Large language models have now demonstrated expert-level knowledge across numerous scientific domains [8, 9]—they possess, in effect, the first ingredient of this cycle. What is less reliably expressed in autonomous settings is the second: the procedural knowledge of *how* to move from domain expertise to reliable new findings, including when to stop, verify, and backtrack. If LLMs can be taught to formulate questions, design and execute experiments, verify results, and reason from evidence to conclusions—much as a graduate student learns from an experienced mentor—then the combination of broad domain knowledge with sound methodology should enable genuine knowledge creation.

Artificial intelligence is already transforming science, but current AI systems encode scientific *content* and task *capability* rather than an end-to-end, auditable scientific *methodology*. At one end, domain-specific systems achieve deep integration with particular fields: AlphaFold embeds protein physics [10, 11], autonomous laboratories encode reaction rules [12, 13], and biomedical platforms such as Biomni [14] orchestrate hundreds of specialized tools across 25 subfields to execute tasks from drug repurposing to molecular cloning. At the other, AI scientist systems pursue research autonomy at increasing scale: The AI Scientist [15] generates and peer-reviews machine learning papers, its successor produced the first AI-generated peer-reviewed publication [16], Kosmos [17] performs 12-hour autonomous discovery sessions equivalent to months of human research, Robin [18] achieved the first fully automated biological discovery, and Google’s AI co-scientist [19] uses multi-agent debate and tournament evolution to generate and rank research hypotheses for biomedical discovery. Between these, bioinformatics agents [20, 21] automate standard computational pipelines, while deep research products [22] synthesize literature at unprecedented scale. These systems represent genuine and rapid progress in what AI can *do* in science. What often remains under-specified is *how* AI should do it: the persistent methodological constraints—evaluation immutability, complete reporting, claim–evidence alignment, alternative-hypothesis exclusion—that distinguish reliable knowledge from plausible-sounding output. Without such constraints, capable systems can more easily accumulate integrity-relevant risks (e.g., metric drift, incomplete reporting, unverified references, and claim–evidence mismatches) [23, 24]. Existing experiment tracking tools [25] and FAIR data principles [26] address computational reproducibility, and multi-agent frameworks improve coordination [27, 28], but these tools do not encode an end-to-end research methodology as an executable set of norms that can be audited turn by turn. Moreover, as with human researchers, the most consequential—and most difficult—step in the entire cycle is formulating a good scientific question. A single LLM mining the literature often tends toward conservative, incremental questions; the methodology formalization we propose addresses this through structured multi-agent deliberation and explicit ideation strategies that generate divergent candidate directions before convergence.

Here we ask whether the articulable components of scientific methodology—the procedural rules, integrity norms, and governance practices that experienced researchers follow—can be operationalized end-to-end as an executable, phase-gated protocol with persistent methodological constraints, and transferred to general-purpose AI agents as a modular specification they execute step by step. The key insight is twofold. First, the complete research process—from question formulation through direction validation, method design, iterative experimentation, failure management, results integration, and evidence-grounded writing— can be decomposed into three complementary constraint layers (procedural workflow, integrity discipline, and project governance) and specified as an explicit protocol rather than an ad hoc interaction. Second, across our end-to-end validation projects the protocol condition externalizes intermediate planning and verification artifacts, enforces phase transitions and explicit backtracks when evidence is stale, and helps surface and correct claim–evidence mismatches relative to matched protocol-free runs. We applied this formalization to six end-to-end projects spanning diverse domains, without domain-specific modification: five protocol-constrained projects executed under the full methodology plus one controlled study that produced two complete papers under matched conditions with and without the protocol (Supplementary Papers 1– 5 and 6A/6B). Across these projects, the protocol-constrained agent produced evidence-backed, auditable research outputs—including closed-form derivations, quantitative ablations that resolve modeling design choices, and algorithmic refactors that preserve the objective while changing the computational primitive— and recovered well-studied biological signals as validity checks. However, it also inherits well-known failure modes of large language models—including simple arithmetic or consistency errors that can be introduced during manuscript drafting and may only be surfaced through multi-round, adversarial review. These validation runs suggest that the formalization itself—not merely the underlying model capability—materially contributes to methodological rigour and auditability, as matched comparison runs without constraints exhibited more frequent integrity-relevant risks. Human researchers may participate at validation gates to contribute scientific judgment that remains difficult to formalize (scientific taste, intuition about significance [5]), but the methodology functions as a self-consistent system: the interlocking constraints guide the AI through the full research lifecycle, with each layer catching distinct failure modes. We instantiate this formalization as an open-source tool, *Amplify*, which implements the protocol and produces auditable artifacts; all validation projects and the controlled study in this paper are executed via Amplify.

## Results

We propose a formalized, executable research methodology for general-purpose AI agents, implemented as a phase-gated protocol with persistent constraints. The protocol structures the full lifecycle into seven phases with explicit return paths, maintains always-on integrity and governance invariants, and embeds role-specialized multi-agent deliberation checkpoints that can trigger revision and backtracking. In a tool-enabled environment (IDE with web search/browsing, literature reading, and a shell for execution), the agent is permitted to iteratively gather evidence, run analyses, verify claims against fresh computations

### Decomposing scientific methodology into functional layers

To formalize scientific methodology, we first needed to determine whether its constituent practices share a structure amenable to decomposition. We analysed the methodological norms enforced by experienced researchers—as documented in integrity guidelines [29, 30], reporting standards [3, 4], and pre-registration frameworks [1, 2]—and observed that they cluster around three distinct functional concerns: *what to do next* (procedural sequencing), *what constraints must always hold* (integrity norms), and *whether the project should continue* (strategic governance).

We use this three-part organization as a practical decomposition: each concern can be specified and enforced via a corresponding constraint layer. This separation is not merely a convenient taxonomy but reflects how methodology functions in practice. In well-functioning human research groups, procedural planning (the PI’s research plan), integrity enforcement (lab culture, peer norms), and strategic oversight (committee review, self-assessment) are typically carried out by different actors through different mechanisms [7]. We formalized each concern as a separate constraint layer applicable to AI agents: a **procedural workflow** of seven phase-gated phases with explicit return paths, an **integrity discipline** of seven persistent constraints, and a **governance** layer of four strategic functions (Fig. 1).

**Figure 1.**
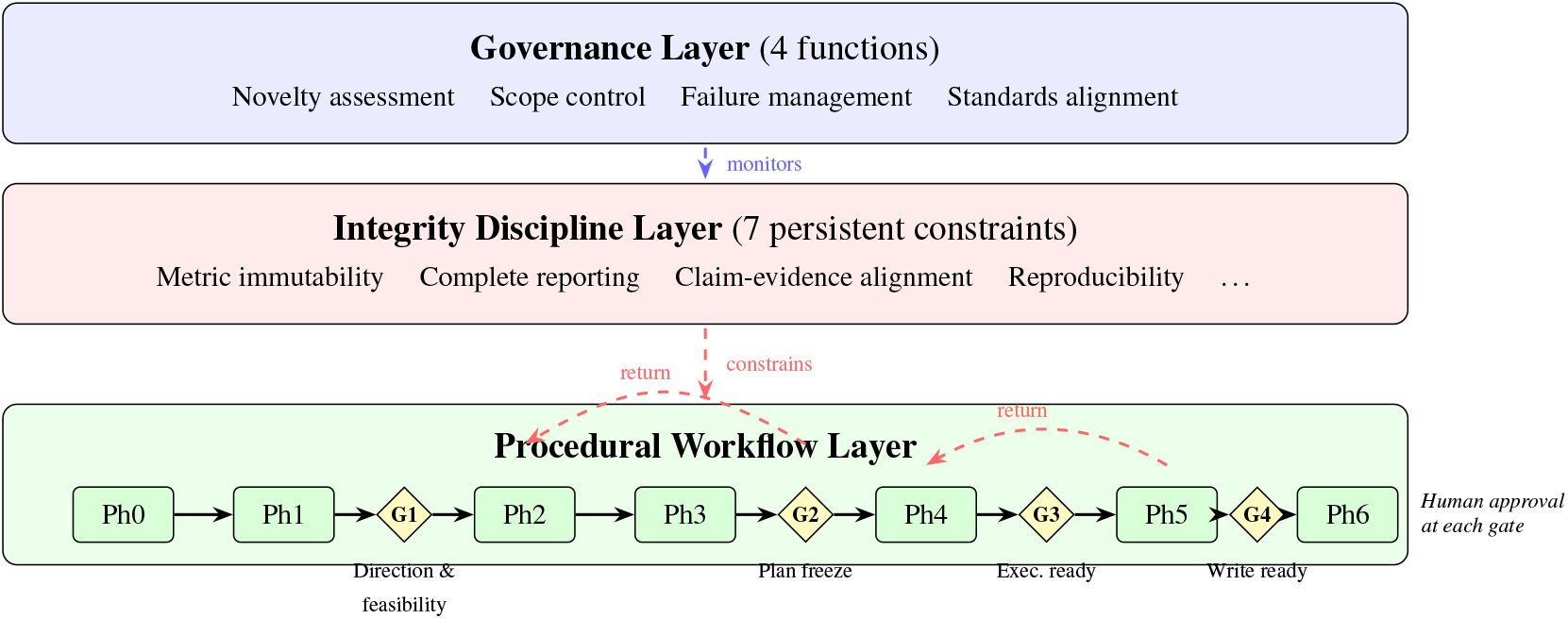
Three-layer methodology formalization with validation gates. Scientific methodology is decomposed into three complementary constraint layers: a procedural workflow (seven phase-gated phases with explicit return paths), an integrity discipline (seven persistent constraints active throughout), and a governance layer (four strategic oversight functions). Four mandatory gates (G1–G4, diamonds) require human approval at critical transitions, preserving scientific judgment while the formalization provides methodological discipline. The integrity and governance layers operate continuously across all phases. and references, and revise artifacts until gate criteria are satisfied. We evaluate the approach through five end-to-end protocol-constrained projects spanning multiple domains and research types, and a matched controlled study (Project 6) in which the same task and environment produced two complete papers with and without the protocol; the six projects and manuscripts are summarized in Supplementary Table S1. Across the validation projects, the protocol externalizes intermediate planning and verification artifacts and makes integrity-relevant interventions auditable. In the controlled study, the protocol-free baseline could still produce a complete manuscript, but the protocol condition more consistently enforced domain-target alignment and verification obligations by requiring those intermediate artifacts and return paths to be made explicit and checkable. Importantly, beyond process-level interventions, these projects produced evidence-backed research outputs that are auditable in the generated papers (Table 2), including analytical derivations, quantitative ablations, algorithmic refactors, and recovery of well-studied biological signals as validity checks.

**Table 1.** Integrity discipline constraints and the failure modes they address. The final column summarizes examples of enforcement drawn from project artifacts and logs.

| Constraint | Failure mode addressed | Observed enforcement |
| --- | --- | --- |
| Metric immutability | Post hoc evaluation changes | Protocol locked at G2; changes require explicit authorization. |
| Complete reporting | Selective reporting | Negative results retained and disclosed (gpuADMIX; ArchaicPainter positive-only emission ablation). |
| Claim-evidence alignment | Unsupported assertions | Paper text reconciled with code/results; mismatches corrected. |
| Alternative-hypothesis exclusion | Confounded causal claims | Alternatives enumerated; discriminating tests required (DESI). |
| Reproducibility | Unreproducible computation | Seeds + environment/log capture + scriptable reruns required. |
| Verification | Unverified status claims | Fresh checks required; references verified; re-compute/re-compile before claims. |
| Figure standards | Substandard visualizations | Readable, consistent figures with explicit inclusion/caption checks before final compilation. |

**Table 2.** Evidence-backed cross-domain research outputs produced under the protocol. Each entry is stated narrowly and paired with an explicit evidence anchor in the corresponding Supplementary Paper.

| Project | Output type | Narrow, checkable claim | Evidence anchor |
| --- | --- | --- | --- |
| gpuADMIX | Algorithmic refactor | The binomial-likelihood EM updates can be rearranged into a small fixed set of GEMM (matrix multiplications) plus elementwise operations, making the original objective GPU-native without changing the model. | Supp. Paper 1 |
| ArchaicPainter | Ablation conclusion | In introgression HMM emission, treating mismatches as informative (bidirectional emission) is necessary for segment discriminability when the reference matches the donor; down-weighting mismatches reduces simulation F1 from 0.480 to 0.154. | Supp. Paper 2 |
| HapGraph | Analytical correction | For cross-population IBD between two modern lineages, the expected mean segment length scales as $50/T$ cM (not $100/T$); with detection threshold $\ell_{\min}$, $E[L \mid L > \ell_{\min}] = 50/T + \ell_{\min}$. | Supp. Paper 4 |
| z2-quantum-mpemba | Closed-form derivation | For product initial states, the initial $Z_2$ entanglement asymmetry admits a closed-form expression bounded by $\ln 2$, providing an exact calibration scale for numerical experiments. | Supp. Paper 5 |
| DESI | Robust observation (+ caveat) | Window-level mean pairwise TMRCA is substantially deeper within AFR (1,229 ka) than EAS (859 ka) and EUR (920 ka) across all 22 autosomes; a FitCoal-parameter stress test suggests mismatch under a limited model class and summary statistics, motivating follow-up with better-matched simulation and likelihood-based fitting. | Supp. Paper 3 |

We evaluate this decomposition through cross-domain application, and organize the Results below by mapping failure modes observed when constraints are absent to the layer(s) designed to prevent them.

### Procedural workflow mitigates the risk of collapsing research into undifferentiated generation

The procedural workflow encodes seven phase-gated phases (domain anchoring, direction exploration, problem validation, method design, experiment execution, results integration, paper writing) with explicit return paths for backtracking, each with defined entry conditions, deliverables, and exit criteria (Fig. 2). Temporal ordering is enforced: experiments cannot begin before the evaluation protocol is frozen, and paper writing cannot begin before results are integrated.

**Figure 2.**
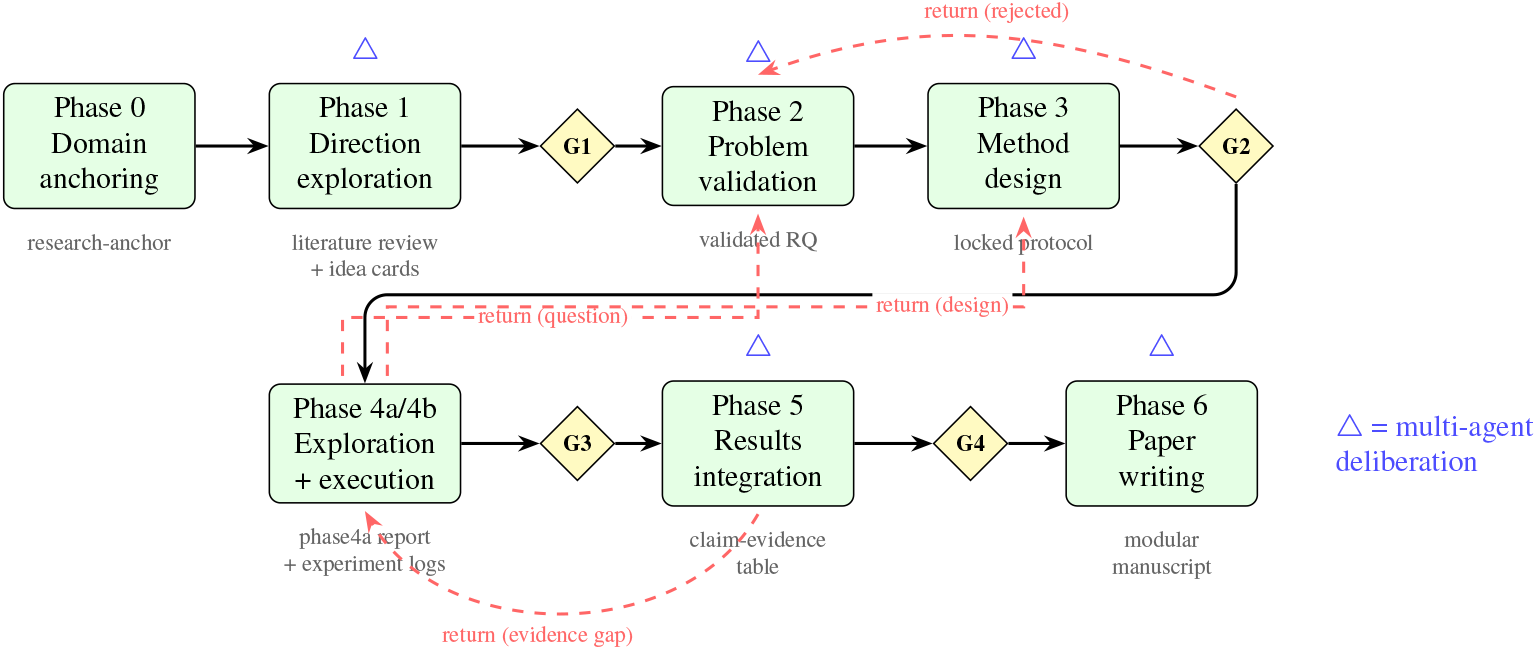
Seven-phase procedural workflow with deliverables. Each phase produces defined artifacts (grey text). Gates (diamonds) require human approval before progression. Blue triangles mark phases where multi-agent deliberation panels are deployed. Phase 4 is explicitly two-stage: Phase 4a exploratory probing that may trigger returns to refine the question or design, and Phase 4b full execution. Red dashed arrows show return paths triggered by governance interventions (e.g., plan rejection at G2), Phase 4a findings (question/design refinement), or evidence gaps identified during integration (Ph5 →Ph4). The workflow enforces temporal ordering: evaluation protocols are locked at G2 before experiments begin, and evidence sufficiency is verified at G4 before writing begins. frameworks [1, 2]—and observed that they cluster around three distinct functional concerns: *what to do next* (procedural sequencing), *what constraints must always hold* (integrity norms), and *whether the project should continue* (strategic governance).

This phase-gated structure addresses what we observe as the most consequential failure mode when AI agents conduct research without procedural constraints. In the protocol-free baseline of our controlled study (Project 6), the agent produced a complete manuscript while explicitly interleaving drafting with ongoing data extraction and analysis—beginning to write and compile the paper before the full analysis had finished, and then revising the draft as additional results arrived. In this baseline condition, intermediate deliverables (e.g., a locked protocol, sufficiency criteria, or an integration blueprint) were not externalized, reducing auditability and making goalpost-moving and stale-claim reuse difficult to detect. With the procedural workflow imposed, the protocol requires explicit literature grounding, direction validation, evaluation-protocol locking before experimentation, and iteration/backtracking when evidence is insufficient—behaviours driven by the constraints rather than by additional model capability.

Among the phases of the research cycle, question formulation (direction exploration and problem validation) proved both the most critical and the most difficult to automate well. In our projects, a single-agent literature review tended to produce conservative, incremental questions—extensions of existing work that minimize risk but also minimize novelty. We address this by combining structured multi-agent deliberation during Phase 1 with six explicit ideation strategies (contradiction mining, assumption challenging, cross-domain transfer, limitation-to-opportunity conversion, counterfactual reasoning, and trend extrapolation). In Project 3 (DESI), multi-agent brainstorming merged two initially separate directions into a more ambitious unified proposal; in Project 2 (ArchaicPainter), the divergent phase generated multiple candidate directions before convergence on the Li & Stephens extension that proved most productive.

### Integrity discipline makes integrity checks explicit and auditable

The integrity discipline comprises seven persistent constraints that activate at defined trigger points and remain active until project completion. Each targets a documented threat to research validity, but critically, each is enforced through explicit artifacts and halting/backtracking behaviour rather than implicit “best effort” diligence (Table 1).

Across the protocol-constrained projects, these constraints were triggered repeatedly and resulted in concrete interventions: negative results were retained and disclosed (e.g., ArchaicPainter’s failed “positive-only” emission variant and gpuADMIX’s documented bugs/limitations), skeptical checks forced explicit alternative-hypothesis testing (DESI), reference and manuscript audits corrected unsupported citations and code–text mismatches, and verification steps enforced fresh checks (including reference verification and final compilation sanity checks) before claims were finalized.

### Governance prevents indefinite continuation of failing approaches

The governance layer encodes four strategic functions: novelty assessment, scope control, failure management, and standards alignment. In our projects, governance repeatedly forced strategic self-assessment to be made explicit and auditable: novelty checks triggered redesigns when an idea was not yet publishable; scope control recorded and justified exclusions; and standards-alignment checks translated evidence sufficiency into concrete venue decisions and documented limitations when baselines or supplements could not be completed.

The failure-management function proved particularly important: it enforces structured reassessment after repeated setbacks and presents explicit pivot/downgrade/stop options, rather than allowing unbounded iteration to remain implicit. This mechanism mirrors the role of thesis-committee reviews and lab-group critiques, but externalizes the decision in a form that can be reviewed after the fact.

### Validation gates provide structured checkpoints

The formalization includes four gates between major phases (G1–G4), each defining explicit criteria that must be satisfied before progression (Fig. 1). **G1** validates research direction, scientific significance, and resource feasibility. **G2** locks the evaluation protocol. **G3** confirms data quality and resource readiness. **G4** verifies that evidence is sufficient for writing.

These gates function as quality checkpoints within the methodology. In our validation projects, a human researcher reviewed gate criteria and provided approval or redirection—and this proved valuable, particularly when the researcher challenged an insufficiently novel plan at G2 (see Vignette 2). However, the gates are architecturally part of the methodology’s procedural logic, not a separate human-dependence mechanism: the criteria they enforce (e.g., “evaluation protocol must be locked before experimentation begins”) are themselves formalized constraints. The primary contribution is the methodology that structures what happens between gates—the phases and their explicit return paths, the integrity discipline, the governance interventions—which transforms AI behaviour regardless of who or what evaluates the gate criteria. Human participation at gates enhances quality by contributing scientific judgment that is difficult to formalize, but the methodology provides the research discipline that makes gate evaluation meaningful in the first place.

### Methodology formalization adapts across research types

A single set of methodological principles must accommodate the genuine diversity of research practice to be considered a valid formalization. Performance-driven method development requires different procedures from story-driven scientific discovery or utility-driven tool evaluation. We accommodated this diversity through differential activation: the integrity discipline and governance layers operate identically across all research types (metric immutability and complete reporting are universal), while the procedural workflow adapts. **Method** projects emphasize evaluation protocol locking, baseline reproduction, and mandatory iteration (minimum three diagnose–hypothesize–fix–measure cycles). **Discovery** projects emphasize analysis storyboard design, alternative hypothesis exclusion, and narrative coherence. **Tool** projects emphasize benchmarking, usability, and documentation. **Hybrid** projects activate both Method and Discovery tracks.

This structure suggests an organizing principle: research methodology has a *universal core* of integrity and governance norms, with *type-specific procedural instantiations* that adapt to the nature of the knowledge claim being made.

### Structured multi-agent deliberation provides repeated, role-differentiated review checkpoints

In human research, critical decisions are repeatedly stress-tested through discussions at multiple points in the workflow (lab meetings, advisor feedback, collaborator critique), not only at the journal-review stage. We operationalize an analogous mechanism by embedding structured multi-agent deliberation at five critical junctures. At each juncture, the system dynamically instantiates role-specialized *sub-agents* conditioned on the research question and current project state (e.g., a domain expert, a skeptical critic, and an editor). Crucially, each sub-agent is given an independent context and evaluates the same artifact (protocol, plan, results integration blueprint, or manuscript section) against a shared rubric without sharing its intermediate reasoning with the others. Deliberation proceeds in structured rounds (maximum five): convergence requires unanimous PASS; otherwise, the artifact is modified and *all* sub-agents re-assess the full artifact. The resulting recommendations can trigger explicit returns to earlier phases (e.g., rerun experiments, redesign analyses, or retract unsupported claims). Unresolved disagreements are escalated to the human researcher.

This draws on evidence that diverse multi-agent panels improve reasoning quality [31–33] and extends the principle from single-query tasks to sustained, multi-phase research where accumulated context matters.

### Cross-domain validation

We applied the complete methodology formalization to research projects spanning diverse scientific domains and research types, without any domain-specific modification (Supplementary Papers; Fig. 5). In each protocol-constrained project, a general-purpose LLM operating under the full constraints conducted the complete research lifecycle. We report here on five protocol-constrained projects spanning population genomics, computational bioinformatics, human evolutionary genetics, computational population genetics, and condensed-matter physics. We additionally report a sixth project (Project 6): a controlled study on the same dataset and task, in which the same AI agent produced two complete manuscripts under matched conditions with and without the protocol (Supplementary Papers 6A/6B).

Across these projects, we focus on a small set of evidence-backed research outputs that are directly auditable in the generated papers: (i) closed-form analytical derivations that can be independently checked, (ii) quantitative ablations that resolve specific modeling design choices, (iii) algorithmic refactors that preserve the objective but change the computational primitive, and (iv) domain analyses that recapitulate known signals while surfacing limitations and inconsistencies transparently. Table 2 summarizes these outputs and provides an evidence anchor (equation/figure/table) for each.

#### Project 1 (gpuADMIX, Type H—Method + Tool)

GPU-accelerated ancestry estimation preserving the exact ADMIXTURE binomial likelihood. A key method insight is that the EM updates can be algebraically reorganized into a small fixed set of GEMM operations plus elementwise steps, so the original binomial objective becomes GPU-native without changing the model. The constrained agent developed a complete software tool (Python/PyTorch), conducted ablation experiments across five random seeds at each of nine *K* values, and produced a publication-ready manuscript. The procedural workflow enforced evaluation protocol locking before experiments, and the integrity discipline required reporting all seeds including failures. A critical integrity event occurred when stored results for *K* ≥ 8 were discovered to originate from an earlier code version; the verification constraint forced re-measurement, which reversed the paper’s narrative from “limited at high *K*” to “competitive at all *K* via multi-seed strategy.” Multi-agent deliberation panels caught two fatal citation errors (one entirely fabricated reference), a formula mismatch between code and text (FISTA momentum vs. simple Nesterov), and a misattributed benchmark figure—all before the manuscript was finalized. The resulting tool achieved a 213*×* speedup over ADMIXTURE and 41*×* over fastmixture while maintaining *Q*-matrix correlation with ADMIXTURE outputs *r*^2^ *>* 0.9999.

#### Project 2 (ArchaicPainter, Type H—Method + Discovery

A three-state Hidden Markov Model extending the Li & Stephens haplotype-matching framework to archaic reference genomes for per-haplotype introgression detection. The constrained agent developed the method, validated it on 50 independent simulation replicates (*F*_1_ = 0.480 *±* 0.095; 56*×* improvement over a density baseline, Wilcoxon *p* = 8.9 *×* 10^−16^), and applied it to chromosome 21 of the 1000 Genomes Project. A central design conclusion is that mismatches to the archaic reference are not mere noise but carry discriminative signal when the reference is a faithful proxy for the donor: down-weighting mismatches via a positive-only emission reduces simulation F1 to 0.154 *±* 0.079. Multi-agent deliberation identified three fatal and four major evidence gaps, forcing the agent back from Phase 5 to Phase 4 for supplemental experiments. The human researcher overrode the requirement for direct tool comparison (HMMix/DAIseg installation failed), and the governance layer ensured this was documented as an acknowledged limitation with a corresponding venue downgrade. Notably, the method independently recovered ∼2% Neanderthal ancestry in European and East Asian populations— consistent with published estimates from independent methods [34]—and detected the complete interferon receptor gene cluster (*IFNAR1, IFNAR2, IFNGR2, IL10RB*) as a Neanderthal-introgressed region in Europeans, corroborating published adaptive introgression findings [35].

#### Project 3 (DESI, Type D—Discovery)

A window-based coalescent-depth analysis quantifying genome-wide mean pairwise TMRCA differences across super-populations. This project exhibited the most consequential governance interventions. At G2, the human researcher challenged the proposed plan (“is there actually a new method here, or are you just running existing tools?”), triggering a complete project redesign. During execution, the AI agent retracted its own interpretation of initial results, explicitly stating “my earlier interpretation was wrong” and “is this enough for *Nature Genetics*? No—not from this angle.” The resulting paper grounds its claims through simulation calibration and chromosome-wide consistency checks: within-AFR mean TMRCA is 1,229.5 *±* 6.4 ka, deeper than within-EAS (858.8 *±* 5.4 ka) and within-EUR (920.2 *±* 5.5 ka), with the AFR−EAS difference (370.7 *±* 8.4 ka) positive on all 22 autosomes. The paper further proposes—based on a parameter scan of a *panmictic* ∼930 ka bottleneck model and window-level summary statistics (e.g., 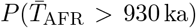 and 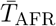)—that the bottleneck may be less severe than Fit-Coal’s published parameterization. However, after manual review we note that this inference depends on the specific summary-statistic comparison and simulation setup, and that the simulated data use a much smaller number of windows than the real-data analysis, which may bias the tail-fraction estimate; we therefore treat this result as a suggestive direction for follow-up with better-matched simulation and likelihood-based fitting rather than a decisive conclusion.

#### Project 4 (HapGraph, Type C—Tool)

A Bayesian admixture graph inference tool unifying F-statistics and inter-population IBD sharing for joint estimation of graph topology and admixture proportions (*α*), with an IBD-based timing model for *T* on fixed topologies. This project demonstrated the integrity discipline’s role in iterative bug discovery and correction. During Phase 4a prototype testing, the verification constraint identified that the F_2_ path-length likelihood approximation was insensitive to admixture—a critical implementation flaw that rendered the tool unable to estimate *α*. Rather than proceeding with a partially working system, the constraint forced a complete rewrite to an ancestry-vector formulation with NNLS branch-length fitting, followed by systematic correction of two additional issues (F_3_ bias correction per Patterson et al. 2012, and IBD source-clade filtering) through three diagnose–fix–measure iterations in Phase 4b. A technically important correction in this project is the cross-population IBD length scale: for two modern lineages separated by *T* generations, the expected mean length is 50*/T* cM, and under a detection threshold *ℓ*_min_ it shifts to 50*/T* + *ℓ*_min_. Multi-agent deliberation on the manuscript draft then caught critical discrepancies between the paper text and the actual code and forced reconciliation before the manuscript could pass the quality gate. On simulation benchmarks (S1–S3, 20 seeds each), HapGraph achieved 100% topology accuracy at *K* ≤ 2 (where *K* denotes the number of admixture edges), *α* MAE of 0.054, and *T* MAE of 10.9 generations with *T* 95% CI coverage of 93%. Applied to 26 populations from the 1000 Genomes Project, the tool identified *K* = 8 admixture events without manual curation; seven of the eight targets correspond to well-studied admixed populations (ASW, ACB, PUR, CLM, MXL, BEB, PJL), and timing estimation for real data was deferred pending whole-genome IBD pre-computation.

#### Project 5 (z2-quantum-mpemba, Type D—Discovery)

A condensed-matter physics study of the Z_2_ quantum Mpemba effect in integrable one-dimensional free-fermion chains (transverse-field Ising model and anisotropic XY chain). The constrained agent derived an exact closed-form expression for the initial entanglement asymmetry and verified it to machine precision, then used exact diagonalization to establish that Z_2_ QME is present across all 23 post-quench field values tested—including the DQPT-free ferromagnetic phase. Statistical analysis of 671 initial-state pairs revealed a strong correlation between the Mpemba crossing time *t*_*M*_ and the initial entanglement-asymmetry imbalance (Spearman *ρ* = +0.90, *p* ≈ 3 *×* 10^−240^), while showing that DQPT singularities modulate but do not govern crossings (KS *p* ≈ 6 *×* 10^−25^; proximity test *p* = 0.27). Finite-size scaling over *N* = 8–18 supported persistence in the thermodynamic limit. Notably, despite modest computational runtime (order tens of minutes on a 128-core CPU server), this project required sustained analytical reasoning and exact-solution verification, consistent with the effort distribution in human theoretical physics.

Across the five protocol-constrained projects (Supplementary Table S1), the procedural workflow completed all seven phases with all four gates enforced in each. The integrity discipline repeatedly halted progression until verification and documentation requirements were satisfied (Table 1), and multi-agent deliberation served as an embedded adversarial review mechanism that surfaced actionable gaps (evidence insufficiency, code–text mismatches, and citation integrity issues) before manuscripts were finalized. Governance interventions—including methodology pivots, scope reductions, and venue-alignment downgrades when evidence was insufficient—were required in multiple projects, illustrating that sustained strategic self-assessment can be operationalized as part of the protocol rather than left to ad hoc prompting.

### Controlled study (Project 6): protocol increases auditability and methodological structure

To isolate the effect of the protocol from underlying model capability, we report a matched controlled study (Project 6) in which the sole manipulated factor is activation of the formalized constraints. The same base model (Claude Opus 4.6) operating in the same IDE (Cursor) was tasked with conducting autonomous scientific research on the 1000 Genomes Project 2022 high-coverage dataset and producing a complete manuscript. In the protocol-free baseline condition, the agent produced a technically solid, indel-focused analysis (target standard: *Genomics*) with a single-file manuscript and a set of analysis scripts. In the Cursor+Amplify condition, the agent produced a modular manuscript and, crucially, externalized the methodological process as auditable protocol artifacts (intake anchor, literature review and gap analysis, explicit planning documents, and an integration blueprint), enabling structured iteration and clearer evidence-to-claim traceability (Table 3). Both complete manuscripts (with protocol vs. protocol-free) are included as Supplementary Papers 6A and 6B.

**Table 3.**
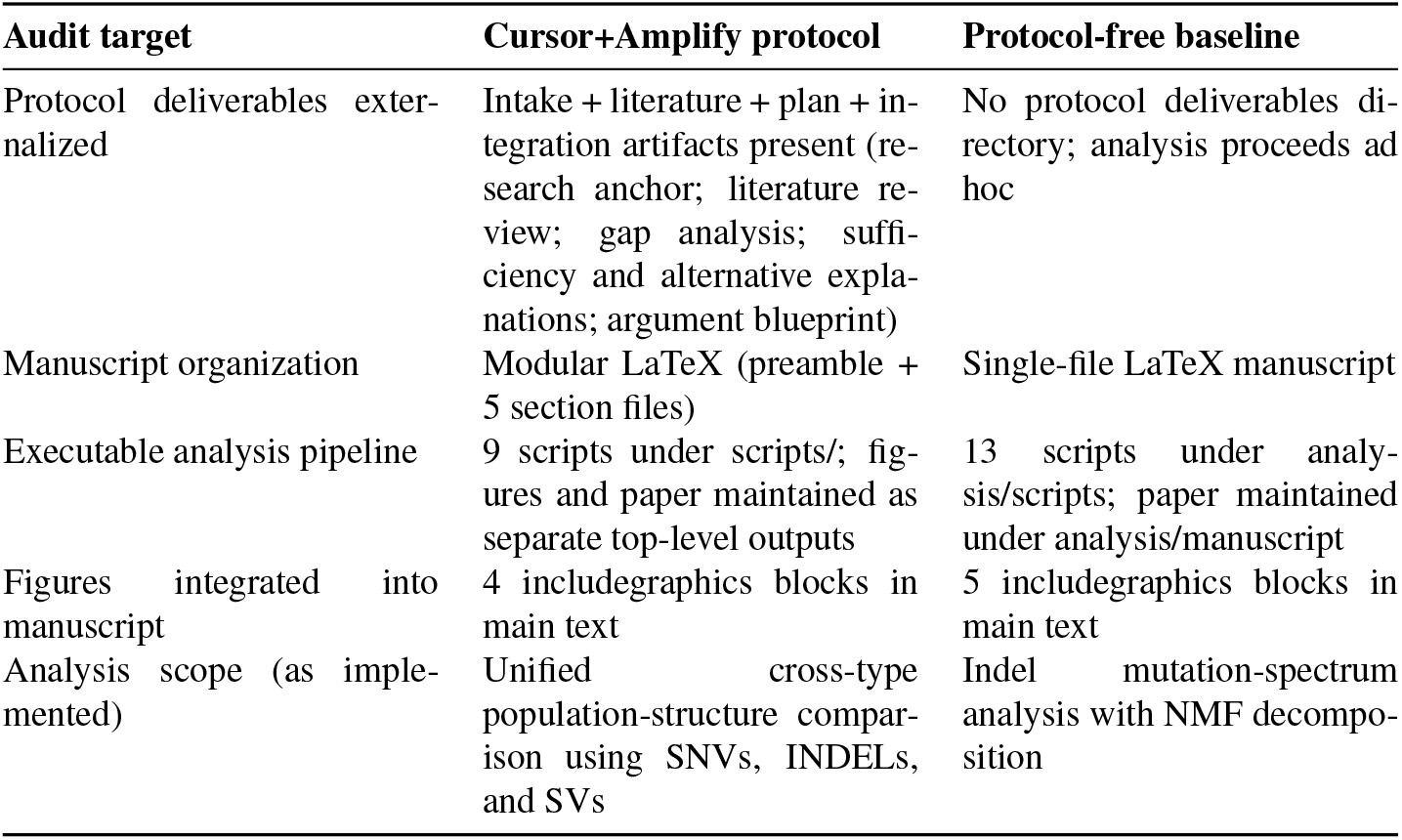
Controlled study: artifact-level comparison of protocol-free baseline versus Cursor+Amplify. Both conditions used the same base model and tool access on the same dataset; differences are assessed via the presence of auditable artifacts and executable outputs.

| Audit target | Cursor+Amplify protocol | Protocol-free baseline |
| --- | --- | --- |
| Protocol deliverables externalized | Intake + literature + plan + integration artifacts present (research anchor; literature review; gap analysis; sufficiency and alternative explanations; argument blueprint) | No protocol deliverables directory; analysis proceeds ad hoc |
| Manuscript organization | Modular LaTeX (preamble + 5 section files) | Single-file LaTeX manuscript |
| Executable analysis pipeline | 9 scripts under scripts/; figures and paper maintained as separate top-level outputs | 13 scripts under analysis/scripts; paper maintained under analysis/manuscript |
| Figures integrated into manuscript | 4 includegraphics blocks in main text | 5 includegraphics blocks in main text |
| Analysis scope (as implemented) | Unified cross-type population-structure comparison using SNVs, INDELs, and SVs | Indel mutation-spectrum analysis with NMF decomposition |

To make the difference in artifact externalization concrete, Fig. 3 summarizes the produced directory structure in each condition.

**Figure 3.**
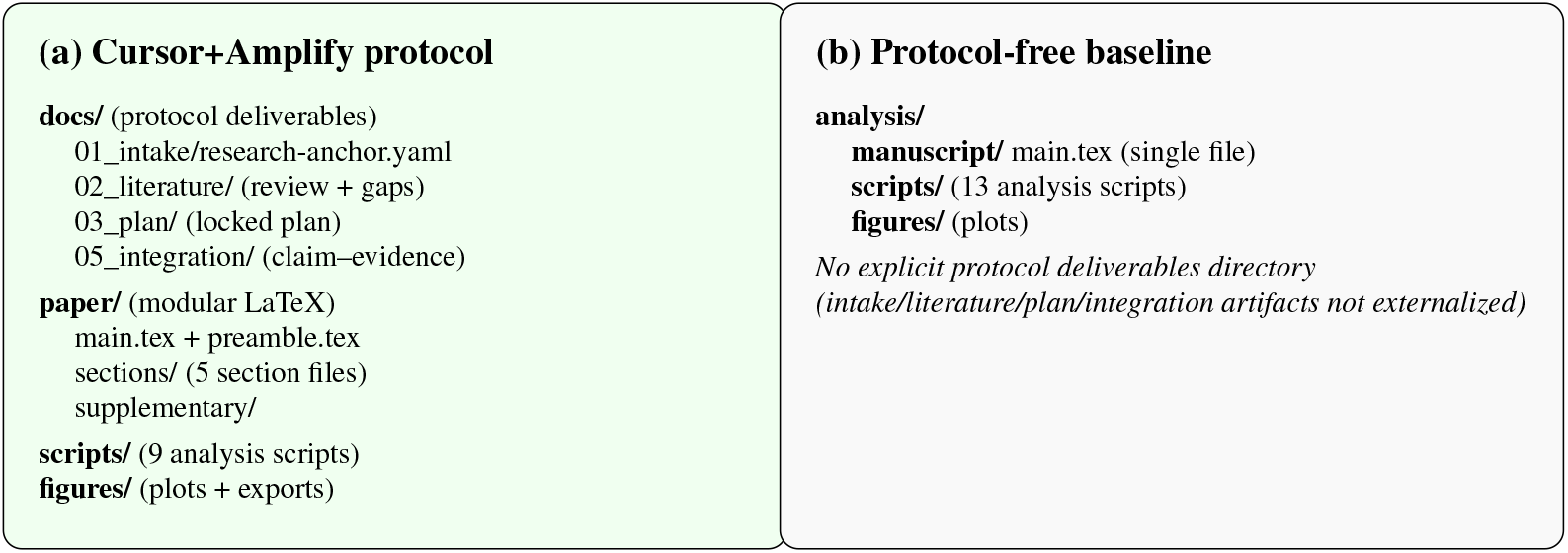
Controlled study: artifact externalization differs under a phase-gated protocol. In the Cursor+Amplify condition, protocol deliverables are externalized as auditable on-disk artifacts (docs/) alongside a modular manuscript (paper/) and an executable analysis pipeline (scripts/). In the protocol-free baseline, analysis proceeds without an explicit protocol artifact directory and outputs are organized around scripts and a single-file manuscript.

**Figure 4.**
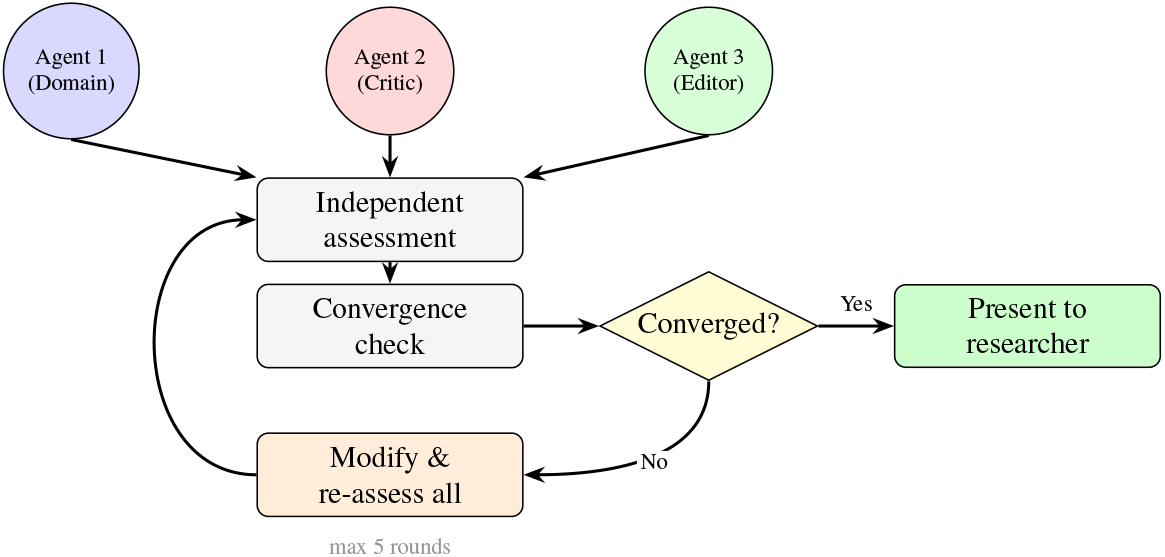
Multi-agent deliberation protocol. Three role-specialized sub-agents are dynamically instantiated based on the research question and current project state. Each sub-agent assesses the same artifact with an independent context before any convergence check, reducing anchoring and enabling adversarial critique. Deliberation proceeds through structured rounds: convergence requires unanimous PASS; otherwise the artifact is modified and all sub-agents re-assess the full artifact. Unresolved disagreements are escalated to the human researcher.

**Figure 5.**
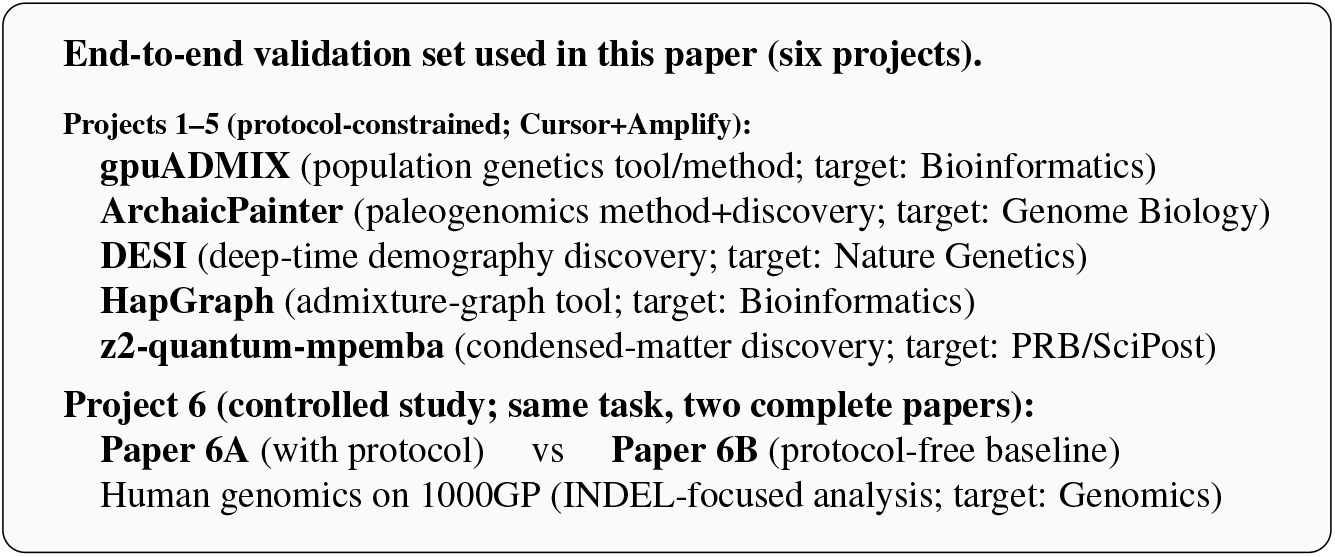
Cross-domain validation set and controlled study. Five projects were executed under the full protocol constraints (Cursor+Amplify) across population genetics, paleogenomics, human evolutionary genetics, computational population genetics, and condensed-matter physics. A sixth project is a controlled study on the same 1000 Genomes dataset and task, producing two complete papers under matched conditions with (Paper 6A) and without (Paper 6B) the protocol.

### Mechanism: how constraints alter behaviour in practice

To illustrate *how* constraints change agent behaviour—not merely that outcomes differ—we trace three representative episodes from the validation projects.

#### Vignette 1: Verification constraint catches stale evidence (gpuADMIX)

During Phase 4, the agent benchmarked gpuADMIX against fastmixture at *K* = 2–10 and initially concluded that gpuADMIX was inferior at *K* ≥ 8. The agent flagged this honestly (anti-cherry-pick constraint: “this needs honest disclosure”). When the human researcher later questioned the claim, the verification constraint required fresh re-measurement rather than reliance on stored results. Re-running revealed that the *K* ≥ 8 numbers originated from an earlier, buggy code version: the true best-of-five gpuADMIX log-likelihood matched or exceeded fastmixture at every *K*. Without the verification constraint, the stale results would have propagated into the manuscript unchallenged. Additionally, multi-agent review of the paper draft caught a completely fabricated reference (a non-existent paper attributed to a real author) and a formula discrepancy between the code (FISTA momentum) and the text (simple Nesterov)—errors that would have survived conventional self-review.

#### Vignette 2: Governance forces project redesign; integrity forces claim retraction (DESI)

At the G2 gate, the human researcher challenged the proposed analysis plan: “Is there actually a new method here, or are you just running existing tools?” The agent honestly acknowledged that the plan was “an analysis pipeline plus new application, not a new method,” triggering a complete project redesign. Later, during Phase 4, the agent’s initial finding—a two-component TMRCA distribution suggesting ancient structure— was questioned by the human. The agent re-analysed the evidence, explicitly stated “my earlier interpretation was wrong,” and downgraded the claim: “Is this enough for *Nature Genetics*? No—not from this angle.” Rather than defending the original interpretation, the integrity discipline required the agent to pursue additional simulation experiments and calibration checks. In the final write-up, a technical bias check with random labels (Model A) yields an AFR−EAS difference of 0.5 *±* 0.3 ka, while a realistic Out-of-Africa model with panmictic deep ancestry (Model OoA) predicts an AFR−EAS difference of 463.9 ka. The empirical window-level AFR−EAS difference (370.7 ka) is then assessed in the context of model-based stress tests, including a FitCoal-parameter scan that compares window-level summary statistics (e.g., 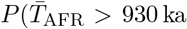) and 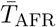) from 600 simulated windows against 27,507 real-data windows, and therefore explicitly scopes conclusions to the tested model class and summary-statistic estimator. The governance layer’s failure-management function—forcing structured reassessment after repeated setbacks—was essential to reaching this outcome rather than abandoning the project or advancing unsupported claims.

#### Vignette 3: Multi-agent deliberation catches code–text mismatches (HapGraph)

During Phase 6 (paper writing), three AI review agents independently audited the HapGraph Methods section against the actual codebase. The adversarial reviewer flagged multiple code–text inconsistencies that would survive conventional self-review (the formulas were plausible and internally consistent, and compilation/unit tests would not catch them). One representative example was the inter-population IBD timing model: the corrected implementation and write-up require 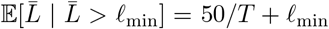, reflecting a breakpoint rate of 1*/*(2*T* ) per Morgan for two modern lineages. The claim-evidence alignment constraint required the writing agent to reconcile every quantitative assertion with the implementation before the section could pass the quality gate, and the multi-round deliberation protocol ensured discrepancies were resolved before the text was finalized. Additionally, the same review cycle identified fabricated bibliography metadata (correct author names paired with wrong titles, volumes, and DOIs) that would have been undetectable without systematic cross-referencing.

### Relationship to existing approaches

The methodology formalization addresses a dimension distinct from the capabilities that existing AI research systems optimize. To clarify this distinction, we categorize recent systems by what they encode and what they omit (Table 4).

**Table 4.** Methodology formalization addresses a dimension distinct from domain capability. Each column represents a category of existing systems; rows indicate whether the category provides each capability. Entries reflect capabilities as reported in the cited system descriptions. Legend: ✓=core feature; Partial=limited or not systematic; —=not reported/primary goal.

|  | This work | Autonomous scientists | AI co-scientist | Domain agents | Deep re-search |
| --- | --- | --- | --- | --- | --- |
| Persistent integrity | ✓ | — | — | — | — |
| Eval. immutability | ✓ | — | — | — | — |
| Complete reporting | ✓ | — | — | — | — |
| Human approval gates | ✓ | Partial | Partial | Partial | Partial |
| Direction ideation | ✓ | ✓ | ✓ | Partial | Partial |
| Structured deliberation | ✓ | Partial | ✓ | — | — |
| Autonomous execution | ✓* | ✓ | ✓ | ✓ | ✓ |
| Tool execution | ✓† | Partial | Partial | ✓ | Partial |
| Domain agnostic | ✓ | Partial | ✓ | — | ✓ |

\* Amplify is designed so that phase transitions are subject to explicit human review/approval at validation gates. When desired, a simple prompt setting can enable a “fast mode” in which the agent proceeds by default using the protocol’s recommended decisions, minimizing human intervention while retaining the same phase structure and constraints.
† Domain tool access is provided by the execution environment (e.g., Cursor-style IDEs that support web search/browsing and a bash shell), allowing agents to install and invoke many domain tools on demand.

#### Domain-specific task automation

Systems such as AutoBA [20], BioMaster [21], and Biomni [14] encode knowledge of *which tools to run and in what order* for specific scientific domains. AutoBA and BioMaster automate bioinformatics pipelines (RNA-seq, ChIP-seq, spatial transcriptomics) by generating and executing analysis plans from minimal user input; Biomni extends this to 25 biomedical subfields by orchestrating 105 software tools, 150 biological protocols, and 59 databases. Similarly, ChemCrow [36] and SciAgents [37] achieve deep integration with chemistry and materials science knowledge, respectively. These systems excel at executing established workflows—a genuine contribution—but do not encode higher-level methodological reasoning: they do not question whether a planned analysis adequately addresses the research question, require evaluation protocols to be locked before execution, or enforce complete reporting of unfavourable outcomes.

#### End-to-end autonomous AI scientists

A rapidly growing category of systems aims to automate the full research cycle. The AI Scientist [15] and its successor AI Scientist v2 [16] generate ideas, write code, run experiments, and produce complete ML papers—the latter achieving the first AI-generated peer-reviewed publication at an ICLR 2025 workshop. Kosmos [17] (FutureHouse) performs extended 12-hour discovery sessions with a structured world model that maintains coherence across ∼200 agent rollouts, producing reports with 79.4% statement accuracy across metabolomics, materials science, neuroscience, and statistical genetics. Robin [18] (FutureHouse) achieved the first fully automated biological discovery—identifying a drug candidate for age-related macular degeneration through iterative hypothesis generation and experimental design. Google’s AI-powered empirical software system [38] uses tree search to optimize scientific code across six benchmarks, achieving expert-level performance in genomics, epidemiology, and neuroscience. Agentomics [39] autonomously develops state-of-the-art ML models for biomedical datasets, outperforming human expert solutions on 11 of 20 benchmarks. These systems demonstrate that LLMs can execute substantive research tasks autonomously—a capability our approach leverages but does not aim to replicate. What they do not provide is persistent methodological discipline: none enforces evaluation immutability, requires complete reporting of negative results, mandates alternative-hypothesis exclusion before causal claims, or provides governance mechanisms to halt failing approaches.

#### Structured hypothesis generation

Google’s AI co-scientist [19] is the closest existing system to our approach in terms of structured process. Built on Gemini 2.0, it employs specialized agents (Generation, Reflection, Ranking, Evolution, Proximity, Meta-review) in a “generate, debate, and evolve” cycle that iteratively improves research hypotheses through tournament evolution, with demonstrated validation in drug repurposing, novel target discovery, and bacterial evolution. The system shares our philosophy that multiagent deliberation and human-AI collaboration improve research quality. However, the two approaches differ fundamentally in *scope* and *what is formalized*. AI co-scientist formalizes the hypothesis generation and refinement stage—producing ranked candidate hypotheses for scientists to validate experimentally. Our formalization covers the *complete research lifecycle*: from question formulation through experiment execution, results integration, and manuscript preparation, with persistent integrity constraints (evaluation immutability, complete reporting, claim-evidence alignment) active throughout. AI co-scientist does not encode constraints on how experiments should be conducted once a hypothesis is selected, does not require complete reporting of negative outcomes, and does not provide governance mechanisms to force pivoting when approaches fail. The relationship is again complementary: AI co-scientist’s sophisticated hypothesis generation could feed into our methodology’s Phase 1 (direction exploration), after which our integrity and governance layers would ensure the selected hypothesis is pursued with sustained methodological discipline.

#### Knowledge synthesis and deep research

Products such as OpenAI’s Deep Research [22] and Google’s Gemini Deep Research automate literature search and report generation at scale. These are valuable for the direction exploration phase of the research cycle but cover only one component of the full methodology— they do not design experiments, lock evaluation protocols, or verify that claims rest on specific evidence.

#### Complementarity

Our formalization is complementary to all these approaches rather than competitive with them. Domain-specific systems encode *what tools to use*; autonomous scientists encode *the capability to execute*; deep research products encode *how to synthesize existing knowledge*; our work encodes *the methodological discipline that makes research outputs reliable*. In principle, the three constraint layers (procedural workflow, integrity discipline, governance) could be applied atop any of these systems—providing the methodological scaffolding that currently separates impressive capability demonstrations from publishable, reproducible science.

## Discussion

The most surprising aspect of these results is not that AI agents can be constrained to follow rules— instruction-following is a known capability of modern LLMs—but that a relatively compact formalization of research methodology suffices to transform the behaviour of general-purpose AI from ad hoc text generation into structured research practice across diverse domains. Two findings stand out. First, the complete research process—how to formulate questions, validate directions, design methods, iterate through failures, recognize when to pivot, integrate results into evidence-grounded narratives—*can* be articulated as a coherent system of interlocking constraints. Second, AI agents, when given this system, can be steered to follow it: in our validation projects, the same models that without constraints can produce complete manuscripts while leaving planning and verification implicit instead conduct multi-phase, methodologically disciplined research with auditable intermediate artifacts. This suggests that the gap between “AI that can write about science” and “AI that can do science responsibly” is not primarily a gap in model capability but a gap in *methodology transfer*: the procedural knowledge that experienced researchers possess was simply never given to the AI.

This finding resonates with a broader insight from constitutional AI [40], which demonstrated that explicit principles steer model behaviour more reliably than implicit training signals. We extend this from AI safety to scientific integrity, and from individual interactions to multi-phase projects with persistent state and accumulated constraints. The three-layer decomposition we propose (workflow, discipline, governance) mirrors how methodology functions in human research groups—the PI’s plan, the lab’s integrity culture, the oversight of committees [7]—and the fact that each layer catches distinct failure modes (see Results Section) suggests that these are not arbitrary categories but reflect genuine functional divisions in how reliable knowledge is produced. Methodological constraints do not replace domain expertise, stronger models, or improved tool integrations. Rather, they make a research agent’s methodological *process* explicit and auditable— externalizing phase progression, turning integrity checks into enforceable obligations with concrete artifacts, and forcing stop/pivot decisions to be written down and justified. A practical implication is that the formalization can be applied directly atop general-purpose LLMs. Because the protocol is model-agnostic, it can benefit automatically from improvements in underlying foundation models (knowledge, reasoning, tool use) while keeping the methodology layer fixed and auditable. In cost terms, this shifts effort from large-scale model training to an executable, reusable protocol that can be shared and iterated as open-source software.

The validation projects also reveal a domain-dependent shift in where effort is required. In data-analysis projects, the dominant cost is often computational execution and debugging of pipelines. In contrast, in the z2-quantum-mpemba condensed-matter project, computational runtime was modest but the agent devoted substantially more effort to analytical reasoning and proof-like verification against exact constraints (closed-form initial conditions, finite-size scaling, and benchmarkable dynamical signatures)—a pattern that aligns with how human theoretical physics often emphasizes reasoning over data processing.

Looking forward, one can view protocol-layer enforcement as a stepping stone rather than an endpoint. In principle, the same constraints could be used as training signals (e.g., via reinforcement learning or preference optimization) so that models internalize parts of the methodology rather than relying on external enforcement; we did not pursue such training due to resource constraints and because our goal here is an auditable, model-agnostic protocol. We also observed meaningful performance differences across foundation models on different phases and tasks. A protocol controller makes it straightforward to exploit this heterogeneity by routing different phases or deliberation sub-agents to models that are better suited for the corresponding cognitive demands (e.g., long-horizon planning, code execution, or mathematical reasoning).

Several risks require candid acknowledgment. First, formalized methodology enforces *process* but cannot assess *substance*: a project can satisfy every constraint and still be scientifically trivial. Moreover, performance remains bounded by the underlying model’s knowledge and reasoning ability; the protocol can force checks and backtracks, but it cannot invent missing domain insight. Human judgment at gates addresses this partially, but users should understand that methodology-constrained AI produces rigorously *structured* research, not independently *validated* science. Second, the current formalization encodes the hypothetico-deductive methodology dominant in quantitative sciences; qualitative research, abductive reasoning, and indigenous knowledge systems follow different methodological traditions not captured here. Extending the formalization to accommodate methodological pluralism is essential future work. Third, constraints are enforced through structured instructions rather than formal verification; LLMs are probabilistic systems, and subtle violations (such as biased interpretation of ambiguous results) may evade detection.

Overall, the results support a practical path to scaling methodological rigour: treat methodology as an auditable, reusable protocol layer that can be applied to general-purpose models today, improved as foundation models advance, and eventually partially internalized through training or deployed via phase- and role-specific model routing.

## Methods

### Approach to methodology formalization

To formalize scientific methodology, we analysed how experienced researchers maintain rigour across the research lifecycle and distilled recurring patterns into explicit, executable principles. Sources included published methodology guides, research integrity literature [29, 30, 41], reporting standards (CONSORT [3], PRISMA [4]), pre-registration frameworks [1, 2], and the accumulated conventions of peer review practice. We sought principles that were (a) domain-agnostic—applicable to computational research regardless of field, (b) enforceable—expressible as verifiable constraints rather than aspirational guidelines, and (c) decomposable—assignable to a specific functional layer without entangling with other concerns .

This process yielded 24 distinct principles organized into the three-layer structure described in Results. Six overarching constitutional rules govern the entire formalization: (1) research type determines all procedural paths; (2) the target publication standards are established at inception and influence all downstream decisions; (3) the AI agent adopts a domain-specific expert identity rather than operating as a generic assistant; (4) methodological justification precedes implementation; (5) evaluation criteria, once locked, cannot be modified without explicit human authorization; and (6) no claim may be made without fresh computational verification.

### Implementation as a phase-gated research protocol

To test whether formalized methodology can be transferred to AI agents as an operational protocol (rather than a single prompt), we implemented each principle as a structured protocol card specifying trigger conditions, required inputs, procedural steps, deliverables, exit criteria, and integration points. Collectively, these cards define a stateful, phase-gated process: phases advance only when entry/exit criteria are met; invariants such as metric immutability persist across phases; and violations trigger halts and explicit return paths for correction. Constraints are categorized as *rigid* (followed without adaptation; e.g., metric immutability, complete reporting) or *flexible* (principles adapted to context; e.g., direction exploration, method design). Rigid constraints include explicit verification steps—for example, the metric immutability constraint requires comparison of any proposed evaluation change against the locked protocol, with mandatory halt on discrepancy.

The protocol operates atop general-purpose LLMs (Claude, GPT-4, Gemini, and others) without fine-tuning or model modification. At runtime, a lightweight protocol controller ensures that the agent is operating under the correct phase and discipline rules by loading the relevant protocol cards into the agent’s working context as structured system-level instructions. Crucially, persistent state is maintained outside the model via logged artifacts and explicit interfaces between phases, so that enforcement is not merely “remembering a prompt” but checking protocol invariants against concrete outputs (e.g., the locked evaluation protocol, recorded seeds, and claim–evidence tables).

#### Controller implementation in the open-source release

Amplify is implemented as a plugin-style skills library plus a small set of runtime hooks and always-on rules that make protocol loading explicit and reproducible. The skills library is declared in a plugin manifest (amplify/.cursor-plugin/plugin.json) and stored as protocol cards under amplify/skills/*/SKILL.md. A SessionStart hook (amplify/hooks/hooks.json calling amplify/hooks/session-start.sh) injects the full content of the using-amplify skill into the system context at the beginning of a chat session, making global workflow constraints (e.g., one-phase-per-turn, gate enforcement, and mandatory skill invocation when triggers apply) available before any phase begins. For Cursor-style rule systems, an equivalent always-on bootstrap rule file (amplify/install/amplify-bootstrap.mdc) provides the same controller constraints when hook-based plugins are not used.

Phase transitions are enforced procedurally: the protocol requires the agent to stop after completing a phase or reaching a gate, present deliverables and a gate checklist, and wait for explicit user approval before proceeding. Discipline and governance cards (e.g., metric-lock, results-verification-protocol) remain active once triggered and explicitly forbid common integrity violations (e.g., post hoc metric changes without authorization; claiming results without fresh verification evidence). Persistent state and locked contracts are externalized as on-disk artifacts—not hidden model context—using templates such as docs/01_intake/research-anchor.yaml and docs/03_plan/evaluation-protocol.yaml (locked after G2), along with experiment logs and claim–evidence alignment tables. This separation between (i) runtime context (protocol text) and (ii) verifiable artifacts (contracts and evidence) is what prevents the system from degenerating into prompt-only compliance and makes the workflow auditable.

### Procedural workflow specification

The procedural workflow encodes seven phase-gated phases with mandatory temporal ordering and explicit return paths for backtracking:

**Phase 0 (Domain Anchoring)** identifies the research domain, subdomain, research type (Method, Discovery, Tool, or Hybrid), and available resources. Ambiguous inputs trigger clarification rather than assumption.

**Phase 1 (Direction Exploration)** executes autonomous literature search (15–30 papers with full-text retrieval), applies six structured ideation strategies (contradiction mining, assumption challenging, cross-domain transfer, limitation-to-opportunity conversion, counterfactual reasoning, trend extrapolation), generates candidate research directions, and refines them through multi-agent brainstorming.

**Phase 2 (Problem Validation)** subjects the selected direction to adversarial questioning through a three-agent deliberation panel, applying a novelty litmus test and feasibility verification. This phase is mandatory for all research types.

**Phase 3 (Method/Framework Design)** branches by type. Method projects lock an evaluation protocol (metrics, datasets, seeds, statistical tests, baselines). Discovery projects design an analysis storyboard with main and supporting lines, sufficiency criteria, and pre-identified alternative explanations.

**Phase 4 (Experiment Execution)** operates in two stages: exploratory validation followed by full execution. Method projects require minimum three diagnose–hypothesize–fix–measure iteration cycles. All experiments use isolated computational environments, fixed random seeds, and logged conditions.

**Phase 5 (Results Integration)** compiles outputs, constructs a claim-evidence alignment table, and deploys a three-agent panel for narrative design. Fatal vulnerabilities block progression.

**Phase 6 (Paper Writing)** produces modular manuscripts with per-section multi-agent polishing, autonomous reference verification, and full-paper review.

### Integrity discipline specification

Seven persistent constraints activate at defined trigger points and remain active until project completion:

*Metric immutability*: after the evaluation protocol is locked, any modification requires justification, proof of necessity, and human authorization.

*Complete reporting*: all experimental seeds reported with mean and standard deviation; all negative results recorded; baselines given equal resources; all specified datasets tested.

*Claim-evidence alignment*: mapping table linking every paper assertion to supporting evidence; unmapped claims flagged for removal.

*Alternative-hypothesis exclusion*: systematic confounder assessment before any causal claim. *Reproducibility*: hypothesize–baseline–experiment–verify–interpret cycle with environment logging. *Verification*: fresh computational evidence required before any status claim.

*Figure standards*: publication-specific visual quality enforced through pre-defined style profiles.

### Multi-agent deliberation protocol

Deliberation sessions follow a standardized convergence protocol. A shared scoring rubric is established before deliberation. Agents assess artifacts independently, revealing assessments simultaneously to prevent anchoring. After each round, the system checks convergence (all agents PASS, no fatal issues). If not converged, modifications are applied and all agents re-assess the complete artifact (not just changes). Deliberation terminates upon convergence, upon reaching the round limit (maximum five), or upon non-convergence (disagreements escalated to the human researcher).

### Validation design

We evaluated the formalized methodology through two complementary approaches. First, we applied the full constraint set to five protocol-constrained research projects spanning diverse scientific domains and research types. For each project, we documented: (1) domain and research type classification; (2) all phases and gates completed; (3) integrity constraint compliance (metric stability, seed reporting, negative result recording); and (4) output artifacts suitable for audit (paper structure, figure inclusion, reference list, and claim–evidence mapping).

Second, to assess whether the constraints themselves (rather than LLM capability alone) account for the observed rigour, we conducted a direct controlled comparison matched on model, task, tool access, and budget, with the sole difference being activation of the methodology constraints (protocol-free baseline versus Cursor+Amplify; Section “Controlled study”). Concretely, in the controlled-study project (Project 6) the agent produced two complete manuscripts under matched conditions with and without the protocol (Supplementary Papers 6A and 6B). We evaluated outputs by auditing generated artifacts rather than subjective scoring (e.g., manuscript modularity, executable scripts, and the presence of explicit protocol deliverables).

#### Controlled study (Project 6): protocol-free baseline versus Cursor+Amplify protocol

To isolate the contribution of the protocol controller from underlying model capability and tool access, we conducted a head-to-head controlled study on an identical task using the 1000 Genomes Project 2022 high-coverage dataset (3,202 individuals, 26 populations). Both conditions used the same IDE environment (Cursor), the same base model (Claude Opus 4.6), and the same local toolchain (bcftools, PLINK2, Python scientific stack), under the same constraint of no deep learning model training. The protocol-free baseline condition (Cursor with no Amplify protocol enabled) was explicitly forbidden from using any Amplify materials and was prompted to autonomously produce original, publishable scientific findings and a complete manuscript. The protocol condition (Cursor+Amplify) enabled the phase-gated methodology protocol and emphasized autonomous execution (running analyses) while producing the protocol deliverables as on-disk artifacts. Human input was restricted to operational continuation and final compilation requests; no methodological guidance, scientific direction, or writing edits were provided. Outcomes were evaluated by auditing generated artifacts rather than subjective scoring: manuscript organization (single-file versus modular sections), analysis scripts and directory structure, and the presence of protocol deliverables (intake anchor, literature review artifacts, planning documents, and integration blueprint).

## Code and data availability

The Amplify protocol specification (24 skills, templates, and installation/bootstrapping materials) is released as open-source software under the MIT licence at https://github.com/EvoClaw/amplify. All code used for the analyses in this paper—including project-specific scripts, figures, and manuscripts for Projects 1–6 (including the controlled-study pair, Papers 6A/6B)—is available in the public repositories under the EvoClaw organization: https://github.com/orgs/EvoClaw/repositories.

This work introduces no proprietary datasets. Empirical genomics analyses use publicly available 1000 Genomes Project data; simulation-based projects generate data programmatically and include the corresponding code for reproduction in the repositories above.

## Author Contributions

Y.Z. conceived the study, designed the protocol and controlled study, implemented the project, performed all experiments and analyses, and wrote the manuscript. J.Z. provided suggestions on biological data analysis and workflow design, and reviewed the manuscript. All authors approved the final version.

## Acknowledgements

Amplify and the accompanying project artifacts are released under the MIT licence to encourage reuse, reproduction, and extension by the research community.

## Footnotes

$ Source code: https://github.com/EvoClaw/amplify.

