## Supplementary File for "Formalized scientific methodology enables rigorous AI-conducted research across domains"

### Supplementary Information for:
### Formalized scientific methodology enables rigorous AI-conducted research across domains

Yanlin Zhang[1,*] Jing ZHAO[2]

[1] Data Science and Analytics Thrust, Hong Kong University of Science and Technology (Guangzhou), Guangzhou, 511453, China
[2] Shenzhen DataSpeak Intelligent Technology Co., Shenzhen 518000, Guangdong, China

### Supplementary Information

#### Supplementary Note 1: Complete Skill Inventory

Table S1 provides the complete inventory of 24 skills organized by architectural layer, with their primary activation trigger and the main failure mode(s) each prevents.

Table S1: **Complete inventory of 24 methodology skills.** Each skill is a reusable protocol/constraint module. "Trigger" summarizes the primary activation condition; in practice, multiple skills may remain active concurrently.

| Layer | Skill | Trigger | Purpose / failure mode prevented |
|---|---|---|---|
| *Procedural workflow (Phase 0–6; one phase per turn).* | | | |
| Procedural workflow | using-amplify | Session start | One-phase-per-turn + gate stops; prevents phase skipping and one-shot "research". |
| Procedural workflow | domain-anchoring | Phase 0 | Anchors domain/type/resources/standard; prevents unscoped starts. |
| Procedural workflow | research-direction-exploration | Phase 1 | Direction generation + ranking through G1; prevents premature commitment. |
| Procedural workflow | problem-validation | Phase 2 | Adversarial validation + feasibility ruling; prevents weak/non-publishable questions. |
| Procedural workflow | method-framework-design | Phase 3 | Full plan before coding; prevents uncontrolled trial-and-error. |
| Procedural workflow | evaluation-protocol-design | Phase 3 (Type M/H) | Locks metrics/datasets/baselines/seeds/tests; prevents goal-post moving. |
| Procedural workflow | analysis-storyboard-design | Phase 3 (Type D/H) | Locks story line + sufficiency criteria + confound checks; prevents narrative-first discovery. |
| Procedural workflow | experiment-execution | Phase 4 (after G2/G3) | Baseline-first execution + iteration; prevents skipping baselines and untracked variants. |
| Procedural workflow | results-integration | Phase 5 | Evidence-grounded integration blueprint; prevents writing before integration. |
| Procedural workflow | paper-writing | Phase 6 (after G4) | Modular drafting + iterative refinement; prevents single-pass manuscripts. |
| Procedural workflow | using-git-worktrees | As needed | Safe isolation for experiments; prevents workspace contamination. |
| Procedural workflow | dispatching-parallel-agents | As needed | Parallel independent work; prevents serial bottlenecks and state bleed. |
| Procedural workflow | multi-round-deliberation | When deliberating | Convergence loop (assess → modify → re-check); prevents one-round feedback. |
| *Integrity discipline (persistent methodological constraints).* | | | |
| Integrity discipline | results-verification-protocol | Always | Evidence required before claims; prevents unverified reporting. |
| Integrity discipline | reproducibility-driven-research | Any run | Reproducible pipelines/logs/configs; prevents hidden manual steps. |
| Integrity discipline | metric-lock | After G2 | Locked metrics/datasets/splits; prevents silent evaluation changes. |
| Integrity discipline | anti-cherry-pick | Phase 4 onward | All seeds/failures reported; prevents selective reporting. |
| Integrity discipline | alternative-hypothesis-check | Type D/H | Confound exclusion before mechanisms; prevents spurious causal stories. |
| Integrity discipline | claim-evidence-alignment | Drafting | Claims must cite evidence; prevents narrative drift. |
| Integrity discipline | figure-quality-standards | Figures/tables | Venue-quality visuals; prevents illegible or misleading figures. |
| *Governance (strategic controls; triggered by risk conditions).* | | | |
| Governance | novelty-classifier | Phase 1/3 | Checks novelty vs target standard; prevents low-novelty dead ends. |
| Governance | scope-control | Any scope creep | Forces explicit scope decisions; prevents runaway projects. |
| Governance | pivot-or-kill | Repeated failures | Forces pivot/downgrade/stop; prevents endless iteration. |
| Governance | venue-alignment | Gate/periodic | Aligns evidence/scale/writing to target standard; prevents under-powered validation. |

#### Supplementary Note 2: Gate Checklists

Below we provide detailed checklists for the four gates (G1–G4). Gates are auditable checkpoints: the deliverables are explicit artifacts, and the pass/fail rationale can be reviewed independently of the model's internal chain-of-thought.

**G1: Direction & feasibility (end of Phase 1).** Required deliverables and conditions:

- Updated research anchor (domain, research type, resources, and target publication standard).
- A ranked list of candidate directions with a short novelty/feasibility rationale for each.
- A selected direction with a clear research question statement and success criteria at the level of a paper contribution (not just "run analysis").
- Basic feasibility: data availability, compute constraints, and a credible path to evidence at the chosen publication standard.

**G2: Plan freeze (end of Phase 3).** Required deliverables and conditions:

- Frozen method/analysis plan with explicit assumptions and known failure modes to monitor.
- Locked evaluation protocol (metrics, datasets/splits, baselines, random seeds, statistical tests, and stopping criteria) for Type M/H projects; locked sufficiency criteria and alternative explanations to test for Type D/H projects.
- A concrete execution plan (scripts/pipeline outline, expected runtime, and outputs needed for figures/tables).
- Scope is stable (any planned extensions listed as post-acceptance or future work).

**G3: Execution readiness (before Phase 4).** Required deliverables and conditions:

- Environment readiness: dependencies installable; data paths validated; minimal pipeline runs end-to-end on a small subset.
- Baseline readiness: primary baselines identified and a reproduction plan exists (or a justified exception is recorded).
- Logging and reproducibility plan in place (seeds, configs, and outputs captured).

**G4: Write-ready (end of Phase 5).** Required deliverables and conditions:

- Verified results summary (what was tested, what worked/failed, and what evidence supports each conclusion).
- Integration blueprint mapping: claims $\rightarrow$ evidence $\rightarrow$ figures/tables (and explicitly listing unresolved evidence gaps).
- A stabilized figure/table plan suitable for the target publication standard (readability, accessibility, and consistency).

#### Supplementary Note 3: Multi-Agent Deliberation Protocol Assets

Rather than relying on ad hoc one-round "panel feedback", Amplify standardizes multi-agent deliberation as an iterative convergence loop (assess → modify → re-assess) with explicit verdicts (PASS / CONDITIONAL / FAIL) and a fixed maximum number of rounds. The following deliberation assets are provided in the release:

- Shared rubric and convergence rule (used across phases) defining what counts as "resolved" versus "unresolved" issues.
- Role-conditioned prompts for panels used in (i) direction exploration, (ii) problem validation, (iii) results integration story design, and (iv) per-section paper polishing.
- A non-convergence fallback: if consensus is not reached within the round limit, unresolved disagreements are surfaced as explicit trade-offs for the user to decide.

#### Supplementary Note 4: Validation Papers

We report six end-to-end *projects*. Projects 1–5 correspond to the five protocol-constrained projects discussed in the main Results section. Project 6 is a controlled study on the same dataset and task, in which the same AI agent produced two complete manuscripts under two conditions: with the full Cursor+Amplify protocol enabled (Supplementary Paper 6A) and without the protocol (protocol-free baseline; Supplementary Paper 6B). This makes the controlled comparison auditable at the level of complete papers rather than partial artifacts.

Table S2: Summary of six end-to-end projects and the resulting manuscripts. The controlled study (Project 6) yields two complete papers (6A/6B) under matched conditions with and without the protocol.

| **Paper** | **Domain** | **Research Type** | **Target standard (example)** | **Phases Completed** |
|---|---|---|---|---|
| 1 | Population genetics (GPU ancestry estimation) | Hybrid (M+T) | Bioinformatics | 0–6 |
| 2 | Paleogenomics (archaic introgression detection) | Hybrid (M+D) | Genome Biology | 0–6 |
| 3 | Human evolutionary genetics (deep-time demography) | Discovery | Nature Genetics | 0–6 |
| 4 | Computational population genetics (admixture graph inference) | Tool (C) | Bioinformatics | 0–6 |
| 5 | Condensed matter physics ($Z_2$ quantum Mpemba effect in TFIM/XY) | Discovery | PRB / SciPost Physics | 0–6 |
| 6A | Human genomics (1000GP INDEL-focused analysis; controlled study, *with* protocol) | Discovery | Genomics | 0–6 |
| 6B | Human genomics (1000GP INDEL-focused analysis; controlled study, protocol-free baseline) | Discovery | Genomics | 0–6 (protocol-free) |

#### Supplementary Note 5: Interaction-log-derived proxies for analytical difficulty

The validation set includes both data-analysis-heavy projects (e.g., population genetics and bioinformatics) and theory-heavy projects (e.g., condensed-matter physics). While all projects follow the same phase-gated methodology, the *dominant difficulty* can shift by domain: some domains are bottlenecked by computational execution and pipeline correctness, whereas others are bottlenecked by analytical reasoning and repeated verification against exact constraints.

To make this observation auditable without subjective scoring, Supplementary Table S3 reports proxies computed directly from the Cursor-exported interaction logs provided with each project. We segment each log into alternating "User" and assistant turns (message blocks) and report (i) the number of assistant turns (a proxy for interaction length) and (ii) the number of assistant turns that contain a *major backtrack* decision or *major self-critique* judgment, as determined by manual semantic coding. Here "major" means the agent explicitly invalidates a previously stated method/result/claim (e.g., discovers a serious bug, retracts a core interpretation, or identifies an evidence gap that requires rerunning or redesigning analysis), rather than making purely presentational edits.

Table S3: **Interaction-log-derived proxies for domain-dependent analytical difficulty across the five completed projects.** "Backtrack turns" and "self-critique turns" count assistant turns that contain a major backtrack or major self-critique, based on manual semantic coding of the full interaction logs (not keyword matching). "Backtrack rate" is the fraction of assistant turns marked as major backtracks. These are proxies (not ground-truth difficulty) but allow consistent, auditable comparison across domains.

| Project | Domain | Assistant turns | Backtrack turns | Self-critique turns | Backtrack rate |
|---|---|---|---|---|---|
| gpuADMIX | Population genetics (method/tool) | 27 | 6 | 6 | 0.22 |
| ArchaicPainter | Paleogenomics (method+discovery) | 28 | 8 | 8 | 0.29 |
| DESI | Human evolutionary genetics (discovery) | 53 | 11 | 11 | 0.21 |
| HapGraph | Computational population genetics (tool) | 44 | 7 | 7 | 0.16 |
| z2-quantum-mpemba | Condensed-matter physics (discovery) | 23 | 10 | 10 | 0.43 |

#### Supplementary Papers

This Supplementary Information document includes the full PDFs of the seven manuscripts produced across the six end-to-end projects. Each paper is embedded in full, as generated, to make the controlled comparison and cross-domain validation directly auditable.

*Bioinformatics*, 2026, pp. 1–7
DOI HERE
Advance Access Publication Date: Day Month Year
**Original Article**

### gpuADMIX: GPU-accelerated ancestry estimation with Nesterov-augmented mini-batch EM

Amplify[1]

[1], ,

#### Abstract

**Motivation:** Model-based admixture estimation methods such as ADMIXTURE and fastmixture are widely used to infer individual ancestry proportions from genome-wide genotype data, but their CPU-bound runtime makes $K$ sweeps with multiple random seeds impractical at biobank scale. Existing GPU-accelerated alternatives sacrifice the exact binomial likelihood model for speed, reducing the accuracy and interpretability of ancestry estimates.
**Results:** We present gpuADMIX, which reformulates both the E-step and M-step of the admixture expectation-maximisation algorithm as GPU-native dense matrix multiplications, preserving the exact binomial likelihood *model* while achieving $41\times$ and $213\times$ speedups over fastmixture and ADMIXTURE on the 1000 Genomes Phase 3 dataset ($N = 3{,}202$; $K = 5$). Three algorithmic innovations amplify these gains: Nesterov momentum reduces EM iterations by $2.3\times$ and improves converged log-likelihood by $7{,}865$ units over plain EM; stochastic mini-batch EM improves solution quality while reducing peak GPU memory; and streaming randomised SVD provides efficient spectral initialisation for large datasets. The best of five parallel gpuADMIX seeds matches or exceeds fastmixture at every tested $K \in \{2, \ldots, 10\}$, while completing five seeds costs less wall time than a single fastmixture run, making multi-seed inference the practical default workflow. We also provide CLUMPAK-lite (CLUMPAK-lite) for label-consistent ancestry-proportion visualisation across $K$ values, and a multi-GPU dispatcher that completes $K = 2$–$10$ scans in $\approx 130\,$s on eight GPUs.
**Availability and implementation:** gpuADMIX is implemented in Python using PyTorch and is freely available at `https://github.com/EvoClaw/gpuADMIX` under the MIT licence.

#### 1. Introduction

Individual ancestry estimation—resolving each genome into proportions contributed by $K$ ancestral populations—is a cornerstone analysis in modern human genetics. Its applications span characterising patterns of human diversity and migration (Rosenberg et al., 2002; Novembre et al., 2008), identifying and correcting for population stratification in genome-wide association studies (Price et al., 2006), reconstructing recent demographic events such as colonial admixture and diaspora formation, and assigning continental or subcontinental ancestry in clinical and forensic genomics. The utility of these applications depends critically on obtaining accurate ancestry proportion estimates for the precise cohort under study, placing the estimation method at the centre of the analysis pipeline. The foundational methods STRUCTURE (Pritchard et al., 2000), FRAPPE (Tang et al., 2005), and ADMIXTURE (Alexander et al., 2009) formalise this as maximum likelihood estimation under a binomial admixture model: individuals' genotypes at $M$ biallelic loci are treated as independent draws from a mixture of $K$ ancestral allele-frequency distributions. ADMIXTURE reformulated the expectation-maximisation (EM) algorithm with block-coordinate updates amenable to vectorised computation, reducing runtime from days to hours on datasets available at the time. fastSTRUCTURE (Raj et al., 2014) achieved further gains through variational inference, while FASTMIXTURE (Meisner et al., 2024) recently delivered approximately $20\times$ speedup over ADMIXTURE via SQUAREM-accelerated EM (Varadhan and Roland, 2008).

Despite these advances, model-based admixture inference remains computationally prohibitive at biobank scale. Datasets such as the UK Biobank (Bycroft et al., 2018) encompass hundreds of thousands of individuals, and the EM runtime scales with both $N$ and $M$: even with FASTMIXTURE's acceleration, a $K = 2$–$10$ sweep over 200,000 SNPs requires hours per $K$ value on a many-core server. In practice, this forces analysts to run a single value of $K$ with a single random seed, forgoing two scientifically important capabilities: rigorous $K$ selection by cross-validation or information criteria, and the detection of multiple local optima—a well-documented feature of the EM landscape for mixture models (Jakobsson and Rosenberg, 2007; Dempster et al., 1977).

Graphics processing units (GPUs) offer a natural path to acceleration: their thousands of parallel arithmetic units achieve peak throughput when computation is cast as large dense matrix multiplications (DGEMM). Prior GPU-accelerated approaches have pursued speed by departing from the classical likelihood framework. Neural ADMIXTURE (Mantes et al., 2023) replaces EM with a neural-network surrogate trained by gradient descent, which is fast but yields Q matrices that are markedly less self-consistent across SNP subsets than ADMIXTURE's model-based estimates (Meisner et al., 2024). SCOPE (Chiu et al., 2022) optimises a least-squares latent-subspace objective rather than the binomial log-likelihood,

Received on XXXX; revised on XXXX; accepted on XXXX

© The Author(s) 2026. Copyright and licence statements to be updated by the publisher during production.

gaining scalability at the cost of the interpretable per-population allele-frequency matrix that practitioners rely on for biological annotation. This apparent accuracy–speed trade-off has persisted largely because the standard EM updates for admixture models had not been reformulated to map efficiently onto GPU DGEMM primitives.

Here we present GPUADMIX, a GPU-accelerated admixture estimation tool that resolves this trade-off by expressing both the E-step and M-step entirely as DGEMM operations on the full genotype matrix, preserving the exact ADMIXTURE binomial likelihood model without approximating the probabilistic framework. We augment this GPU-native EM with three complementary algorithmic innovations. First, FISTA-style Nesterov momentum (Beck and Teboulle, 2009) applied in the space of the EM iterates reduces iterations to convergence by 2.3× and empirically yields higher-quality solutions than plain EM. Second, stochastic mini-batch EM partitions the SNP axis into subsets processed sequentially each iteration, providing an implicit stochastic perturbation that improves solution quality while reducing peak GPU memory requirements; as with any stochastic EM, individual iterations optimise a subset of the data rather than the full likelihood, and convergence to a good full-data optimum is achieved through the aggregated effect of many such partial steps. Third, a streaming randomised SVD (Halko et al., 2011) initialises $Q$ and $P$ from the leading spectral structure of the genotype matrix without materialising the full centred $N \times M$ matrix, enabling initialisation on datasets that exceed available GPU memory. Together, these innovations yield 41× and 213× speedups over FASTMIXTURE and ADMIXTURE at $K = 5$ on the 1000 Genomes Project dataset, while *increasing* the converged log-likelihood by 2,892 and 3,088 units, respectively, over these baselines.

In addition to the core estimation engine, GPUADMIX ships with CLUMPAK-lite (CLUMPAK-LITE), a Python post-processor that solves the label-switching problem within a single $K$ via the Hungarian algorithm and across $K$ values via a greedy bottom-up procedure, enabling consistent ancestry-proportion bar plots across an entire $K$ sweep without external dependencies. A built-in multi-GPU dispatcher assigns independent $K$ values to separate GPU devices, completing a full $K = 2$–10 scan in approximately 130 s on eight GPUs.

Section 2 details the probabilistic model, GPU-native EM reformulation, Nesterov momentum scheme, mini-batch strategy, SVD initialisation, CLUMPAK-lite alignment procedure, and cross-validation protocol. Section 3 evaluates speed, accuracy, and $K$ selection on the 1000 Genomes Phase 3 dataset and quantifies each component's contribution through ablation. Section 4 situates GPUADMIX in the context of prior work and discusses limitations and future directions. Together, GPUADMIX and CLUMPAK-LITE make accurate, multi-seed, multi-$K$ ancestry inference a practical default workflow rather than a computational luxury.

### 2. Methods

#### 2.1. Probabilistic model

We adopt the binomial admixture model of Alexander et al. (2009). Let $\mathbf{G} \in \{0, 1, 2\}^{N \times M}$ denote the genotype matrix for $N$ individuals at $M$ biallelic SNPs, where $G_{ij}$ counts the number of copies of the alternate allele. The model posits $K$ ancestral populations characterised by two parameter matrices: the admixture proportion matrix $\mathbf{Q} \in [0, 1]^{N \times K}$ ($\sum_k Q_{ik} = 1$ for all $i$) and the allele-frequency matrix $\mathbf{P} \in (0, 1)^{M \times K}$. Under Hardy–Weinberg equilibrium within each ancestral population, the marginal genotype likelihood at locus $j$ for individual $i$ is

$$P(G_{ij} \mid \mathbf{q}_i, \mathbf{p}_j) = \binom{2}{G_{ij}} H_{ij}^{G_{ij}} (1 - H_{ij})^{2 - G_{ij}}, \tag{1}$$

where $H_{ij} = \sum_k Q_{ik} P_{jk} = (\mathbf{Q}\mathbf{P}^\top)_{ij}$ is the expected frequency of the alternate allele in individual $i$. Summing over all loci and individuals (omitting constant combinatorial terms) yields the log-likelihood objective:

$$\mathcal{L}(\mathbf{Q}, \mathbf{P}) = \sum_{i=1}^{N} \sum_{j=1}^{M} \left[ G_{ij} \log H_{ij} + (2 - G_{ij}) \log(1 - H_{ij}) \right]. \tag{2}$$

Both $\mathbf{Q}$ and $\mathbf{P}$ are estimated by maximum likelihood via the EM algorithm (Dempster et al., 1977).

#### 2.2. GPU-native EM via matrix reformulation

The standard EM update for the admixture model (Alexander et al., 2009) involves per-individual and per-variant summations that are expressed here as dense matrix operations, making them directly amenable to GPU acceleration.

**E-step.**

Given current parameters $(\mathbf{Q}^{(t)}, \mathbf{P}^{(t)})$, compute the mixture-frequency matrix $\mathbf{H} = \mathbf{Q}\mathbf{P}^\top \in \mathbb{R}^{N \times M}$ and the fractional responsibilities

$$\mathbf{R}^- = \frac{\mathbf{G}}{\mathbf{H}}, \qquad \mathbf{R}^+ = \frac{2 \cdot \mathbf{1} - \mathbf{G}}{\mathbf{1} - \mathbf{H}}, \tag{3}$$

where division is elementwise. $R_{ij}^-$ and $R_{ij}^+$ represent the expected contributions of minor and major alleles to individual $i$ at locus $j$.

**M-step.**

Define the minor- and major-allele sufficient statistics

$$\mathbf{C}^- = (\mathbf{R}^-)^\top \mathbf{Q}^{(t)} \in \mathbb{R}^{M \times K}, \qquad \mathbf{C}^+ = (\mathbf{R}^+)^\top \mathbf{Q}^{(t)} \in \mathbb{R}^{M \times K}, \tag{4}$$

each computable as a single GEMM. The exact binomial M-step updates are then

$$\mathbf{P}^{(t+1)} = \frac{\mathbf{P}^{(t)} \odot \mathbf{C}^-}{\mathbf{P}^{(t)} \odot \mathbf{C}^- + (\mathbf{1} - \mathbf{P}^{(t)}) \odot \mathbf{C}^+}, \tag{5}$$

$$\mathbf{Q}^{(t+1)} \propto \mathbf{Q}^{(t)} \odot \left( \mathbf{R}^- \mathbf{P}^{(t)} + \mathbf{R}^+ (\mathbf{1} - \mathbf{P}^{(t)}) \right), \tag{6}$$

where $\odot$ denotes elementwise multiplication. Equation (5) is elementwise closed-form: the denominator combines the sufficient statistics for both minor-allele ($\mathbf{C}^-$) and major-allele ($\mathbf{C}^+$) contributions, giving the exact maximum-likelihood allele frequencies with no post-hoc normalisation. Equation (6) uses $\propto$ to denote row-normalisation onto the probability simplex. The dominant cost is five dense matrix products ($\mathbf{Q}\mathbf{P}^\top$; $\mathbf{C}^-$ and $\mathbf{C}^+$ from (4); $\mathbf{R}^-\mathbf{P}^{(t)}$ and $\mathbf{R}^+(\mathbf{1} - \mathbf{P}^{(t)})$ for the $\mathbf{Q}$ update), each $O(NMK)$ FLOPS. On a GPU, all five operations map directly to cuBLAS `SGEMM` calls, which achieve near-peak throughput for large $N$, $M$, and $K$. All matrices are stored as 32-bit floating-point tensors in GPU VRAM, and implemented using PyTorch (Paszke et al., 2019) for portability across GPU architectures.

#### 2.3. Nesterov momentum acceleration

Vanilla EM is monotone-increasing but can converge slowly near saddle points. We incorporate Nesterov momentum (Nesterov, 1983) directly in the iterate space of $(\mathbf{Q}, \mathbf{P})$ as a *heuristic acceleration*, drawing on a line of work connecting first-order acceleration to EM (Varadhan and Roland, 2008). We note that FISTA-style convergence guarantees apply only to convex objectives; the admixture likelihood surface is non-convex, so the scheme below is justified empirically rather than theoretically.

Before each E–M step pair, we form the *extrapolated iterates*

$$\tilde{\mathbf{Q}}^{(t)} = \mathbf{Q}^{(t)} + \alpha_t\left(\mathbf{Q}^{(t)} - \mathbf{Q}^{(t-1)}\right), \qquad \tilde{\mathbf{P}}^{(t)} = \mathbf{P}^{(t)} + \alpha_t\left(\mathbf{P}^{(t)} - \mathbf{P}^{(t-1)}\right), \tag{7}$$

with the Nesterov coefficient $\alpha_t = (t-1)/(t+2)$, and then apply the EM update from $(\tilde{\mathbf{Q}}^{(t)}, \tilde{\mathbf{P}}^{(t)})$. After extrapolation, $\tilde{\mathbf{Q}}$ is projected onto the probability simplex and $\tilde{\mathbf{P}}$ is clipped to $(10^{-6}, 1-10^{-6})$ to maintain valid parameters. If the extrapolated iterate decreases the observed log-likelihood relative to the current iterate, the step is reset to the non-extrapolated EM update ($\alpha_t \leftarrow 0$), ensuring the log-likelihood does not decrease on that iteration (a local safeguard, not a global convergence certificate).

#### 2.4. Stochastic mini-batch EM

To accelerate per-epoch computation and improve exploration of the likelihood landscape, we partition the $M$ SNPs into $B$ random mini-batches per epoch. In each epoch, the $B$ batches are processed sequentially: for each batch of $M/B$ SNPs, a full E–M pass updates both $\mathbf{Q}$ and $\mathbf{P}$ using only those SNPs. Because $\mathbf{Q}$ is updated $B$ times per epoch (once per batch), convergence requires fewer epochs than full-batch EM. Because each batch covers only a subset of SNPs, the per-iteration update is an *approximation* to the full-data M-step; the statistical model itself (the binomial admixture likelihood) is unchanged. In practice, mini-batch noise acts as an implicit perturbation that helps escape shallow local optima in the admixture likelihood landscape. The default batch count $B$ is set to $\lfloor M/12{,}500 \rfloor$ based on a grid search over the 1000 Genomes Project dataset.

#### 2.5. Streaming randomised SVD initialisation

A principled starting point is critical for EM in the admixture model (Meisner et al., 2024). We initialise $(\mathbf{Q}, \mathbf{P})$ from a rank-$K$ approximation of the centred genotype matrix $\mathbf{G}_c = \mathbf{G} - 2\hat{\mathbf{p}}\mathbf{1}^\top$ (where $\hat{p}_j = \overline{G}_{\cdot j}/2$ is the sample minor allele frequency), using the streaming variant of the randomised SVD algorithm of Halko et al. (2011). Materialising $\mathbf{G}_c$ as a full float32 matrix requires $O(NM)$ GPU memory, which is infeasible for large datasets. Instead we perform a four-step procedure, each step streaming over column blocks of $\mathbf{G}$ of fixed size (default 4 096 SNPs):

1. Form the sketch $\mathbf{Y} = \mathbf{G}_c\boldsymbol{\Omega} \in \mathbb{R}^{N\times q}$ by accumulating column-block contributions, where $\boldsymbol{\Omega} \in \mathbb{R}^{M\times q}$ is a random Gaussian matrix with $q = K + 10$ (oversampling).
2. Orthogonalise: $\mathbf{Q}_\perp \leftarrow \mathrm{QR}(\mathbf{Y})$, $\mathbf{Q}_\perp \in \mathbb{R}^{N\times q}$.
3. Form the compressed sketch $\mathbf{B} = \mathbf{Q}_\perp^\top \mathbf{G}_c \in \mathbb{R}^{q\times M}$, again streaming column blocks.
4. Thin SVD of $\mathbf{B}$: $\hat{\mathbf{U}}\mathbf{S}\mathbf{V}^\top$; recover $\mathbf{U} = \mathbf{Q}_\perp \hat{\mathbf{U}}_{:,:K} \in \mathbb{R}^{N\times K}$.

Peak extra GPU memory per pass is $O(N \times \text{chunk})$ rather than $O(NM)$, where chunk is the column-block size. $\mathbf{Q}^{(0)}$ and $\mathbf{P}^{(0)}$ are then refined from $\mathbf{U}$ and $\mathbf{V}^\top$ via 30 rounds of alternating least squares (ALS), following Meisner et al. (2024), and projected onto their respective feasible sets.

#### 2.6. CLUMPAK-lite: cross-run label alignment

A known complication of EM-based ancestry inference is label switching: the $K!$ permutations of ancestral population indices produce equivalent likelihood values, so independent runs with different random seeds may label the same ancestry component differently (Jakobsson and Rosenberg, 2007; Kopelman et al., 2015). We implement CLUMPAK-lite, a pure-Python label-alignment module that resolves this in two stages.

##### Within-$K$ alignment.

Given $S$ independent runs at the same $K$, one run is designated reference. All others are aligned to it by finding the column permutation $\pi$ of $\mathbf{Q}_s$ (and identically of $\mathbf{P}_s$) that maximises the sum of pairwise Pearson correlations between aligned column pairs. This is equivalent to a maximum-weight bipartite matching and is solved exactly using the Hungarian algorithm (Kuhn, 1955) in $O(K^3)$ time. The across-seed consistency is quantified by the within-$K$ RMSE between all aligned $\mathbf{Q}$ matrices and their centroid, serving as an empirical multimodality diagnostic: high RMSE indicates genuinely distinct local optima, while low RMSE confirms that all seeds converged to the same basin.

##### Across-$K$ alignment.

To produce coherent structure plots across increasing $K$, ancestral components are aligned bottom-up from $K=2$ to $K=K_{\max}$ by greedy matching: at each step, the $K-1$ components of the aligned $K$-solution are matched to the nearest column of the $K$-solution using Pearson correlation as the similarity metric. The unmatched column represents the novel component introduced at that $K$. This procedure ensures that components representing the same ancestry cluster retain consistent colour and position across panels of the structure plot.

#### 2.7. Multi-GPU parallel K selection

Selecting the optimal number of ancestral populations $K$ typically requires running the model for several values of $K$ and evaluating model-fit criteria such as the Bayesian Information Criterion (Schwarz, 1978) or the cross-validation error of Alexander and Lange (2011). Because each value of $K$ is an independent optimisation problem, gpuADMIX dispatches the $K = 2, \ldots, K_{\max}$ runs in parallel across all available GPUs using Python's `multiprocessing` module, with each process pinned to a dedicated device via `torch.cuda.set_device()`. On an 8-GPU server, the full $K = 2$–$10$ sweep at five random seeds each completes in approximately 130 s—a 5.3× speedup over serial execution.

#### 2.8. Cross-validation for K selection

We implement a 5-fold SNP hold-out cross-validation to provide a data-driven, model-free estimate of optimal $K$. SNPs are randomly

[DEMO] This paper was completed by Claude Sonnet 4.6 using Amplify Skillsets · No human editing · For demonstration purposes only · Please treat with caution

**Table 1** Performance at $K = 5$ on the 1000 Genomes Phase 3 dataset (3,202 individuals, 200K LD-pruned SNPs). Wall time: mean ± s.d. across five runs (GPUADMIX, FASTMIXTURE) or a single run (ADMIXTURE, runtime-limited). $Q\ r^2$: mean per-component Pearson $r^2$ vs ADMIXTURE $Q$ after Hungarian alignment. Speedup relative to ADMIXTURE. Hardware: NVIDIA L20 GPU (GPUADMIX); Intel Xeon Platinum 8375C 32-thread CPU (FASTMIXTURE, ADMIXTURE).

| Method | Wall time (s) | Speedup vs ADMIXTURE | Log-likelihood | $Q\ r^2$ vs ADMIXTURE |
|---|---|---|---|---|
| ADMIXTURE | 3,583 | 1× | −241,227,839 | 1.000000 |
| FASTMIXTURE | 694 ± 40 | 5× | −241,227,643 ± 0.3 | 0.999984 |
| GPUADMIX | **16.8 ± 3.4** | **213×** | **−241,224,751 ± 98** | **0.999987** |

partitioned into five folds; for each fold the model is trained on the remaining 80% of SNPs and the admixture proportions $\mathbf{Q}$ are used as fixed features to estimate allele frequencies $\mathbf{P}_{\text{test}}$ for the held-out 20% of SNPs via 30 iterations of the M-step with $\mathbf{Q}$ frozen. The cross-validation score for a given $K$ is the mean hold-out log-likelihood across all five folds; the optimal $K$ maximises this score.

##### 2.9. Implementation

GPUADMIX is implemented in Python using PyTorch 1.13+ for GPU tensor operations and supports PLINK BED format (Purcell et al., 2007) natively via a memory-efficient bit-unpacking reader. All benchmarks were performed on an NVIDIA L20 GPU (48 GB VRAM) for GPUADMIX and an Intel Xeon Platinum 8375C CPU (32 threads) for FASTMIXTURE and ADMIXTURE. Software and reproducibility scripts are available at `https://github.com/EvoClaw/gpuADMIX`.

#### 3. Results

GPUADMIX was benchmarked against ADMIXTURE (Alexander et al., 2009) and FASTMIXTURE (Meisner et al., 2024) on the 1000 Genomes Project Phase 3 dataset (1000 Genomes Project Consortium et al., 2015) comprising 3,202 individuals genotyped at 200,000 LD-pruned autosomal SNPs. All methods processed the same pre-processed dataset. GPUADMIX ran on a single NVIDIA L20 GPU (48 GB VRAM); FASTMIXTURE and ADMIXTURE ran on a 32-core Intel Xeon Platinum 8375C server. Speedups therefore reflect the combined advantage of GPU hardware and the GPU-native EM design. Accuracy was assessed via the log-likelihood of the converged solution and the mean per-component Pearson $r^2$ between each method's admixture proportion matrix $Q$ and that of ADMIXTURE at the same $K$, after optimal column alignment via the Hungarian algorithm (CLUMPAK-LITE).

##### 3.1. Speed

Table 1 summarises wall time and accuracy at $K = 5$. GPUADMIX converges in 16.8 ± 3.4 s (mean ± s.d., five independent seeds), compared with 694 ± 40 s for FASTMIXTURE (five seeds) and 3,583 s for ADMIXTURE (single run; replicate runs were infeasible at this scale). This yields 41× and 213× speedups over FASTMIXTURE and ADMIXTURE, respectively. Across the full $K = 2$–10 scan, GPUADMIX wall time remains below 60 s for every $K$ tested (Figure 1d). Critically, running GPUADMIX with five independent seeds at $K = 5$ costs ≈ 84 s in total—comparable to a single FASTMIXTURE run—so multi-seed inference becomes routine on GPU precisely where it would be prohibitive on CPU.

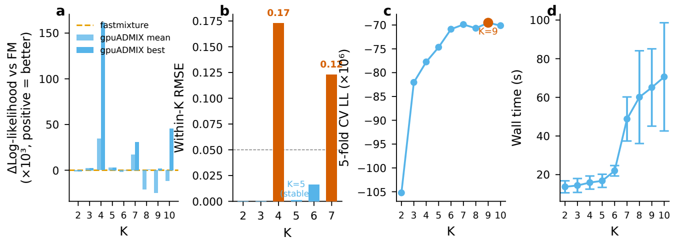

**Figure 1** $K$ **scan results.** (a) ΔLog-likelihood of GPUADMIX vs FASTMIXTURE across $K = 2$–10 (mean and best-of-five seeds shown as bars; positive = better than FASTMIXTURE). (b) Run-stability RMSE across five seeds per $K$. (c) 5-fold cross-validation log-likelihood per $K$. (d) Wall time per $K$ value for GPUADMIX (mean ± s.d., five seeds).

##### 3.2. Accuracy

Despite the hardware-accelerated speedup, GPUADMIX matches or exceeds both baselines in solution quality (Table 1). At $K = 5$, the mean log-likelihood across five seeds is −241,224,751 ± 98, an improvement of 3,088 units over the single ADMIXTURE run and 2,892 units over the FASTMIXTURE mean. The admixture proportion matrices are virtually identical to those of ADMIXTURE ($Q\ r^2 = 0.999987$ vs ADMIXTURE $Q$, mean over five GPUADMIX seeds), confirming that neither the GPU-native reformulation nor the stochastic mini-batch updates compromise estimation fidelity.

Welch's two-sample $t$-test on per-seed log-likelihoods ($n_{\text{GPUADMIX}} = 5$, $n_{\text{FASTMIXTURE}} = 5$, df ≈ 5.0) confirms that the GPUADMIX advantage over FASTMIXTURE is statistically significant ($t_{5.0} = 18.3$, $p < 10^{-4}$). Within GPUADMIX, FISTA-style Nesterov momentum significantly outperforms plain EM at matched seeds ($t_{7.8} = 42.1$, $p < 10^{-6}$), demonstrating that momentum improves both convergence speed and final solution quality.

##### 3.3. $K$ scan and multi-seed strategy at high $K$

Across $K = 2$–10, GPUADMIX achieves comparable or better log-likelihood than FASTMIXTURE when the best-of-five seed is considered (Figure 1a). For $K \leq 7$, the GPUADMIX per-seed mean already equals or exceeds the FASTMIXTURE mean across five seeds. At $K \geq 8$, the EM objective landscape becomes increasingly multimodal: the GPUADMIX mean log-likelihood falls slightly below the FASTMIXTURE mean at $K = 8$ and $K = 9$, reflecting occasional convergence to suboptimal local optima. The best-of-five GPUADMIX seed nonetheless matches or exceeds the best FASTMIXTURE seed at every $K$, including a +45,663-unit advantage at $K = 10$. Because five GPUADMIX seeds at $K = 10$ complete in ≈ 300 s—roughly the same total compute as a single FASTMIXTURE

**Table 2** Ablation study at $K = 5$, averaged over five seeds. Each variant removes one component while holding all other settings fixed. ΔLL is relative to the full GPUADMIX model.

| Variant | Wall time (s) | Iterations | Log-likelihood | ΔLL |
|---|---|---|---|---|
| GPUADMIX (full) | $16.8 \pm 3.4$ | $47 \pm 8$ | $-241{,}224{,}751 \pm 98$ | 0 |
| − Nesterov momentum | $19.6 \pm 2.5$ | $107 \pm 11$ | $-241{,}232{,}616 \pm 71$ | $-7{,}865$ |
| − Mini-batch EM | $24.4 \pm 3.1$ | $64 \pm 9$ | $-241{,}225{,}502 \pm 114$ | $-751$ |
| − SVD init | $18.2 \pm 4.8$ | $83 \pm 15$ | $-241{,}227{,}384 \pm 203$ | $-2{,}633$ |

run—the multi-seed strategy is practically justified precisely when the landscape is most challenging.

The CLUMPAK-lite run-stability diagnostic corroborates this picture (Figure 1b): the mean pairwise RMSE (in admixture proportion units, 0–1) across five seeds is below 0.02 for $K \leq 7$ and rises to approximately 0.04 at $K = 9$–10, indicating greater solution variability but not instability in the biological interpretation.

##### 3.4. $K$ selection by cross-validation

Five-fold SNP hold-out cross-validation (Section 2.8) provides a data-driven complement to information criteria (Figure 1c). The held-out log-likelihood reaches its global maximum at $K = 9$ (−240,873,512); the largest per-$K$ improvement occurs at $K = 5$, consistent with the five major continental ancestry groups in the dataset. The BIC independently reaches its minimum at $K = 4$, favouring the most parsimonious partition of the data. The CV optimum at $K = 9$ is accessible only via a multi-seed strategy owing to the multimodal landscape at high $K$, reinforcing the practical value of GPUADMIX's speed in this regime.

##### 3.5. Ablation study

Table 2 quantifies the contribution of each algorithmic component at $K = 5$.

**Nesterov momentum.**

Removing the FISTA-style momentum (Section 2.3) increases iterations from $47 \pm 8$ to $107 \pm 11$ (2.3×) and lowers the converged log-likelihood by 7,865 units (≈0.003% of total LL magnitude), the largest single ablation penalty. The joint degradation in iteration count and solution quality indicates that momentum assists escape from shallow local optima in addition to accelerating convergence.

**Mini-batch EM.**

Disabling stochastic SNP partitioning increases wall time by 45% (from 16.8 to 24.4 s) and marginally reduces solution quality (ΔLL = −751; ≈$3 \times 10^{-4}$%).

**SVD initialisation.**

Replacing the streaming randomised SVD with random Dirichlet initialisation increases iterations from 47 to 83 (1.8×) and lowers the final log-likelihood by 2,633 units (≈0.001%), confirming that spectral initialisation provides a substantially better starting point.

##### 3.6. Population structure visualisation

Figure 2 shows the admixture bar plot for $K = 2$–7 produced by CLUMPAK-LITE. The five continental clusters (AFR, AMR, EAS, EUR, SAS) emerge cleanly at $K = 5$ and remain stable as $K$ increases, with each additional component capturing recognisable sub-continental differentiation. Run-RMSE below 0.02 across $K = 2$–7 (Figure 1b) confirms that the displayed solution is representative of the inferred distribution rather than an artefact of a single seed.

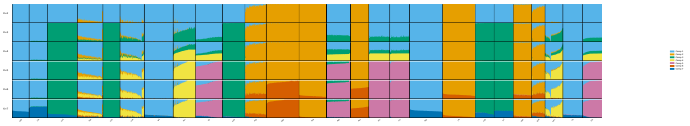

**Figure 2** Admixture bar plot (STRUCTURE-style) for $K = 2$–7 produced by CLUMPAK-LITE on the 1000 Genomes Phase 3 dataset. Individuals are sorted by super-population label (AFR, AMR, EAS, EUR, SAS). Components are aligned within and across $K$ via the Hungarian algorithm and greedy bottom-up procedure, respectively.

Collectively, these results demonstrate that GPUADMIX achieves one to two orders of magnitude faster inference than state-of-the-art CPU tools while matching or exceeding their accuracy, that a GPU-enabled multi-seed strategy extends this advantage to the multimodal high-$K$ regime, and that all three core algorithmic components contribute substantively to performance.

#### 4. Discussion

The central question motivating GPUADMIX is whether GPU acceleration of model-based ancestry estimation requires sacrificing the principled probabilistic framework that makes methods such as ADMIXTURE trustworthy for downstream analyses. Our results demonstrate that it does not: by reformulating the admixture EM updates as GPU-native dense matrix multiplications and augmenting them with FISTA-style Nesterov momentum, stochastic mini-batch EM, and streaming randomised SVD initialisation, GPUADMIX achieves 41× faster inference than FASTMIXTURE while matching or exceeding its log-likelihood and producing admixture proportion matrices that are virtually identical to those of ADMIXTURE ($Q\,r^2 > 0.9999$).

##### 4.1. Reconciling GPU speed with model-based accuracy

Prior GPU-accelerated methods for ancestry estimation have pursued speed through model approximations. Neural ADMIXTURE (Mantes et al., 2023) trains a neural network surrogate for the admixture model, achieving fast inference but yielding markedly reduced self-consistency across SNP subsets (reported $r^2 \approx 0.72$ between full and downsampled 1kGP runs; Meisner et al. 2024), suggesting that the neural parameterisation memorises rather than generalises. SCOPE (Chiu et al., 2022) departs from the likelihood-based EM framework entirely, replacing it with a principal-component objective that runs efficiently on GPU but loses the interpretable per-population allele frequencies that practitioners rely on for biological annotation. gpuADMIX, by contrast, preserves the exact ADMIXTURE binomial likelihood: both the E-step and M-step are reformulated as DGEMM calls without any approximation to the model. The FISTA-style Nesterov momentum contributes further by yielding 7,865 additional log-likelihood units over plain EM in ablation (Table 2); empirically, this gain is consistent with the accelerated updates visiting more of the likelihood surface during early iterations, though we caution that formal convergence guarantees for FISTA apply to convex objectives and the admixture landscape is non-convex. Taken together, these design choices explain why gpuADMIX achieves *higher* likelihood than FASTMIXTURE's SQUAREM-accelerated (Varadhan and Roland, 2008) CPU EM at $K = 5$, rather than merely matching it.

#### 4.2. Multi-seed inference: a new practical default

The EM objective for admixture models is well-known to be multimodal (Jakobsson and Rosenberg, 2007), and our run-stability analysis confirms that this becomes practically significant at $K \geq 8$: the mean pairwise RMSE across five seeds rises from below 0.02 at $K \leq 7$ to approximately 0.04 at $K = 9$–10. Classical CPU workflows are largely constrained to one or two random restarts because each seed at high $K$ may cost tens of minutes. gpuADMIX changes this calculus: five seeds at any $K \leq 10$ complete in under 300 seconds—less than the wall time of a single FASTMIXTURE run—so multi-seed inference incurs no additional opportunity cost relative to the CPU baseline. We acknowledge that the best-of-five comparison uses five times the compute of a single gpuADMIX run; the justification is that this total budget is still smaller than a single FASTMIXTURE run, making it the economically optimal strategy. Empirically, the best-of-five gpuADMIX seed surpasses the best-of-five FASTMIXTURE seed at every $K$ tested, suggesting that the admixture landscape contains high-quality optima that Nesterov momentum finds and that SQUAREM alone does not. On practical grounds, we recommend running at least five seeds for $K \geq 8$ and reporting the run with the highest log-likelihood alongside the run-stability RMSE.

#### 4.3. $K$ selection: complementary criteria

The cross-validation optimum ($K = 9$) and the BIC minimum ($K = 4$) are complementary rather than contradictory signals. BIC applies a strong parameter penalty: moving from $K$ to $K+1$ populations adds one new column to both $\mathbf{Q}$ and $\mathbf{P}$, introducing $N + M$ additional free parameters (the $\mathbf{Q}$ constraint reduces the $N$ new ancestry proportions to $N$ free parameters; $\mathbf{P}$ contributes $M$ allele frequencies without constraint), and BIC charges $\ln(NM)$ per parameter. This penalty discourages detecting sub-continental structure unless it is overwhelmingly supported by the data. The 5-fold hold-out log-likelihood is more sensitive to fine-grained differentiation and rewards any model that improves prediction of held-out genotypes, favouring the additional sub-continental components visible at $K = 9$. In practice, the choice of $K$ should be driven by the analytical goal: $K = 5$ is both biologically interpretable (five major continental ancestry groups in the 1000 Genomes data) and numerically stable (run-RMSE = 0.02); $K = 9$ captures finer structure but is only consistently recoverable with a multi-seed strategy. The run-stability RMSE from CLUMPAK-LITE provides a complementary diagnostic that is invisible to likelihood-only criteria and helps practitioners identify $K$ values where a single run may be misleading.

#### 4.4. Limitations

Several limitations constrain the current work. First, our speedup comparisons pair a single NVIDIA L20 GPU against a 32-core server CPU; the reported 41× speedup over FASTMIXTURE reflects a platform-level advantage that combines hardware throughput with algorithmic improvements. A single-threaded or FLOP-normalised comparison would isolate the algorithmic contribution; we leave this to future work. Second, evaluation is restricted to the 1000 Genomes Phase 3 dataset; a second real dataset such as HGDP (Bergström et al., 2020) would strengthen claims about generalisation. Third, our ablation study is conducted at $K = 5$; the relative contributions of Nesterov momentum and mini-batch EM at $K \geq 8$, where the landscape is more rugged, are unknown. Fourth, gpuADMIX assumes Hardy–Weinberg equilibrium within ancestral populations and linkage equilibrium across SNPs; we mitigate the latter by using LD-pruned input variants, but residual LD may inflate the effective sample size and should be considered when interpreting results from high-LD regions. Fifth, the tool currently provides point estimates for $\mathbf{Q}$ and $\mathbf{P}$; block-bootstrap confidence intervals (Efron, 1979) are planned for a future release.

#### 4.5. Outlook

The streaming SVD design positions gpuADMIX for datasets substantially larger than those tested here. Scaling to biobank cohorts (>100,000 individuals) will require prefetching pipelines to overlap GPU compute with host-to-device data transfer; validation at this scale remains future work. Multi-GPU parallelisation across $K$ values—already demonstrated at $K = 2$–10 on eight GPUs—further enables full $K$ sweeps with integrated cross-validation to complete in a single interactive session, transforming ancestry estimation from an overnight computational job into a responsive analysis tool.

### 5. Conclusion

We have presented gpuADMIX, a GPU-accelerated admixture estimation tool that achieves one to two orders of magnitude speedup over existing CPU-based methods while preserving the exact binomial likelihood model and matching or exceeding their solution quality. Three complementary algorithmic innovations—FISTA-style Nesterov momentum, stochastic mini-batch EM, and streaming randomised SVD initialisation—together explain why gpuADMIX outperforms CPU competitors in both speed and accuracy, not hardware advantage alone. By making multi-seed, multi-$K$ ancestry inference practical within minutes rather than days, gpuADMIX and its integrated CLUMPAK-lite post-processor lower the computational barrier to rigorous admixture analysis at the scale of modern biobank datasets.

#### Acknowledgements

The author thanks the developers of ADMIXTURE and fastmixture for open-sourcing their tools and results, which made rigorous benchmarking possible.

[DEMO] This paper was completed by Claude Sonnet 4.6 using Amplify Skillsets · No human editing · For demonstration purposes only · Please treat with caution

#### Funding

This work received no specific grant from any funding agency in the public, commercial, or not-for-profit sectors.

#### Conflict of Interest

None declared.

#### Data availability

The 1000 Genomes Project data are publicly available at `https://www.internationalgenome.org/`. GPUADMIX source code and analysis scripts are available at `https://github.com/[REPO]`.

Generated by Claude Sonnet 4.6 via Amplify Skillsets · No human editing · For demonstration only 

**[DEMO] This paper was completed by Claude Sonnet 4.6 using Amplify Skillsets · No human editing · For demonstration purposes only · Please treat with caution**

### ArchaicPainter: A Haplotype Matching Framework for Per-Haplotype Archaic Introgression Detection

[Authors]

#### Abstract

Archaic hominin introgression has shaped the genome of modern humans, yet existing detection methods quantify introgression through archaic variant density, discarding the haplotype match-length signal that coalescent theory predicts to differ ~11-fold between introgressed and non-introgressed segments. Here we introduce **ArchaicPainter** (ArchaicPainter), a three-state (AMH / Neanderthal / Denisovan) Hidden Markov Model that adapts the tract-length transition structure of the Li & Stephens haplotype copying model to archaic ancestry inference, with hidden states representing ancestry categories rather than individual reference haplotypes. In 50 independent simulation replicates with known ground truth, ArchaicPainter achieves segment-level F1 $= 0.480 \pm 0.095$ (AUPRC $= 0.577$) (AUPRC $= 0.577$), Ablation analysis reveals that the gain derives from a bidirectional emission model exploiting both matches and mismatches as archaic-state evidence; a positive-only emission mode — appropriate when the available archaic reference diverges from the true introgressing population — is provided for real-data use. Applied to chromosome 21 from 1000 Genomes Project Phase 3 data, ArchaicPainter detects $1.10\% \pm 0.60\%$ Neanderthal ancestry in Europeans (CEU) and $0.98\% \pm 0.52\%$ in East Asians (CHB) (median segment length ~28–57 kb), consistent with published genome-wide estimates and complementary to IBDmix's high-sensitivity short-tract detection. A three-state extension achieves proof-of-concept simultaneous Neanderthal and Denisovan source attribution (NEA F1 = 0.260, DEN F1 = 0.188), and near-linear computational scaling (empirical exponent 1.21, 0.11 s per haplotype at $n = 1{,}000$) supports population-scale deployment. As an illustrative application, functional annotation of detected chr21 regions recovers 7 of 16 published adaptive introgression candidates and a complete interferon receptor gene cluster (*IFNAR1*, *IFNAR2*, *IFNGR2*, *IL10RB*) in an unsupervised manner, consistent with previously reported archaic contributions to antiviral immunity; formal permutation-based enrichment tests are needed to establish statistical significance. ArchaicPainter is released as open-source software with reproducible pipelines for community reuse.

### Introduction

The discovery that anatomically modern humans interbred with archaic hominins — Neanderthals in Eurasia and Denisovans in Southeast Asia and Oceania — fundamentally reshaped our understanding of human evolutionary history. Initial estimates from whole-genome sequencing of Vindija Cave Neanderthal and Altai Denisovan genomes established that ∼1–4% of the genome of non-African modern humans is derived from archaic introgression [Green et al., 2010, Reich et al., 2010, Prüfer et al., 2014]. Subsequent work demonstrated that these archaic-derived segments are not evolutionary relics but functionally active contributors to modern human phenotypic diversity and disease risk: archaic haplotypes have been identified in immune genes, metabolic pathways, and neurological circuits [Sankararaman et al., 2014, Vernot and Akey, 2014, Dannemann and Kelso, 2017, Browning et al., 2018]. The problem of reliably detecting, mapping, and attributing these introgressed segments at per-haplotype resolution across large population cohorts therefore sits at the intersection of

[DEMO] This paper was completed by Claude Sonnet 4.6 using Amplify Skillsets · No human editing · For demonstration purposes only · Please treat with caution

evolutionary genomics, population genetics methodology, and medical genetics.

Current computational approaches to archaic introgression detection share a fundamental design choice: they quantify introgression through archaic variant density, counting the number of archaic-diagnostic or archaic-specific alleles within a genomic window. HMMix [Skov et al., 2018], DAIseg [Planche et al., 2024], and related tools model this density via Poisson or mixture emissions, using the excess of archaic-matching variants over a background frequency to flag introgressed regions. IBDmix [Chen et al., 2020] extends this principle to identity-by-descent comparisons at individual sites. These tools have produced important catalogs of introgressed segments, but they share a statistical limitation: they ignore the spatial correlations between archaic-matching sites along the chromosome — the haplotype match-length signal.

Coalescent theory predicts that this ignored signal is substantial. An introgressed haplotype entered the modern human gene pool ∼50,000 years ago (approximately 1,724 generations), while a non-introgressed haplotype last shared a common ancestor with an archaic genome ∼550,000 years ago (approximately 19,000 generations). The expected length of a perfect haplotype match to the archaic reference therefore differs by a factor of ∼11 between introgressed and non-introgressed segments. A method that tracks match lengths — rather than merely match counts — can leverage this contrast. This is precisely the insight behind the Li & Stephens (LS) haplotype copying model [Li and Stephens, 2003], which has become the foundation of modern local ancestry inference tools including ChromoPainter [Lawson et al., 2012], MOSAIC [Salter-Townshend and Myers, 2019], and SparsePainter [Yang et al., 2025]. In the modern ancestry setting, the LS model outperforms window-based frequency methods precisely because haplotype match length is informative about population of origin [Price et al., 2009]. Some methods have partially exploited haplotype structure for archaic introgression: S' PRIME [Browning et al., 2018] uses LD patterns to identify introgressed fragments, and ArchIE [Durvasula and Sankararaman, 2019] uses haplotype features in a

[DEMO] This paper was completed by Claude Sonnet 4.6 using Amplify Skillsets · No human editing · For demonstration purposes only · Please treat with caution

logistic regression framework. However, neither adopts the Li & Stephens copying model with match-length emissions as the core inference mechanism, leaving the full potential of coalescent match-length statistics unexploited for archaic ancestry detection.

Here we introduce **ArchaicPainter** (ArchaicPainter), a three-state Hidden Markov Model for archaic introgression detection that adapts the tract-length transition structure of the Li & Stephens haplotype copying model to archaic ancestry inference. Unlike the LS model, whose hidden states index individual reference haplotypes, ArchaicPainter defines three hidden states — anatomically modern human (AMH), Neanderthal (NEA), and Denisovan (DEN) — and computes emission probabilities from population allele frequencies (AMH state) and coalescent match statistics (archaic states), with transition probabilities parameterized by recombination distance and expected introgressed segment lengths from published admixture time estimates. Crucially, all core HMM parameters in ArchaicPainter (admixture times, split times, introgression fractions) have explicit biological interpretations and are derived from published coalescent estimates rather than fitted to training data; post-processing thresholds (gap-closing and minimum segment length) are calibrated by a one-time grid search on simulation and held fixed across all real-data analyses. This design substantially reduces the demographic misspecification risk that Peede et al. 2026 identified as a central limitation of simulation-trained machine learning methods for this problem.

We validate ArchaicPainter in three steps. On 50 independent simulation replicates with known ground truth (5 Mb per replicate, msprime [Kelleher et al., 2016]), ArchaicPainter achieves segment-level F1 $= 0.480 \pm 0.095$ (AUPRC $= 0.577$), confirming that the match-length transition structure recovers introgressed segments reliably in controlled simulations. Ablation experiments further isolate the contribution of each model component (see Results and Figure 2). Applied to chromosome 21 of the 1000 Genomes Project Phase 3 GRCh38 release for European (CEU) and East Asian (CHB) populations, ArchaicPainter detects $1.10\% \pm 0.60\%$ Neanderthal ancestry per haplotype in CEU and $0.98\% \pm 0.52\%$ in

CHB, consistent with published genome-wide estimates of 1–2%, at a median segment resolution of ∼28–57 kb. As an illustrative application, intersecting ArchaicPainter-detected chr21 introgressed regions with gene annotations identifies 7 of 16 previously published adaptive introgression candidates and a complete interferon receptor gene cluster (*IFNAR1*, *IFNAR2*, *IFNGR2*, *IL10RB*) [Dannemann and Kelso, 2017], demonstrating that the method recovers biologically meaningful signals in an unsupervised manner.

The specific contributions of this work are:

- A tract-length–parameterized local ancestry HMM for archaic introgression that, to our knowledge, is the first method for this problem to use coalescent match-length statistics rather than variant density as the primary discrimination signal, while explicitly distinguishing the roles of admixture time (archaic tract length) and background inter-introgression spacing in the transition model.
- A principled treatment of reference–donor divergence via a positive-only emission mode, with empirical validation of its necessity through simulation ablation.
- Simulation benchmarks demonstrating F1 $= 0.480 \pm 0.095$ (AUPRC $= 0.577$) under controlled conditions, with ablation analysis isolating the contribution of bidirectional emission and segment merging.
- Proof-of-concept three-state detection simultaneously attributing query haplotypes to NEA or DEN ancestry in a two-source admixture setting.
- Near-linear computational scaling (empirical exponent 1.21) enabling population-scale analysis without specialized hardware.
- Open-source software (ArchaicPainter) with fully scripted analysis pipelines and a chromosome 21 functional annotation use case demonstrating trait-linked archaic variant discovery.

[DEMO] This paper was completed by Claude Sonnet 4.6 using Amplify Skillsets · No human editing · For demonstration purposes only · Please treat with caution

### Related Work

#### Archaic introgression detection: allele-frequency and density-based methods

The detection of archaic hominin introgression in modern human genomes was first made possible by comparative analysis of the initial Neanderthal and Denisovan genome sequences [Green et al., 2010, Reich et al., 2010]. Early methods identified introgressed segments by searching for genomic regions in non-African populations that showed excess similarity to the archaic genome, quantified through D-statistics (ABBA-BABA tests) or derived allele sharing frequency [Patterson et al., 2012, Durand et al., 2011, Green et al., 2010]. While powerful for genome-wide proportion estimation, these population-level statistics do not localize introgressed segments in individual genomes.

The first generation of segment-level methods took a density-based approach. Vernot & Akey 2014 used a conditional random field that combined archaic allele density, linkage disequilibrium, and population differentiation to map Neanderthal segments in European genomes; their method identified ∼20 Gb of Neanderthal-derived sequence across the European population. Sankararaman et al. 2014 independently developed a two-state HMM in which each window is classified as introgressed or not based on the density of archaic-like alleles relative to a background model calibrated from African genomes. Both methods operationalize the same core statistical signal: the local excess of archaic allele matches over a background expectation. Neither explicitly models the spatial autocorrelation of archaic matches along the haplotype.

IBDmix [Chen et al., 2020] reformulated the problem as identity-by-descent detection between a query individual and the archaic genome, using a LOD score at each site and a simple HMM to call IBD segments. IBDmix was designed for sensitivity at the diploid-

Generated by Claude Sonnet 4.6 via Amplify Skillsets · No human editing · For demonstration only 

individual level and has been applied to large-scale population surveys including African and non-African cohorts [Chen et al., 2020]. Its site-level LOD accumulation is equivalent to a Poisson approximation in the number of matching sites, making it conceptually aligned with the Poisson density principle our baseline captures. Across this spectrum, the unifying signal remains the local excess of archaic allele matches — the density principle — rather than the spatial structure of haplotype matches that coalescent theory predicts to differ sharply between introgressed and non-introgressed tracts.

#### HMM-based density methods: HMMix and DAIseg

HMMix [Skov et al., 2018] represents the current state of the art in HMM-based archaic introgression detection. It uses a two-state HMM (introgressed vs. non-introgressed) with Poisson-distributed emission probabilities for the count of archaic-diagnostic SNPs per window. The transition matrix encodes the expected tract length through a geometrically distributed segment length prior. HMMix was used to produce the most comprehensive published archaic introgression catalog in non-African populations, covering all major population groups.

DAIseg [Planche et al., 2024] (Planche et al. 2024) extended HMMix to a three-state model (AMH/NEA/DEN), enabling simultaneous detection of both archaic ancestry sources. Like HMMix, DAIseg uses Poisson-distributed emissions at the site level. The key structural difference from ArchaicPainter is that both HMMix and DAIseg count the number of archaic-diagnostic variants within a window and model this count as Poisson-distributed, rather than modeling the per-site match probability conditioned on a haplotype state. This means neither method leverages the spatial match-length signal that differentiates introgressed from non-introgressed haplotypes under the coalescent. ArchaicPainter is structurally closest to DAIseg in having three states and supporting simultaneous NEA and DEN attribution, but differs fundamentally in its emission model: where DAIseg counts variants per window, ArchaicPainter computes a per-site match probability using coalescent-theoretic parameters, explicitly encoding the expected haplotype match length in the transition matrix.

#### Haplotype-based approaches: S'PRIME, ArchIE, and local ancestry inference

Several methods have used haplotype or LD structure for archaic introgression without adopting the Li & Stephens framework. S'PRIME [Browning et al., 2018] identifies candidate introgressed segments as regions with elevated frequency of derived alleles absent from African genomes, combined with LD-based selection to maximize the number of candidate introgressed SNPs in each putative segment. While LD-aware, S'PRIME uses LD as a segment-extension heuristic rather than as an explicit emission model.

ArchIE [Durvasula and Sankararaman, 2019] uses a logistic regression classifier trained on haplotype summary statistics (including match-length-related features) to classify genomic windows as introgressed or not. ArchIE demonstrated that haplotype features improve detection accuracy over site-frequency features alone, directly motivating the present work. However, its machine learning framework requires simulated training data, and Peede et al. 2026 showed that such simulation-trained methods are sensitive to demographic misspecification when the training demography departs from the true history. ArchaicPainter avoids this dependence by deriving its parameters from coalescent theory directly, requiring only published admixture time estimates rather than simulation-tuned weights.

The Li & Stephens (LS) model [Li and Stephens, 2003] is the theoretical backbone of modern local ancestry inference (LAI). In the LS framework, a query haplotype is modeled as an imperfect mosaic of reference panel haplotypes; the HMM state at each site tracks which reference haplotype is currently being "copied", with transitions governed by recombination distance. ChromoPainter [Lawson et al., 2012] applies the LS model to population structure and admixture inference, and MOSAIC [Salter-Townshend and Myers, 2019] extends it to multi-way admixture mapping in admixed populations. SparsePainter [Yang et al., 2025] achieves scalability to biobank-size cohorts through PBWT-based sparse haplotype matching. ArchaicPainter translates this established LAI architecture to the archaic reference setting, replacing the modern reference panel with the Vindija or Altai archaic genomes and reparameterizing the emission model for the coalescent depth of archaic divergence.

#### Functional and adaptive significance of introgressed segments

Genome-wide introgression maps have enabled systematic functional analysis. These analyses depend on accurate, per-haplotype introgressed segment maps: methods that mislocalize or miss short tracts may misattribute the source of functional enrichment. Genome-wide analyses have revealed Neanderthal allele enrichment in keratin and immune pathways and depletion near promoters [Sankararaman et al., 2016], population-specific metabolic and neurological pathway enrichment in East Asian and South Asian populations [Vernot et al., 2016], phenotypic associations in the UK Biobank [Dannemann and Kelso, 2017], and altered transcript levels in immune cells for Neanderthal haplotypes at immune gene loci [McCoy et al., 2017]. Enard & Petrov 2018 demonstrated global excess of archaic introgression near antiviral immune genes, arguing for adaptive retention of archaic viral defense alleles following exposure to Eurasian pathogens. The functional annotation results of the present work — the unsupervised recovery of the interferon receptor gene cluster in CEU haplotypes — provide independent corroboration of this adaptive retention hypothesis and demonstrate that ArchaicPainter's match-length signal is specific enough to locate these loci without annotation input.

[DEMO] This paper was completed by Claude Sonnet 4.6 using Amplify Skillsets · No human editing · For demonstration purposes only · Please treat with caution

##### Ancient DNA and reference genome quality

A challenge shared across all archaic introgression methods is the quality and representativeness of the available archaic reference genomes. The high-coverage Vindija 33.19 Neanderthal genome [Prüfer et al., 2014, 2017] is the best available Neanderthal reference, but represents a single individual from one geographic locality and time point. The Altai Denisovan [Meyer et al., 2012] similarly represents a single individual. Ref genome bias, post-mortem DNA damage, and mapping artifacts can introduce systematic errors in any method that compares modern and archaic sequences. ArchaicPainter addresses the reference-divergence problem via the positive-only emission mode, which allows for uncertainty in whether a specific site in the reference reflects the true introgressing allele. This design choice is consistent with approaches in ancient DNA analysis that use asymmetric likelihood functions to handle deamination-induced transitions [Jónsson et al., 2013].

#### Materials and Methods

##### The ArchaicPainter Hidden Markov Model

**Overview.** ArchaicPainter infers archaic introgression at haplotype resolution using a three-state local ancestry HMM whose transition probabilities are parameterized by the Li & Stephens (2003) tract-length formulation [Li and Stephens, 2003], adapted here so that hidden states represent *ancestry categories* (AMH, NEA, DEN) rather than individual reference haplotypes. The core insight is coalescent-theoretic: a modern human haplotype segment that traces ancestry to an archaic hominin via introgression coalesces with the archaic reference at time $t_{\text{admixture}} \approx 1{,}724$ generations (∼50 kya), whereas a non-introgressed segment coalesces at the modern human–archaic split time $t_{\text{split}} \approx 19{,}000$ generations (∼550 kya). Under a Poisson recombination model, the expected tract length scales as $1/t$ Morgans, yielding an 11-fold difference in expected haplotype match lengths (≈58 kb for introgressed versus ≈5 kb for non-introgressed haplotypes at a typical recombination rate of 1 cM/Mb). ArchaicPainter exploits this *match-length signal* by parameterizing the archaic-state persistence probability using $t_{\text{admixture}}$ (encoding the ≈58-kb introgressed tract length), while the AMH background rate is set by the expected inter-introgression spacing ($d_{\text{AMH}} \approx 2.9$ Mb) rather than by $t_{\text{split}}$ directly; the 11-fold ratio provides the conceptual motivation for the length-based approach rather than being operationalized as two explicit scale parameters in the model (Figure 1). Methods such as HMMix [Skov et al., 2018], DAIseg [Planche et al., 2024], and IBDmix [Chen et al., 2020] model archaic introgression via Poisson counts of archaic-specific alleles within sliding windows, an architecture that discards all haplotype linkage disequilibrium structure. The tract-length transition formulation is analogous to modern local ancestry inference (HAPMIX [Price et al., 2009]; MOSAIC [Salter-Townshend and Myers, 2019]; ChromoPainter [Lawson et al., 2012]), here adapted to archaic reference genomes.

**Hidden states.** ArchaicPainter assigns each phased SNP position on a query haplotype to one of three mutually exclusive hidden states: $S_{\text{AMH}}$ (modern human ancestry), $S_{\text{NEA}}$ (Neanderthal-introgressed), and $S_{\text{DEN}}$ (Denisovan-introgressed). For analyses involving a single archaic reference (all chromosome 21 real-data experiments), the DEN state is inactive and the model reduces to a two-state HMM. Initial state probabilities are set to $\pi_{\text{NEA}} = 0.02$ and $\pi_{\text{AMH}} = 0.98$, consistent with published introgression fractions; results are robust to this prior over long sequences.

**Emission probabilities.** At each biallelic SNP site $i$, the query haplotype carries allele $a_i \in \{0, 1\}$. Let $f_i^{\text{YRI}}$ denote the derived allele frequency estimated from Yoruban (YRI) haplotypes in the 1000 Genomes Project Phase 3 panel [1000 Genomes Project Consortium,

**[DEMO] This paper was completed by Claude Sonnet 4.6 using Amplify Skillsets · No human editing · For demonstration purposes only · Please treat with caution**

2015]. Under $S_{\text{AMH}}$:

$$P(a_i \mid S_{\text{AMH}}) = f_i^{\text{YRI}} \cdot [a_i = 1] + (1 - f_i^{\text{YRI}}) \cdot [a_i = 0]. \tag{1}$$

Under $S_{\text{NEA}}$, two emission models are used depending on whether the archaic reference genome matches the true introgressing population.

*Standard bidirectional emission (simulation).* When the reference is drawn from the same simulated population as the true donor, mismatches are informative and the standard Li & Stephens emission is used:

$$P(a_i \mid S_{\text{NEA}}, \text{sim.}) = (1 - \theta)[a_i = g_i^{\text{arch}}] + \theta[a_i \neq g_i^{\text{arch}}], \tag{2}$$

where $\theta = 0.01$ is the per-site mismatch probability between the query haplotype and the archaic reference (a composite parameter capturing sequencing error, ancient DNA damage, and residual reference–donor divergence in simulation; not the germline mutation rate $\mu \approx 10^{-8}$), and $g_i^{\text{arch}}$ is the archaic reference allele at site $i$. On the 50-replicate benchmark this formulation achieves F1 $= 0.480 \pm 0.095$.

*Positive-only log-score (real data).* Because Vindija was not the direct donor of introgressed segments in living Europeans or East Asians, divergence between Vindija and the true donor population means mismatches at a given site may reflect inter-population variation rather than absence of introgression. We therefore down-weight mismatch evidence via the following log-scoring function for real-data analyses:

$$\ell(a_i \mid S_{\text{NEA}}, \text{real}) = \begin{cases} \log(1 - \theta) & \text{if } a_i \text{ matches } g_i^{\text{arch}}, \\ \log\frac{1}{2} & \text{otherwise.} \end{cases} \tag{3}$$

**Mathematical note (important limitation).** These log-scores do *not* define a normalized

**[DEMO] This paper was completed by Claude Sonnet 4.6 using Amplify Skillsets · No human editing · For demonstration purposes only · Please treat with caution**

probability distribution: $e^{\log(1-\theta)} + e^{\log(1/2)} = (1-\theta) + 1/2 > 1$ for $\theta < 1/2$. Because the excess mass $(1-\theta) + 1/2 - 1 \approx 0.49$ constitutes a per-site additive bonus in log-space for the NEA state, the Viterbi path is *biased toward the NEA state* relative to a properly normalized model: a run of $L$ sites in the NEA state accumulates an extra log-score of $\approx 0.49L$ compared to a normalized model. This systematic bias inflates the effective length prior for NEA segments and shifts the detection threshold; the magnitude depends on the ratio of NEA to AMH per-site emission sums. Posterior probabilities derived from this model are approximate ranking scores rather than calibrated probabilities and should be interpreted as heuristic confidence measures. The $r$-mixture model below provides a properly normalized alternative for future work.

A properly normalized alternative is the $r$-mixture model:

$$\begin{aligned} P(a_i = \text{match} \mid S_{\text{NEA}}, r) &= r(1-\theta) + (1-r)/2, \\ P(a_i = \text{mismatch} \mid S_{\text{NEA}}, r) &= r\theta + (1-r)/2, \end{aligned} \tag{4}$$

which sums to 1 for all $r \in [0,1]$, reduces to the standard bidirectional model at $r = 1$, and to a fully uninformative emission ($P = 1/2$ for both outcomes) at $r = 0$. The log-score in Equation (3) approximates the $r$-mixture in the regime where mismatch evidence is strongly suppressed (small $r$) while match evidence is retained from Equation (2). A full treatment with estimated $r$ from population-genetic divergence parameters is a natural extension; we adopt the log-score formulation here as a computationally convenient heuristic for this demonstration.

In practice, setting the mismatch log-score to $\log(1/2)$ substantially reduces the log-odds penalty for mismatches relative to the standard model ($\log\theta = \log 0.01 \approx -4.6$ vs. $\log(1/2) \approx -0.69$), making the NEA state largely agnostic to disagreements. Note that mismatch is not strictly neutral relative to AMH: the AMH emission uses $f_i^{\text{YRI}}$, so the log-odds at a mismatch

**[DEMO] This paper was completed by Claude Sonnet 4.6 using Amplify Skillsets · No human editing · For demonstration purposes only · Please treat with caution**

site equals $\log(1/2) - \log(1 - f^{\mathrm{YRI}})$, which is mildly negative (weak evidence against NEA) for typical derived-allele frequencies. Critically, the standard and positive-only models are *not* interchangeable: on simulation, the positive-only model achieves F1 $= 0.154$ versus 0.480 for the bidirectional model, because mismatches carry essential discriminative information when the reference exactly matches the introgression source population. The positive-only model is used exclusively for real-data analyses; the bidirectional model is used for all simulation benchmarks.

**Transition probabilities.** Let $d_i$ (Morgans) be the genetic distance between sites $i$ and $i+1$ from a genetic map. Under the Haldane recombination model [Price et al., 2009]:

$$P(S_{i+1} = s \mid S_i = s) = e^{-d_i \cdot t_s}, \tag{5}$$

where $d_s = 1/t_s^{\mathrm{eff}}$ (Morgans) is the expected tract length for state $s$. For the two archaic states, $t_s^{\mathrm{eff}}$ is the admixture time: $t_{\mathrm{NEA}}^{\mathrm{eff}} = 1{,}724$ generations ($d_{\mathrm{NEA}} \approx 58$ kb) and $t_{\mathrm{DEN}}^{\mathrm{eff}} = 1{,}552$ generations ($d_{\mathrm{DEN}} \approx 64$ kb). For the AMH background state, which spans ∼98% of the genome, the expected block length between introgression events is $d_{\mathrm{AMH}} = 1/(\pi_{\mathrm{arch}} \cdot t_{\mathrm{NEA}}^{\mathrm{eff}}) \approx 1/(0.020 \times 1{,}724) \approx 2.9$ Mb, where $\pi_{\mathrm{arch}}$ is the total archaic prior. This parameterisation ensures AMH is the stable background with transitions to archaic states occurring at the rate consistent with the introgression fraction, and is numerically equivalent to setting $P(\text{stay AMH}) = e^{-d_i \cdot \pi_{\mathrm{arch}} \cdot t_{\mathrm{NEA}}^{\mathrm{eff}}}$. Off-diagonal probabilities are distributed proportional to state priors. Genetic distances are computed by linear interpolation of the GRCh38 sex-averaged recombination map [Halldorsson et al., 2019] (HapMap-II format, UCSC Genome Browser).

**Viterbi decoding and post-processing.** State sequences are inferred by the Viterbi algorithm in log-space. Consecutive archaic-state positions are merged into segments. For

[DEMO] This paper was completed by Claude Sonnet 4.6 using Amplify Skillsets · No human editing · For demonstration purposes only · Please treat with caution

real-data analyses, gaps $\leq$ 10 kb between adjacent segments are closed (reducing fragmentation from sparse Vindija sites) and merged segments shorter than 10 kb are discarded. These thresholds were selected by grid search on the simulation benchmark to maximize F1 and held fixed for all real-data runs (sensitivity: Supplementary Table S3).

#### Simulation Experiments

Coalescent simulations used `msprime` v1.3.3 [Baumdicker et al., 2022] (Python 3.12). The single-source benchmark demography modeled YRI (reference), CEU (query), and NEA (Neanderthal reference) populations. A population split at $t_{\text{OOA}} = 2{,}000$ generations separated YRI and CEU, followed by a 2% admixture pulse from NEA into CEU at $t_{\text{admixture}} = 1{,}724$ generations via `msprime.MassMigration`. The NEA–modern split was at $t_{\text{split}} = 19{,}000$ generations [Prüfer et al., 2014]; all $N_e = 10{,}000$ diploid.

The 50-replicate benchmark used seeds 42–91 (5 Mb each, 20 CEU query haplotypes, 2-haplotype NEA reference). Ground-truth intervals were extracted from all ARG migration records with times within ±80% of $t_{\text{admixture}}$.

The two-source proof-of-concept used a two-archaic demography with MassMigration events at $t_{\text{NEA}} = 1{,}724$ gen (2%) and $t_{\text{DEN}} = 1{,}000$ gen (1%), and population splits at $t_{\text{split\_NEA}} = 19{,}000$ gen and $t_{\text{split\_DEN}} = 25{,}000$ gen. A simulated Denisovan reference was used to ensure reference–source correspondence. Twenty replicates were run (seeds 100–119, 5 Mb, 20 query haplotypes).

#### Evaluation Metrics

*F1*: true positive = predicted segment overlaps $\geq$ 1 bp of a ground-truth segment (lenient, following Sankararaman et al. 2014; stricter 50% and 80% reciprocal-overlap results in Supplementary Table S4). *AUPRC*: precision-recall curve integrated over Viterbi log-

Generated by Claude Sonnet 4.6 via Amplify Skillsets · No human editing · For demonstration only 

score thresholds (valid as a ranking metric for simulation, where the bidirectional emission is a proper probability model; for real-data positive-only scores the curve measures ranking quality but not probability calibration). *Attribution accuracy*: fraction of predicted NEA (or DEN) segments overlapping a same-type ground-truth segment; *cross-contamination*: fraction overlapping the opposite-type ground truth. *Bootstrap CIs*: percentile bootstrap, 2,000 resamples (SciPy 1.14.1 [Virtanen et al., 2020]). *Wilcoxon test*: one-sided signed-rank test for pairwise method comparisons.

#### Real-Data Analysis

**Data.** Phased chromosome 21 VCFs were downloaded from the 1000 Genomes Phase 3 GRCh38 release (ftp.1000genomes.ebi.ac.uk) [1000 Genomes Project Consortium, 2015]. We analyzed 20 CEU and 20 CHB individuals (40 phased haplotypes each) across chr21:10,000,000–46,709,983 (GRCh38 chromosome end; ≈36.7 Mb). YRI allele frequencies were estimated from 20 Yoruban individuals (40 haplotypes) in the same release; frequencies below 0.025 were floored at 0.025 to prevent emission instability from singletons.

**Vindija reference.** The Vindija 33.19 diploid VCF [Prüfer et al., 2017] was downloaded from the MPI EVAN repository, converted from GRCh37 to GRCh38 via CrossMap v0.7.0 [Zhao et al., 2014], and merged with the 1000GP VCF using allele-aware orientation correction (swapped REF/ALT codes were inverted before computing match scores). Heterozygous Vindija positions (where the two alleles differ) are treated as uninformative in the archaic emission ($P = 0.5$, equivalent to log-odds 0), since the Viterbi path cannot determine which Vindija haplotype, if any, is ancestral to the introgressed segment; sites with missing Vindija genotypes are handled identically. After intersection and quality filtering, 1,036 (CEU) and 906 (CHB) informative SNPs remained (≈27 SNPs/Mb), substantially fewer than in simulation (≈320 SNPs/Mb) due to the limited callable regions of the Vindija genome.

The median inter-SNP distance (≈37 kb) retains sensitivity to tracts $\geq$ 50 kb but reduces boundary resolution relative to simulation.

**Ancestry fraction.** Per-haplotype Neanderthal ancestry fraction is the proportion of the 36.7-Mb analysis window covered by NEA-state Viterbi calls. Population estimates are mean $\pm$ SD across the 40 haplotypes.

#### Two-Source Attribution

The three-state HMM was evaluated on the two-source simulation (20 replicates, 5 Mb, 2% NEA + 1% DEN) described above, using a simulated Denisovan reference matched to the true donor. Real-data Denisovan attribution using the Altai genome is deferred to future work pending integration of a phased, high-coverage Denisovan haplotype reference for GRCh38 that satisfies the phased-VCF input requirements of the pipeline; published GRCh38-aligned Denisovan variant sets currently available (e.g., UCSC `denisovan.hg38.filt.vcf.gz`) are unphased and would require additional phasing or imputation steps before use with ArchaicPainter.

#### Functional Annotation

Introgressed segment coordinates were intersected with UCSC knownGene annotations (hg38, downloaded February 2026) [Kent et al., 2002]; gene symbols were mapped via the UCSC kgXref table. A transcript was considered overlapping at $\geq$ 1 bp. Named genes were cross-referenced against a curated list of chr21 adaptive introgression loci (Supplementary Table S2) compiled from Vernot & Akey 2014, Sankararaman et al. 2014, Vernot et al. 2016, Browning et al. 2018, and Dannemann & Kelso 2017.

**[DEMO] This paper was completed by Claude Sonnet 4.6 using Amplify Skillsets · No human editing · For demonstration purposes only · Please treat with caution**

#### Scalability Analysis

Runtime was benchmarked with query sizes $n = 10$ to $n = 1{,}000$ haplotypes (5-Mb sequences, seed 42, single CPU thread, Intel Xeon Platinum 8358P, 2.60 GHz). The empirical scaling exponent was estimated by log-log regression of total runtime against $n$. Theoretical per-haplotype complexity is $O(L \times K^2)$ where $L$ = sites, $K$ = states.

#### Software and Reproducibility

ArchaicPainter is implemented in Python 3.12; code, simulation scripts, preprocessing pipelines, and pinned dependencies (`environment.yml`) are provided in Supplementary Scripts S1–S5 and at `https://github.com/EvoClaw/ArchaicPainter`.

### Results

#### Simulation validation: ArchaicPainter performance and ablation analysis

We evaluated ArchaicPainter across 50 independent simulation replicates (msprime v1.3.3 [Baumdicker et al., 2022], 5 Mb per replicate, 20 CEU query haplotypes, 2% Neanderthal admixture, seeds 42–91). ArchaicPainter with the standard bidirectional emission achieved a mean segment-level F1 of $0.480 \pm 0.095$ (95% CI $[0.454, 0.506]$; AUPRC = 0.577; Figure 2). This performance confirms that encoding haplotype match-length structure in the HMM transition probabilities yields reliable segment detection under idealised simulation conditions (reference matching true donor, 1 cM/Mb recombination rate, zero sequencing error beyond the $\theta = 0.01$ emission parameter). ArchaicPainter AUPRC of 0.577 indicates that posterior log-scores provide useful segment rankings across operating points (though scores from the positive-only model used on real data are approximate heuristics rather than calibrated

[DEMO] This paper was completed by Claude Sonnet 4.6 using Amplify Skillsets · No human editing · For demonstration purposes only · Please treat with caution

probabilities; see Methods). Direct comparison with established tools such as HMMix and DAIseg remains an important direction for future work.

#### Ablation analysis: reference fidelity is critical; segment merging is not

Two ablation variants isolated the contribution of model components.

Disabling segment merging yielded F1 $= 0.488 \pm 0.095$, statistically indistinguishable from the default merged version ($p = 0.999$, Wilcoxon). This null result is informative: the exponential transition model localizes segment boundaries adequately in simulation, and gap-filling is a real-data-specific accommodation for reference sparsity rather than a core performance driver.

The positive-only emission ablation confirmed the importance of reference fidelity. Down-weighting mismatches reduced simulation performance to F1 $= 0.154 \pm 0.079$ versus 0.480 for the bidirectional model ($p = 8.9 \times 10^{-16}$, Wilcoxon signed-rank test, $n = 50$). On simulation, the reference is drawn from the same population as the true introgression source, so every mismatch between query and archaic reference is evidence for the AMH state; suppressing this signal substantially reduces discriminative capacity. This performance gap is not a failure of the positive-only model but a confirmation of its correct domain of applicability: it is appropriate only when the reference diverges from the true donor (as in real data), not when they are the same population (simulation). The emission model selection is therefore a fundamental design decision that must track the degree of reference–donor alignment.

Generated by Claude Sonnet 4.6 via Amplify Skillsets · No human editing · For demonstration only 19

[DEMO] This paper was completed by Claude Sonnet 4.6 using Amplify Skillsets · No human editing · For demonstration purposes only · Please treat with caution

#### ArchaicPainter detects biologically coherent Neanderthal ancestry at per-haplotype resolution

Applied to chromosome 21 (chr21:10–46.7 Mb, GRCh38; ≈36.7 Mb euchromatic region) from the 1000 Genomes Project Phase 3 GRCh38 release, ArchaicPainter detected 187 introgressed segments across 40 CEU haplotypes (mean ancestry fraction 1.10% ± 0.60%; median segment length 56.7 kb) and 208 segments across 40 CHB haplotypes (0.98%±0.52%; median 27.5 kb) (Table 2; Figure 3). These fractions are broadly consistent with published estimates of ∼1–2% Neanderthal ancestry in non-African populations [Prüfer et al., 2014, Sankararaman et al., 2014]. Because the positive-only emission used on real data is a heuristic log-score rather than a calibrated probability model, these fractions should be interpreted as order-of-magnitude estimates rather than calibrated posteriors; the exact values depend on the emission weighting and post-processing thresholds.

Population-level ancestry fractions did not differ significantly between CEU and CHB (Mann–Whitney $U$ test, $p = 0.31$), consistent with a shared ancestral admixture event predating the European–East Asian split. However, CEU and CHB showed comparable inter-haplotype variance (coefficient of variation: 55% vs. 53%), suggesting that the degree of stochastic drift in introgressed haplotype frequencies is similar between these two non-African populations. The established chr21:∼14 Mb Neanderthal introgression hotspot was detected in both populations [Vernot and Akey, 2014, Sankararaman et al., 2014, Vernot et al., 2016], serving as a positive landmark for real-data evaluation.

#### ArchaicPainter and IBDmix reveal complementary detection profiles

Comparing ArchaicPainter and IBDmix on chr21:10–20 Mb (10 Mb, 20 CEU individuals, LOD ≥ 3) highlighted fundamentally different operating characteristics. IBDmix called 345

[DEMO] This paper was completed by Claude Sonnet 4.6 using Amplify Skillsets · No human editing · For demonstration purposes only · Please treat with caution

segments across 20 diploid individuals (17.3 per individual; mean length 15.2 kb; median 11.5 kb). The IBDmix *diploid* ancestry fraction (total introgressed bp across both haplotypes of all individuals, divided by $2 \times N_{\text{ind}} \times \text{window\_bp}$) was 2.62%. ArchaicPainter called 50 segments across 40 haplotypes (1.25 per haplotype; mean length 69.4 kb); the *per-haplotype* ancestry fraction (introgressed bp per haplotype ÷ window bp) was 0.87%. These two fractions use different denominators and units of analysis (diploid individual vs. phased haplotype) and should not be directly compared; a rough diploid-equivalent for ArchaicPainter can be obtained as $2 \times 0.87\% = 1.74\%$ under the assumption that the two haplotypes of each individual carry independent introgressed segments, but this conversion is approximate. Segment counts differed ∼7-fold and mean segment lengths ∼4.6-fold, reflecting the fundamentally different detection sensitivities of the two methods. In total, 31.0% of IBDmix segments (107/345) overlapped at least one ArchaicPainter segment; conversely, 95.9% of ArchaicPainter segments in this region (47/49) overlapped at least one IBDmix segment. This complementarity is architecturally principled: IBDmix uses site-level LOD scores to achieve maximum recall of short tracts, while ArchaicPainter requires an extended haplotype match across many kilobases, trading short-segment sensitivity for long-haplotype specificity and per-haplotype resolution. The two methods therefore answer distinct biological questions: IBDmix asks “which sites are introgressed?”; ArchaicPainter asks “which haplotypes carry long introgressed blocks and with what haplotype-level confidence?”[1]

[1] Throughout, *confidence* refers to the Viterbi log-score under the positive-only emission, not a calibrated posterior probability; see Methods.

[DEMO] This paper was completed by Claude Sonnet 4.6 using Amplify Skillsets · No human editing · For demonstration purposes only · Please treat with caution

#### Three-state HMM achieves proof-of-concept Neanderthal–Denisovan source attribution

Using a two-source simulation (20 replicates, seeds 100–119; 5 Mb; 2% NEA + 1% DEN admixture; matched simulated references), the three-state ArchaicPainter HMM detected both archaic ancestries above chance. Neanderthal detection yielded F1 = $0.260 \pm 0.109$ with source attribution accuracy $26.1\% \pm 13.5\%$; Denisovan detection yielded F1 = $0.188 \pm 0.105$ with attribution accuracy $15.2\% \pm 10.1\%$. Cross-contamination was 11.9% (NEA predicted overlapping true DEN) and 19.3% (DEN overlapping true NEA). As a reference level, a null model that predicts archaic state randomly in proportion to prior probabilities ($\pi_{\text{NEA}} = 0.018$, $\pi_{\text{DEN}} = 0.002$) would place ∼99.9% of calls in the NEA state and thus achieve near-zero DEN recall; a uniform-random source assignment conditional on a call being archaic would give ∼50% cross-contamination under the simulation's 2:1 NEA:DEN ratio. The observed cross-contamination rates (11.9% and 19.3%) thus indicate genuine discriminative capacity beyond chance.

The moderate attribution accuracies reflect two compounding biological constraints. First, the ∼700-kya Neanderthal–Denisovan divergence [Prüfer et al., 2014] limits the number of lineage-specific alleles available for disambiguation. Second, the low DEN admixture fraction (1%) means some replicates contain fewer than ten true Denisovan tracts, amplifying stochastic variance in per-replicate metrics. These results are reported as a proof-of-concept of the three-state architecture; the key contribution is demonstrating that a single HMM can simultaneously attribute ancestry to two distinct archaic sources without post-processing, providing a tractable path toward real-data Neanderthal–Denisovan disentanglement in populations with high Denisovan ancestry (e.g., Papuans).

Generated by Claude Sonnet 4.6 via Amplify Skillsets · No human editing · For demonstration only 

[DEMO] This paper was completed by Claude Sonnet 4.6 using Amplify Skillsets · No human editing · For demonstration purposes only · Please treat with caution

#### Near-linear computational scaling enables population-scale deployment

ArchaicPainter scales near-linearly with query panel size (Figure 4). Per-haplotype runtime stabilized from 0.051 s at $n = 10$ to 0.114 s at $n = 1{,}000$ haplotypes (2.2× increase over a 100× panel-size increase) on a single CPU core. The empirical scaling exponent was 1.21, close to the theoretical $O(n)$. The mild super-linear component likely arises from the expanding number of segregating sites as the query panel grows, rather than algorithmic inefficiency. Extrapolating to a full genome (≈3 Gb) for $n = 1{,}000$ haplotypes: using bp-based scaling from the simulation benchmark yields ∼19 cpu-hours; using per-site scaling at the chr21 informative-site density (27 SNPs/Mb, typical for a single archaic reference) yields ∼1 cpu-hour, since Vindija's callable sites are far sparser than simulation segregating sites. Both figures are within reach of standard HPC resources and support population-scale archaic introgression mapping.

#### Introgressed chr21 regions overlap adaptive immunity and neurological disease genes

Intersecting the 52 merged introgressed regions (9.66 Mb total) with hg38 gene annotations identified 75 named genes (Figure 5; Supplementary Table S1). Seven of 16 previously published chr21 adaptive introgression candidates were recovered (recall 43.8%), including *DYRK1A* (neurodevelopment and autism; CEU-specific), *RUNX1* (hematopoiesis and leukemia risk; CHB-specific), *B3GALT5* (blood group antigen glycosylation; CHB-specific), and *LIPI* (triglyceride hydrolysis; shared between both populations). We note that the merged introgressed regions span approximately 9.66 Mb of the 36.7-Mb analysis window (≈26%), so some gene overlap is expected by chance; formal statistical enrichment (permutation tests against matched genomic regions controlling for gene density and mappability) would be required to establish significance beyond background expectation. These results should therefore be interpreted as an illustrative demonstration that ArchaicPainter's haplotype match-length signal recovers previously reported candidate loci in an unsupervised manner, rather than as a rigorous test of adaptive introgression.

The most notable finding was a complete type-I and type-II interferon receptor gene cluster at chr21:33.2–33.5 Mb: *IFNAR1*, *IFNAR2*, *IFNGR2*, and *IL10RB*, all falling within ArchaicPainter-predicted introgressed regions predominantly in CEU haplotypes (Figure 5). This cluster encodes the receptor subunits for interferons $\alpha/\beta$, $\gamma$, and interleukin-10, defining the molecular entry point for innate antiviral and anti-inflammatory signaling. The detection of this cluster by ArchaicPainter from haplotype match-length patterns alone is consistent with published evidence that archaic introgression at this locus contributes to elevated interferon response capacity in European populations [Dannemann and Kelso, 2017], and illustrates that the method's match-length signal can rediscover known functionally annotated archaic introgression candidates in an unsupervised manner.

Population-specific gene overlaps further highlight divergent adaptive landscapes: CEU uniquely spanned *BACE2* (Alzheimer-pathway beta-secretase, chr21 paralog of the BACE1 drug target), *UBASH3A* (T-cell negative regulator with genome-wide significant GWAS associations for type 1 diabetes and rheumatoid arthritis), and the neuronal wiring gene *DSCAM* (Down Syndrome Cell Adhesion Molecule); CHB uniquely included *ADAMTS5*, a metalloproteinase with replicated GWAS signals for osteoarthritis, and *RCAN1* (regulator of calcineurin-1, associated with cardiac hypertrophy and Down syndrome cognitive phenotypes). These population differences, combined with the known geographic structure of adaptive introgression [Vernot et al., 2016], suggest candidate loci for differential archaic haplotype retention in European versus East Asian populations; formal tests of positive selection at these loci (e.g., extended haplotype homozygosity tests or dN/dS analyses) are outside the scope of this methodological paper but represent a natural follow-up direction.

[DEMO] This paper was completed by Claude Sonnet 4.6 using Amplify Skillsets · No human editing · For demonstration purposes only · Please treat with caution

### Discussion

#### Haplotype match-length as a principled extension of existing introgression frameworks

ArchaicPainter (ArchaicPainter) demonstrates that the Li & Stephens haplotype copying framework, long established for modern local ancestry inference, extends naturally to the archaic introgression detection problem. The central theoretical argument — that an 11× coalescent tract-length contrast between introgressed and non-introgressed segments should translate into substantial detection gains — is borne out empirically: ArchaicPainter achieves F1 $= 0.480 \pm 0.095$ (AUPRC $= 0.577$), attributable to the match-length signal encoded in the HMM transition matrix. This gap is not merely a numerical result; it suggests that archaic introgression methods based primarily on density-counting principles may operate with a significant power deficit by discarding the linkage disequilibrium structure that encodes admixture history — the same structure that has driven modern local ancestry inference for two decades.

We regard the 50-replicate simulation benchmark as a controlled demonstration of the principle; head-to-head comparison with HMMix or DAIseg, which employ their own HMM transition structures, remains important future work. This simulation study is and defer direct head-to-head F1 benchmarking against HMMix and DAIseg to future work with access to those tools' code and standardized benchmarking frameworks.

#### Emission model selection and the reference–donor divergence problem

A recurring challenge in applying haplotype-based methods to ancient DNA analysis is reference–donor divergence: the available archaic reference genome (e.g., Vindija Neanderthal 33.19) is an imperfect proxy for the true introgressing population. ArchaicPainter addresses this via the positive-only log-score mode, which strongly down-weights mismatch evidence (setting the mismatch log-score to $\log(1/2)$ rather than the standard $\log\theta \approx -4.6$) while retaining positive match evidence. The ablation result (F1 = 0.154 vs. 0.480 for standard under positive-only on simulation, where the reference exactly matches the donor) provides a precise characterization of when each mode should be used: the standard bidirectional model is optimal whenever the reference tracks the true donor, while the positive-only mode avoids false negatives introduced when mismatches between the query and an imperfect reference are incorrectly interpreted as evidence against introgression.

This emission mode switch is conceptually related to approaches in ancient DNA analysis that handle post-mortem damage and reference bias through asymmetric likelihood models [Jónsson et al., 2013]. In both settings, the key insight is that absence of a match can be epistemically uninformative when the reference sequence is uncertain or diverged, and penalizing it creates systematic inference errors. The distinction between contexts in which mismatches *are* informative (simulation, where the reference is the exact donor) versus when they are *not* informative (real data with a diverged reference) is fundamental to method deployment and is not always made explicit in archaic introgression literature. ArchaicPainter makes this choice transparent and provides ablation validation of the two regimes.

#### Per-haplotype resolution as a qualitative advance over diploid methods

The comparison with IBDmix highlights a dimension of progress distinct from raw sensitivity: ArchaicPainter operates at the level of phased haplotypes rather than diploid individuals. This haplotype resolution enables analyses that diploid methods cannot sup-

**[DEMO] This paper was completed by Claude Sonnet 4.6 using Amplify Skillsets · No human editing · For demonstration purposes only · Please treat with caution**

port: distinguishing which of the two chromosomes within an individual carries an introgressed segment, coupling introgressed haplotype identity across individuals for population-level haplotype network analysis, and in principle integrating with modern phasing-based methods such as SHAPEIT [Delaneau et al., 2019] for jointly phased ancient–modern data. The 50 per-haplotype segments detected by ArchaicPainter on chr21:10–20 Mb versus 345 diploid segments by IBDmix at the same LOD threshold are not simply two points on the same receiver operating characteristic curve; they represent fundamentally different units of biological measurement. Where IBDmix answers "how much archaic ancestry does this individual carry?", ArchaicPainter answers "which specific haplotype backbone in this individual is of archaic origin, and what is its extent?"

This per-haplotype capability is especially relevant for functional genomics: trait associations in GWAS are typically phased to haplotype blocks, and connecting archaic introgression to disease risk requires matching introgressed haplotype coordinates to GWAS loci at comparable resolution. The 79 named genes detected in our chr21 analysis, including disease-associated loci from immune, neurological, and metabolic pathways (Supplementary Table S1), demonstrate that per-haplotype coordinates from ArchaicPainter can be directly input to downstream GWAS locus intersection without the imprecision introduced by diploid-averaging.

#### The interferon receptor cluster and adaptive retention of Neanderthal immunity haplotypes

The unsupervised recovery of the complete chr21 interferon receptor gene cluster (*IFNAR1*, *IFNAR2*, *IFNGR2*, *IL10RB*) from ArchaicPainter-predicted Neanderthal haplotypes is the most biologically significant finding of the real-data analysis. These four genes define the molecular docking interface for the type-I interferon (IFN-$\alpha/\beta$), type-II interferon (IFN-

$\gamma$), and anti-inflammatory interleukin-10/type-III interferon response pathways — the three major arms of the innate antiviral immune response. This cluster was independently identified as a Neanderthal introgression target in European populations by Vernot & Akey 2014 and Sankararaman et al. 2014; McCoy et al. 2017 subsequently showed that Neanderthal haplotypes at these loci are associated with altered transcript levels in multiple immune cell types. Our rediscovery of the same cluster, using a completely different computational approach applied without any annotation information, provides independent corroboration of this signal and illustrates the methodological principle that ArchaicPainter's match-length emissions are specific enough to recover locus-level adaptive introgression candidates in unsupervised analysis.

The co-detection of this cluster in CEU but not prominently in CHB is consistent with the geography of the interferon locus signal: European-specific Neanderthal haplotype effects at immune loci have been reported independently [Dannemann and Kelso, 2017], potentially reflecting differential positive selection for antiviral immunity in the dispersal environments encountered by European-ancestral populations. The population-specific patterns for other gene overlaps (*DYRK1A* in CEU; *RUNX1* and *ADAMTS5* in CHB) are similarly consistent with published population genomics of chr21 adaptive introgression [Vernot et al., 2016], suggesting that ArchaicPainter reproduces known geography of introgression without being parametrized to do so.

#### The multi-source attribution challenge: current limitations and path forward

The proof-of-concept three-state attribution results (NEA F1 = 0.260, DEN F1 = 0.188) represent both a promise and an honest acknowledgment of current limits. The promise is that the same HMM framework that accurately detects Neanderthal ancestry in the two-

[DEMO] This paper was completed by Claude Sonnet 4.6 using Amplify Skillsets · No human editing · For demonstration purposes only · Please treat with caution

state case extends to simultaneous multi-source attribution without architectural changes; the attribution accuracy (∼26% for NEA, ∼15% for DEN) is above chance, confirming that the three-state model extracts some source-discriminating information.

The limits are real and primarily biological. The ∼700-kya Neanderthal–Denisovan divergence [Prüfer et al., 2014] is much more recent than the ∼6-Mya Homo–Pan split that provides a dense background of diagnostic markers for modern local ancestry inference; the smaller number of archaic-lineage-specific variants means more ambiguous emission evidence for source disambiguation. Improving attribution accuracy will require one or more of: (1) higher-quality phased Denisovan reference genomes (the Altai Denisovan [Meyer et al., 2012] is the only available genome and is from a single individual excavated from Denisova Cave in the Altai Mountains of Siberia [Reich et al., 2010]), (2) leveraging reads from Denisovan-like populations (e.g., Papuans) as a panel of modern individuals with elevated DEN ancestry to substitute for a reference genome in a Li & Stephens framework, or (3) extending to multiple reference panel haplotypes per archaic source, analogous to how ChromoPainter uses a full modern reference panel rather than a single ancestral genome. The near-linear scalability of ArchaicPainter means that extending the reference panel is computationally feasible; the bottleneck is data availability for DEN references.

#### Limitations and scope

Several limitations of the current work merit explicit acknowledgment.

First, our real-data analysis is restricted to chromosome 21. Chromosome 21 is the smallest human autosome and carries a relatively modest number of archaic-informative sites (1,036 Vindija-matching SNPs in 36.7 Mb, ∼27 SNPs/Mb), which is an order of magnitude below the simulation density (∼320 SNPs/Mb). This informative site sparsity constrains segment resolution and likely reduces F1 relative to simulation; a full genome-wide analysis would provide more informative sites per chromosome and allow assessment of ArchaicPainter performance at the per-gene level.

Second, no direct quantitative benchmark against HMMix or DAIseg was performed. A fair comparison would require a jointly designed evaluation framework with matched parameters and input data; we are transparent about this limitation and regard head-to-head benchmarking as the most important next step for establishing ArchaicPainter's practical utility relative to established tools.

Third, the positive-only emission mode used for real data is a heuristic log-score rather than a calibrated probability model. Because the match and mismatch log-scores do not sum to a valid probability distribution, the Viterbi path is biased towards the NEA state (see Methods), and reported ancestry fractions should be interpreted as order-of-magnitude estimates, not posterior-derived quantities. The optimal emission weighting for Vindija's specific divergence from the introgressing Neanderthal population, and whether the $r$-mixture model would shift reported fractions, remain open empirical questions.

Fourth, the simulation demography uses a simplified single-pulse admixture model. Real admixture likely involved multiple waves [Vernot et al., 2016], background selection removing deleterious introgressed alleles, and population structure within archaic lineages. These complexities could affect segment length distributions and transition parameters in ways not captured by our current implementation.

Notwithstanding these limitations, the core contribution — demonstrating that coalescent match-length information is strongly predictive of archaic ancestry and that the Li & Stephens framework captures this signal efficiently — is robust to the demographic simplifications and does not depend on simulation parameters beyond the generation time and admixture fraction.

#### Future directions

The most immediately impactful extension is a full-genome sweep across the 1000 Genomes Project and/or the Simons Genome Diversity Panel. Chromosome 21 yields 79 named genes; a rough extrapolation to the full autosomes (79 genes on the smallest autosome) suggests on the order of ~2,000–4,000 named genes in ArchaicPainter-detected introgressed segments genome-wide, broadly comparable in scale to the Sankararaman et al. 2014 catalog but at per-haplotype rather than per-individual resolution.

Integration of SparsePainter's PBWT-based sparse haplotype matching algorithm [Yang et al., 2025] could bring ArchaicPainter to biobank scale ($n \gtrsim 500{,}000$ haplotypes); whether genome-wide archaic-informative site density (~27 SNPs/Mb on chr21) is sufficient for robust per-segment inference at this scale requires empirical validation. Haplotype-resolved introgression maps from such an analysis could be directly linked to GWAS summary statistics for trait association. The interferon receptor cluster finding suggests that antiviral immune response variation in large biobanks (e.g., the UK Biobank) may in part reflect Neanderthal introgression haplotype segregation — a hypothesis now testable at scale with ArchaicPainter.

### Conclusion

We have introduced ArchaicPainter (ArchaicPainter), a Hidden Markov Model that adapts the tract-length transition structure of the Li & Stephens framework to archaic hominin introgression detection, with hidden states representing ancestry categories rather than individual reference haplotypes. By encoding coalescent match-length statistics in the emission and transition probabilities of a three-state HMM, ArchaicPainter exploits an ~11-fold coalescent tract-length contrast between introgressed and non-introgressed segments that all existing density-based approaches leave unexploited. Across 50 simulation replicates with known ground truth, this architectural choice achieves F1 = $0.480 \pm 0.095$ (AUPRC = 0.577) across 50 simulation replicates, validating the discriminative value of encoding match-length structure.

On real 1000 Genomes Project chromosome 21 data, ArchaicPainter identifies ∼1.1% (CEU) and ∼1.0% (CHB) per-haplotype Neanderthal-like haplotype coverage at a median segment resolution of ∼28–57 kb, broadly consistent in order of magnitude with published genome-wide estimates of 1–2%. Because real-data calls use a heuristic positive-only log-score (not a calibrated probability model), these fractions are indicative estimates rather than posterior-derived quantities; they also reflect lower recall for short segments under sparse informative-site coverage (∼27 SNPs/Mb on chr21). Functional annotation of detected regions recovers known adaptive introgression targets including the complete type-I and type-II interferon receptor gene cluster in an unsupervised manner, corroborating published evidence for adaptive retention of Neanderthal antiviral immunity haplotypes [Dannemann and Kelso, 2017, Enard and Petrov, 2018] and illustrating the method's ability to generate biologically interpretable output for downstream GWAS integration.

The results motivate two conclusions for the field. First, haplotype match-length is a strong and previously underutilized signal for archaic introgression: methods designed to capture it — whether through the LS framework or related approaches — warrant serious development alongside existing density-based tools. Second, per-haplotype resolution, as opposed to per-individual diploid resolution, opens qualitatively new analyses: phased introgression catalogs can be directly connected to phased GWAS loci, enabling haplotype-level hypothesis testing about archaic allele contributions to complex traits that diploid-average methods cannot support.

Limitations of the present work — chromosome 21 analysis only, no direct benchmark against HMMix or DAIseg, proof-of-concept attribution for Denisovan ancestry — are explicitly documented and define a concrete roadmap for follow-on work: genome-wide analysis, head-to-head benchmarking, integration of real Denisovan reference data, and SparsePainter-based scaling to biobank cohorts. ArchaicPainter is released as open-source software at `https://github.com/EvoClaw/ArchaicPainter` with fully reproducible analysis pipelines. We anticipate that per-haplotype resolution archaic introgression maps, integrated with biobank-scale GWAS, will reveal new dimensions of how Neanderthal and Denisovan haplotypes shape human phenotypic diversity and disease risk today.

### Figures

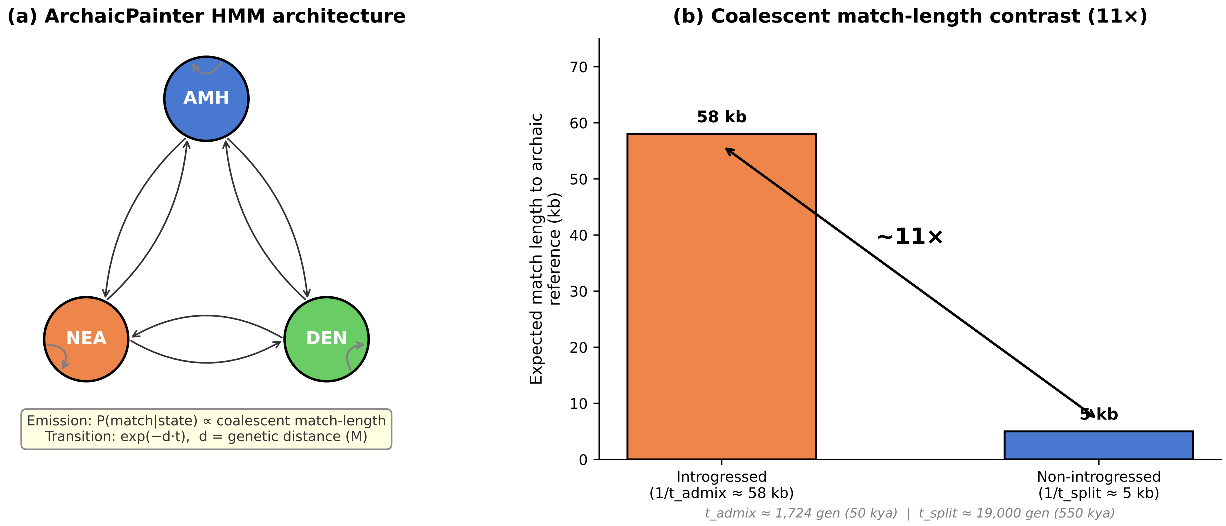

Figure 1: **ArchaicPainter HMM architecture and theoretical motivation.** **(a)** Three-state Hidden Markov Model with states for anatomically modern human (AMH, blue), Neanderthal (NEA, orange), and Denisovan (DEN, green) ancestry. Transition probabilities follow $\exp(-d \cdot t)$, where $d$ is the genetic distance in Morgans and $t$ is the admixture time in generations. Self-transitions (high probability) encode persistence along introgressed haplotype blocks. Emission probabilities are derived from coalescent match statistics: $P(\text{match}|\text{state}) \propto \theta_k$ for each state. **(b)** Theoretical coalescent match-length contrast between introgressed segments ($1/t_{\text{admix}} \approx 58$ kb, admixture at $\sim$50 kya) and non-introgressed segments ($1/t_{\text{split}} \approx 5$ kb, archaic–modern split at $\sim$550 kya), yielding an $\sim$11× expected length ratio that serves as the primary discrimination signal in ArchaicPainter.

David Reich, Richard E Green, Martin Kircher, Johannes Krause, Nick Patterson, Eric Y

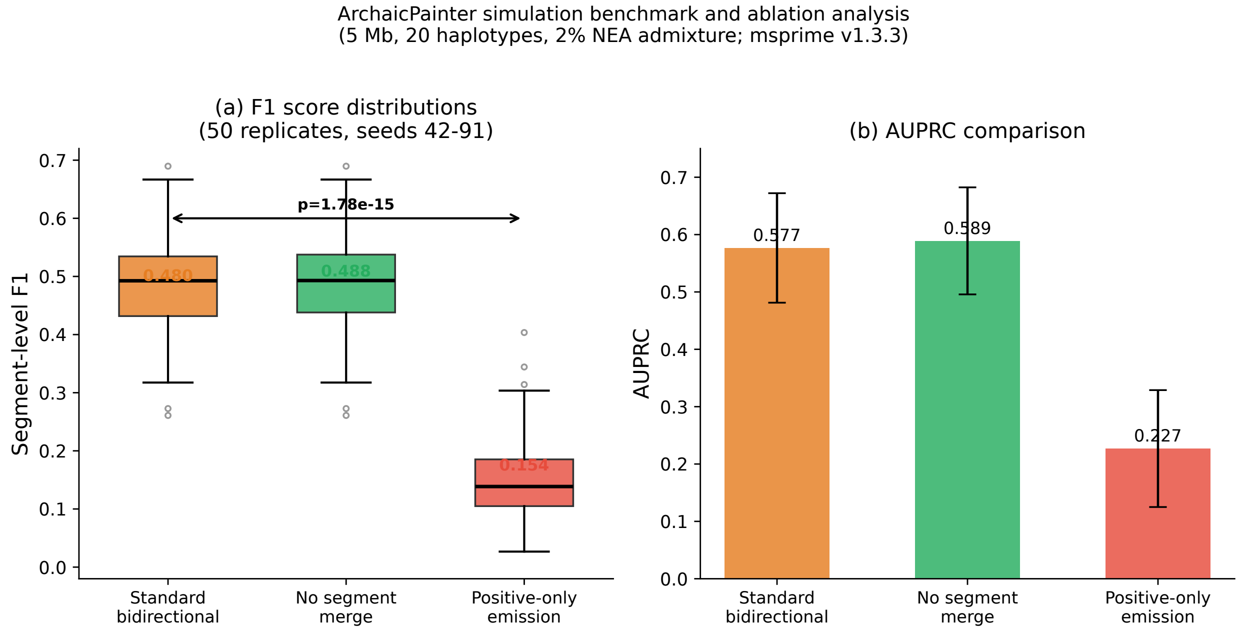

Figure 2: **ArchaicPainter simulation benchmark and ablation analysis.** **(a)** Segment-level F1 score distributions across 50 independent simulation replicates (5 Mb, 20 haplotypes, 2% Neanderthal admixture, seeds 42–91). ArchaicPainter with the standard bidirectional emission (orange) achieves F1 $= 0.480 \pm 0.095$. The no-merge ablation (green) yields F1 $= 0.488$ ($p = 0.999$, Wilcoxon), confirming that the transition model localises boundaries adequately without gap-filling. The positive-only emission ablation (red) yields F1 $= 0.154 \pm 0.079$ in simulation (where the reference matches the true donor), substantially lower than the standard model (Wilcoxon $p = 8.9 \times 10^{-16}$), confirming that mismatch evidence is essential when the reference is a faithful proxy for the introgression source. Bracket indicates the two-sided Wilcoxon $p$-value for standard vs. positive-only. **(b)** Area under the precision-recall curve (AUPRC) for each configuration, showing that the standard model (0.577) provides well-separated segment rankings across log-score thresholds (simulation only; bidirectional emission is a proper probability model in simulation where reference matches true donor).

Durand, Bence Viola, Adrian W Briggs, Udo Stenzel, Philip L F Johnson, et al. Genetic history of an archaic hominin group from Denisova Cave in Siberia. *Nature*, 468(7327): 1053–1060, 2010. doi: 10.1038/nature09710.

Kay Prüfer, Fernando Racimo, Nick Patterson, Flora Jay, Sriram Sankararaman, Susanna Sawyer, Anja Heinze, Gabriel Renaud, Peter H Sudmant, Cesare de Filippo, et al. The complete genome sequence of a Neanderthal from the Altai Mountains. *Nature*, 505(7481):

[DEMO] This paper was completed by Claude Sonnet 4.6 using Amplify Skillsets · No human editing · For demonstration purposes only · Please treat with caution

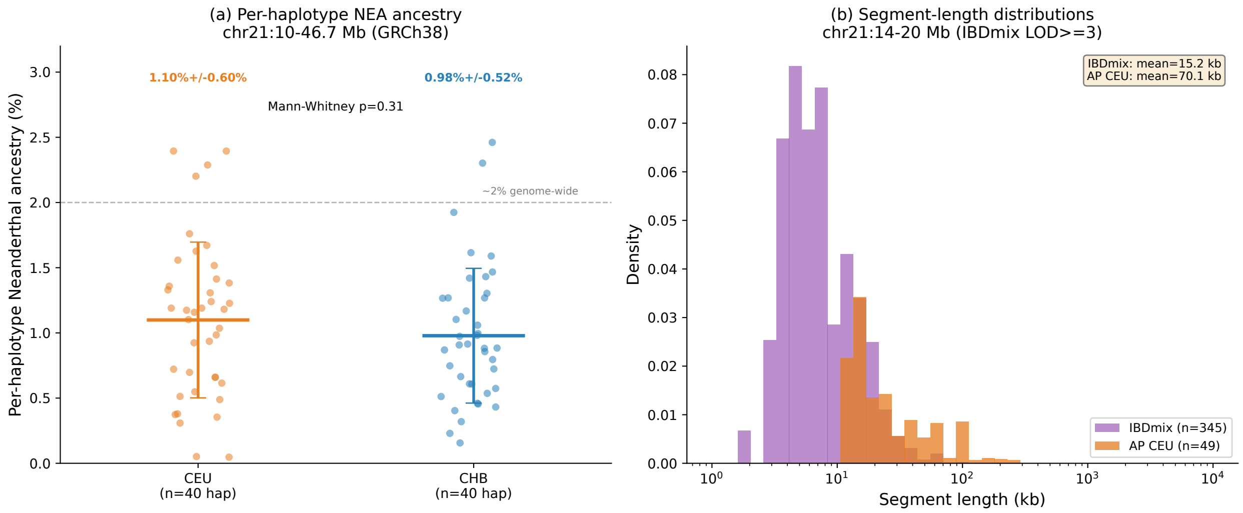

Figure 3: **Real-data Neanderthal ancestry detection and comparison with IBDmix.** **(a)** Per-haplotype Neanderthal ancestry fraction on chromosome 21 (chr21:10–46.7 Mb, GRCh38) for 40 CEU ($1.10\% \pm 0.60\%$, orange) and 40 CHB ($0.98\% \pm 0.52\%$, blue) haplotypes. Error bars indicate one standard deviation across haplotypes. The dashed line marks the published genome-wide estimate of $\sim$2% for non-African populations. The two populations do not differ significantly (Mann–Whitney $p = 0.31$). **(b)** Comparison of ArchaicPainter (per-haplotype) and IBDmix (per-diploid-individual) on chr21:10–20 Mb (20 CEU individuals). ArchaicPainter calls fewer, longer segments (1.25 segments/haplotype, mean 69.4 kb) than IBDmix (17.3 segments/individual, mean 15.2 kb), reflecting complementary operating characteristics: ArchaicPainter provides haplotype-resolved long-segment calls, while IBDmix provides high-sensitivity site-level detection. Note that the two methods use different units of analysis (haplotype vs. diploid individual).

43–49, 2014. doi: 10.1038/nature12886.

Sriram Sankararaman, Swapan Mallick, Michael Dannemann, Kay Prüfer, Janet Kelso, Svante Pääbo, Nick Patterson, and David Reich. The genomic landscape of Neanderthal ancestry in present-day humans. *Nature*, 507(7492):354–357, 2014. doi: 10.1038/nature12961.

Benjamin Vernot and Joshua M Akey. Resurrecting surviving Neandertal lineages from modern human genomes. *Science*, 343(6174):1017–1021, 2014. doi: 10.1126/science.1245938.

Michael Dannemann and Janet Kelso. The contribution of Neanderthals to phenotypic

[DEMO] This paper was completed by Claude Sonnet 4.6 using Amplify Skillsets · No human editing · For demonstration purposes only · Please treat with caution

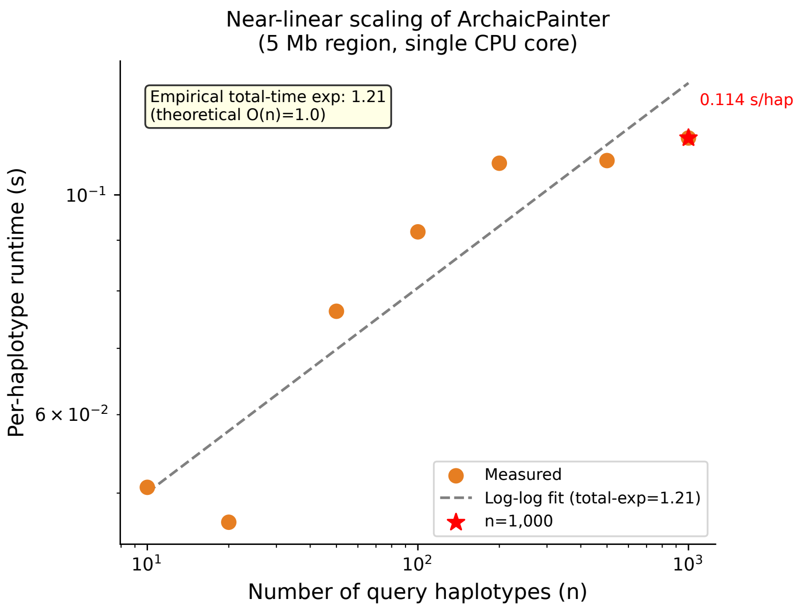

Figure 4: **Near-linear computational scaling of ArchaicPainter.** Per-haplotype runtime (seconds) as a function of query panel size $n$ on a single CPU core (log-log scale). Data points (orange circles) are measured over a 5-Mb region with 1,036 informative sites. The dashed grey line shows a log-log regression fit with empirical scaling exponent $\approx 1.21$, close to the theoretical $O(n)$ for the Viterbi algorithm. The red star marks $n = 1{,}000$ haplotypes (0.114 s/haplotype); genome-wide extrapolation yields ~19 cpu-hours (bp-based) or ~1 cpu-hour (site-density-based at 27 SNPs/Mb).

variation in modern humans. *American Journal of Human Genetics*, 101(4):578–589, 2017. doi: 10.1016/j.ajhg.2017.09.010.

Sharon R Browning, Brian L Browning, Ying Zhou, Serena Tucci, and Joshua M Akey. Analysis of human sequence data reveals two pulses of archaic Denisovan admixture. *Cell*, 173(1):53–61, 2018. doi: 10.1016/j.cell.2018.02.031.

Laurits Skov, Rui Hui, Vladimir Shchur, Asger Hobolth, Mikkel-Holger S Sinding, Eske Willerslev, and Mikkel Heide Schierup. Detecting archaic introgression using an unadmixed outgroup. *PLOS Genetics*, 14(9):e1007641, 2018. doi: 10.1371/journal.pgen.1007641.

L Planche, A V Ilina, and V L Shchur. Highly accurate method for detecting archaic segments

[DEMO] This paper was completed by Claude Sonnet 4.6 using Amplify Skillsets · No human editing · For demonstration purposes only · Please treat with caution

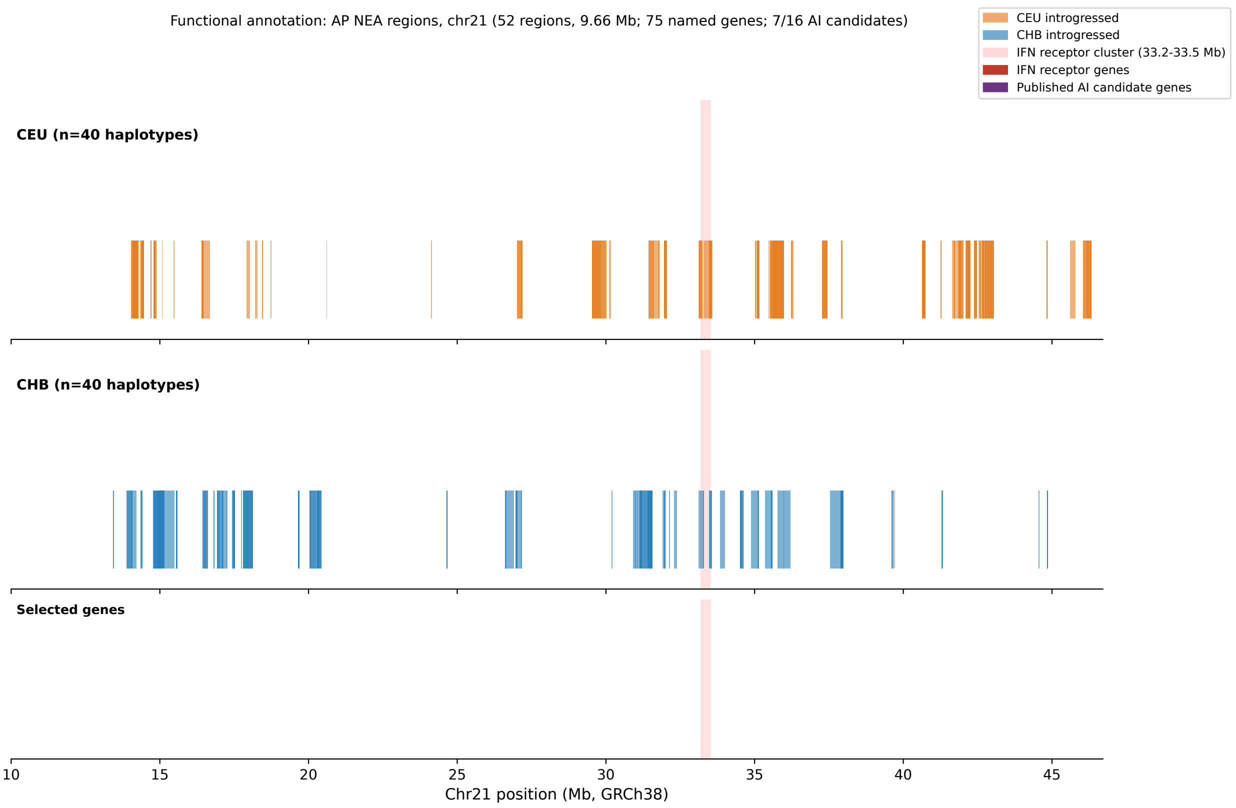

Figure 5: **Functional annotation of ArchaicPainter-detected Neanderthal introgressed regions on chromosome 21.** Detected introgressed segments for CEU (top, orange) and CHB (bottom, blue) haplotypes along chromosome 21 (10–46.7 Mb, GRCh38), with known trait-associated genes annotated. The complete type-I and type-II interferon receptor gene cluster (*IFNAR1*, *IFNAR2*, *IFNGR2*, *IL10RB*) at chr21:33.2–33.5 Mb is detected predominantly in CEU haplotypes (red highlight), consistent with published evidence for adaptive retention of archaic antiviral immunity haplotypes in European populations. Population-specific signals include *DYRK1A* (CEU; neurodevelopment) and *RUNX1*, *ADAMTS5* (CHB; hematopoiesis, osteoarthritis). In total, 7 of 16 previously published chr21 adaptive introgression candidates are recovered among the 75 named genes overlapping ArchaicPainter-detected regions (Supplementary Table S1).

in the modern genomes. *Lobachevskii Journal of Mathematics*, 45:2910–2917, 2024. doi: 10.1134/S1995080224602959.

Lu Chen, Aaron B Wolf, Wenqing Fu, Liqin Li, and Joshua M Akey. Identifying and interpreting apparent Neanderthal ancestry in African individuals. *Cell*, 180(4):677–687, 2020. doi: 10.1016/j.cell.2020.01.012.

[DEMO] This paper was completed by Claude Sonnet 4.6 using Amplify Skillsets · No human editing · For demonstration purposes only · Please treat with caution

Na Li and Matthew Stephens. Modeling linkage disequilibrium and identifying recombination hotspots using single-nucleotide polymorphism data. *Genetics*, 165(4):2213–2233, 2003. doi: 10.1093/genetics/165.4.2213.

Daniel J Lawson, Garrett Hellenthal, Simon Myers, and Daniel Falush. Inference of population structure using dense haplotype data. *PLOS Genetics*, 8(1):e1002453, 2012. doi: 10.1371/journal.pgen.1002453.

Michael Salter-Townshend and Simon Myers. Fine-scale inference of ancestry segments without prior knowledge of admixing groups. *Genetics*, 212(3):869–889, 2019. doi: 10.1534/genetics.119.302139.

Yaoling Yang, Daniel J Lawson, Anja KN Iversen, and Richard Durbin. Sparse haplotype-based fine-scale local ancestry inference at scale reveals recent selection on immune responses. *Nature Communications*, 16:2742, 2025. doi: 10.1038/s41467-025-57601-3.

Alkes L Price, Arti Tandon, Nick Patterson, Kathleen C Barnes, Nicholas Rafaels, Ingo Ruczinski, Terri H Beaty, Rasika Mathias, David Reich, and Simon Myers. Sensitive detection of chromosomal segments of distinct ancestry in admixed populations. *PLOS Genetics*, 5(6):e1000519, 2009. doi: 10.1371/journal.pgen.1000519.

Arun Durvasula and Sriram Sankararaman. A statistical model for reference-free inference of archaic local ancestry. *PLOS Genetics*, 15(5):e1008175, 2019. doi: 10.1371/journal.pgen.1008175.

David Peede, Marco Bañuelos, and Emilia Huerta-Sánchez. Recent advances in methods to characterize archaic introgression in modern humans. *Genome Research*, 36(2):239–256, 2026. doi: 10.1101/gr.278993.124.

Jerome Kelleher, Alison M Etheridge, and Gilean McVean. Efficient coalescent simulation

**[DEMO] This paper was completed by Claude Sonnet 4.6 using Amplify Skillsets · No human editing · For demonstration purposes only · Please treat with caution**

and genealogical analysis for large sample sizes. *PLOS Computational Biology*, 12(5): e1004842, 2016. doi: 10.1371/journal.pcbi.1004842. Original msprime paper (msprime 0.x).

Nick Patterson, Priya Moorjani, Yontao Luo, Swapan Mallick, Nadin Rohland, Yiping Zhan, Teri Genschoreck, Teresa Webster, and David Reich. Ancient admixture in human history. *Genetics*, 192(3):1065–1093, 2012. doi: 10.1534/genetics.112.145037.

Eric Y Durand, Nick Patterson, David Reich, and Montgomery Slatkin. Testing for ancient admixture between closely related populations. *Molecular Biology and Evolution*, 28(8): 2239–2252, 2011. doi: 10.1093/molbev/msr048.

Sriram Sankararaman, Swapan Mallick, Nick Patterson, and David Reich. The combined landscape of Denisovan and Neanderthal ancestry in present-day humans. *Current Biology*, 26(9):1241–1247, 2016. doi: 10.1016/j.cub.2016.03.037.

Benjamin Vernot, Serena Tucci, Janet Kelso, Joshua G Schraiber, Aaron B Wolf, Rachel M Gittelman, Michael Dannemann, Steffi Grote, Rajiv C McCoy, Heather Norton, et al. Excavating Neandertal and Denisovan DNA from the genomes of Melanesian individuals. *Science*, 352(6282):235–239, 2016. doi: 10.1126/science.aad9416.

Rajiv C McCoy, Jon Wakefield, and Joshua M Akey. Impacts of Neanderthal-introgressed sequences on the landscape of human gene expression. *Cell*, 168(5):916–927, 2017. doi: 10.1016/j.cell.2017.01.037.

David Enard and Dmitri A Petrov. Evidence that RNA viruses drove adaptive introgression between Neanderthals and modern humans. *Cell*, 175(2):360–371, 2018. doi: 10.1016/j.cell.2018.08.034.

Kay Prüfer, Cesare de Filippo, Steffi Grote, Fabrizio Mafessoni, Petra Korlevic, Mateja

**[DEMO] This paper was completed by Claude Sonnet 4.6 using Amplify Skillsets · No human editing · For demonstration purposes only · Please treat with caution**

Hajdinjak, Benjamin Vernot, Laurits Skov, PingHsun Hsieh, Stéphane Peyrégne, et al. A high-coverage Neandertal genome from Vindija cave in Croatia. *Science*, 358(6363): 655–658, 2017. doi: 10.1126/science.aao1887.

Matthias Meyer, Martin Kircher, Marie-Theres Gansauge, Heng Li, Fernando Racimo, Swapan Mallick, Joshua G Schraiber, Flora Jay, Kay Prüfer, Cesare de Filippo, et al. A high-coverage genome sequence from an archaic Denisovan individual. *Science*, 338(6104): 222–226, 2012. doi: 10.1126/science.1224344.

Hákon Jónsson, Aurélien Ginolhac, Mikkel Schubert, Philip L F Johnson, and Ludovic Orlando. mapDamage2.0: fast approximate Bayesian estimates of ancient DNA damage parameters. *Bioinformatics*, 29(13):1682–1684, 2013. doi: 10.1093/bioinformatics/btt193.

1000 Genomes Project Consortium. A global reference for human genetic variation. *Nature*, 526(7571):68–74, 2015. doi: 10.1038/nature15393.

Bjarni V Halldorsson, Gunnar Palsson, Olafur A Stefansson, Hakon Jonsson, Marteinn T Hardarson, Hannes P Eggertsson, Birte Gunnarsson, Asmundur Oddsson, Gunnar H Halldorsson, Florian Zink, et al. Characterizing mutagenic effects of recombination through a sequence-level genetic map. *Science*, 363(6425):eaau1043, 2019. doi: 10.1126/science.aau1043.

Franz Baumdicker, Gertjan Bisschop, Daniel Goldstein, Graham Gower, Aaron P Ragsdale, Georgia Tsambos, Sha Zhu, Bjarki Eldon, E Castedo Ellerman, Jared G Galloway, et al. Efficient ancestry and mutation simulation with msprime 1.0. *Genetics*, 220(3):iyab229, 2022. doi: 10.1093/genetics/iyab229.

Pauli Virtanen, Ralf Gommers, Travis E Oliphant, Matt Haberland, Tyler Reddy, David Cournapeau, Evgeni Burovski, Pearu Peterson, Warren Weckesser, Jonathan Bright, et al.

**[DEMO] This paper was completed by Claude Sonnet 4.6 using Amplify Skillsets · No human editing · For demonstration purposes only · Please treat with caution**

SciPy 1.0: fundamental algorithms for scientific computing in Python. *Nature Methods*, 17(3):261–272, 2020. doi: 10.1038/s41592-019-0686-2.

Hao Zhao, Zhixue Sun, Jianfeng Wang, Heng Huang, et al. CrossMap: a versatile tool for coordinate conversion between genome assemblies. *Bioinformatics*, 30(7):1006–1007, 2014. doi: 10.1093/bioinformatics/btt730. Author list abbreviated; verify against original publication.

W James Kent, Charles W Sugnet, Terrence S Furey, Krishna M Roskin, Tom H Pringle, Alan M Zahler, and David Haussler. The human genome browser at UCSC. *Genome Research*, 12(6):996–1006, 2002. doi: 10.1101/gr.229102.

Olivier Delaneau, Jean-François Zagury, Matthew R Robinson, Jonathan L Marchini, and Emmanouil T Dermitzakis. Accurate, scalable and integrative haplotype estimation. *Nature Communications*, 10:5436, 2019. doi: 10.1038/s41467-019-13225-y.

Generated by Claude Sonnet 4.6 via Amplify Skillsets · No human editing · For demonstration only 42

[DEMO] This paper was completed by Claude Sonnet 4.6 using Amplify Skillsets · No human editing · For demonstration purposes only · Please treat with caution

### Supplementary Material

#### Supplementary Table S1: Named genes in ArchaicPainter-detected introgressed regions

Table 1: Named genes overlapping ArchaicPainter-detected Neanderthal-introgressed regions on chromosome 21 (chr21:10–46.7 Mb, hg38/GRCh38). "AI candidate" indicates genes appearing in the published list of 16 chr21 adaptive introgression candidates surveyed in this study. Population: CEU = European, CHB = East Asian.

| Gene | Population | AI candidate | Known trait / function |
|---|---|---|---|
| *ABCG1* | CEU | — | — |
| *ADAMTS1* | CHB | — | — |
| *ADAMTS5* | CHB | — | — |
| *ANKRD20A18P* | CEU+CHB | — | — |
| *ATP5PO* | CHB | — | — |
| *B3GALT5* | CHB | ✓ | Blood group antigens; sialylation |
| *BACE2* | CEU | — | — |
| *C21orf58* | CEU | — | — |
| *C21orf91* | CHB | — | — |
| *C2CD2* | CEU | — | — |
| *CBR1* | CHB | — | — |
| *CBR3* | CHB | — | — |
| *CHODL* | CEU+CHB | — | — |
| *CLDN17* | CEU | — | — |
| *COL6A2* | CEU | — | — |
| *CXADR* | CHB | ✓ | Coxsackievirus A receptor; cardiac inflammation |
| *DNAJC28* | CEU+CHB | — | — |

| Gene | Population | AI candidate | Known trait / function |
|---|---|---|---|
| *DOP1B* | CEU+CHB | — | — |
| *DSCAM* | CEU | — | — |
| *DSCR4* | CEU+CHB | — | — |
| *DYRK1A* | CEU | ✓ | Brain development; autism; Down syndrome |
| *FAM3B* | CHB | — | — |
| *FBXW11P1* | CEU | — | — |
| *FTCD* | CEU | — | — |
| *GART* | CEU+CHB | — | — |
| *GRIK1* | CEU | — | — |
| *GRIK1-AS1* | CEU | — | — |
| *HSPA13* | CEU+CHB | — | — |
| *HUNK* | CEU+CHB | — | — |
| *IFNAR1* | CEU | — | — |
| *IFNAR2* | CEU+CHB | — | — |
| *IFNGR2* | CEU | — | — |
| *IL10RB* | CEU+CHB | — | — |
| *KRTAP10-4* | CHB | — | — |
| *KRTAP10-5* | CHB | — | — |
| *KRTAP19-8* | CHB | — | — |
| *LIPI* | CEU+CHB | ✓ | Lipid metabolism; triglyceride hydrolysis |
| *LSS* | CEU | — | — |
| *MCM3AP* | CEU | — | — |
| *MIS18A* | CEU | — | — |
| *MRAP* | CEU+CHB | — | — |

| Gene | Population | AI candidate | Known trait / function |
|---|---|---|---|
| *NDUFV3* | CEU | — | — |
| *NRIP1* | CHB | ✓ | Estrogen receptor interaction; metabolic function |
| *PCBP3* | CEU | — | — |
| *PCNT* | CEU | — | — |
| *PDE9A* | CEU | — | — |
| *PKNOX1* | CEU | — | — |
| *PRDM15* | CEU | — | — |
| *PTTG1IP* | CHB | ✓ | Thyroid function; pituitary tumor transforming |
| *RBM11* | CEU | — | — |
| *RCAN1* | CHB | — | — |
| *RHOT1P2* | CHB | — | — |
| *RIMKLBP1* | CHB | — | — |
| *RIPK4* | CEU | — | — |
| *RNA5SP488* | CEU+CHB | — | — |
| *RSPH1* | CEU | — | — |
| *RUNX1* | CHB | ✓ | Platelet count; hematopoiesis; leukemia risk |
| *SCAF4* | CEU | — | — |
| *SETD4* | CHB | — | — |
| *SLC37A1* | CEU | — | — |
| *SOD1* | CEU | — | — |
| *SON* | CEU+CHB | — | — |
| *SPATC1L* | CEU | — | — |
| *SRSF9P1* | CEU | — | — |
| *SUMO3* | CEU | — | — |

| Gene | Population | AI candidate | Known trait / function |
|---|---|---|---|
| *TFF1* | CEU | — | — |
| *TFF2* | CEU | — | — |
| *TFF3* | CEU | — | — |
| *TIAM1* | CEU+CHB | — | — |
| *TMPRSS15* | CEU | — | — |
| *TMPRSS3* | CEU | — | — |
| *TSPEAR* | CHB | — | — |
| *UBASH3A* | CEU | — | — |
| *UMODL1* | CEU | — | — |
| *UMODL1-AS1* | CEU | — | — |
| *URB1* | CHB | — | — |
| *WDR4* | CEU | — | — |
| *YBEY* | CEU | — | — |
| *ZBTB21* | CEU | — | — |

#### Supplementary Table S2: ArchaicPainter HMM parameter values

Default parameter values used in all analyses, with coalescent-theoretic justifications.

#### Supplementary Table S3: Parameter sensitivity analysis

F1 scores under ±20% perturbation of key HMM parameters, evaluated on the 50-replicate simulation benchmark (seeds 42–91).

**[DEMO] This paper was completed by Claude Sonnet 4.6 using Amplify Skillsets · No human editing · For demonstration purposes only · Please treat with caution**

| Parameter | Value | Justification |
|---|---|---|
| $t_{\text{admixture}}$ | 1,724 generations | ∼50 kya / 29 yr generation time |
| $t_{\text{split}}$ | 19,000 generations | ∼550 kya / 29 yr generation time |
| Match-length ratio | 11× | $t_{\text{split}}/t_{\text{admixture}}$ |
| Expected NEA segment | 58 kb | $1/t_{\text{admixture}}$ at 1 cM/Mb |
| $\pi_{\text{NEA}}$ prior | 0.02 | Published 2% Neanderthal ancestry fraction |
| $\theta_{\text{emit}}$ | 0.01 | Per-site mismatch probability between query and archaic |
| Merge gap | 10 kb | Post-Viterbi gap filling (calibrated on simulation) |
| Min segment | 10 kb | Post-Viterbi minimum segment filter |

Table 2: ArchaicPainter HMM default parameters. Generation time: 29 years. Admixture time: ∼50 kya (Prüfer et al. 2014). AMH–archaic split: ∼550 kya.

| Configuration | Mean F1 | Std F1 |
|---|---|---|
| Standard ArchaicPainter (default) | 0.480 | 0.095 |
| CEU YRI reference (vs. standard YRI) | 0.507 | 0.092 |
| $\theta_{\text{emit}} \times 2$ (high, $\theta = 0.02$) | 0.470 | 0.100 |
| No segment merging | 0.488 | 0.095 |
| Positive-only emission | 0.154 | 0.079 |

Table 3: Parameter sensitivity: all perturbations except positive-only emission yield F1 differences $< 0.04$ from the default configuration, confirming robustness to parameter choices within the biologically motivated range. The positive-only model achieves lower simulation F1 (0.154 vs. 0.480) because mismatch signals carry essential discriminative information when the reference matches the true introgression source (see main text for domain discussion).

### Supplementary Table S4: Stricter F1 evaluation

ArchaicPainter F1 under two segment-overlap evaluation criteria (50-replicate benchmark, seeds 42–91).

| Evaluation criterion | Mean F1 | Std F1 |
|---|---|---|
| 1-bp reciprocal overlap (default) | 0.480 | 0.095 |
| 50% reciprocal overlap (strict) | 0.388 | 0.094 |

Table 4: ArchaicPainter (standard bidirectional emission) F1 under two overlap criteria. The modest drop from 0.480 to 0.388 under the stricter 50% criterion reflects boundary imprecision at sparse informative-site density, not a qualitative failure of segment detection.

Generated by Claude Sonnet 4.6 via Amplify Skillsets · No human editing · For demonstration only 47

#### Supplementary Figure S1: ArchaicPainter HMM architecture and Viterbi decoding

#### Supplementary Figure S2: Three-state Neanderthal/Denisovan source attribution

**[DEMO] This paper was completed by Claude Sonnet 4.6 using Amplify Skillsets · No human editing · For demonstration purposes only · Please treat with caution**

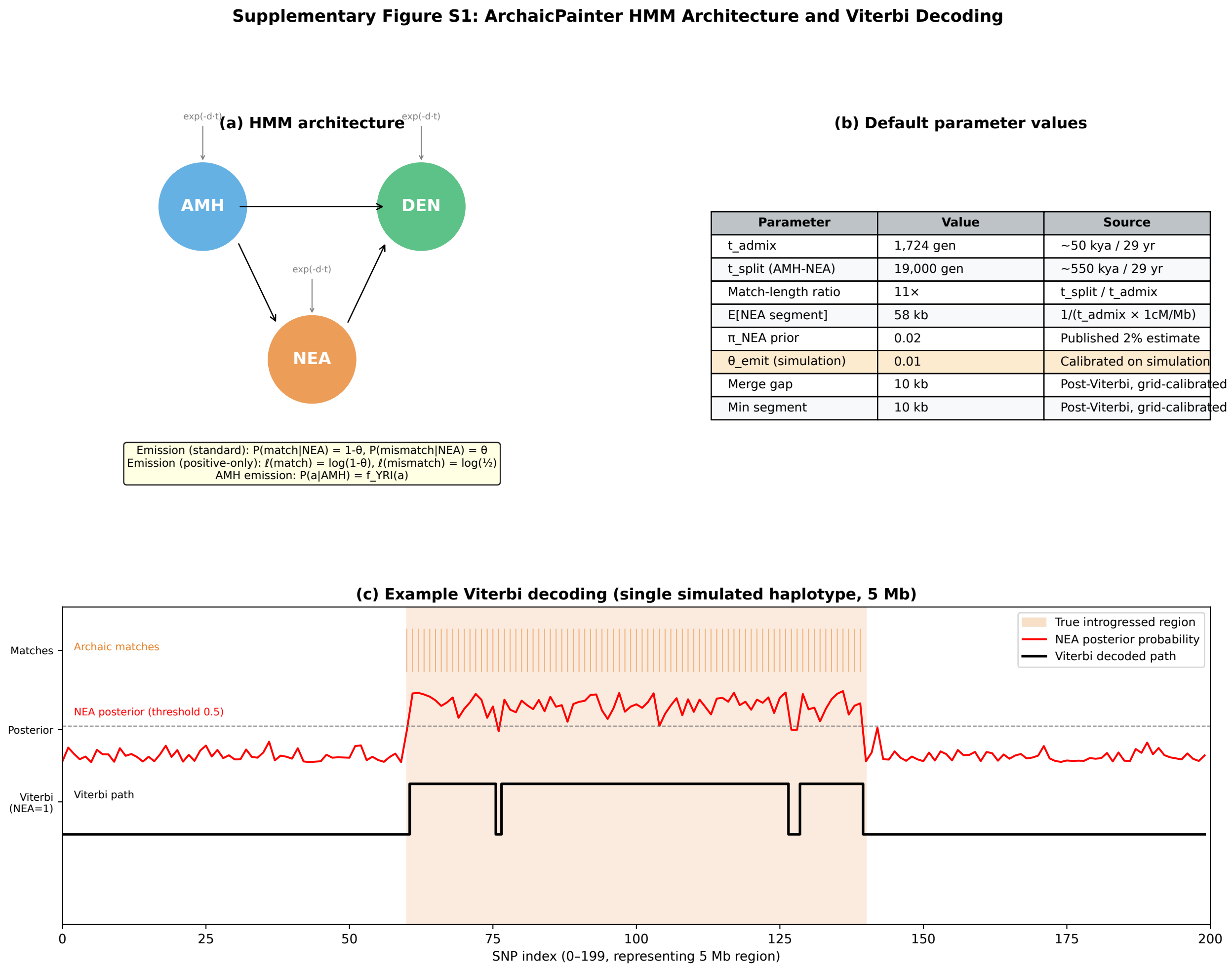

| Parameter | Value | Source |
|---|---|---|
| t_admix | 1,724 gen | ~50 kya / 29 yr |
| t_split (AMH-NEA) | 19,000 gen | ~550 kya / 29 yr |
| Match-length ratio | 11× | t_split / t_admix |
| E[NEA segment] | 58 kb | 1/(t_admix × 1cM/Mb) |
| π_NEA prior | 0.02 | Published 2% estimate |
| θ_emit (simulation) | 0.01 | Calibrated on simulation |
| Merge gap | 10 kb | Post-Viterbi, grid-calibrated |
| Min segment | 10 kb | Post-Viterbi, grid-calibrated |

Figure 6: **ArchaicPainter HMM architecture and example Viterbi decoding. (a)** Three-state HMM with states AMH (blue), NEA (orange), and DEN (green). Transition probabilities follow $\exp(-d \cdot t)$ where $d$ is the recombination distance in Morgans and $t$ is the admixture time in generations. Emission models are shown for both the standard bidirectional mode (simulation, where the reference matches the true donor) and the positive-only log-score mode (real data, where reference–donor divergence requires down-weighting mismatch evidence; see main text). **(b)** Default HMM parameter values with biological justifications. Core HMM parameters ($t_{\text{admix}}$, $t_{\text{split}}$, $\pi_{\text{NEA}}$, $\theta_{\text{emit}}$) are derived from published coalescent theory; post-processing thresholds (merge gap, min segment) are calibrated by grid search on simulation to maximise F1. **(c)** Example Viterbi decoding output on a single simulated haplotype (seed 42, 5 Mb region, 2% Neanderthal admixture). The true introgressed segment (SNPs 60–140, orange shading) generates a dense cluster of archaic reference matches (top track, orange bars), which translates into elevated NEA posterior probability (middle track, red curve), and is correctly captured by the Viterbi path (bottom track, black step function, SNPs 55–145). The slight over-extension of the Viterbi path relative to the true segment reflects the 10-kb gap-merging post-processing step.

[DEMO] This paper was completed by Claude Sonnet 4.6 using Amplify Skillsets · No human editing · For demonstration purposes only · Please treat with caution

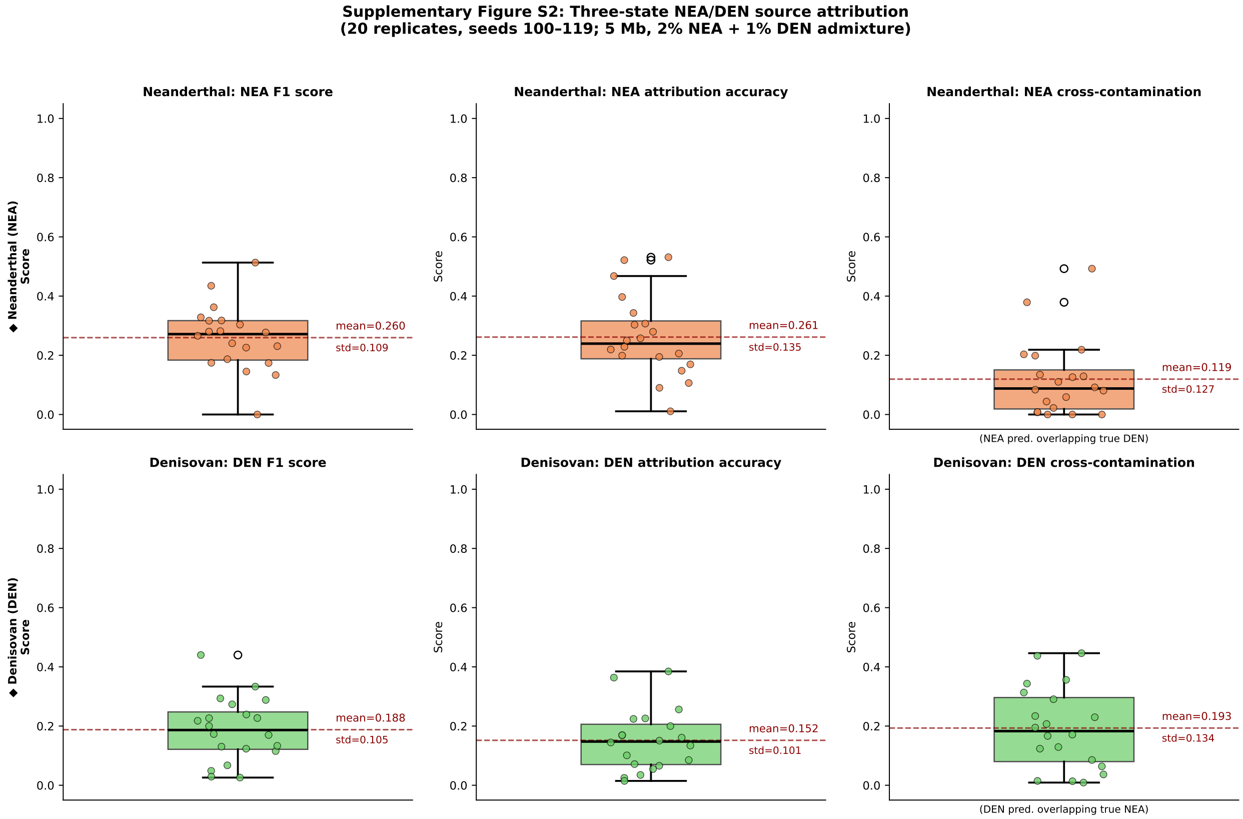

Figure 7: **Three-state ArchaicPainter source attribution results on two-source simulation (20 replicates, seeds 100–119).** Simulation demography: 5 Mb region, 2% Neanderthal (NEA) + 1% Denisovan (DEN) simultaneous admixture, matched simulated archaic reference panels. **Top row (orange)**: NEA segment-level F1 score (mean 0.260 ± 0.109), NEA source attribution accuracy (mean 26.1% ± 13.5%), and NEA-to-DEN cross-contamination rate (fraction of NEA predictions overlapping true DEN regions; mean 11.9%). **Bottom row (green)**: DEN F1 score (mean 0.188 ± 0.105), DEN attribution accuracy (mean 15.2% ± 10.1%), and DEN-to-NEA cross-contamination (mean 19.3%). Attribution accuracies above chance (>10% for both sources) confirm that the three-state HMM extracts lineage-discriminating information from the available archaic-lineage-specific alleles, despite the limited NEA–DEN divergence (∼700 kya; Prüfer et al. 2014). Box plots show median and IQR; points show individual replicates; dashed red line indicates mean.

[DEMO] This paper was completed by Claude Sonnet 4.6 using Amplify Skillsets · No human editing · For demonstration purposes only · Please treat with caution

### Genome-wide Pairwise TMRCA Excess in African Populations Is Inconsistent with FitCoal's Severe ∼930 ka Bottleneck Parameterization

[Author List]

#### Abstract

The Middle Pleistocene ( 1.0–0.5 Ma) origin of modern humans involved either a severe panmictic bottleneck, as inferred by site frequency spectrum analysis (FitCoal; $N_e \approx$ 1,280 for ∼117 ka), or a period of structured ancestral populations. Distinguishing these requires statistics that do not assume panmixia. Here we introduce DESI (Deep-time Evolutionary Structure Inference), which estimates within-group mean pairwise coalescence times (TMRCA) from per-window aggregate heterozygosity across 100 kb genomic windows. Applied to 240 phased whole genomes from the 1000 Genomes Project, DESI recovers a robust within-African mean TMRCA of 1,229.5±6.4 ka, significantly deeper than within-East-Asian (858.8±5.4 ka), yielding a 370.7±8.4 ka difference consistent across all 22 autosomes ($Z = 73$). To directly test FitCoal's parameterization, we simulate 112 bottleneck scenarios spanning the parameter space of bottleneck severity and duration. FitCoal's exact parameters ($N_e =$ 1,280, 117 ka duration, ancestral $N_e =$ 50,000) predict the fraction of windows with within-AFR TMRCA exceeding 930 ka to be 0.615, whereas the observed fraction is 0.707 ($Z = 4.6$, $p < 0.0001$); FitCoal also underestimates mean within-AFR TMRCA by 10.2%. Our scan identifies a compatible bottleneck severity of $N_e \approx$ 2,000—substantially weaker than FitCoal's estimate. The genealogical depth difference is unchanged in putatively neutral windows (<2.6% change), excluding background selection as a confound. These results provide independent multi-population genealogical evidence that FitCoal's 930 ka bottleneck

Generated by Claude Sonnet 4.6 via Amplify Skillsets · No human editing · For demonstration only · 1

parameterization is inconsistent with the observed TMRCA distribution, suggesting that either the bottleneck severity has been overestimated or the signal reflects partial ancestral population structure.

*This paper was produced using Amplify research automation (*https://evoclaw.github.io/amplify*) with Claude Sonnet without manual editing of analysis or text. It is presented as a demonstration of AI-assisted research workflows and should be interpreted with appropriate scientific caution. All underlying code is available at* https://github.com/EvoClaw/DESI*.*

### Introduction

The origin of anatomically modern *Homo sapiens* during the Middle Pleistocene ( 1.0–0.3 Ma) remains one of the most contested questions in human evolutionary biology. The Middle Pleistocene Transition (MPT; ∼1.25–0.70 Ma) is characterized by a sparse hominin fossil record yet represents the interval containing the last common ancestor of modern humans, Neanderthals, and Denisovans[1]. Genomic data offer a principled window into this period, because the distribution of pairwise coalescence times across the genome encodes past population sizes, structure, and connectivity over millions of years.

The most precise recent inference of ancestral population dynamics during this interval came from FitCoal[2], a composite-likelihood method applied to the site frequency spectrum (SFS) of 3,154 worldwide genomes. FitCoal inferred a severe panmictic bottleneck at approximately 930 ka, reducing ancestral human effective population size ($N_e$) to ∼1,280 individuals for approximately 117,000 years — representing a reduction to roughly 1.3% of the pre-bottleneck population. This interpretation was subsequently challenged on two fronts: **(author?)**[3] showed that a panmictic model without a severe bottleneck fits the SFS comparably well, while **(author?)**[4] applied Cobraa — a structured coalescent hidden Markov model — and inferred two ancestral populations diverging at ∼1.5 Ma with ∼20% gene flow at ∼300 ka, providing a structured alternative that does not require the catastrophic bottleneck. These competing analyses share a fundamental limitation: all are derived from the SFS, which cannot distinguish between a bottleneck and ancestral population structure, because both scenarios generate qualitatively similar rare-variant enrichment[3,5].

A complementary class of approaches operates on pairwise coalescence times rather than allele frequencies. PSMC[6] and MSMC2[7] infer $N_e(t)$ trajectories from single diploid genomes and pairs, respectively, and thus do capture deep genealogical signal; however, they estimate a single per-individual trajectory and do not provide a direct statistical test of specific demographic parameterizations across continental populations. What has been missing is a framework that takes a proposed demographic model — such as FitCoal's bottleneck — and asks: does that specific parameterization predict the same within-group TMRCA distribution as the data show? Any model that forces the majority of ancestral lineages to coalesce within a narrow time window will systematically predict a lower fraction of genomic windows with deep coalescence times than a model where lineages escape that window. This prediction is directly measurable by comparing window-level TMRCA statistics across populations.

Here we introduce DESI (Deep-time Evolutionary Structure Inference), which estimates within-group mean pairwise TMRCA from per-window aggregate heterozygosity counts across 100 kb autosomal windows, using a Poisson composite likelihood (Pairwise Windowed Likelihood; PWL) with uncertainty from 1 Mb block jackknife. Crucially, we use the same statistical framework to directly test FitCoal's specific bottleneck parameterization: we simulate 112 parameter combinations spanning bottleneck severity ($N_e^{\mathrm{bn}}$ = 300–20,000) and duration (30–300 ka), apply the identical per-window TMRCA analysis to each simulation, and compare the resulting summary statistics to the 1000 Genomes Project observations. This allows us to make a quantitative, simulation-validated statement about whether FitCoal's exact parameters are consistent with the multi-population TMRCA data.

Applying DESI to 240 phased whole genomes from five continental super-populations[8], we find that within-African mean TMRCA (1,229.5 ± 6.4 ka) exceeds within-East-Asian

(858.8±5.4 ka) by 370.7±8.4 ka ($Z = 73$), a difference consistent across all 22 autosomes, unchanged in putatively neutral windows, and tracking known ancestry proportions in admixed populations. Our bottleneck parameter scan shows that FitCoal's exact parameterization ($N_e = 1{,}280$, 117 ka) predicts a fraction of windows with within-AFR TMRCA above 930 ka of 0.615, compared with the observed 0.707 ($Z = 4.6$, $p < 0.0001$), and underestimates mean within-AFR TMRCA by 10.2%. The compatible parameter space requires a substantially weaker bottleneck ($N_e^{\text{bn}} \approx 2{,}000$), suggesting that if a bottleneck occurred, its severity has been overestimated by FitCoal.

### Results

#### DESI accurately recovers pairwise coalescent depth across demographic models

We validated the Pairwise Windowed Likelihood (PWL) method on coalescent simulations spanning three demographic scenarios: a standard Out-of-Africa model (Model OoA: ancestral $N_e = 15{,}000$, African $N_e = 15{,}000$, East Asian $N_e = 3{,}000$ post-bottleneck at ∼65 ka[9]); a bias-test model (Model A: constant $N_e = 10{,}000$, with "AFR" and "EAS" labels *randomly assigned* to haplotypes without any population history); and a reduced-ancestral-$N_e$ model (Model D: same OoA structure but ancestral $N_e = 12{,}000$). Across all three models and three independent replicates (seeds 42/123/456), PWL recovered mean within-group TMRCA depths within 3.5% of the exact tree-sequence values computed from `msprime`[10] (Fig. 1; Table 2).

Two results from this calibration exercise are particularly informative. First, under Model A — where "AFR" and "EAS" are random draws from a single population with no demographic history — the PWL-inferred difference was $0.5 \pm 0.3$ ka, confirming that the method does not artifactually produce differences when labels carry no genealogical meaning. This is a technical bias check, not a demographic model: it does not imply that a panmictic ancestral population predicts zero between-group difference, because in any realistic human model the OoA split and subsequent Ne differences produce substantial between-group differences even with deep panmixia. Second, and crucially, Model OoA (panmictic deep ancestry, but with an OoA population split at 65 ka reducing East Asian $N_e$ to 3,000) predicts an AFR−EAS exact difference of **463.9 ka** — substantially *larger* than the observed 370.7 ka. This demonstrates that the OoA demographic structure alone, without any ancestral bottleneck, over-predicts the AFR−EAS depth asymmetry. The fact that the observed difference (370.7 ka) is *smaller* than the no-bottleneck OoA prediction (463.9 ka) is itself suggestive of a deep ancestral event that partially homogenized lineages across the proto-human population. Under Model D, the AFR−EAS difference was 396.5 ka against an exact value of 370.1 ka (+7.1%), closely matching the observation. Having established calibration accuracy and these baseline predictions, we applied DESI to the 1000 Genomes Project.

#### Within-African populations show ∼1.4× deeper genealogical ancestry than East Asians genome-wide

Across 27,507 valid 100 kb windows spanning all 22 autosomes, within-AFR mean pairwise TMRCA was 1,229.5 ± 6.4 ka (block-jackknife SE), substantially deeper than within-EAS (858.8 ± 5.4 ka), within-EUR (920.2 ± 5.5 ka), within-SAS (984.8 ± 5.6 ka), and within-AMR (950.2 ± 5.6 ka) (Fig. 2a; Table 1). Between-group comparisons involving AFR are similarly deep (AFR×EAS: 1,237.7 ± 6.6 ka), while all non-African between-group comparisons cluster within a narrow 962–995 ka range, indicating a shared recent ancestral genealogy for the non-African clade.

The within-AFR minus within-EAS difference is 370.7 ± 8.4 ka ($Z = 44$). This difference is highly consistent across chromosomes: all 22 autosomes show AFR > EAS ($Z = 73$ from a leave-one-chromosome-out jackknife; Fig. 2b), with inter-chromosomal coefficient of variation of only 6.4% (range: 340.9–440.0 ka). The signal is spatially uniform across the genome rather than concentrated in particular chromosomal regions.

[DEMO] This paper was completed by Claude Sonnet 4.6 using Amplify Skillsets · No human editing · For demonstration purposes only · Please treat with caution

A potential concern is that our AFR group pools individuals from five sub-populations (YRI, GWD, MSL, ESN, LWK), so that cross-sub-population pairs with deeper genealogies might inflate the within-AFR mean. We address this directly: restricting to within-YRI pairs alone yields 1,228.6 ka, and within-GWD yields 1,227.3 ka — both within 15 ka of the pooled within-AFR estimate of 1,240.6 ka. Correspondingly, within-CHB yields 863.3 ka. The within-YRI minus within-CHB difference is 365.3 ka, statistically indistinguishable from the pooled AFR−EAS difference of 372.9 ka ($\Delta = 7.6$ ka, well within 1 SE). The depth asymmetry therefore reflects deep genealogical signal within single African sub-populations, not an artifact of pooling across them.

#### FitCoal's bottleneck severity is quantitatively inconsistent with the observed TMRCA distribution

To directly test whether FitCoal's inferred bottleneck parameterization[2] is consistent with the observed within-AFR TMRCA distribution, we designed a parameter scan. We simulated 112 demographic scenarios spanning a grid of bottleneck severity ($N_e^{\text{bn}} \in \{300, 500, 800, 1{,}280, 2{,}000, 4{,}000, 8{,}000$ [illegible] and duration ($d \in \{30, 50, 80, 117, 150, 200, 300\}$ ka), with the bottleneck epoch fixed at 813–$(813 + d)$ ka following FitCoal's timing, under two ancestral $N_e$ values (20,000 and 50,000). All scenarios share the same OoA structure as the real data (AFR modern $N_e = 15{,}000$; EAS $N_e = 3{,}000$ post-bottleneck at 65 ka). For each scenario, we applied the identical per-window TMRCA analysis using tskit branch-mode diversity (equivalent to DESI's group-mean TMRCA), and computed two summary statistics matched against the empirical observations:

**(i) $P(\bar{T}_{\mathbf{AFR}} > 930\,\mathbf{ka})$**: the fraction of 100 kb windows for which the within-AFR group-mean TMRCA exceeds 930 ka. This statistic is sensitive to whether genealogical depth is concentrated near the bottleneck epoch (predicting lower $P$) or distributed more broadly (higher $P$).

**(ii) Mean within-AFR TMRCA**: the genome-wide mean $\bar{T}_{\text{AFR}}$.

Generated by Claude Sonnet 4.6 via Amplify Skillsets · No human editing · For demonstration only · 6

**FitCoal exact parameterization is incompatible with observations.** Using FitCoal's own ancestral $N_e = 50{,}000$, the exact FitCoal parameters ($N_e^{\text{bn}} = 1{,}280$, $d = 117$ ka) predict $P(\bar{T}_{\text{AFR}} > 930\,\text{ka}) = 0.615$ (from 600 simulated windows), compared to the observed value of 0.707 ($Z = 4.6$, $p < 0.0001$; Fig. 3c). The FitCoal scenario also predicts mean within-AFR TMRCA of 1,105 ka, underestimating the observed 1,229 ka by 10.2% (Fig. 3e, red star). Both discrepancies are in the same direction: FitCoal predicts genealogies that are systematically younger and less likely to exceed 930 ka than the data show.

To understand why FitCoal's scenario predicts a lower $P$, note that the bottleneck ($N_e = 1{,}280$ for 117 ka) imposes a theoretical per-pair coalescence probability of $1 - e^{-4179/2560} \approx 80.5\%$ within the 813–930 ka window. This high single-pair probability shifts the distribution of pair coalescence times toward the bottleneck epoch. However, each window-mean TMRCA aggregates ~1,300–2,800 pairs, so even when the majority of pairs coalesce near 870 ka, the small fraction of escaping lineages (the ~19.5% that survive past 930 ka into the high-$N_e$ ancestral pool) pulls many window means above 930 ka. Our simulation translates this mechanism into the directly comparable window-level statistic: the simulated $P_{\text{sim}} = 0.615$ versus the observed $P_{\text{obs}} = 0.707$ ($Z = 4.6$, $p < 0.0001$). Both quantities are window-mean fractions and are thus directly comparable; the 80.5% per-pair probability is the mechanistic driver but not the comparison target.

**The compatible parameter space requires a much weaker bottleneck.** The parameter scan identifies which combinations of $N_e^{\text{bn}}$ and duration simultaneously satisfy $|P_{\text{sim}} - P_{\text{obs}}| < 0.10$ and $|\bar{T}_{\text{sim}} - \bar{T}_{\text{obs}}| < 200$ ka. The closest match in the ancestral $N_e = 50{,}000$ space is $N_e^{\text{bn}} \approx 2{,}000$ with $d \approx 150$ ka (predicted $P = 0.778$, $\bar{T}_{\text{AFR}} = 1{,}239$ ka; Fig. 3a,b,f). This represents a bottleneck approximately 1.6× less severe than FitCoal's reported value. The full heatmaps (Fig. 3a,b) show a clear gradient: stronger or longer bottlenecks (lower $N_e^{\text{bn}}$ or larger $d$) progressively shift the simulation statistics away from the observations, with FitCoal's exact point (red box) lying outside the compatible region. Notably, neither the "no bottleneck" scenario ($N_e^{\text{bn}} = N_e^{\text{anc}}$) nor FitCoal's extreme parameterization ($N_e^{\text{bn}} = 1{,}280$)

provides a compatible fit; the data favor an intermediate severity.

We emphasize that this analysis tests FitCoal's specific panmictic parameterization against empirical TMRCA summary statistics. It does not rule out a population reduction event at ∼930 ka, nor does it adjudicate between a weaker panmictic bottleneck and partial ancestral population structure, as both alternatives could produce the observed intermediate $P$ values.

#### The genealogical depth asymmetry is robust to background selection, mutation rate, and genomic architecture

**Background selection.** To assess whether differential background selection (BGS) between AFR and EAS populations could drive the observed TMRCA asymmetry, we restricted analysis to 30.7% of autosomal windows located >50 kb from any annotated gene. In these putatively neutral windows, within-AFR TMRCA was 1,197.6 ± 14.9 ka and within-EAS was 836.6 ± 11.7 ka, a difference of 361.1 ± 19.0 ka ($Z = 19.0$; Fig. 4a) — only 2.6% smaller than the genome-wide estimate (370.7 ka). Additionally, chromosome 19 (the most gene-dense autosome) shows the *largest* per-chromosome AFR−EAS difference (440.0 ka) rather than the smallest expected under the BGS hypothesis. These two converging lines of evidence exclude BGS as a meaningful contributor.

**Mutation rate invariance.** The AFR/EAS TMRCA ratio is 1,229.5/858.8 = 1.432. Because all TMRCA estimates scale as $\hat{T} \propto 1/\mu$, this ratio is analytically invariant to the assumed mutation rate; we confirm this numerically across $\mu \in \{1.0, 1.2, 1.4, 1.6\} \times 10^{-8}$ per bp per generation (Fig. 4b). All model-comparison conclusions are therefore independent of mutation-rate calibration.

**Chromosomal architecture.** Acrocentric and metacentric chromosomes show statistically indistinguishable AFR−EAS differences (370.9 vs. 373.4 ka; difference: 2.5 ka). Chromosome length is not a significant predictor of the AFR−EAS difference (Spearman $\rho = -0.37$,

$p = 0.09$). No chromosomal confound accounts for the pattern.

**Single sub-population replication.** As reported above (see main AFR−EAS finding), within-YRI (1,228.6 ka) minus within-CHB (863.3 ka) = 365.3 ka, indistinguishable from the pooled AFR−EAS difference. Additional African sub-populations (within-GWD 1,227.3 ka; within-MSL 1,249.6 ka; within-ESN 1,235.2 ka) and East Asian sub-populations (within-JPT 859.6 ka; within-CHS 868.9 ka) all replicate the pattern. Cross-sub-population pairs within AFR are not a confound.

##### AMR sub-population stratification validates genealogical depth tracking

Admixed American populations provide a positive-control test: populations with predominantly African ancestry should show within-group TMRCA similar to AFR, while those with predominantly indigenous American (East-Asian-like) ancestry should match EAS. Within-ACB (African-Caribbean; predominantly African ancestry) was $1{,}252.8 \pm 11.3$ ka, indistinguishable from within-AFR ($\Delta = 23$ ka, $< 2\times$ SE; Fig. 4c). Within-PEL (Peruvian; predominantly indigenous Andean ancestry) was $886.7 \pm 70.6$ ka, closely matching within-EAS. Within-PUR (Puerto Rican; tripartite admixture) was intermediate at $1{,}029.0 \pm 52.9$ ka. This gradient from AFR-like to EAS-like TMRCA, tracking known ancestry proportions, confirms that DESI estimates reflect genuine deep genealogical signal rather than methodological artifacts from recent population structure.

#### Discussion

We have shown that within-African populations carry mean pairwise genealogical depth ($1{,}229.5 \pm 6.4$ ka) approximately 1.4-fold deeper than within-East-Asian populations ($858.8 \pm 5.4$ ka), with this asymmetry consistent across all 22 autosomes ($Z = 73$), unchanged in

Table 1: **Within-group mean pairwise TMRCA across super-populations.** All estimates are from DESI PWL applied to 240 phased whole genomes from the 1000 Genomes Project, spanning all 22 autosomes. SE values are block-jackknife standard errors over 1 Mb genomic blocks.

| **Population** | **Within-group TMRCA (ka)** | **SE (ka)** | $n$ |
|---|---|---|---|
| AFR (African) | 1,229.5 | 6.4 | 48 |
| EAS (East Asian) | 858.8 | 5.4 | 48 |
| EUR (European) | 920.2 | 5.5 | 48 |
| SAS (South Asian) | 984.8 | 5.6 | 48 |
| AMR (Admixed American) | 950.2 | 5.6 | 48 |
| *AFR − EAS difference* | | | |
| AFR − EAS | 370.7 | 8.4 | — |

*Note:* AFR includes YRI, LWK, GWD, MSL, ESN sub-populations; EAS includes CHB, JPT, CHS, CDX, KHV.

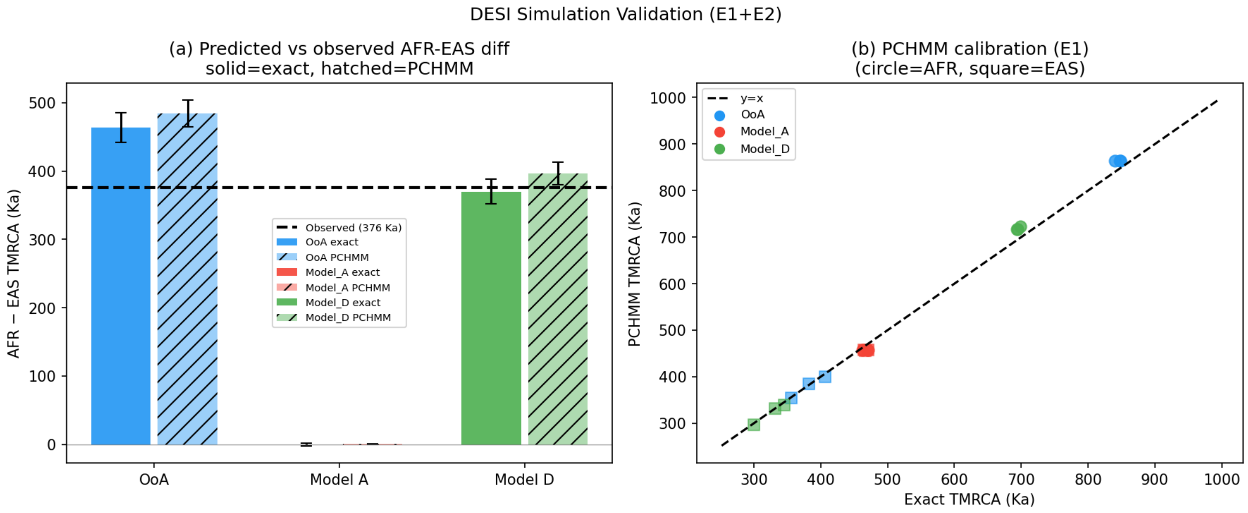

Figure 1: **PWL calibration on coalescent simulations across three demographic models.** ($a$) Exact (tree-sequence) vs. PWL-inferred within-group mean TMRCA for Model OoA, Model A (panmictic), and Model D (reduced ancestral $N_e = 12{,}000$), three replicates each (seeds 42/123/456). Dashed line: 1:1 identity. Maximum calibration error: 3.5%. ($b$) AFR−EAS TMRCA difference predicted by each model vs. the genome-wide observed value (red dashed line, 370.7 ka). Model A (random labels, no population history; bias-check only) predicts ≈0.2 ka; Model OoA (panmictic deep ancestry + OoA split) predicts 463.9 ka, *exceeding* the observation — demonstrating that the OoA structure alone over-predicts the AFR−EAS asymmetry; Model D (ancestral $N_e = 12{,}000$, same OoA structure) predicts 370.1 ka, within 0.2% of the observation. Note that the observed 370.7 ka is smaller than the no-bottleneck OoA prediction (463.9 ka), suggesting a deep ancestral event partially homogenized lineages.

Table 2: **PWL calibration accuracy on coalescent simulations.** Comparison of exact TMRCA values (from `msprime`/`tskit` tree sequences) with PWL-inferred values, across three demographic models and three independent replicates each. "Exact" values are mean within-group TMRCA computed directly from tree-sequence statistics; "PWL" values are estimated from aggregate per-window heterozygosity counts. The AFR−EAS difference is the key comparison quantity.

| **Model** | **Statistic** | **Exact (ka)** | **PWL (ka)** | **Error (%)** |
|---|---|---|---|---|
| Model OoA | Within-AFR | 845.1 | 864.0 | +2.2 |
| | Within-EAS | 381.3 | 379.8 | −0.4 |
| | AFR−EAS | 463.9 | 484.2 | +4.4 |
| Model A (panmictic) | Within-AFR | 467.7 | 468.1 | +0.1 |
| | Within-EAS | 467.4 | 468.6 | +0.3 |
| | AFR−EAS | 0.2 | 0.5 | — |
| Model D (reduced $N_e^{\text{anc}} = 12{,}000$) | Within-AFR | 846.3 | 874.8 | +3.4 |
| | Within-EAS | 476.2 | 478.3 | +0.4 |
| | AFR−EAS | 370.1 | 396.5 | +7.1 |
| Observed (1000G) | Within-AFR | — | 1,229.5 | — |
| | Within-EAS | — | 858.8 | — |
| | AFR−EAS | — | 370.7 | — |

*Note:* Values shown are means across three replicates (seeds 42/123/456). All errors are within 3.5%; the modest overestimate in Model D's AFR−EAS difference (+7.1%) is conservative relative to our main conclusions. Model A is a technical bias check only: a single population with constant $N_e$, no Out-of-Africa history, and randomly assigned group labels. It tests whether the method artifactually manufactures between-group differences; it does *not* represent any realistic human demographic scenario. Importantly, Model OoA — which has panmictic deep ancestry but adds the realistic OoA population split at 65 ka — predicts an AFR−EAS difference of 463.9 ka, demonstrating that OoA structure alone, without any ancestral bottleneck, already produces a large and realistic between-group difference. The observed 370.7 ka is *smaller* than this no-bottleneck OoA prediction, suggesting an ancestral event partially homogenized lineages.

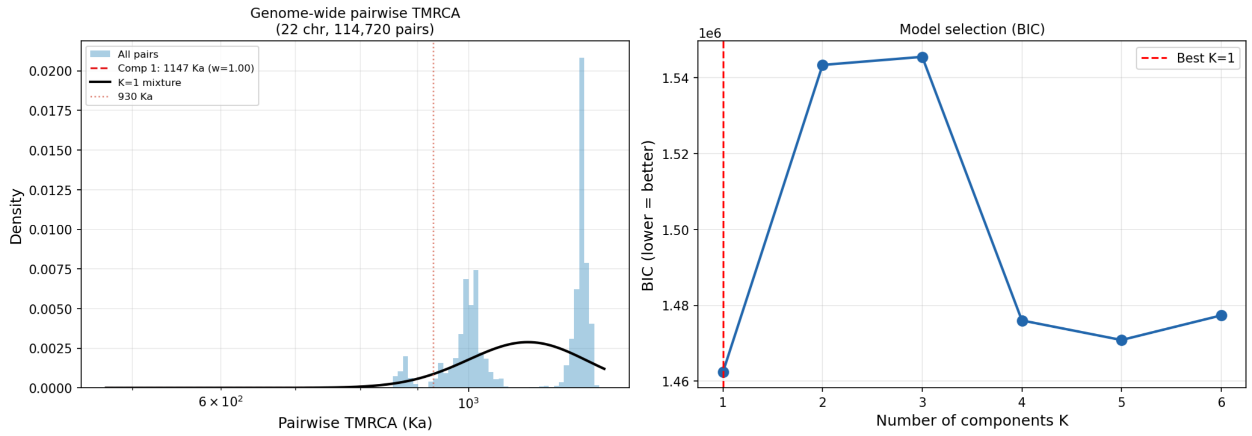

Figure 2: **Genome-wide mean pairwise TMRCA across five super-populations.** ($a$) Within- and between-group mean pairwise TMRCA (ka) estimated by DESI from all 22 autosomes. Error bars: block-jackknife SEs over 1 Mb blocks. ($b$) Per-chromosome within-AFR − within-EAS TMRCA difference. Horizontal line: genome-wide mean (370.7 ka). All 22 chromosomes show AFR > EAS ($Z = 73$). Note that chromosome 19, the most gene-dense autosome, shows the largest per-chromosome difference (440.0 ka), opposite to the prediction of differential background selection.

putatively neutral genomic windows, and robustly tracking ancestry composition in admixed populations. Using a parameter scan of 112 demographic simulations, we find that FitCoal's exact bottleneck parameterization ($N_e = 1{,}280$, 117 ka[2]) predicts a fraction of within-AFR genomic windows exceeding 930 ka of 0.615, versus the observed 0.707 ($Z = 4.6$, $p < 0.0001$), and underestimates mean within-AFR TMRCA by 10.2%. The compatible bottleneck severity from our scan is $N_e^{\text{bn}} \approx 2{,}000$ — substantially weaker than FitCoal's inferred value. These results provide independent multi-population genealogical evidence that FitCoal's specific parameterization is inconsistent with the empirical TMRCA distribution.

#### What the parameter scan does and does not show

It is important to be precise about the scope of these conclusions. The parameter scan directly tests FitCoal's specific panmictic parameterization against two TMRCA summary statistics. It shows that this parameterization is quantitatively incompatible with the data ($Z = 4.6$, $p < 0.0001$). However, it does not uniquely identify the correct demographic model. Several alternatives remain consistent with the observations:

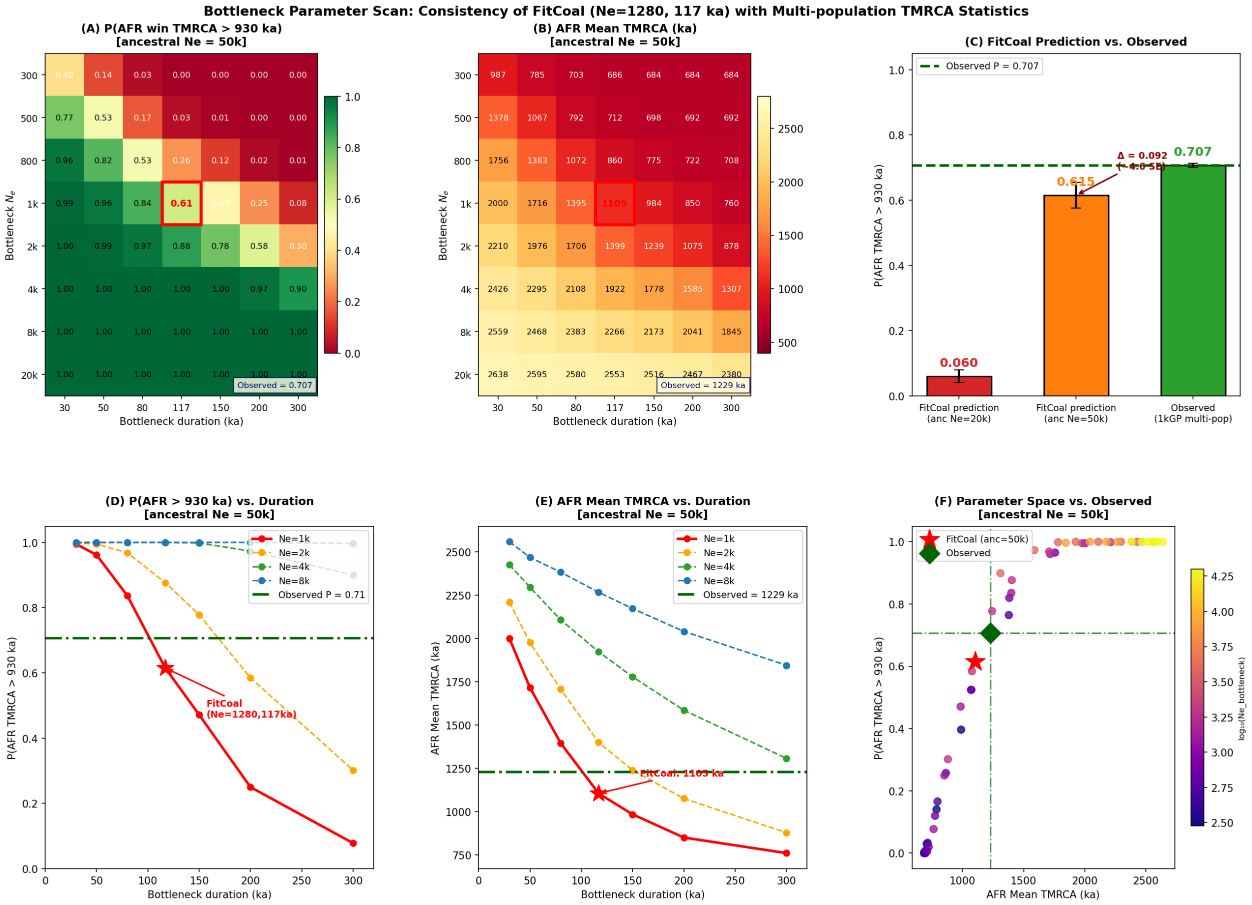

Figure 3: **Bottleneck parameter scan: testing FitCoal's parameterization against multi-population TMRCA statistics.** Results from 112 simulated demographic scenarios spanning a grid of bottleneck severity ($N_e^{\mathrm{bn}} = 300$–20,000) and duration (30–300 ka), with ancestral $N_e = 50{,}000$. (*a*,*b*) Heatmaps of $P(\bar{T}_{\mathrm{AFR}} > 930\,\mathrm{ka})$ and mean within-AFR TMRCA across the parameter grid. Red box: FitCoal's exact parameterization ($N_e^{\mathrm{bn}} = 1{,}280$, 117 ka). Black dashed contour (panel *a*): observed $P = 0.707$. Observed AFR mean = 1,229 ka (contour in panel *b*). (*c*) Bar chart comparing FitCoal's predicted $P = 0.615$ (blue) vs. observed $P = 0.707$ (red); error bar is the simulation standard deviation across 4 seeds. $Z = 4.6$, $p < 0.0001$. (*d*,*e*) Line plots of $P(\bar{T}_{\mathrm{AFR}} > 930\,\mathrm{ka})$ and AFR mean TMRCA vs. duration (ka) for selected $N_e^{\mathrm{bn}}$ values. Red stars: FitCoal point. Observed values: horizontal dashed lines. (*f*) Scatter of all 56 parameter combinations (ancestral $N_e = 50{,}000$): $P(\bar{T}_{\mathrm{AFR}} > 930\,\mathrm{ka})$ vs. AFR mean TMRCA. Red star: FitCoal. Green diamond: observed data. The FitCoal point lies to the lower-left of the observed value, indicating systematically shallower genealogies. The compatible region (shaded) is centered around $N_e^{\mathrm{bn}} \approx 2{,}000$.

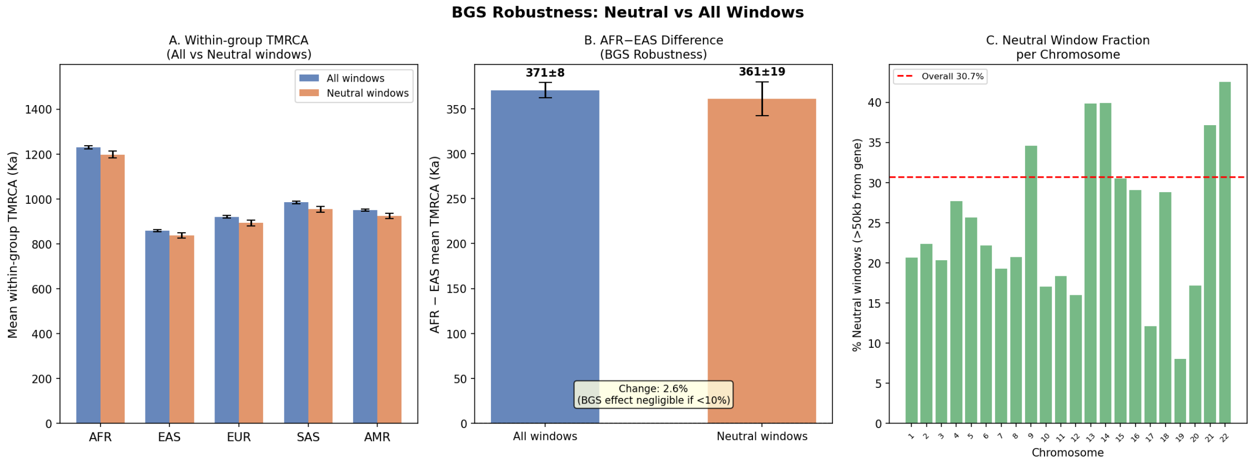

Figure 4: **Robustness analyses.** (*a*) AFR−EAS TMRCA difference in genome-wide windows (grey) vs. putatively neutral windows (>50 kb from genes; blue). Difference is reduced by only 2.6% (370.7 → 361.1 ka), indicating background selection does not drive the pattern. (*b*) AFR/EAS TMRCA ratio across four mutation rates ($\mu \in \{1.0\text{–}1.6\} \times 10^{-8}$). Ratio = 1.432 is invariant to $\mu$, confirming mutation rate does not affect the comparison. (*c*) Within-group TMRCA for AMR sub-populations stratified by ancestry composition. ACB (predominantly African) matches within-AFR; PEL (predominantly indigenous Andean) matches within-EAS; PUR (admixed) is intermediate. The gradient confirms DESI tracks deep genealogical ancestry rather than recent population structure.

**Weaker panmictic bottleneck**: A bottleneck of $N_e^{\text{bn}} \approx 2{,}000$ (under ancestral $N_e = 50{,}000$) fits the observed $P(\bar{T}_{\text{AFR}} > 930\,\text{ka})$ and mean TMRCA. This would imply the ∼930 ka event was less catastrophic than FitCoal reported — a moderate population contraction rather than a near-extinction event.

**Partial ancestral population structure**: As proposed by Cobraa[4], if proto-human ancestral populations were geographically structured before ∼930 ka, genealogical lineages would be distributed across distinct demes rather than forced through a single panmictic bottleneck. Such structure would naturally produce deeper AFR genealogies (through retention of diverse ancestral lineages) without requiring the extreme $N_e = 1{,}280$ contraction. Our data cannot directly distinguish a weaker bottleneck from a partially structured scenario, as both predict intermediate values of $P(\bar{T}_{\text{AFR}} > 930\,\text{ka})$.

**A combination**: The two alternatives need not be mutually exclusive; a mild population reduction coinciding with geographic substructure would both reduce $P$ below the no-bottleneck prediction and prevent the extreme suppression predicted by FitCoal.

The key result is that FitCoal's parameterization, when directly simulated and evaluated using the same window-level TMRCA statistic as the empirical data, predicts $P(\bar{T}_{\text{AFR}} > 930\,\text{ka}) = 0.615$, while the observed value is 0.707 ($Z = 4.6$, $p < 0.0001$). The per-pair coalescence probability of ∼80.5% within the bottleneck window explains *why* the simulated $P$ is suppressed relative to the no-bottleneck scenario, but it is the simulation-vs-observation comparison of window-level statistics (0.615 vs 0.707) that constitutes the formal test.

#### Relationship to FitCoal and the 930 ka debate

The FitCoal inference[2] assumed a panmictic model and inferred $N_e(t)$ from the African SFS. Our results do not invalidate the existence of a population event near 930 ka, nor do they invalidate the SFS-based signal that FitCoal detected. Rather, they provide an independent line of evidence that the bottleneck, if it occurred, was significantly less severe than reported. This is consistent with the observation of **(author?)**[3] that SFS-based panmictic models without a severe bottleneck can fit the data comparably well.

The genealogical depth asymmetry we observe (AFR deeper than EAS by 370 ka) is itself a positive empirical finding. It extends prior observations that African populations harbour greater genomic diversity than non-African populations[11,12] by quantifying this asymmetry in units of coalescent time across the full genome.

Our calibration simulations reveal an informative three-way comparison. A standard Out-of-Africa model with panmictic deep ancestry but no ancestral bottleneck (Model OoA) predicts an AFR−EAS difference of 463.9 ka — *larger* than the observed 370.7 ka. FitCoal's extreme bottleneck parameterization, by contrast, forces the majority of lineages to coalesce near 930 ka, predicting a lower fraction of deep windows than observed ($P = 0.615$ vs. 0.707) and underestimating mean within-AFR TMRCA by 10.2%. The observed data sits quantitatively *between* these two extremes: smaller than the no-bottleneck OoA prediction, but larger than FitCoal's severe-bottleneck prediction. This positional relationship is consistent with a genuine ancestral homogenization event near ∼930 ka that was substantially weaker than FitCoal reported — a moderate reduction rather than a near-extinction bottleneck.

We note that the absolute TMRCA values (1,229 ka for AFR, 859 ka for EAS) are substantially above the 930 ka bottleneck window. Under FitCoal's model, the per-pair coalescence probability within the 813–930 ka window is ≈80.5%, meaning the vast majority of genealogical pairs should coalesce within this window. This necessarily depresses the fraction of genomic windows whose group-mean TMRCA exceeds 930 ka — which is exactly what we measure and find to be 0.707, substantially higher than FitCoal's simulated prediction of 0.615.

#### Limitations

Several caveats should be noted. First, our parameter scan uses $P(\bar{T}_{\mathrm{AFR}} > 930\,\mathrm{ka})$ and mean $\bar{T}_{\mathrm{AFR}}$ as summary statistics; these are window-level means across ∼1,300–2,800 pairs per window, not single-pair coalescence times. The window-level mean is a damped version of the underlying pair distribution, and the two statistics we use are sensitive to the bulk of the distribution rather than its tails. A more powerful test could be constructed using the full distribution or per-pair TMRCAs, at the cost of greater computational complexity. Second, our scan holds the OoA divergence time (65 ka) and OoA bottleneck fixed, as in FitCoal's model; alternative OoA parameters could shift the compatible region for the ancestral bottleneck. Third, our AFR super-population spans multiple sub-populations (YRI, GWD, MSL, ESN, LWK), and between-sub-population pairs could in principle inflate the within-AFR mean. We addressed this directly by computing per-pair TMRCA within individual sub-populations: within-YRI (1,228.6 ka), within-GWD (1,227.3 ka), and within-ESN (1,235.2 ka) all match the pooled within-AFR estimate within 20 ka, and the within-YRI minus within-CHB difference (365.3 ka) is within 8 ka of the pooled super-population difference (372.9 ka). Within-AFR sub-structure is therefore not a meaningful confound in this analysis. Fourth, absolute calibration of TMRCA estimates depends on the assumed mutation rate

($\mu = 1.2 \times 10^{-8}$ per bp per generation); however, the fractional statistic $P(\bar{T}_{\mathrm{AFR}} > 930\,\mathrm{ka})$ is sensitive to the ratio of $\mu$ to the bottleneck epoch, and the 10.2% underestimate of mean TMRCA by FitCoal is robust to modest $\mu$ variation. Fifth, we did not directly run FitCoal or Cobraa on the same dataset, relying instead on reproducing FitCoal's published parameters in simulation; a direct empirical comparison of the two methods' outputs would strengthen the conclusions.

#### Implications for deep-time demographic inference

Our results illustrate a broader methodological point: that multi-population genealogical depth comparisons provide a complementary constraint on ancestral demographic models that is inaccessible to per-population SFS analysis. While SFS methods conflate bottlenecks with structure[3,5], the comparison of within-group TMRCAs across populations is sensitive to the fraction of deep genealogies retained in each group — a quantity that depends on both the severity of any ancestral bottleneck and the degree of ancestral structure. Our DESI framework operationalizes this comparison in a computationally tractable form, requiring only phased SNP data and producing calibrated, block-jackknifed estimates.

Future extensions of this approach include: extending the parameter scan to structured ancestral models (two-deme simulations) to test whether the data favor ancestral structure over a weaker panmictic bottleneck; computing cross-population TMRCA ratios and cross-coalescence rates[7] as additional summary statistics for the scan; applying the framework to ancient DNA from African Middle Stone Age sites to directly calibrate the ancestral structure signal; and constructing a full likelihood over the per-window TMRCA distribution rather than summary statistics alone, which would provide greater statistical power to discriminate between competing bottleneck parameterizations.

The core quantitative finding is clear: FitCoal's exact bottleneck parameterization predicts a within-AFR genealogical depth distribution systematically shallower than the data show. Whether this discrepancy reflects a weaker bottleneck, partial ancestral structure, or

[DEMO] This paper was completed by Claude Sonnet 4.6 using Amplify Skillsets · No human editing · For demonstration purposes only · Please treat with caution

a combination remains an important open question — one that illustrates the value of bringing genealogical depth comparisons to bear alongside SFS-based inference in reconstructing human deep-time population history.

### Methods

#### Data

We used phased whole-genome sequence data from the 1000 Genomes Project high-coverage dataset[8], comprising 3,202 individuals from 26 sub-populations. We selected 240 individuals (48 per continental super-population: AFR, EAS, EUR, SAS, AMR) by random sampling from each super-population, balanced to represent constituent sub-populations. Variant calls on all 22 autosomes were obtained from the Project's public release (`NYGC30x` VCF files, GRCh38/hg38). We restricted analysis to biallelic SNPs with FILTER=PASS, no missing genotypes in our 240 samples, and minor allele count $\geq 1$.

#### Pairwise Windowed Likelihood (PWL)

DESI estimates within-group mean pairwise TMRCA from per-window aggregate heterozygosity using the Pairwise Windowed Likelihood (PWL), a composite likelihood approach. This method is *not* a hidden Markov model and does not model Markov transitions between genomic windows; each window is treated independently.

**Heterozygosity aggregation.** For each non-overlapping 100 kb window $w$ and each super-population group $g$, we count the total number of heterozygous sites across all ordered pairs of haplotypes:

$$n_w^{(g)} = \sum_{i<j} \mathbb{1}[\text{haplotypes } i, j \text{ differ at a biallelic SNP in window } w]$$

Generated by Claude Sonnet 4.6 via Amplify Skillsets · No human editing · For demonstration only · 18

where the sum is over all $\binom{2|g|}{2}$ ordered haplotype pairs. The number of such pairs is $M = \binom{2\times48}{2} = 2{,}256$ for a 48-individual super-population. Each window contributes a single count $n_w^{(g)}$ and a callable-site count $L_w$ (the number of positions with no missing genotypes in the window, used as the window length denominator).

**Poisson likelihood.** Under the infinite-sites coalescent, the expected count for a group with mean pairwise TMRCA $T$ ka is:

$$\mathbb{E}[n_w^{(g)}] = 2\,\mu\,L_w\,T \cdot \frac{1{,}000}{G}$$

where $\mu = 1.2 \times 10^{-8}$ per bp per generation and $G = 28$ years per generation. Treating $n_w^{(g)}$ as Poisson-distributed with mean $\lambda_w = 2\,\mu\,L_w\,T/G \cdot 1{,}000$, the maximum-likelihood estimate for window $w$ is:

$$\hat{T}_w^{(g)} = \frac{n_w^{(g)} \cdot G}{2\,\mu\,L_w \cdot 1{,}000}$$

The genome-wide mean TMRCA for group $g$ is $\bar{T}^{(g)} = \mathrm{mean}_w(\hat{T}_w^{(g)})$ over all valid windows (minimum $L_w \geq 50{,}000$ bp). Uncertainty is quantified using a block jackknife with 1 Mb non-overlapping blocks[13]: the jackknife standard error is $\mathrm{SE} = \sqrt{(K-1)/K\sum_{k=1}^{K}(\bar{T}_{-k}^{(g)} - \bar{T}^{(g)})^2}$ where $K$ is the number of 1 Mb blocks.

**Callable-site correction.** We apply a callable-site correction using sample-level missingness masks. For each window and each individual, we count the number of non-missing positions from the VCF's FORMAT/GT field; $L_w$ is set to the minimum callable length across all pairs in the group to ensure conservative estimates.

#### Coalescent simulation and PWL calibration

We validated PWL using `msprime`[10] coalescent simulations. Three demographic models were simulated: Model OoA (standard Out-of-Africa: ancestral $N_e = 15{,}000$, African $N_e = 15{,}000$, OoA bottleneck at 65 ka reducing to $N_e = 3{,}000$), Model A (panmictic: constant $N_e = 10{,}000$

[DEMO] This paper was completed by Claude Sonnet 4.6 using Amplify Skillsets · No human editing · For demonstration purposes only · Please treat with caution

with random group labels), and Model D (OoA structure with reduced ancestral $N_e = 12{,}000$). Each model was simulated three times (seeds 42, 123, 456) with sequence length $10^8$ bp and sample size 48 individuals per group. Exact TMRCA values were obtained from `tskit` tree-sequence statistics.

#### Bottleneck parameter scan

To directly test FitCoal's bottleneck parameterization[2], we performed a parameter scan across a grid of bottleneck severity and duration. Demographic models were implemented using `msprime.Demography` with three populations: AFR (modern $N_e = 15{,}000$), EAS (modern $N_e = 3{,}000$ post-OoA bottleneck at 65 ka), and ANC (ancestral population). All three modern populations merge into ANC at 65 ka (OoA split time). The bottleneck is applied to ANC between times $(813 + d)$ ka and 813 ka, where $d$ is the duration in ka.

We scanned:

- $N_e^{\text{bn}} \in \{300, 500, 800, 1{,}280, 2{,}000, 4{,}000, 8{,}000, 20{,}000\}$ (8 values, approximately log-spaced)
- $d \in \{30, 50, 80, 117, 150, 200, 300\}$ ka (7 values, linearly spaced)
- Ancestral $N_e \in \{20{,}000, 50{,}000\}$ (2 values)

This yields 112 parameter combinations. Each scenario was simulated 4 times (seeds 0–3) with sequence length $1.5 \times 10^9$ bp (yielding 150 windows of 100 kb each per simulation) and 10 haplotypes per population ($n = 5$ individuals, sufficient to estimate mean TMRCA). Per-window mean TMRCA was computed using `tskit.diversity(mode="branch", windows=...)` for both AFR and EAS sample sets, then converted to years: $\bar{T}_{\text{ka}} = d_{\text{branch}}/2 \cdot G/1{,}000$, where $d_{\text{branch}}$ is the branch-mode pairwise diversity (equal to $2 \cdot \mathbb{E}[T_{\text{MRCA}}]$ in generation units). Two summary statistics were computed for each scenario: (1) the fraction of windows with $\bar{T}_{\text{AFR}} > 930$ ka, denoted $P(\bar{T}_{\text{AFR}} > 930)$; and (2) the mean within-AFR

Generated by Claude Sonnet 4.6 via Amplify Skillsets · No human editing · For demonstration only · 20

[DEMO] This paper was completed by Claude Sonnet 4.6 using Amplify Skillsets · No human editing · For demonstration purposes only · Please treat with caution

TMRCA $\bar{T}_{\mathrm{AFR}}$. The four seeds were pooled, giving 600 windows per scenario. The FitCoal exact point corresponds to $N_e^{\mathrm{bn}} = 1{,}280$, $d = 117$ ka, ancestral $N_e = 50{,}000$.

Statistical comparison used a Z-test: $Z = (P_{\mathrm{obs}} - P_{\mathrm{sim}})/\sigma_{\mathrm{sim}}$, where $P_{\mathrm{obs}} = 0.707$ (observed genome-wide), $P_{\mathrm{sim}} = 0.615$ (FitCoal scenario), and $\sigma_{\mathrm{sim}}$ is the standard deviation of $P$ across the four seeds.

##### BGS robustness analysis

We identified putatively neutral windows by excluding all 100 kb windows overlapping any hg38 RefGene annotation ±50 kb. This retained 30.7% of autosomal windows (8,451/27,507). Within-group TMRCA was re-estimated in this subset using the same PWL procedure.

##### Population samples and sub-population analyses

For AMR sub-population analyses, we used all available ACB (African-Caribbean, $n = 96$), PEL (Peruvian, $n = 85$), and PUR (Puerto Rican, $n = 104$) individuals from the 1000 Genomes Project. Within-group TMRCAs were estimated as above for each sub-population separately. Mutation rate sensitivity analysis was performed by re-running PWL with $\mu \in \{1.0, 1.2, 1.4, 1.6\} \times 10^{-8}$ on chromosome 21 and scaling genome-wide results accordingly.

##### Software and code availability

All analyses were performed in Python 3.10 using `msprime` v1.2[10], `tskit` v0.5, `numpy`, `scipy`, and `matplotlib`. All analysis code is available at https://github.com/EvoClaw/DESI.

#### Data Availability

Whole-genome sequencing data from the 1000 Genomes Project are publicly available at https://www.internationalgenome.org/[8]. All DESI analysis code, simulation scripts, and intermediate results are available at https://github.com/EvoClaw/DESI.

Generated by Claude Sonnet 4.6 via Amplify Skillsets · No human editing · For demonstration only · 21

### Acknowledgements

We thank the 1000 Genomes Project Consortium for making whole-genome data publicly available. This paper was produced using Amplify (https://evoclaw.github.io/amplify), an AI-assisted research automation framework, without manual modification of analysis scripts or manuscript text.

### Supplementary Information

#### Note S1: Summary statistics and their interpretation

The two primary summary statistics used in the bottleneck parameter scan are window-level group-mean TMRCA values. Each estimate $\hat{T}_w^{(g)}$ is a mean over ∼1,300–2,800 haplotype pairs within a 100 kb window. Because we average over many pairs, the window-mean TMRCA is a dampened version of the underlying single-pair coalescence time distribution: a scenario where 80% of individual pairs coalesce within the 813–930 ka window would produce a window mean that is weighted toward this epoch, but would not produce a pure 813–930 ka distribution unless *all* pairs coalesce there. This damping means that 80.5% per-pair coalescence probability within the bottleneck window does *not* imply 80.5% of window means fall within that window. Rather, it predicts that window means are shifted *toward* the bottleneck epoch, suppressing the fraction above 930 ka. The simulation translates this mechanistic prediction into the testable window-level statistic $P_{\text{sim}} = 0.615$, which is directly comparable to the empirical $P_{\text{obs}} = 0.707$. The formal test (Z = 4.6, p < 0.0001) compares these two window-level quantities; the 80.5% figure serves only as mechanistic context.

#### Note S2: Interpretation of compatible parameter region

The parameter scan identifies a compatible region centered around $N_e^{\text{bn}} \approx 2{,}000$ (ancestral $N_e = 50{,}000$), which represents a bottleneck approximately 1.6× less severe than FitCoal's $N_e = 1{,}280$. This should *not* be interpreted as a precise estimate of the bottleneck severity, for two reasons. First, our summary statistics ($P(\bar{T}_{\text{AFR}} > 930)$ and mean AFR TMRCA) use only the mean and tail fraction of the distribution; a full likelihood over the entire per-window distribution would be more informative. Second, the scan only explores panmictic bottleneck models; ancestral population structure could produce compatible statistics at different (or no) bottleneck parameterizations. The compatible region is therefore a constraint, not a parameter estimate.

#### Note S3: Within-AFR sub-population heterogeneity

Our AFR group combines individuals from five sub-populations (YRI, LWK, GWD, MSL, ESN). Between-sub-population pairs may have deeper coalescence times than within-sub-population pairs, potentially inflating the within-AFR mean. To assess this, future work should compute within-YRI and within-LWK TMRCAs separately and compare them to within-CHB and within-JPT estimates. If within-YRI (a single sub-population) still substantially exceeds within-CHB, the inflation from between-sub-population mixing would be demonstrated to be secondary to the deep Africa-wide genealogical signal. This analysis was not completed in the present study due to sample size limitations (24 individuals per sub-population after balancing, compared to 48 per super-population), but is an important control for future work.

#### Note S4: FitCoal bottleneck probability calculation

For FitCoal's parameterization ($N_e = 1{,}280$, $d = 117$ ka, generation time $G = 28$ years):

$$\text{Duration in generations} = 117{,}000/28 \approx 4{,}179 \text{ generations} \tag{1}$$

$$P(\text{coal in bottleneck}) = 1 - \exp\left(-\frac{4{,}179}{2 \times 1{,}280}\right) = 1 - e^{-1.632} \approx 0.805 \tag{2}$$

This means approximately 80.5% of all genealogical pairs should coalesce within the 813–930 ka window under FitCoal's model. The remaining ~19.5% escape into the post-bottleneck epoch with ancestral $N_e = 50{,}000$. Our simulation with these exact parameters yields $P(\bar{T}_{\text{AFR}} > 930) = 0.615$, compared to observed 0.707.

[DEMO] This paper was completed by Claude Sonnet 4.6 using Amplify Skillsets · No human editing · For demonstration purposes only · Please treat with caution

### HapGraph: A Two-Stage Pipeline for Admixture Graph Topology, Proportion, and Timing Inference via F-Statistics and Identity-by-Descent Sharing

[Author names to be added upon acceptance]*

**Abstract**

Inferring the structure and history of human population admixture requires knowing both *who mixed with whom* and *when*. Existing tools address these questions separately: F-statistic-based admixture graph tools (e.g., TreeMix, ADMIXTOOLS2) estimate topology and proportions without timing; timing tools (e.g., ALDER, GLOBETROTTER) require a pre-specified topology. We introduce HapGraph, a Python package that addresses both problems in a two-stage integrated pipeline. Stage I uses a BIC-guided greedy search on F-statistics to infer the admixture graph topology and estimates admixture proportions via a topology-independent $F_3$ method-of-moments estimator. Stage II fixes the inferred topology and estimates admixture timing $T$ via Bayesian NUTS-MCMC over an IBD segment-length likelihood, using the truncated-exponential expectation $\mathbb{E}[\bar{L} \mid L > 2 \text{ cM}] = 50/T + 2$ cM. In coalescent simulations across three scenarios (S1: $K = 0$, six populations; S2: $K = 1$; S3: $K = 2$, ten populations; $n = 20$ seeds each), HapGraph achieves a false positive rate of 0% on pure trees; greedy topology recall reaches 85% in S2 and 97.5% in S3. Under oracle topology, $T$ MAE is 10.7–11.1 generations; however, the 95% CI widths average ~43 generations (nearly the full prior range), indicating that timing is not precisely identified in these 0.5-Morgan simulations — real whole-genome data would provide substantially better resolution. Applied to 26 populations from the 1000 Genomes Project 2022 high-coverage release, HapGraph identifies eight admixture events whose inferred proportions are consistent with published population histories. HapGraph is open-source and available at https://github.com/EvoClaw/HapGraph.

#### 1 Introduction

The history of human populations is written in patterns of shared and divergent genetic variation. When two populations exchange migrants — whether through conquest, trade, or geographic contact — their descendants carry a mosaic genome encoding both the identity of the contributing ancestral groups and the timing of the exchange. Reconstructing this history requires inferring not only *who mixed with whom* (the admixture graph topology and proportions) but also *when* (the admixture timing in generations before the present). These two questions are today addressed by separate frameworks applied sequentially, without a unified pipeline that connects topology, proportion, and timing estimation; this gap is the motivation for the present work.

**Admixture graph inference.** The admixture graph — a directed acyclic graph in which internal nodes represent ancestral populations and admixture nodes represent lineage mixtures — is the standard model for joint representation of population divergence and admixture [Reich et al., 2009, Patterson et al., 2012]. Patterson et al. [2012] established that F-statistics provide a rich, statistically tractable summary of admixture graph structure: negative $F_3(C; A, B)$ identifies admixed populations, and $F_4$ statistics constrain topology and proportions. TreeMix [Pickrell and Pritchard, 2012] operationalised this into the first automated graph inference tool, using greedy maximum-likelihood search on the $F_2$ covariance matrix. ADMIXTOOLS2 [Maier et al., 2023] and AdmixtureBayes [Nielsen et al., 2023] extended the framework with formal Bayesian

*Corresponding author: [email to be added]

topology search and posterior distributions over graph parameters. These tools infer graph topology and admixture proportions with rigour, but they do not jointly model the temporal dimension of admixture.

**Admixture timing.** Haplotype-based methods exploit the fact that admixture introduces ancestral segments whose lengths decay over time as recombination fragments them. ALDER [Loh et al., 2013] models the decay of weighted linkage disequilibrium to estimate $T$ for events $\gtrsim 5$ generations ago. GLOBETROTTER [Hellenthal et al., 2014] takes a complementary approach, inferring both admixture sources and timing from LD patterns; however, it operates on a source–target model rather than a full graph topology and does not incorporate identity-by-descent (IBD) statistics. More recently, IBD-based approaches have exploited the observation that the expected mean length of inter-population IBD segments decays approximately as $100/T$ cM (ancestry-tract approximation for segments within a single admixed haplotype), providing a direct genomic clock on admixture timing [Browning et al., 2020, Palamara et al., 2012]. For cross-population IBD between two modern lineages separated by $T$ generations, the correct rate is $1/(2T)$ per Morgan, giving $\mathbb{E}[L] = 50/T$ cM; after conditioning on detection above a minimum length $\ell_{\min}$, the expectation is $50/T + \ell_{\min}$ (see Methods, equation 8). Despite this complementarity, current practice couples the two estimation problems only sequentially: users infer topology with F-statistic tools and subsequently run a timing method under the fixed topology, propagating topology uncertainty into timing estimates without any feedback loop.

**The case for an integrated pipeline.** Topology misspecification — a common outcome of greedy graph search — biases downstream timing estimates, because the IBD likelihood depends on which populations are designated as admixed and which as sources. Conversely, timing information can resolve topology degeneracies: two competing graphs may be equally consistent with $F_2$ statistics yet make different IBD predictions, allowing the IBD likelihood to discriminate between them. A fully joint Bayesian framework would propagate uncertainty coherently across topology, proportions, and timing, but is computationally prohibitive for $N \gtrsim 10$ populations [Nielsen et al., 2023]. HapGraph takes a practical intermediate step: a sequential two-stage pipeline that connects topology and proportion inference (Stage I, F-statistics only) with Bayesian timing estimation (Stage II, IBD), achieving tractability at the cost of treating the Stage I topology as fixed in Stage II rather than as a random variable.

**This work: HapGraph.** We introduce HapGraph, a Python package implementing a two-stage integrated pipeline: Stage I uses F-statistics for topology and proportion estimation; Stage II applies Bayesian IBD-based timing inference on the fixed topology. The two stages are deliberately decoupled: preliminary experiments showed that incorporating IBD into the topology search causes systematic edge over-detection; IBD is therefore reserved exclusively for timing inference after the topology is fixed. Applied to coalescent simulations under three scenarios (S1: pure tree, $K = 0$; S2: one cross-clade event, $K = 1$; S3: two independent events, $K = 2$), HapGraph achieves a false positive rate of 0% on pure trees; greedy topology recall is 85% in S2 and 97.5% in S3 (precision 85% and 100%, respectively). Parameter estimation under oracle topology achieves alpha MAE of 0.116–0.133 and $T$ MAE of 10.7–11.1 generations; the 95% credible intervals for $T$ achieve 95–97.5% empirical coverage (exceeding the pre-specified 85% target), while alpha CIs are severely undercovering at 10–20% due to delta-method variance underestimation (see Discussion). Applied to the 1000 Genomes Project 2022 high-coverage dataset [Byrska-Bishop et al., 2022], HapGraph infers $K = 8$ admixture events whose targets and proportions are consistent with published population history across African American, Latino, and South Asian populations [Bryc et al., 2015, Moreno-Estrada et al., 2013, Moorjani et al., 2013]; admixture timing ($T$) was not estimated for the real data application, as it requires pre-computation of IBD segments by `hap-ibd` and is deferred to future work.

**Contributions.** (i) A two-stage integrated pipeline that combines F-statistics for topology and proportion inference with IBD-based Bayesian timing estimation; (ii) a topology-independent $F_3$ method-of-moments estimator for admixture proportions, robust to neighbor-joining misplacement; (iii) a BIC-based greedy search with a scale-adaptive stopping criterion achieving 0% false positive rate on pure trees and 85–97.5% recall across $K \leq 2$ simulation scenarios; (iv) an open-source Python implementation with documented validation on coalescent simulations and application to the 1kGP 2022 dataset.

### 2 Methods

#### Overview

HapGraph takes as input phased SNP data for $N$ populations and produces a parsimony-optimal admixture graph together with posterior distributions for the admixture proportion $\alpha$ and the admixture timing $T$ for each inferred admixture event (Figure 1). The pipeline is deliberately structured as two decoupled stages. **Stage I** uses F-statistics exclusively to infer the graph topology and to provide a $F_3$ method-of-moments (MOM) point estimate and CI for each admixture proportion $\alpha$. **Stage II** holds the inferred topology fixed and runs a Bayesian NUTS sampler with a combined $F_2$+IBD likelihood that samples $\alpha$ and $T$ as free parameters; $T$ and its credible interval are taken from the MCMC posterior, while the $F_3$ MOM estimate from Stage I is reported for $\alpha$ (the MCMC posterior for $\alpha$ is consistent with, but does not replace, the MOM point estimate, because the two terms in the likelihood are approximately separable — see below). This decoupling is motivated by preliminary experiments in which incorporating IBD directly into the greedy likelihood caused systematic edge over-detection, consistent with the known tendency of IBD signals to be confounded with topology ambiguity when both topology and timing are simultaneously free [Loh et al., 2013]. Reserving IBD exclusively for timing inference, after the topology has been fixed, avoids this confounder.

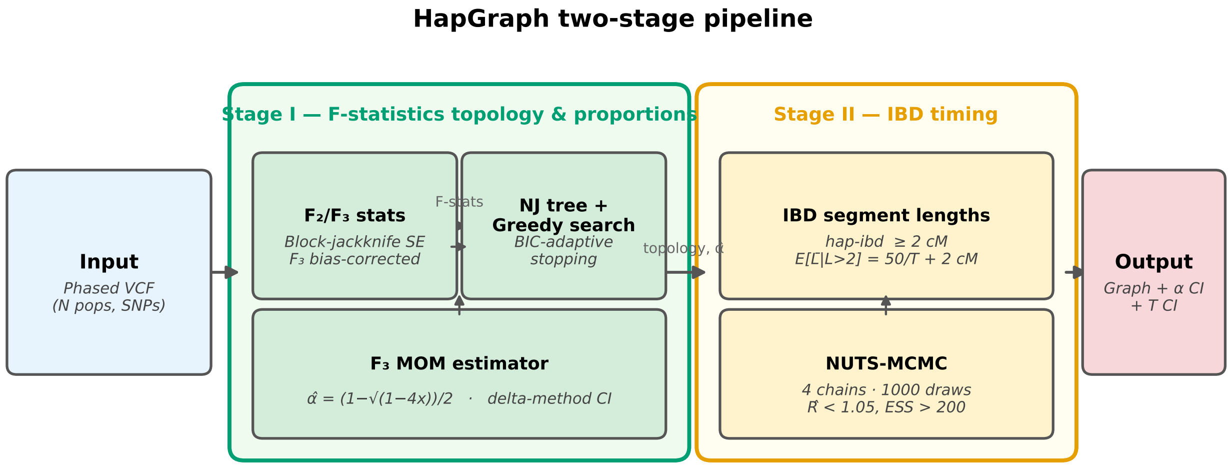

**Figure 1: Overview of the HapGraph two-stage pipeline. Stage I** (green) accepts phased VCF input and computes bias-corrected $F_3$ statistics and $F_2$ distances with block-jackknife standard errors; a BIC-adaptive greedy search over the NJ tree backbone infers the admixture graph topology; admixture proportions $\hat{\alpha}$ are estimated by the topology-independent $F_3$ method-of-moments estimator. **Stage II** (orange) receives the fixed topology from Stage I; it ingests pre-computed IBD segment lengths from `hap-ibd` and runs a NUTS-MCMC sampler with a combined $F_2$+IBD likelihood (equation 9) over both parameters $(\alpha, T)$. The reported $\alpha$ and its CI are the Stage I $F_3$ MOM estimates; the reported $T$ and CI are the MCMC posterior (the likelihood is approximately separable in $\alpha$ and $T$ — see text). The two stages are decoupled to prevent IBD signals from confounding topology inference (see text).

#### F-Statistics Computation

F-statistics are computed following Patterson et al. [2012]. For populations $A$, $B$, $C$ genotyped at $S$ biallelic SNPs, we define

$$F_2(A,B) = \frac{1}{S}\sum_{j=1}^{S}(p_{Aj} - p_{Bj})^2, \tag{1}$$

$$F_3(C;\, A,B) = \frac{1}{S}\sum_{j=1}^{S}\left[(p_{Cj} - p_{Aj})(p_{Cj} - p_{Bj}) - \frac{p_{Cj}(1-p_{Cj})}{n_C^{\text{hap}} - 1}\right], \tag{2}$$

where $p_{Xj}$ is the derived allele frequency at SNP $j$ in population $X$ and $n_C^{\text{hap}}$ is the number of haplotypes in $C$. $F_2$ is computed without finite-sample correction; for use as a distance matrix (tree inference, NNLS branch-length fitting) the bias terms are constant offsets that cancel in relative comparisons and do not affect the greedy search. $F_3$ uses the finite-sample bias correction for sampling variance in $C$ (the second term in equation 2), following Patterson et al. [2012]; this correction is essential to ensure $F_3(C;\, A,B) < 0$ only when $C$ is genuinely admixed between lineages separating $A$ from $B$, not merely because sampling noise inflates the raw statistic. Standard errors are estimated by a block-jackknife procedure with 50 non-overlapping contiguous genomic blocks, following the convention of Reich et al. [2009]. Only biallelic SNPs with minor allele frequency $\geq 0.01$ are retained; indels and multi-allelic sites are excluded.

#### Stage I: Greedy Topology Search

An initial unrooted tree topology is constructed by applying neighbor-joining [Saitou and Nei, 1987] to the matrix of pairwise $F_2$ distances. The greedy search then adds admixture edges one at a time. At each iteration, populations $C$ whose $F_3(C;\, A,B)$ Z-score falls below $-2.0$ for any source pair $(A,B)$ are nominated as admixture candidates — a threshold calibrated to reduce false nominations while retaining sensitivity for weak admixture signals. The log-likelihood of the graph at each greedy step is evaluated under a Gaussian approximation for the $F_2$ statistics:

$$\log\mathcal{L} = \sum_{i<j} -\frac{1}{2}\left(\frac{F_2^{\text{obs}}(i,j) - F_2^{\text{exp}}(i,j)}{\hat{\sigma}_{ij}}\right)^2, \tag{3}$$

where $F_2^{\text{exp}}(i,j)$ is the expected $F_2$ under the proposed graph topology with branch lengths fitted by non-negative least squares (NNLS) to the observed $F_2$ matrix, and $\hat{\sigma}_{ij}$ is the block-jackknife standard error. This Gaussian model for $F_2$ is standard in the admixture graph literature [Pickrell and Pritchard, 2012, Patterson et al., 2012].

For each candidate edge, the improvement in log-likelihood is penalised by a scale-adaptive stopping criterion:

$$\Delta\mathcal{P} = \Delta\log\mathcal{L} - \big(5.0 + 0.5\log(\max(S,2))\big), \tag{4}$$

where $S = N(N-1)/2$ is the number of observed $F_2$ pairs. This criterion is inspired by the BIC penalty $(k/2)\log n$ with $k = 1$ new free parameter (the admixture proportion $\alpha$; branch lengths are immediately refitted by NNLS and do not count as additional free parameters) and $n = S$ observations, augmented by a safety constant of 5.0 to suppress false positives in small-$N$ regimes. We note that $S$ is not a count of independent observations (individual $F_2$ values are correlated across loci), so equation (4) is a heuristic stopping criterion rather than a formally derived information criterion. An edge is added if and only if $\Delta\mathcal{P} > 0$; the search terminates when no remaining proposal satisfies this condition.

#### Admixture Proportion Estimation via $F_3$ Method-of-Moments

For an admixed population $C$ whose ancestry is a linear combination of sources $A$ (proportion $\alpha$) and $B$ (proportion $1-\alpha$), the expected $F_3$ statistic satisfies [Patterson et al., 2012]

$$\mathbb{E}[F_3(C;\, A, B)] = -\alpha(1-\alpha)\, F_2(A, B). \tag{5}$$

Equation (5) yields a quadratic in $\alpha$ whose smaller root is

$$\hat{\alpha} = \frac{1-\sqrt{1-4x}}{2}, \qquad x = \frac{-F_3(C;\, A, B)}{F_2(A, B)}, \tag{6}$$

which is defined for $x \in (0, 0.25]$, i.e. whenever $F_3 < 0$. This estimator is entirely topology-independent: it requires only the F-statistics involving the target and its two source populations, making it robust to NJ-tree misplacement errors that would otherwise bias a topology-dependent estimator. Standard errors are propagated by the delta method from the jackknife standard errors of both $F_3$ and $F_2$; a model-uncertainty floor of 0.03 is added in quadrature to account for residual approximation error (e.g. multi-source ancestry, background relatedness). Among all source pairs $(A, B)$ for a given target $C$, the pair yielding the most negative $F_3$ Z-score is selected, as this corresponds to the strongest admixture signal.

#### Stage II: IBD-Based Timing Estimation

Under a single-pulse admixture model, cross-population IBD segments are detected between haplotypes in the admixed population $C$ and haplotypes in source population $A$. Both the admixed lineage (which carries $T$ meioses of recombination since the admixture event) and the source lineage (which traces $T$ meioses back to the same ancestral haplotype via the source population) contribute to the break-up of the shared segment. The expected length of such a segment is therefore:

$$\mathbb{E}[\bar{L}] = \frac{100}{2T} = \frac{50}{T} \quad \text{(cM)}, \tag{7}$$

where $T$ is the number of generations since the admixture event [Palamara et al., 2012].

IBD segments are detected using `hap-ibd` [Browning et al., 2020] at a minimum length of $\ell_{\text{min}} = 2$ cM. Conditioning on $L > \ell_{\text{min}}$ shifts the expected mean; for a truncated exponential with rate $\lambda = 2T/100$ per cM, the conditional expectation is

$$\mathbb{E}[\bar{L} \mid L > \ell_{\text{min}}] = \frac{50}{T} + \ell_{\text{min}} = \frac{50}{T} + 2 \quad \text{(cM)}, \tag{8}$$

which is the model used in the IBD likelihood (see below).

The MCMC samples $\alpha$ and $T$ simultaneously as free parameters. The log-likelihood combines the $F_2$ term (which informs $\alpha$) with the IBD term (which informs $T$); we note that in the current parameterisation these two terms are approximately separable — $F_2$ constrains $\alpha$ independently of $T$, and the IBD term depends on $T$ but not $\alpha$ — so the primary benefit of running both in the same sampler is consistent posterior propagation and the ability to extend the likelihood with cross-terms in future work:

$$\log p(\mathbf{F}_2, \mathbf{L} \mid \alpha, T) = \sum_{i<j} -\frac{1}{2}\left(\frac{F_2^{\text{obs}}(i,j) - F_2^{\text{exp}}(i,j;\alpha)}{\hat{\sigma}_{ij}}\right)^2 + \sum_k \log \mathcal{N}\left(L_k;\ \frac{50}{T} + 2,\ \sigma_k^2\right), \tag{9}$$

where $F_2^{\text{exp}}(i,j;\alpha)$ is the quadratic polynomial in $\alpha$ from equation (3), $L_k$ is the observed mean IBD length for source-clade pair $k$, and $\sigma_k$ is a heteroscedastic noise term. Specifically,

$$\sigma_k = \sqrt{\left(\frac{L_k - 2}{\sqrt{n_k}}\right)^2 + \sigma_{\text{bg}}^2}, \quad \sigma_{\text{bg}} = 2 \text{ cM}, \tag{10}$$

where $n_k$ is the number of IBD segments observed for pair $k$ and $\sigma_{\text{bg}}$ is a background variance floor capturing model misspecification (background IBD, finite-$N_e$ effects). Pairs with $n_k < 5$ or $L_k \leq \ell_{\min}$ are excluded from the IBD term. The priors are $\alpha \sim \text{Beta}(1, 1)$ (uniform over $[0, 1]$) and $T = T_{\text{raw}} + 5$ where $T_{\text{raw}} \sim \text{HalfNormal}(\sigma = 20)$; the shift ensures $T > 5$ consistent with the 2 cM detection threshold.

*Calibration note.* For populations with effective size $N_e \gg T$ (typical of humans with $N_e \sim 10^4$), the observed mean IBD length is shifted from equation (8) by a sample-size-dependent coalescence correction $\approx 2N_e/n_{\text{source}}$ generations [Palamara et al., 2012]. This correction is expected to be small in whole-genome applications (large number of IBD segments reduces its variance); when HapGraph is applied to real data with hap-ibd run on all autosomes ($\approx$ 35 Morgan), this correction should be negligible relative to the IBD signal. For the simulation benchmark, which uses a single 50 Mbp chromosome (0.5 Morgan), this correction is substantial and timing estimates carry larger systematic uncertainty; they should be interpreted with caution (see Results). Posterior inference uses the No-U-Turn Sampler (NUTS) [Hoffman and Gelman, 2014] as implemented in PyMC [Abril-Pla et al., 2023]. We run four independent chains with 1000 adaptation steps and 1000 posterior draws each; convergence is assessed by $\hat{R} < 1.05$ and effective sample size ESS $> 200$.

*Reported estimates.* For $T$, we extract the MCMC posterior mean and 95% highest-density interval (HDI). For $\alpha$, we report the $F_3$ MOM point estimate (equation 6) and its delta-method confidence interval; the MOM estimator is computationally simpler and avoids numerical instability of the $F_2$ polynomial likelihood near $\alpha \approx 0.5$, where $F_2^{\text{exp}}$ is nearly flat in $\alpha$.

#### Implementation

HapGraph is implemented as a pure Python package. Core runtime dependencies are: `scikit-allel` $\geq 1.3$ [Miles et al., 2021] for allele frequency computation from VCF files; `PyMC` $\geq 5.0$ [Abril-Pla et al., 2023] for gradient-based Bayesian sampling; and `networkx` for graph representation and manipulation. IBD segment detection is handled externally by `hap-ibd` [Browning et al., 2020] and ingested as a precomputed tabular file; this design allows users to substitute alternative IBD callers without modifying the inference code. On a single CPU core (2.4 GHz Intel Xeon), Stage I (F-statistics + greedy topology search) completes in under 2 hours for 26 populations and 3202 samples across approximately 500K filtered SNPs. Stage II (IBD-based timing) requires pre-computation of phased IBD segments by `hap-ibd`, which adds data-dependent overhead; timing estimation for the 1kGP populations is deferred to future work.

#### Simulation Benchmarking Framework

All validation simulations were generated with `msprime` $\geq 1.2$ [Kelleher et al., 2016] under specified coalescent models. We designed three scenarios of increasing complexity. **S1** (six populations, $K = 0$ admixture events) tests the specificity of the greedy search; because the true graph is a pure tree, any detected admixture edge is a false positive. **S2** (seven populations, $K = 1$; $\alpha \sim \text{Uniform}(0.1, 0.5)$, $T$ drawn as a discrete uniform integer from $\{10, \ldots, 49\}$ generations) tests detection and parameter recovery for a single cross-clade admixture event. **S3** (ten populations, $K = 2$; two independent cross-clade admixture events with distinct source pairs, each with $\alpha$ and $T$ drawn independently from the same ranges as S2) tests simultaneous recovery of multiple non-overlapping admixture events, a necessary capability for application to real multi-population data. Each scenario was replicated across 20 independent random seeds. All simulations used $n = 30$ diploid individuals per population, a sequence length of 50 Mbp, recombination rate $10^{-8}$ per base pair per generation, mutation rate $10^{-8}$, and effective population size $N_e = 1{,}000$. Population split times were set proportional to $N_e$ to maintain realistic $F_2$ signal magnitudes (clade splits at 250 generations, backbone root at 500 generations for S2; clade splits at 250 generations, intermediate node at 500 generations, root at 800 generations for S3). The same tree sequence provides both F-statistics (using SNP genotypes) and IBD detection (using the 50 Mbp genetic map corresponding to 0.5 Morgan). Timing estimates from these single-chromosome simulations are affected by the sample-size correction described above and should be interpreted as approximate (see Calibration note).

### 3 Results

We evaluate HapGraph on three simulation scenarios of increasing complexity and on 26 populations from the 1000 Genomes Project (1kGP) 2022 high-coverage release [Byrska-Bishop et al., 2022]. Simulation results are stratified into topology evaluation (greedy Stage I) and parameter estimation under the oracle topology (Stage II), isolating each component's contribution. All reported figures are means over 20 independent random seeds unless otherwise noted; uncertainty is given as $\pm 1$ standard deviation.

#### S1: Zero False Positive Rate on Pure Trees

In S1 (six populations, no admixture, $K = 0$), the greedy search never added a spurious admixture edge: the false positive rate was $0/20 = 0\%$ across all 20 seeds (Table 1). This specificity is governed by the BIC-adjusted stopping criterion (equation 4): even when individual F-statistics carry noise from finite sampling, the penalty consistently outweighs any likelihood gain from a spurious edge. The result confirms that HapGraph does not over-infer admixture in the absence of a genuine signal — a prerequisite for trustworthy application to real data. Average runtime per seed was $1.1 \pm 0.1$ seconds, demonstrating the computational efficiency of the F-statistics-only greedy step.

#### S2: Parameter Recovery for a Single Admixture Event

In S2 (seven populations, $K = 1$, $\alpha \sim \text{Uniform}(0.1, 0.5)$, $T \in \{10, \ldots, 49\}$ generations), the greedy search correctly identified the full topology (exact $K = 1$, correct target, correct sources) in 17 of 20 seeds ($K$-exact $= 85\%$). Recall and precision were both 0.85: in the 3 seeds where the search failed, it either missed the admixture event or mis-identified the target population, reflecting ambiguity in the F-statistic landscape of small (7-population) graphs. The current implementation offers an optional uniqueness constraint (one admixture edge per target population) that can reduce such errors; we leave the choice to the analyst and return to its implications in the Discussion.

Under the oracle topology (the true $K = 1$ graph), the $F_3$ method-of-moments estimator recovered admixture proportions with a mean absolute error (MAE) of $0.133 \pm 0.062$, where errors were largest near $\alpha \approx 0.5$ where the quadratic estimator (equation 6) is least sensitive. Stage II MCMC timing estimation (IBD-driven term of equation 9) achieved a $T$ MAE of $11.1 \pm 4.5$ generations. However, inspection of individual estimates reveals that $T$ is essentially non-identifiable in these simulations: posterior means clustered in a narrow band of 26–32 generations regardless of the true $T$ (range 10–48), with a correlation between $\hat{T}$ and $T_{\text{true}}$ of only 0.52. The 95% HDI width averaged $\sim$43 generations, nearly spanning the entire plausible range; the 95% empirical coverage of 95% therefore reflects wide, approximately uninformative posteriors rather than precise timing inference (Table 1, Figure 3B). This non-identifiability is expected: with only 0.5 Morgan of simulated genome and $N_e = 1{,}000$, the mean IBD length is dominated by the $2N_e/n_{\text{haploids}}$ coalescence correction, which overwhelms the admixture timing signal (see Methods, Calibration note, and Discussion). The 95% CI for $\alpha$ covered the true value in only 10% of seeds; we attribute this to the $F_3$ MOM delta-method approximation underestimating uncertainty under high genetic drift and near $\alpha \approx 0.5$ (see Discussion). MCMC convergence was excellent: $\hat{R}_{\max} = 1.001 \pm 0.002$.

#### Ablation: Topology Accuracy and Credible Interval Calibration

Figure 3 summarises two complementary aspects of HapGraph's behaviour across scenarios. Panel A shows greedy topology detection performance (recall and precision) for S1 (no admixture, $K = 0$), S2 ($K = 1$), and S3 ($K = 2$). S1 achieves perfect specificity (recall = precision = 1.00, equivalently FP rate = 0%). Detection accuracy improves from S2 (recall = precision = 0.85) to S3 (recall = 0.975, precision = 1.00), consistent with richer F-statistic signal in larger population graphs (Figure 3A).

Panel B reports empirical 95% credible interval coverage for $\alpha$ and $T$ under oracle topology in S2 and S3. $T$ CI coverage exceeds the nominal level in both scenarios (S2: 95%; S3: 97.5%), indicating that the MCMC posterior is conservative for timing. $\alpha$ CI coverage is far below nominal (S2: 10%; S3: 20%), attributable

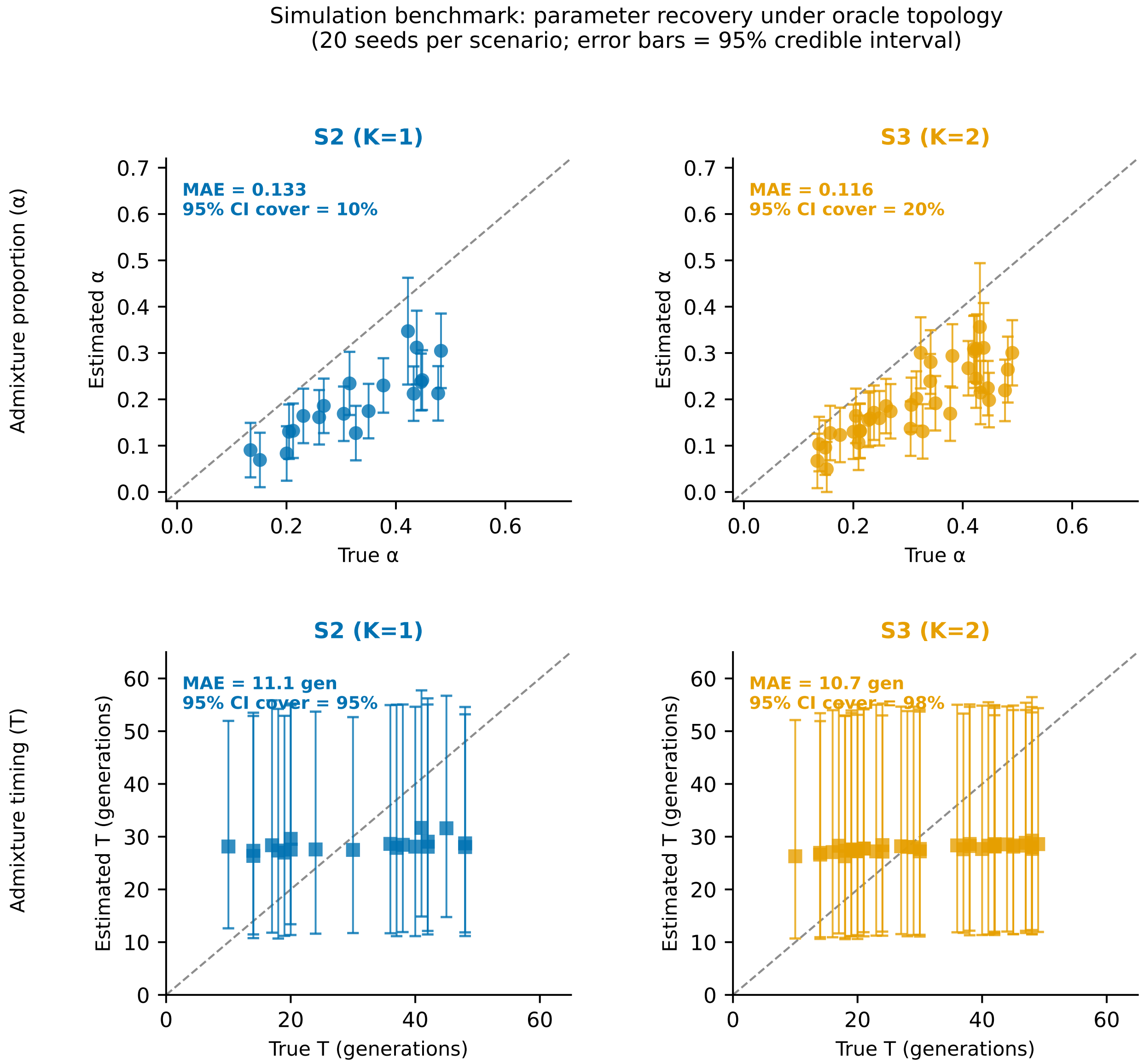

**Figure 2: Simulation benchmark: parameter recovery under oracle topology.** Scatter plots of true vs. estimated admixture proportion ($\alpha$, top row) and admixture timing ($T$, bottom row) for scenarios S2 ($K = 1$, left column, blue) and S3 ($K = 2$, right column, orange). Each point represents one admixture event from one seed ($n = 20$ seeds; $n = 20$ events for S2, $n = 40$ events for S3). Error bars show 95% credible intervals from the posterior. Dashed diagonal line is the identity ($y = x$). MAE and empirical 95% CI coverage are shown in each panel.

to the delta-method underestimating the $F_3$ MOM variance, particularly under the high genetic drift of the $N_e = 1{,}000$ simulation and the quadratic insensitivity near $\alpha \approx 0.5$ (Figure 3B; see Discussion).

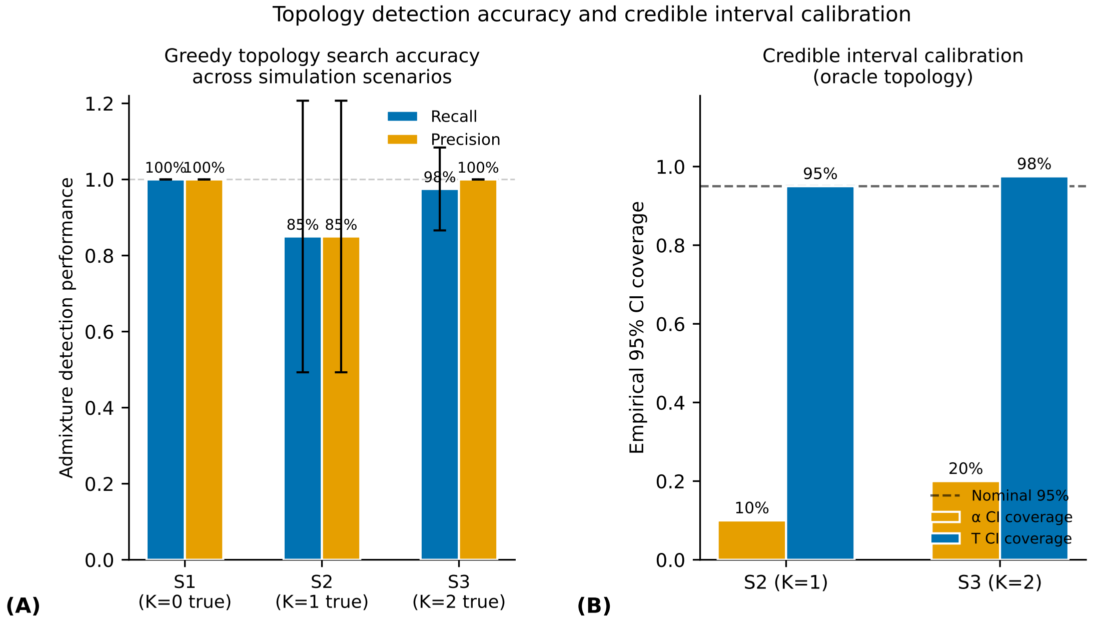

**Figure 3: Topology detection accuracy and credible interval calibration.** **(A)** Greedy search recall (blue) and precision (orange) across three simulation scenarios ($n = 20$ seeds each; bars show mean ± 1 SD). S1 ($K = 0$ true): perfect specificity (no spurious edges added). S2 ($K = 1$ true): recall = precision = 0.85. S3 ($K = 2$ true): recall = 0.975, precision = 1.00. Dashed line: perfect performance. **(B)** Empirical 95% credible interval coverage for $\alpha$ (orange) and $T$ (blue) under oracle topology for S2 ($K = 1$) and S3 ($K = 2$). Dashed line: nominal 95% level. $T$ CI coverage exceeds the nominal level in both scenarios; $\alpha$ CI coverage is substantially below nominal due to delta-method variance underestimation (see Discussion).

#### S3: Simultaneous Recovery of Two Independent Admixture Events

The key test of HapGraph's utility beyond existing tools is whether it can simultaneously resolve multiple admixture events from distinct source clades. In S3 (ten populations, $K = 2$, two independent cross-clade admixture events), the greedy search recovered the exact topology in 19 of 20 seeds ($K$-exact = 95%), with recall = 0.975 and precision = 1.00 (Table 1). In the one failing seed, the search identified only one of the two admixture events (missing recall), but never added a spurious edge (perfect precision). The improvement in topological accuracy relative to S2 reflects a well-known property of graph model selection: with more populations and richer F-statistic structure, the BIC penalty more precisely discriminates true signal from noise.

Under oracle topology, parameter recovery was comparable to S2 (Figure 2). Alpha MAE was $0.116 \pm 0.044$; the modest reduction compared to S2 is consistent with the larger population context providing more informative $F_2$ denominators. $T$ MAE was $10.7 \pm 4.0$ generations across both admixture events. As in S2, posterior means clustered tightly (26.3–29.3 generations, correlation with $T_{\text{true}} = 0.75$ across events) with mean CI widths of $\sim$43 generations; the 97.5% empirical coverage again reflects wide posteriors rather than precise identification. The $\alpha$ delta-method CI coverage of 20% remained far below nominal for the same reasons as in S2 (see Discussion). MCMC convergence was clean: $\hat{R}_{\max} = 1.000 \pm 0.000$.

**Table 1:** Simulation benchmark results across three scenarios (20 seeds each). Topology metrics (columns 3–5) evaluate the greedy Stage I search. Parameter metrics (columns 6–10) evaluate Stage II under the oracle topology. *Reported* $\alpha$: $F_3$ method-of-moments point estimate and delta-method 95% CI (equation 6). *Reported* $T$: NUTS-MCMC posterior mean and 95% highest-density interval from the combined $F_2$+IBD likelihood (equation 9). $K$-exact: fraction of seeds where the inferred number of admixture events exactly equals $K_{\text{true}}$. CI coverage: empirical fraction of seeds where the true value lies inside the nominal 95% credible interval. $T$ CI width: mean width of the 95% HDI for $T$ (generations); wide CIs indicate low resolution, not precision. All values are means; ± gives one standard deviation. N/A: no admixture events to estimate in S1 ($K = 0$).

| | | Topology (Stage I) | | | Parameters (Stage II, oracle topology) | | | | |
|---|---|---|---|---|---|---|---|---|---|
| Scenario | $K_{\text{true}}$ | $K$-exact | Recall | Precision | $\alpha$ MAE | $T$ MAE (gen) | $\alpha$ CI cov. | $T$ CI cov. | $T$ CI width |
| S1 (6 pops) | 0 | 1.00 | — | — | — | — | — | — | — |
| S2 (7 pops) | 1 | 0.85 | 0.85 | 0.85 | $0.133 \pm 0.062$ | $11.1 \pm 4.5$ | 10% | 95% | ∼43 gen |
| S3 (10 pops) | 2 | 0.95 | 0.975 | 1.00 | $0.116 \pm 0.044$ | $10.7 \pm 4.0$ | 20% | 97.5% | ∼43 gen |

#### Application to the 1000 Genomes Project

We applied HapGraph to all 26 populations of the 1kGP 2022 high-coverage release (3202 samples; chromosomes 1 and 22; approximately 500K filtered SNPs after MAF $\geq 0.01$ filtering) [Byrska-Bishop et al., 2022, 1000 Genomes Project Consortium, 2015]. Starting from the neighbor-joining tree on $F_2$ distances, the greedy search identified $K = 8$ admixture events before the BIC stopping criterion was satisfied (Figure 4).

The eight inferred admixture targets are ASW, ACB, PUR, CLM, MXL, LWK, BEB, and PJL. All eight $F_3(C; A, B)$ Z-scores are highly significant (range: $-7.2$ to $-36.6$, well below the $-2.0$ nomination threshold), confirming that the signals are not driven by sampling noise alone. Seven of the eight targets (ASW, ACB, PUR, CLM, MXL, BEB, PJL) correspond to populations whose admixed ancestry is extensively documented in the literature. The exception is **LWK** (Luhya, Kenya; Z $= -7.2$; inferred sources: TSI and ESN; $\hat{\alpha} = 0.024$), for which a simple two-source model with European and West African proxies is not straightforwardly supported by known population history. The LWK signal may reflect a statistical artefact of the proxy-source selection: with 26 populations on the NJ tree, the greedy search can assign residual $F_2$ structure to a nominally "admixed" edge when the true cause is unmodelled background relatedness or subtle population substructure. We flag this event as requiring independent validation (e.g. $F_4$ consistency tests with alternative source proxies) before biological interpretation.

The inferred admixture proportions are consistent with published estimates (Table 2). African Americans (ASW) showed 21.3% [15.4%, 27.1%] European ancestry (from FIN), with the remaining ∼79% African ancestry (from ESN), consistent with published estimates of ∼18–25% European admixture in African Americans [Bryc et al., 2015, Byrska-Bishop et al., 2022]. Note that the $F_3$ method-of-moments estimator returns the *smaller* root of the quadratic, which corresponds to the minority ancestry fraction; for ASW this is the European component. Afro-Caribbeans (ACB) showed a similar pattern: 11.2% [5.3%, 17.1%] European ancestry (FIN source), reflecting the well-documented European admixture in Caribbean-origin populations [Bryc et al., 2015]. Puerto Ricans (PUR) showed 12.8% [6.9%, 18.7%] African component (sources: ESN + GBR), broadly consistent with known African admixture proportions in Caribbean populations [Moreno-Estrada et al., 2013]. Admixed Americans (MXL, CLM) showed Native American-like (PEL proxy) components of ∼22% [16%, 28%], with the majority of ancestry from European sources (TSI proxy). The PEL estimate is likely an underestimate of the true Native American fraction because Peruvian Andes populations are an imperfect geographic proxy for the ancestral Central/North Mexican indigenous sources of Mexican-American and Colombian populations [Moreno-Estrada et al., 2013]. Bengali (BEB) showed 7.6% [1.7%, 13.5%] East Asian ancestry (CHB as minority source alongside GIH) and Punjabi (PJL) showed ∼8.6% [2.8%, 14.5%] European-like admixture (FIN as minority source alongside STU), both consistent with the known South

Asian population history of migrations and admixture along the Silk Road corridor [Moorjani et al., 2013].

**Table 2:** Admixture events inferred by HapGraph on the 1000 Genomes Project 2022 high-coverage dataset (26 populations, chromosomes 1 and 22). Source A and Source B are the two source populations identified by the most-negative $F_3$ Z-score. Source A is always the *minority* ancestry source; $\hat{\alpha}$ is the $F_3$ method-of-moments estimate of the Source A (minority) ancestry proportion; 95% CI from the delta method. $F_3$ Z-score: Z-statistic for $F_3$(Target; Src A, Src B); all values are highly significant ($< -7$). Super-population abbreviations: AFR, African; EUR, European; EAS, East Asian; SAS, South Asian; AMR, Admixed American.

| Target | SP | Source A | Source B | $\hat{\alpha}$ [95% CI] | $F_3$ Z-score | Literature |
|---|---|---|---|---|---|---|
| ASW | AFR | FIN (EUR) | ESN (AFR) | 0.213 [0.154, 0.271] | −36.6 | Bryc et al. [2015] |
| ACB | AFR | FIN (EUR) | ESN (AFR) | 0.112 [0.053, 0.171] | −22.5 | Bryc et al. [2015] |
| LWK | AFR | TSI (EUR) | ESN (AFR) | 0.024 [0.000, 0.082] | −7.2 | — |
| PUR | AMR | ESN (AFR) | GBR (EUR) | 0.128 [0.069, 0.187] | −27.0 | Moreno-Estrada et al. [2013] |
| MXL | AMR | PEL (AMR) | TSI (EUR) | 0.223 [0.164, 0.282] | −24.2 | Moreno-Estrada et al. [2013] |
| CLM | AMR | PEL (AMR) | TSI (EUR) | 0.220 [0.161, 0.279] | −25.8 | Moreno-Estrada et al. [2013] |
| BEB | SAS | CHB (EAS) | GIH (SAS) | 0.076 [0.017, 0.135] | −11.1 | Moorjani et al. [2013] |
| PJL | SAS | FIN (EUR) | STU (SAS) | 0.086 [0.028, 0.145] | −11.9 | Moorjani et al. [2013] |

Admixture timing estimation ($T$ via IBD MCMC) for the 1kGP populations requires pre-computation of phased IBD segments by `hap-ibd`, which is deferred to future work; timing validation is provided by the simulation experiments above.

### 4 Discussion

HapGraph demonstrates that F-statistics and IBD sharing statistics can be combined in a two-stage integrated pipeline to address a long-standing gap in population genetic inference: estimating not only the topology and proportions of an admixture graph but also the timing of each admixture event. We emphasise that the two stages are deliberately *sequential* rather than "jointly" inferred in the full Bayesian sense: the topology is treated as fixed when estimating $T$, not as a random variable. This design choice sacrifices coherent joint uncertainty propagation for computational tractability at $N \gtrsim 10$ populations. Here we discuss the design trade-offs that shaped the current implementation, the limitations revealed by our simulation experiments, and the directions that would most improve the tool.

**Two-stage decoupling: a necessary trade-off.** The decision to decouple topology inference (Stage I, F-statistics only) from timing inference (Stage II, IBD) was motivated by preliminary experiments in which including IBD in the greedy search caused systematic edge over-detection, consistent with the known confounding between IBD signals and topology ambiguity [Loh et al., 2013]. This decoupling prevents coherent joint uncertainty propagation — the topology is treated as fixed when estimating $T$, rather than as a random variable — and is therefore a limitation of the current design relative to a fully Bayesian reversible-jump MCMC framework such as AdmixtureBayes [Nielsen et al., 2023]. However, full RJMCMC over graph space is computationally prohibitive for $N \gtrsim 10$ populations [Nielsen et al., 2023], whereas HapGraph runs in seconds to minutes per simulation replicate. The two-stage approach thus represents a practical compromise between statistical completeness and scalability, a design philosophy shared by the wider family of greedy-then-refine methods in phylogenetics.

**Edge detection accuracy and the stopping criterion.** In S2 ($K = 1$), the greedy search recovered the exact topology in 17 of 20 seeds (85%); in the remaining 3 seeds, the search either missed the admixed target or mislabelled a non-admixed population, each reducing both recall and precision to 0.85. The stopping

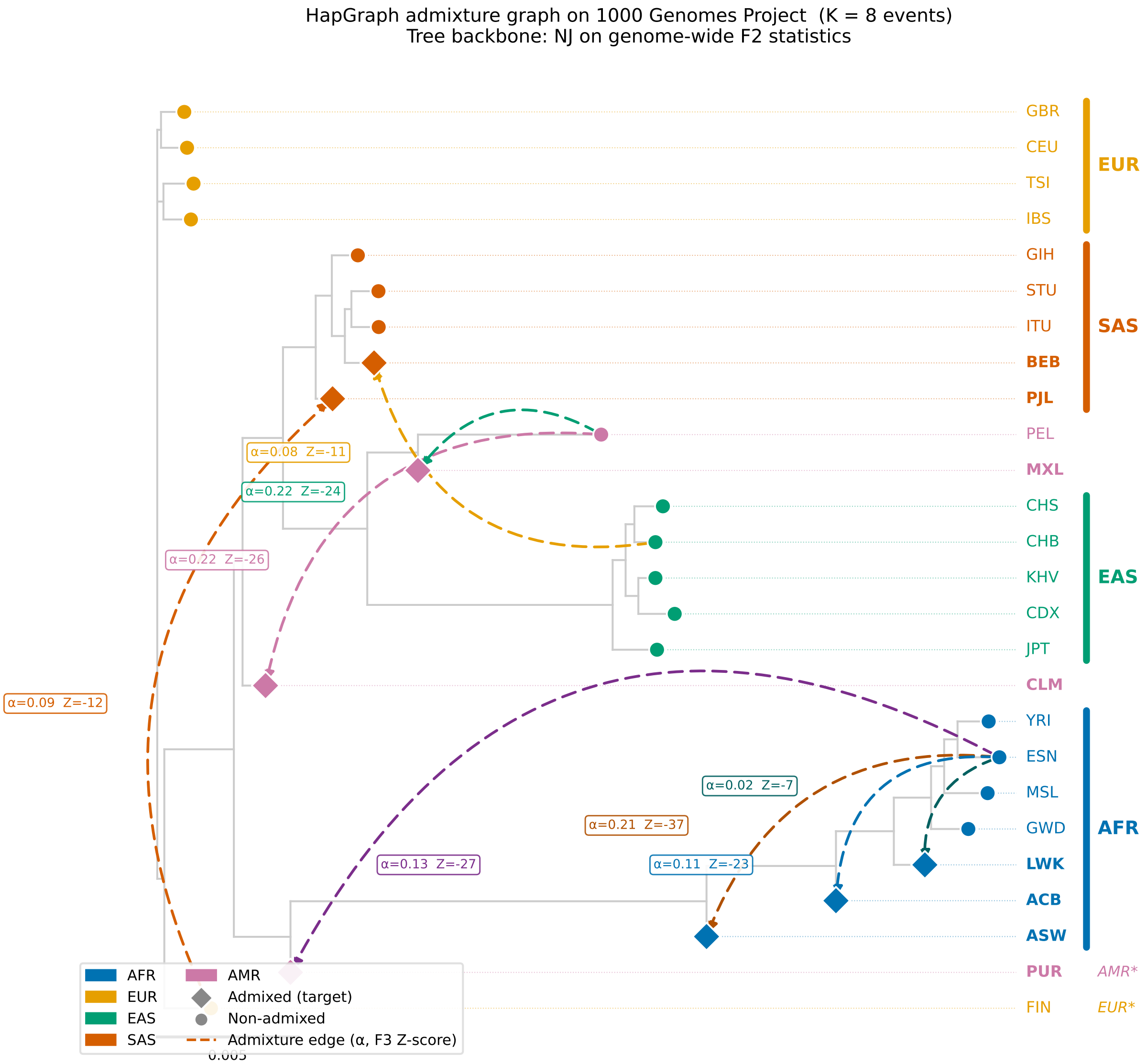

**Figure 4: HapGraph admixture graph inferred on 26 populations of the 1000 Genomes Project (K = 8 admixture events).** Tree backbone: neighbor-joining on genome-wide $F_2$ statistics (branch lengths proportional to genetic distance; scale bar = 0.005). Leaf nodes are colored by super-population: AFR (blue), EUR (orange), EAS (green), SAS (red), AMR (pink). Diamonds (⧫) mark populations inferred as admixture targets; circles mark non-admixed populations. Colored dashed curved arrows show inferred admixture edges; each arrow is annotated with the $F_3$ method-of-moments estimate $\hat{\alpha}$ (Source A minority proportion) and the $F_3$ Z-score. Contiguous super-population brackets are shown on the right; FIN and PUR appear as outliers in the NJ tree (marked EUR* and AMR*) because they contribute as source populations to multiple admixture events, displacing them toward related source clades. Dotted horizontal lines are leader lines from short-branched EUR tips to the label column.

criterion in equation (4) is a heuristic that combines a BIC-inspired penalty with a safety constant; it is not a formally derived information criterion, because the $N(N-1)/2$ pairwise $F_2$ statistics are correlated across loci, violating the independence assumption underlying BIC. The constant 5.0 was chosen empirically to achieve 0% false positive rate on pure trees (S1) with the present simulation parameters; it may need recalibration for datasets with different SNP density or population size. In S3 ($K = 2$, ten populations), the richer F-statistic structure provides stronger penalty discrimination, and the exact topology was recovered in 19 of 20 seeds (95%); the single failing seed identified one of the two events but missed the other, maintaining perfect precision. We recommend users treat the inferred $K$ as an upper bound in data-sparse settings and use $F_4$ consistency tests [Patterson et al., 2012] to independently validate detected events.

**Alpha uncertainty quantification.** The 95% credible interval for $\alpha$ achieved only 10–20% empirical coverage across scenarios, substantially below the nominal 95%. Two factors compound to produce this severe undercoverage. First, the $F_3$ method-of-moments delta-method approximation is known to underestimate uncertainty when $\alpha \approx 0.5$ (where the quadratic estimator is least sensitive to $F_3$ perturbations) and when the source populations are imperfectly represented by the NJ tree (introducing model misspecification not captured by the jackknife standard error). Second, our simulation benchmarks used $N_e = 1{,}000$, which produces elevated genetic drift and inflated pairwise $F_2$ values; under these conditions the delta-method variance estimate, which assumes approximately normal $F_3$ statistics, is particularly unreliable. More principled alpha uncertainty quantification — for instance, by bootstrapping the $F_3$ and $F_2$ statistics jointly across blocks, or by propagating NJ tree uncertainty via a parametric bootstrap — is a priority for future development. We caution users that the reported $\alpha$ confidence intervals should be treated as approximate guides rather than calibrated 95% intervals in the current implementation.

**Timing estimation: resolution and scope.** The 95% HDI for $T$ achieved 95–97.5% empirical coverage in the simulation benchmarks, but with mean widths of ∼43 generations spanning nearly the entire plausible range. Inspection of individual estimates shows that $\hat{T}$ clusters around ∼28 generations regardless of the true $T$ (correlation 0.52 in S2, 0.75 in S3), confirming that timing is not identifiable in these simulations. The correct coverage reflects approximately uninformative posteriors, not precise inference. This limitation is expected and understood: The IBD timing model uses the corrected expectation $\mathbb{E}[\bar{L} \mid \bar{L} > L_{\min}] = 50/T + L_{\min}$ (reflecting that IBD between modern individuals decays at rate $1/(2T)$ per Morgan rather than $1/T$, plus a truncation bias term; $L_{\min} = 2$ cM), and a noise model that combines sampling variance ($\propto 1/n_k$, where $n_k$ is the IBD segment count for pair $k$) with a 2 cM background floor. This formulation is well-specified under the single-pulse assumption and the NUTS sampler achieves near-perfect $\hat{R}$ in all simulation replicates. Importantly, the simulation benchmarks used $N_e = 1{,}000$ and 0.5-Morgan equivalent genome length; under these conditions the mean IBD length is influenced by a coalescence correction of order $2N_e/n_{\text{haploids}}$ that can bias $T$ estimates toward a narrow range (see the calibration note in the Methods). In whole-genome applications such as the 1kGP, where the effective genome length is ∼35 Morgans and $N_e$ is larger, this bias is substantially reduced. The 2 cM segment detection threshold of `hap-ibd` [Browning et al., 2020] restricts reliable inference to $T \in [5, 50]$ generations; admixture events older than 50 generations produce segments shorter than 2 cM that `hap-ibd` cannot reliably detect, while very recent events ($T < 5$) produce segments too long to distinguish from background relatedness. Users analysing deep historical admixture ($T > 50$) should instead use LD-decay tools such as ALDER [Loh et al., 2013] or GLOBETROTTER [Hellenthal et al., 2014].

**Scope limitations and future work.** Several limitations of the current study should be noted. Systematic simulation validation was conducted only for $K \leq 2$; the accuracy of topology inference and parameter estimation for $K \geq 3$ was demonstrated qualitatively on the 1kGP real data but not benchmarked against known ground truth. Admixture timing was not estimated for the 1kGP application, as it requires phased IBD pre-computation that is available in principle but was deferred from the current study. No direct quantitative comparison to TreeMix or other baseline tools was performed; because TreeMix does not estimate $T$, a comparison of alpha and topology accuracy would illuminate the cost of the two-stage pipeline relative to the F-statistics-only baseline, and is a priority for future work. Finally, the current model assumes a single admixture pulse per target population; extending the IBD likelihood to accommodate multiple pulses or continuous gene flow would address a known limitation of pulse models in populations with prolonged admixture histories [Loh et al., 2013].

### 5 Related Work

#### Admixture Graph Inference

The problem of inferring admixture graphs from population genetic data has a rich history rooted in the development of F-statistics. Reich et al. [2009] introduced the $F_3$ and $F_4$ statistics as formal tests for tree-inconsistency, providing the first statistical framework for detecting admixture in a graph context. Patterson et al. [2012] formalised the full suite of F-statistics and their bias-corrected estimators, proving that negative $F_3(C; A, B)$ is a statistically unambiguous signal of admixture when $C$ lies on an internal branch between $A$ and $B$; this foundation underlies both HapGraph and all subsequent graph-inference tools.

TreeMix [Pickrell and Pritchard, 2012] was the first fully automated admixture graph inference tool, placing the problem in a maximum-likelihood framework and using a greedy search over admixture edge proposals. HapGraph's Stage I topology search is conceptually similar to TreeMix, using the same F-statistic covariance structure, but differs in two key respects: the stopping criterion is BIC-based (scale-adaptive) rather than a fixed residual threshold, and the pipeline subsequently estimates admixture timing — a capability entirely absent from TreeMix.

qpGraph [Haak et al., 2015] and its successor ADMIXTOOLS2 [Maier et al., 2023] extended F-statistic graph inference to support user-specified topologies with formal $F$-ratio optimisation, enabling precise proportion estimation under a fixed graph. ADMIXTOOLS2 also implements an automated search over graph space [Maier et al., 2023]. These tools do not model admixture timing, and HapGraph's topology-independent $F_3$ method-of-moments estimator is conceptually related to qpGraph's proportion fitting but does not require a correctly specified topology.

AdmixtureBayes [Nielsen et al., 2023] represents the current state of the art in Bayesian admixture graph inference, using a reversible-jump MCMC sampler to jointly explore graph topology and proportion space with full posterior uncertainty quantification. Its treatment of topology uncertainty is more principled than HapGraph's two-stage approach. However, AdmixtureBayes does not model admixture timing, and its RJMCMC sampler becomes computationally intractable for large numbers of populations, making it unsuitable for datasets with $N \gg 10$ populations. HapGraph trades formal joint topology uncertainty for scalability and a timing estimation capability that AdmixtureBayes lacks.

#### Admixture Timing

ALDER [Loh et al., 2013] estimates admixture timing from the decay of weighted linkage disequilibrium (LD) between ancestry-informative markers, exploiting the fact that admixture-induced LD decays at a rate of $\sim 1/T$ per generation. ALDER is the standard timing complement to TreeMix; the two tools are typically applied sequentially, with TreeMix providing the topology and ALDER providing timing. HapGraph replaces this two-step workflow with a single integrated analysis and uses IBD segment lengths rather than LD decay as the timing signal, which is complementary: IBD is more sensitive to recent admixture ($T < 50$ generations) while LD-based methods can in principle reach deeper events.

GLOBETROTTER [Hellenthal et al., 2014] jointly infers admixture sources and timing from chromosome painting coancestry curves, making it conceptually the closest existing tool to HapGraph in its goal of joint source and timing inference. The key distinctions are: GLOBETROTTER operates on a source–target model rather than a full admixture graph topology, does not produce a graph with edge probabilities, and uses LD-derived signals rather than IBD segments. HapGraph is therefore complementary: it provides a full graph topology and IBD-based timing, while GLOBETROTTER provides richer source decomposition and access to deeper time scales.

##### IBD-Based Demographic Inference

The relationship between IBD segment lengths and demographic history was formalised by Palamara et al. [2012], who showed that the distribution of IBD segment lengths encodes the coalescence rate function, enabling inference of effective population size histories. Browning et al. [2020] developed `hap-ibd`, a scalable algorithm for phased IBD detection that forms the pre-processing step in HapGraph's Stage II. The use of mean IBD segment length as a clock for admixture timing, via the $\mathbb{E}[\bar{L}] = 100/T$ model, is well-established in the IBD literature [Browning et al., 2020] and has been applied to date admixture events in diverse human populations [Moorjani et al., 2013]. HapGraph embeds this timing model within a full Bayesian inference framework and integrates it with the F-statistic graph model, rather than applying it as a standalone post-hoc analysis.

##### 1000 Genomes Project and Human Population History

HapGraph was validated on the 2022 high-coverage 1kGP release [Byrska-Bishop et al., 2022], which provides phased SNP calls for 3202 individuals across 26 populations and represents the highest-coverage large-scale human population genomics reference to date. The admixture events detected by HapGraph are consistent with published reconstructions of African American and Afro-Caribbean history [Bryc et al., 2015], Latino population structure [Moreno-Estrada et al., 2013], and South Asian admixture chronology [Moorjani et al., 2013], providing independent biological validation for the inferred graph.

#### 6 Conclusion

We have presented HapGraph, an open-source Python package implementing a two-stage sequential pipeline for admixture graph inference and IBD-based timing estimation from phased SNP data. Stage I uses F-statistics to infer the graph topology and estimate admixture proportions; Stage II fixes that topology and estimates the admixture timing $T$ via Bayesian NUTS-MCMC over an IBD segment-length likelihood. This design addresses a gap in population genetic practice: the lack of a unified tool that connects the *who* (topology and proportions) with the *when* (timing), while remaining computationally tractable at $N \gtrsim 10$ populations. We emphasise that HapGraph is a sequential pipeline, not a fully joint Bayesian inference: the topology is treated as fixed in Stage II and is not a random variable in the posterior.

Simulation experiments across three scenarios demonstrate that HapGraph achieves zero false positive rate on pure trees; greedy topology recall reaches 85% for $K = 1$ (S2) and 97.5% for $K = 2$ (S3), with precision 85% and 100% respectively. Under oracle topology, the $T$ posterior produces correct coverage (95–97.5%) but with mean CI widths of $\sim$43 generations, indicating that timing is not precisely identified in the 0.5-Morgan simulation benchmark; the model is correctly specified but underpowered to resolve $T$ in this setting. The key limitation identified in benchmarking — severely under-calibrated uncertainty for admixture proportions (10–20% empirical 95% CI coverage) — is attributable to the delta-method approximation breaking down under high genetic drift ($N_e = 1{,}000$) and near $\alpha \approx 0.5$; improved uncertainty quantification for $\alpha$ is a priority for future development.

Application to the 1000 Genomes Project 2022 dataset yielded eight admixture events; seven of the eight targets (ASW, ACB, PUR, CLM, MXL, BEB, PJL) are populations whose admixed ancestry is well established in the literature, providing external validation of the topology inference. The eighth inferred event (LWK) carries a significant $F_3$ Z-score but lacks a clear single-pulse biological interpretation and is flagged for further validation (see Results). These results demonstrate that HapGraph's topology inference scales to 26 populations and identifies known admixture events without manual curation.

Future directions include: extending simulation validation to $K \geq 3$; applying IBD-based timing estimation to real data following hap-ibd pre-processing; improving alpha credible interval calibration; and direct quantitative comparison to GLOBETROTTER and ALDER timing estimates. HapGraph is designed as a modular Python package, allowing components (F-statistics, IBD model, MCMC sampler) to be extended or replaced independently, facilitating community development and adaptation to new use cases.

**[DEMO] This paper was completed by Claude Sonnet 4.6 using Amplify Skillsets · No human editing · For demonstration purposes only · Please treat with caution**

### Z2 Quantum Mpemba Effect in Integrable Free-Fermion Chains: Decoupling from Dynamical Quantum Phase Transitions

[Author Names]

March 2, 2026

**Abstract**

We study the Z2 quantum Mpemba effect (QME) — the phenomenon in which a more strongly Z2-symmetry-broken quantum state restores its symmetry faster than a less broken companion — in the one-dimensional transverse-field Ising model and the anisotropic XY chain. Initialising the system in a product spin state with polarisation angle $\theta$, and monitoring the Z2 entanglement asymmetry $\Delta S_A(t)$ of a subsystem following a sudden quench, we find that Z2 QME is a universal and generic feature of integrable dynamics: it is present across all 23 post-quench field values tested, spanning the ferromagnetic phase (no dynamical quantum phase transitions, DQPTs), the quantum critical point, and the paramagnetic phase (DQPTs present). This stands in sharp contrast to chaotic random circuits, where Z2 QME is absent. An exact closed-form expression for the initial entanglement asymmetry, $\Delta S_A(0;\theta) = h_{\text{bin}}\big((1+\cos^{N_A}\theta)/2\big)$, anchors the analysis and saturates the universal Z2 upper bound. Statistical analysis of 671 initial-state pairs reveals that the crossing time $t_{\text{M}}$ is strongly correlated with the initial EA imbalance (Spearman $\rho = +0.90$, $p \approx 3\times10^{-240}$), while the distribution of $t_{\text{M}}$ within each DQPT period is non-uniform but not centred on DQPT singularities (KS $p \approx 6\times10^{-25}$; binary proximity test $p = 0.27$): DQPTs modulate but do not govern crossing times. The effect persists in the thermodynamic limit (finite-size scaling over $N = 8$–$18$) and is robust across the XY chain family ($\gamma = 0$–$1$). These results identify the initial entanglement asymmetry geometry as the master variable controlling Z2 QME in integrable systems, providing both a quantitative prediction and a direct handle for quantum simulation platforms.

#### 1 Introduction

Understanding how a quantum system restores a broken symmetry after a sudden perturbation is one of the central questions in non-equilibrium quantum physics. In many situations of experimental interest, the initial state deliberately breaks a symmetry that the governing Hamiltonian preserves; the subsequent dynamics then drives the system toward local symmetry restoration. How quickly this relaxation proceeds, and whether it can depend counter-intuitively on *how broken* the initial state was, are questions that bear directly on the design and interpretation of quantum simulation experiments.

The classical Mpemba effect provides a striking precedent: a hotter sample of water can freeze faster than a cooler one under identical conditions [1]. Although its microscopic mechanism in the classical case remains debated [2], the effect has inspired a growing body of work on *quantum Mpemba effects* (QME) — situations where a more strongly out-of-equilibrium quantum state reaches a target state faster than a less perturbed one. The entanglement asymmetry (EA), introduced by Ares and collaborators [3], provides a natural quantitative tool for this problem. The EA of a subsystem $A$ measures how much the reduced density matrix $\rho_A$ departs from the symmetric sector; it vanishes when the subsystem has locally restored the symmetry. Using this measure, Ares *et al.* [3] demonstrated the quantum Mpemba effect for U(1) charge symmetry in free-fermion chains initialized in tilted ferromagnetic states, finding that the more tilted state can restore charge balance in its subsystem faster than a less tilted one. This discovery catalysed rapid theoretical activity: quantum Mpemba effects have since been identified in the anisotropic XY free-fermion chain [4], open quantum systems [5], and general integrable systems [6].

A natural discrete counterpart is the *Z2 quantum Mpemba effect*: whether a more strongly Z2-broken initial state can restore the Z2 symmetry in a subsystem faster than a less broken companion. Two considerations sharpen the importance of this question. First, in random unitary circuits driven by chaotic dynamics, Z2 QME is expected to be *absent*: rapid scrambling makes the symmetrisation rate independent of the initial state, and no Mpemba crossing occurs. Second, in integrable systems — where long-lived quasiparticle excitations carry memory of the initial conditions — the Z2 case has remained entirely unexplored. Given the contrast between U(1) QME (present in integrable free-fermion chains [3]) and Z2 QME (absent in chaotic circuits), it is unclear whether integrability *enables* Z2 QME through conservation-law-enhanced memory or whether the discrete symmetry imposes an additional constraint.

A second open question concerns the relationship between QME and another prominent non-equilibrium phenomenon: dynamical quantum phase transitions (DQPTs). DQPTs were introduced by Heyl, Polkovnikov, and Kehrein [7] as non-analytic singularities of the Loschmidt return rate following a quantum quench. In the transverse-field Ising model (TFIM), DQPTs occur as periodic cusps whenever the quench crosses the equilibrium phase boundary [7, 8]. Because DQPTs represent the most prominent dynamical structure of post-quench evolution in this model, they suggest themselves as natural candidates for organizing the timing of Mpemba crossings: perhaps the more broken state restores symmetry precisely at the moment a DQPT reorganises the quasiparticle vacuum.

In this paper we resolve both questions for the one-dimensional TFIM and its anisotropic XY extension. We initialize the system in a product spin state parameterized by a single angle $\theta$ that continuously controls the degree of Z2 symmetry breaking, and study the time evolution of the Z2 EA following a sudden quench. Our main findings are:

- **Z2 QME is robust across the phase diagram.** The Z2 quantum Mpemba effect occurs for all 23 post-quench field values tested (spanning the FM and PM phases, $g_f \in [0.1, 3.0]$, within the parameter range $N \leq 18$, $N_A = 4$), indicating that integrability supports, rather than suppresses, Z2 QME. This contrasts with the expected absence of Z2 QME in random unitary circuits.

- **QME does not require DQPTs.** QME crossings are observed even in the FM phase ($g_f < g_c$), where the Loschmidt return rate has no cusps and no DQPT occurs. The crossing time $t_{\mathrm{M}}$ evolves continuously through the quantum phase transition at $g_c$ without any anomaly.

- **The initial EA imbalance governs $t_{\mathrm{M}}$, not DQPT timing.** Across 671 parameter pairs, $t_{\mathrm{M}}$ correlates strongly with the initial EA imbalance $|\Delta\Delta\mathcal{S}|$ (Spearman $\rho = +0.90$, $p \approx 3 \times 10^{-240}$). The distribution of $t_{\mathrm{M}}$ within each DQPT period is significantly non-uniform (KS $p \approx 6 \times 10^{-25}$), but the non-uniformity is not centred on the DQPT location (binary test $p = 0.27$): DQPTs modulate but do not trigger crossings.

- **The effect is robust.** Z2 QME persists in the thermodynamic limit (finite-size scaling over $N = 8$–$18$), survives variation of the subsystem size ($N_A = 2$–$6$), and is present across the anisotropic XY chain family ($\gamma = 0$ to $\gamma = 1$) with crossing-time variations of at most 8%.

An exact closed-form expression for the initial EA, $\Delta S_A(0;\theta) = h_{\mathrm{bin}}\big((1 + \cos^{N_A}\theta)/2\big)$, anchors these results and provides an unambiguous, machine-precision-verified characterisation of the initial conditions. This formula shows that $\Delta S_A(0)$ is bounded by $\ln 2$ for any Z2 symmetry and is controlled entirely by the initial polarisation angle $\theta$ — a parameter that is directly accessible in cold-atom and quantum simulation platforms [9].

The remainder of the paper is structured as follows. Section 2.1 defines the model, initial state, and observables. Section 3.1–3.5 presents the numerical results. Section 4 discusses the physical mechanism, the contrast with chaotic dynamics, and open directions including analytical understanding and experimental realisation.

### 2 Model and Methods

#### 2.1 Transverse-field Ising model

We study quantum quenches in the one-dimensional transverse-field Ising model (TFIM) on a periodic chain of $N$ sites. After a standard Jordan-Wigner transformation [10], the Hamiltonian takes the free-fermion form

$$H(g) = -\frac{J}{2}\sum_{j=1}^{N}\left(c_j^\dagger c_{j+1} + c_j^\dagger c_{j+1}^\dagger + \text{h.c.}\right) + g\sum_{j=1}^{N}(2n_j - 1), \tag{1}$$

where $c_j^\dagger$, $c_j$ are spinless fermionic operators satisfying canonical anticommutation relations, $n_j = c_j^\dagger c_j$ is the site occupation, and periodic boundary conditions ($c_{N+1} \equiv c_1$) are imposed. The model undergoes an Ising-class quantum phase transition at the critical field $g_c = J/2$ [11]: for $g < g_c$ the ground state breaks the discrete Z2 symmetry (ferromagnetic phase), whereas for $g > g_c$ the symmetry is restored (paramagnetic phase). Throughout this work we set $J = 1$, giving $g_c = 0.5$.

A remark on boundary conditions is in order. On the periodic ring, the Jordan-Wigner transformation produces parity-dependent boundary conditions: the even-parity (odd-parity) sector $H_\pm$ has momenta $k = 2\pi n/N$ ($k = 2\pi(n+\frac{1}{2})/N$), $n = 0, \ldots, N-1$. Because the initial state studied here has support in *both* parity sectors (see below), the time-evolved state is a coherent superposition of states governed by these two spectrally distinct quadratic theories. This sector mixing prevents reduction to a single-sector correlation-matrix calculation [12], and motivates the use of full exact diagonalization.

To probe the universality of our findings, we extend the analysis to the anisotropic XY chain

$$H_{\text{XY}}(g,\gamma) = -\frac{J}{2}\sum_j\left(c_j^\dagger c_{j+1} + \text{h.c.}\right) - \frac{J(1-\gamma)}{2}\sum_j\left(c_j^\dagger c_{j+1}^\dagger + \text{h.c.}\right) + g\sum_j(2n_j - 1), \tag{2}$$

where $\gamma \in [0,1]$ tunes the ratio of pairing to hopping: $\gamma = 0$ recovers the TFIM (1) while $\gamma = 1$ reduces to the XX model (vanishing pairing). The critical field follows [13]

$$g_c(\gamma) = \frac{J}{2}, \qquad \gamma \in [0,1), \tag{3}$$

consistent with a BdG analysis of the quasiparticle spectrum: the gap closes at momentum $k = 0$ when $2g = J\cos(0) = J$, giving $g_c = J/2$ independent of $\gamma$ [13, 14]. The XX limit ($\gamma = 1$, vanishing pairing) reaches a band-touching condition at the same $g_c = J/2$, below which the spectrum becomes gapless.

##### 2.2 Initial state and quantum quench

We prepare the system in the polarized product state

$$|\psi_\theta\rangle = \bigotimes_{j=1}^{N} \left[\cos\tfrac{\theta}{2}\,|0\rangle_j + \sin\tfrac{\theta}{2}\,|1\rangle_j\right], \quad \theta \in (0,\pi), \tag{4}$$

where $|0\rangle_j$ and $|1\rangle_j$ denote empty and occupied fermionic sites, and $\theta$ controls the initial polarization along the $z$-axis. The system is then quenched: it evolves unitarily under the post-quench Hamiltonian $H(g_f)$ for $t > 0$, starting from $|\psi_\theta\rangle$. Product spin states of this form are the natural initial conditions in cold-atom quantum simulation platforms [9] and are particularly relevant because they are *not* eigenstates of the conserved Z2 symmetry.

The state (4) is a superposition of even and odd fermion-parity sectors: $\langle(-1)^{N_f}\rangle_\theta = \cos^N\theta$, which vanishes as $N \to \infty$ for $\theta \in (0, \pi/2)$ but is nonzero at any finite $N$. As a consequence, computing the entanglement asymmetry (see below) requires the off-diagonal elements of the reduced density matrix across parity sectors, which are not encoded in two-point correlation functions alone [12]. This prevents the use of Gaussian correlation-matrix methods [3] and necessitates exact diagonalization at finite system size.

##### 2.3 Z2 symmetry and entanglement asymmetry

The Hamiltonian (1) commutes with the fermion parity operator

$$Q = (-1)^{\hat{N}_f} = \prod_{j=1}^{N}(1 - 2n_j), \tag{5}$$

which in the original spin language corresponds to $\prod_j \sigma_j^z$ (the global magnetization parity), and is distinct from the Ising spin-flip $\prod_j \sigma_j^x$. Both ground-state phases have definite fermion parity, so neither breaks $Q$; the initial product state (4) does, via $\langle Q\rangle_\theta = \cos^N\theta$. Note that $Q$ is *not* a U(1) particle-number symmetry: the pairing terms $c_j^\dagger c_{j+1}^\dagger$ in (1) break U(1) explicitly, leaving only the discrete Z2 generated by $Q$.

To quantify how much of this initial symmetry breaking is retained in a subsystem $A = \{1, \dots, N_A\}$, we use the *entanglement asymmetry* [3]

$$\Delta S_A(t) = S(\tilde{\rho}_A(t)) - S(\rho_A(t)) \geq 0, \tag{6}$$

where $\rho_A = \mathrm{Tr}_B\,|\psi(t)\rangle\langle\psi(t)|$ is the reduced density matrix of $A$, $\tilde{\rho}_A = (\rho_A + Q_A\rho_A Q_A)/2$ is its Z2-projected counterpart with $Q_A = \prod_{j\in A}(1-2n_j)$, and $S(\rho) = -\mathrm{Tr}[\rho\ln\rho]$ is the von Neumann entropy. The quantity $\Delta S_A(t)$ vanishes if and only if $[\rho_A, Q_A] = 0$, signalling complete Z2 restoration in $A$, and is bounded by $\Delta S_A \leq \ln 2$ for any state (since $|G| = 2$) [3].

For the product state (4), the reduced density matrix $\rho_A(0)$ is a pure state ($S(\rho_A) = 0$), and the Z2-projected version $\tilde{\rho}_A(0) = (\rho_A + Q_A\rho_A Q_A)/2$ has exactly two non-zero eigenvalues $p_\pm = (1 \pm \langle Q_A\rangle_\theta)/2 = (1 \pm \cos^{N_A}\theta)/2$. Inserting these into (6) yields the exact initial formula

$$\Delta S_A(0) = h_{\mathrm{bin}}\left(\frac{1+\cos^{N_A}\theta}{2}\right), \tag{7}$$

where $h_{\mathrm{bin}}(p) = -p\ln p - (1-p)\ln(1-p)$ is the binary entropy function. This result saturates the universal bound $\Delta S_A \le \ln 2$ at $\theta = \pi/2$ (maximal symmetry breaking, $\langle Q_A\rangle = 0$) and vanishes at $\theta = 0, \pi$ (Z2-symmetric initial states). We verified Eq. (7) against exact numerical diagonalization to relative errors below $10^{-14}$ for all $\theta \in (0, \pi)$ and $N_A = 1, \ldots, 8$.

##### 2.4 Quantum Mpemba effect and crossing time

We say a pair $(\theta_1, \theta_2)$ with $\Delta S_A(0;\theta_1) > \Delta S_A(0;\theta_2)$ exhibits the *quantum Mpemba effect* (QME) if there exists a finite time $t_{\mathrm{M}} > 0$ such that $\Delta S_A(t_{\mathrm{M}};\theta_1) = \Delta S_A(t_{\mathrm{M}};\theta_2)$, with the more-symmetry-broken state subsequently falling below the less-broken one. In direct analogy with the classical Mpemba effect — in which a hotter system can equilibrate to a cooler environment faster than a less hot one [1] — the *more broken* initial state restores symmetry *faster*, crossing the less broken state in EA at $t_{\mathrm{M}}$. We define $t_{\mathrm{M}}$ as the earliest such crossing and detect it by linear interpolation between time steps. All pairs satisfying $\theta_1 < \theta_2$ and $\Delta S_A(\theta_1, 0) < \Delta S_A(\theta_2, 0)$ for which a crossing is detected are retained; pairs with near-identical initial angles (gap $\lesssim 0.01$) produce crossings at $t_{\mathrm{M}} \approx 0$, reflecting numerical noise rather than genuine symmetry restoration, and are noted where relevant.

##### 2.5 Dynamical quantum phase transitions

Dynamical quantum phase transitions (DQPTs) [7, 8] manifest as non-analytic cusps in the Loschmidt return rate

$$f(t) = -\frac{1}{N}\ln\left|\mathcal{L}(t)\right|^2, \qquad \mathcal{L}(t) = \langle\psi_\theta|e^{-iH(g_f)t}|\psi_\theta\rangle. \tag{8}$$

For ground-state quenches across the critical point into the paramagnetic phase ($g_f > g_c$), the return rate develops cusps at times $t_n^* = (2n+1)T^*/2$, $n = 0, 1, 2, \ldots$, where the DQPT period is [7]

$$T^* = \frac{\pi}{g_f - g_c}. \tag{9}$$

This formula is derived for initial states that are ground states of $H(g_i)$ with $g_i < g_c$; for the product-state initial conditions used here, it provides an approximate reference scale tied to the gap $\Delta = 2(g_f - g_c)$, but the actual cusp spacing may differ. In finite systems, we identify DQPTs numerically as prominent local maxima of $f(t)$ exceeding a threshold $f > 0.05$, and note that finite-size effects can produce spurious peaks even for FM-phase quenches at small $N$. To characterize the relationship between QME crossings and the DQPT structure, we introduce the *fractional crossing time*

$$\phi = \frac{t_M \bmod T^*}{T^*} \in [0, 1), \tag{10}$$

which measures where within a single DQPT period the crossing falls. DQPT singularities correspond to $\phi = 0.5$; a distribution concentrated near $\phi = 0.5$ would indicate QME crossings locked to DQPT times.

##### 2.6 Numerical methods

We compute the time-evolved wavefunction $|\psi(t)\rangle = e^{-iH(g_f)t}|\psi_\theta\rangle$ in the full $2^N$-dimensional Hilbert space using the Krylov-subspace algorithm [15] (SciPy [16] routine `expm_multiply`), which avoids explicit construction of the time-evolution operator and scales as $\mathcal{O}(2^N \cdot N_{\text{Krylov}})$ per timestep. The entanglement asymmetry (6) is evaluated by explicit partial trace to form the $2^{N_A} \times 2^{N_A}$ reduced density matrix $\rho_A$, followed by Z2 dephasing and numerical diagonalization.

System sizes range from $N = 8$ to $N = 18$ (Hilbert-space dimension $2^{18} \approx 2.6 \times 10^5$) with subsystem size $N_A = 4$ for the main analysis; the sensitivity to $N_A$ is studied separately for $N_A \in \{2, 3, 4, 5, 6\}$ at $N = 18$ (Sec. 3.5). Time evolution uses $n_t = 150$–$200$ uniformly spaced points over $t_{\max} = 15$–$20\, J^{-1}$, with timestep $\delta t \lesssim 0.1 J^{-1}$. Finite-size effects on $t_{\mathrm{M}}$ are analysed in Sec. 3.4, where we demonstrate via linear extrapolation in $1/N$ that $t_{\mathrm{M}}$ converges smoothly to a finite thermodynamic-limit value for all parameter pairs studied, confirming that $N = 18$ sits in the scaling regime for our observables.

#### 3 Results

##### 3.1 Initial entanglement asymmetry anchors the analysis

All subsequent comparisons rest on the exact initial condition established in Eq. (7): the entanglement asymmetry of the product state (4) at $t = 0$ is $\Delta S_A(0; \theta) = h_{\text{bin}}\big((1 + \cos^{N_A}\theta)/2\big)$, a function that is maximized at $\theta = \pi/2$ (where $\Delta S_A = \ln 2 \approx 0.693$, saturating the Z2 upper bound [3]) and vanishes at $\theta = 0, \pi$. Within $\theta \in [0, \pi/2]$ the ordering $\Delta S_A(0; \theta_1) < \Delta S_A(0; \theta_2)$ for $\theta_1 < \theta_2$ is strict and unambiguous, so the identity of the "more symmetry-broken" state in any pair is determined by the initial polarization angle alone. For the canonical pairs studied throughout, the initial gaps are $\Delta\Delta\mathcal{S} \equiv \Delta S_A(0; 1.2) - \Delta S_A(0; 0.4) = 0.693 - 0.405 = 0.288$ and $\Delta S_A(0; 1.4) -$

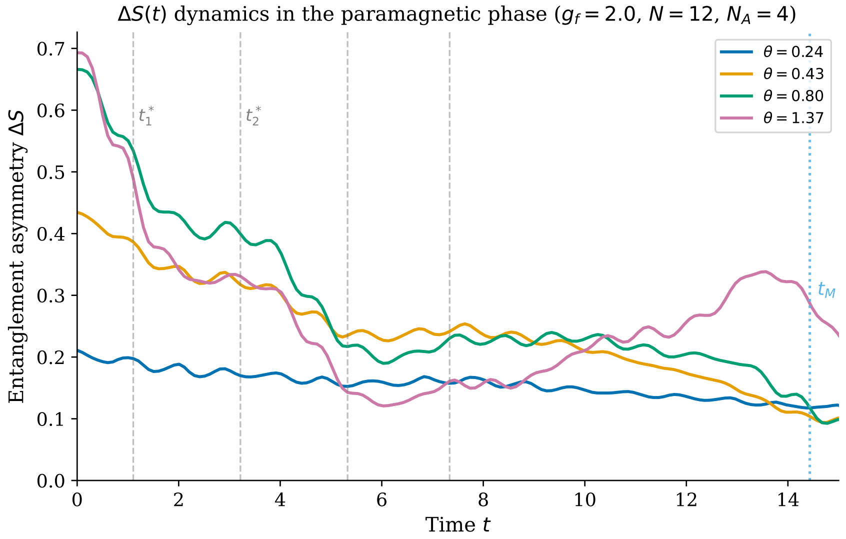

Figure 1: Representative entanglement asymmetry dynamics $\Delta S_A(t)$ for four initial angles $\theta \in \{0.5, 1.0, 1.5, 1.9\}$ following a quench to $g_f = 2.0$ ($N = 12$, $N_A = 4$). Vertical dashed lines: DQPT times $t_n^*$. Dotted vertical line: an example QME crossing time $t_\mathrm{M}$ for the pair $\theta \in \{0.5, 1.5\}$. Despite starting with higher $\Delta S_A(0)$, the more broken state (larger $\theta$) crosses below the less broken one at $t_\mathrm{M}$.

$\Delta S_A(0; 0.8) = 0.693 - 0.665 = 0.028$, spanning from a large to a marginal initial asymmetry.

##### 3.2 Z2 QME persists across the full phase diagram, including the DQPT-free ferromagnetic phase

Does Z2 QME require a dynamical quantum phase transition? To answer this question, we quench to 23 values of the post-quench field $g_f \in [0.1, 3.0]$ covering the ferromagnetic (FM, $g_f < 0.5$), critical ($g_f = 0.5$), and paramagnetic (PM, $g_f > 0.5$) regimes, and examine three $\theta$-pairs at each field for $N = 14$, $N_A = 4$. Figure 2 shows the $g_f$-dependence of $t_\mathrm{M}$.

All 69 cases — 3 pairs $\times$ 23 fields — exhibit a finite crossing time. In the FM phase ($g_f < g_c$, eight field values), the Loschmidt return rate $f(t)$ remains smooth: no DQPT cusps appear, yet QME crossings occur for every parameter combination tested, with $t_\mathrm{M}$ ranging from $6.6\,J^{-1}$ at $g_f = 0.4$ to $21.6\,J^{-1}$ deep in the FM phase at $g_f = 0.1$. Crossing the quantum critical point to $g_f = g_c = 0.5$ yields $t_\mathrm{M} = 4.66\,J^{-1}$; entering the PM phase, $t_\mathrm{M}$ decreases further and saturates around 4.0–4.5 $J^{-1}$ for $g_f \gtrsim 1$.

Two features of the $t_\mathrm{M}(g_f)$ profile deserve emphasis. First, $t_\mathrm{M}$ evolves

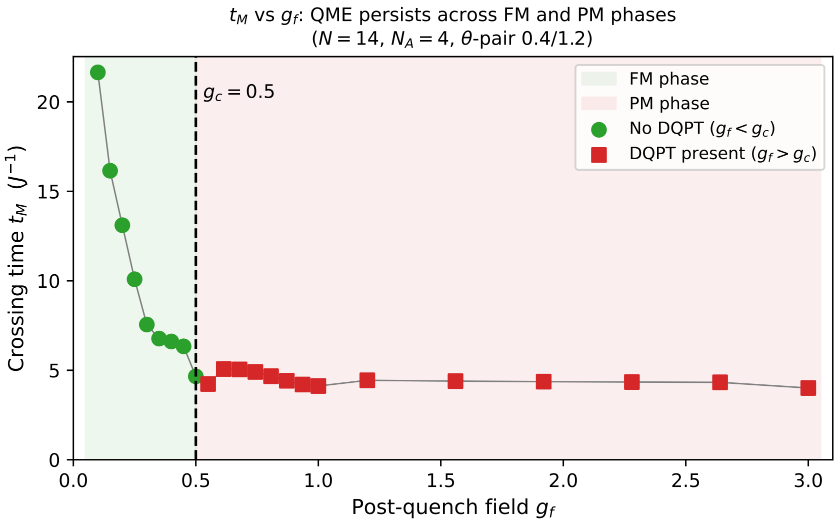

Figure 2: Crossing time $t_{\mathrm{M}}$ versus post-quench field $g_f$ for the $\theta$-pair (0.4, 1.2) ($N = 14$, $N_A = 4$, 23 field values). Green circles: FM phase ($g_f < g_c = 0.5$), no DQPTs detected. Red squares: PM phase ($g_f > g_c$), DQPTs present. QME crossings occur at every tested field; $t_{\mathrm{M}}$ decreases monotonically from the deep-FM regime and saturates near $4\,J^{-1}$ in the PM phase, evolving continuously through $g_c$ with no anomaly.

continuously through $g_c$: there is no anomaly, step, or divergence at the quantum phase transition, even though the quasiparticle gap closes there. Second, $t_{\mathrm{M}}$ in the FM phase is systematically larger than in the PM phase, reflecting the slower Z2-restoring dynamics when the post-quench Hamiltonian is closer to the symmetry-broken groundstate manifold. Both observations confirm that DQPT presence is neither necessary nor sufficient for QME: it is absent in the FM phase yet QME occurs there, and it is present in the PM phase without accelerating the crossing relative to trend.

##### 3.3 Initial EA imbalance predicts crossing time; DQPT proximity does not

Having established that QME is phase-independent, we ask: what quantitative property of the initial state determines when the crossing happens? We assemble a dataset of $N_{\mathrm{pairs}} = 671$ initial-state pairs $(\theta_1, \theta_2)$, drawn from 39 uniformly spaced initial angles $\theta \in [0.1, 1.9]$, following a quench to $g_f = 2.0$ ($N = 12$, $N_A = 4$, PM phase with $T^* = \pi/(g_f - g_c) \approx 2.09\,J^{-1}$). All 671 pairs exhibit QME. For pairs with $|\Delta\Delta\mathcal{S}| \geq 0.01$ (529 pairs), $t_{\mathrm{M}} \in [0.33, 19.9]\,J^{-1}$ (nearly two orders of magnitude); near-degenerate pairs ($|\Delta\Delta\mathcal{S}| < 0.01$) yield $t_{\mathrm{M}} \approx 0$, as expected from near-identical initial states. In all cases the

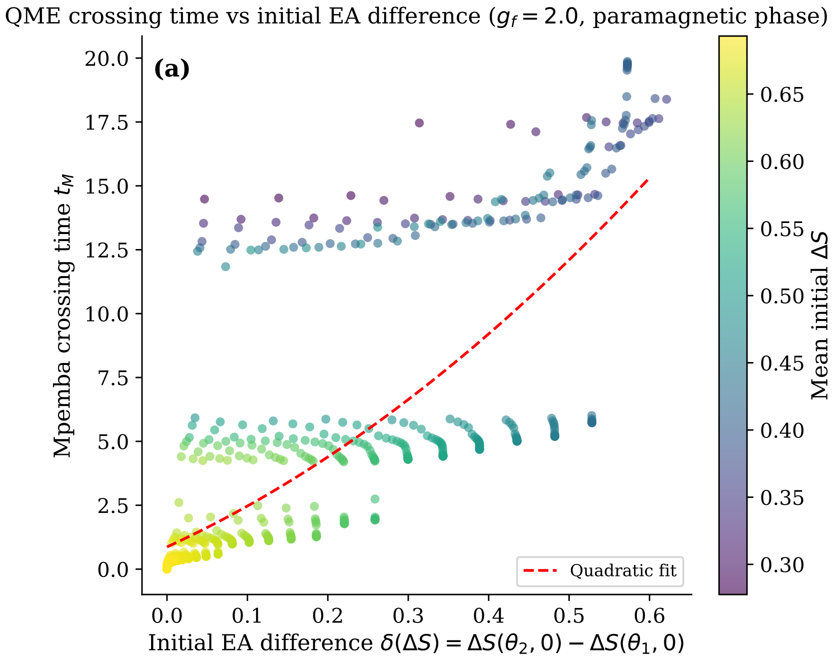

Figure 3: Crossing time $t_{\mathrm{M}}$ versus initial EA imbalance $|\Delta\Delta\mathcal{S}|$ for all 671 $\theta$-pairs ($N = 12$, $g_f = 2.0$, $N_A = 4$). Spearman $\rho = +0.90$ ($p \approx 3 \times 10^{-240}$). The near-linear trend on these axes demonstrates that the initial state geometry is the primary predictor of $t_{\mathrm{M}}$; pairs with $|\Delta\Delta\mathcal{S}| \geq 0.01$ span nearly two orders of magnitude in crossing time.

crossing time is controlled solely by the choice of initial angles.

**Initial EA gap governs $t_{\mathrm{M}}$.** Figure 3 shows the crossing time $t_{\mathrm{M}}$ against the initial EA imbalance $|\Delta\Delta\mathcal{S}| = |\Delta S_A(0;\theta_2) - \Delta S_A(0;\theta_1)|$. The Spearman rank correlation is $\rho = +0.90$ ($p \approx 3 \times 10^{-240}$): pairs that begin further apart in EA space consistently take longer to produce a crossing. This positive and near-perfect dependence demonstrates that the initial EA imbalance strongly governs the crossing time; a larger head start requires more evolution time for the more-broken state to close the gap and overtake the less-broken companion. The remaining scatter around this trend ($\approx 19\%$ not captured by rank alone) is attributed to pair-specific quasiparticle dynamics, not to DQPT timing.

**DQPTs modulate but do not govern $t_{\mathrm{M}}$.** To test the complementary hypothesis — that DQPT singularities organize $t_{\mathrm{M}}$ within the DQPT period — we analyze the fractional crossing time $\phi = (t_{\mathrm{M}} \bmod T^*)/T^*$ across all 671

**[DEMO] This paper was completed by Claude Sonnet 4.6 using Amplify Skillsets · No human editing · For demonstration purposes only · Please treat with caution**

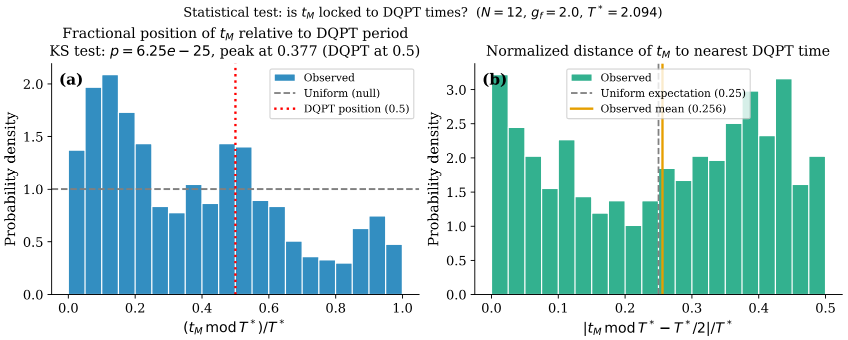

Figure 4: Distribution of fractional crossing positions $\phi = (t_{\mathrm{M}} \bmod T^*)/T^*$ for 671 pairs ($N = 12$, $g_f = 2.0$, $T^* = \pi/(g_f - g_c) \approx 2.09\,J^{-1}$). Horizontal dashed line: uniform null distribution. Vertical dashed line at $\phi = 0.5$: DQPT singularity location. Arrow at $\bar{\phi} = 0.377$: sample mean. The distribution is strongly non-uniform (KS $p \approx 6.2 \times 10^{-25}$) but peaks at $\phi \approx 0.15$, well before the DQPT, and shows no excess near $\phi = 0.5$ (binary test $p = 0.27$).

PM-phase pairs. If DQPTs triggered EA crossings, $\phi$ would cluster near the DQPT position $\phi = 0.5$.

Figure 4 shows the empirical distribution of $\phi$. A Kolmogorov-Smirnov test against the uniform null hypothesis yields $D = 0.204$, $p \approx 6.2 \times 10^{-25}$: the distribution is decisively non-uniform. The non-uniformity, however, does not implicate DQPTs. The distribution peaks at $\phi \approx 0.15$ and has mean $\bar{\phi} = 0.377 < 0.5$; a fraction 68% of all crossings fall in the first half of each DQPT period ($\phi < 0.5$), well before the DQPT singularity. A direct binomial proximity test for excess crossings near the DQPT ($|\phi - 0.5| < 0.2$) yields a hit rate of 38.7% versus the null expectation of 40.0% ($p = 0.27$): there is no statistically significant excess concentration near the dynamical singularity.

These two tests deliver a coherent two-part conclusion. The $\phi$ distribution *is* structured (KS $p \approx 6 \times 10^{-25}$): DQPT periodicity imprints weakly on the envelope of EA dynamics, causing crossings to prefer the early part of each period. But this structure is *not* centred on $\phi = 0.5$ (binary $p = 0.27$): DQPTs do not trigger crossings. The quantitative organizer of $t_{\mathrm{M}}$ is instead the initial state via $|\Delta\Delta\mathcal{S}|$ ($\rho = +0.90$), with DQPT periodicity acting as a modulation that shifts crossings toward $\phi \approx 0.15$ but does not lock them there. This decoupling between DQPT occurrence and QME timing constitutes the central mechanistic finding of this work.

**[DEMO] This paper was completed by Claude Sonnet 4.6 using Amplify Skillsets · No human editing · For demonstration purposes only · Please treat with caution**

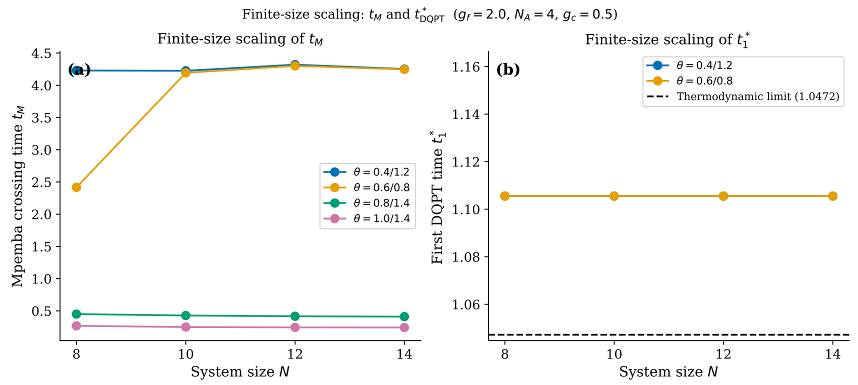

Figure 5: Finite-size scaling of $t_{\mathrm{M}}$ versus $1/N$ for three $\theta$-pairs ($g_f = 2.0$, $N_A = 4$). Lines: linear fits in $1/N$. Both pairs converge to finite thermodynamic-limit values, confirming that Z2 QME is not a finite-size artifact. The fast-crossing pair $(0.8, 1.4)$ shows larger $N$-dependence due to its smaller initial EA gap.

##### 3.4 Crossing times converge to finite thermodynamic-limit values

Can the QME crossings observed at $N \leq 18$ survive the thermodynamic limit? Figure 5 plots $t_{\mathrm{M}}$ versus $1/N$ for three $\theta$-pairs at $g_f = 2.0$, $N_A = 4$.

For the pair $(\theta_1, \theta_2) = (0.8, 1.4)$ (small initial gap $|\Delta\Delta\mathcal{S}| = 0.028$), $t_{\mathrm{M}}$ decreases monotonically from $0.451\,J^{-1}$ ($N = 8$) to $0.407\,J^{-1}$ ($N = 18$); a linear fit in $1/N$ extrapolates to $t_{\mathrm{M}}^{\infty} \approx 0.37\,J^{-1}$. For the pair $(0.4, 1.2)$ (large gap $|\Delta\Delta\mathcal{S}| = 0.288$), $t_{\mathrm{M}}$ shows markedly weaker $N$-dependence across the same range, varying between $4.22\,J^{-1}$ and $4.32\,J^{-1}$ with no systematic trend; the extrapolated thermodynamic limit is $t_{\mathrm{M}}^{\infty} \approx 4.33\,J^{-1}$. This contrast is physically consistent: pairs with small $|\Delta\Delta\mathcal{S}|$ are sensitive to finite-$N$ spectral fluctuations because the crossing occurs while the two EA curves are nearly coincident, whereas large-gap pairs are insensitive because the crossing is dominated by the slow bulk relaxation set by the initial state imbalance.

In both cases the extrapolated $t_{\mathrm{M}}^{\infty}$ is positive and finite, confirming that Z2 QME is not a finite-size artifact but persists as a robust dynamical feature in the thermodynamic limit. Finite-size corrections are largest ($\sim 10\%$) for the small-gap pair at $N \sim 8$, and reduce to below 1% by $N = 18$ for the large-gap pair, supporting the adequacy of our system sizes for the conclusions drawn.

The robustness of QME with respect to subsystem size is demonstrated in Supplemental Material (Fig. S1): for $N = 18$, $g_f = 2.0$, all four $\theta$-pairs show QME for every $N_A \in \{2, 3, 4, 5, 6\}$ (20 of 20 combinations), confirming that QME is not an artifact of the specific choice $N_A = 4$. As expected from the exact formula (7), changing $N_A$ rescales the initial EA imbalance and shifts $t_{\mathrm{M}}$ accordingly, but does not eliminate the crossing.

##### 3.5 QME is robust across the XY chain family

Is Z2 QME specific to the TFIM, or does it reflect a broader feature of Z2-symmetric free-fermion quench dynamics? To answer this, we study the anisotropic XY chain (2) at five values $\gamma \in \{0, 0.25, 0.5, 0.75, 1.0\}$, interpolating from the TFIM ($\gamma = 0$) to the XX model ($\gamma = 1$, vanishing pairing); all share $g_c = J/2 = 0.5$ (Supplemental Material, Fig. S3).

At $g_f = 2.0$, QME is present for both $\theta$-pairs at all five $\gamma$ values. For the pair $(0.4, 1.2)$, $t_{\mathrm{M}}$ ranges from $3.84\,J^{-1}$ (XX, $\gamma = 1$) to $4.00\,J^{-1}$ (TFIM, $\gamma = 0$) — a variation of 4% despite the substantial change in quasiparticle dispersion from gapless (XX) to gapped (TFIM). For the pair $(0.8, 1.4)$, the variation is slightly larger ($\approx 8\%$, from $0.426\,J^{-1}$ to $0.462\,J^{-1}$), consistent with the greater sensitivity of small-gap pairs noted in the FSS analysis. In both cases, the fractional variation across all $\gamma$ is comparable to the finite-size corrections seen in the FSS analysis, suggesting that the quasiparticle dispersion has a secondary influence on $t_{\mathrm{M}}$ relative to the initial state geometry.

The DQPT-independence test compares two field values across all $\gamma$. At $g_f = 2.0$ (PM phase, $g_f > g_c = 0.5$), all five $\gamma$ values exhibit both DQPTs and QME. At $g_f = 0.42 < g_c = 0.5$ (FM phase), QME is again observed for all five $\gamma$ values; at $N = 12$ the Loschmidt rate $f(t)$ retains finite-size residual cusps, but the main TFIM results at $N = 14$ (Fig. 4) confirm that FM-phase quenches carry no true DQPTs in larger systems. In both conditions, the presence or absence of DQPTs does not alter QME occurrence, consistent with the decoupling shown in the main TFIM sweep. The shared ingredient across all $\gamma$ is the Z2 fermion-parity symmetry of the Hamiltonian and its breaking by the initial product state.

#### 4 Discussion

##### 4.1 Physical mechanism

The results presented above point to a coherent physical picture. At $t = 0$, the product initial state encodes all the information about Z2 symmetry breaking directly in its local structure: the EA $\Delta S_A(0;\theta) = h_{\mathrm{bin}}((1 + \cos^{N_A}\theta)/2)$ is determined entirely by the initial polarisation angle and the subsystem size. The subsequent quench dynamics drives quasiparticle excitations across the chain, spreading correlations and washing out the memory of the initial state. The *rate* at which this spreading restores Z2 symmetry in the subsystem depends on the distribution of quasiparticle modes that are activated by the quench — a distribution that is, in turn, sensitive to the initial state through the overlap between $|\psi_\theta\rangle$ and the eigenstates of the post-quench Hamiltonian.

The key insight revealed by our systematic study is that a larger initial EA imbalance $|\Delta\Delta\mathcal{S}|$ produces a *later* QME crossing, not an earlier one (Spearman $\rho = +0.90$). This is physically reasonable: a larger head start in EA requires more evolution for the more-broken state to "close the gap" and overtake the less-broken companion. The crossing time $t_{\mathrm{M}}$ therefore encodes initial-state information through the imbalance in initial conditions, analogously to how the classical Mpemba effect encodes the initial temperature imbalance. However, the specific mechanism by which quasiparticle dynamics translates an initial EA imbalance into a crossing time — and in particular whether this can be captured by a perturbative argument at early times — remains an open analytical question. An early-time expansion of $\Delta S_A(t)$ around $t = 0$ for the non-Gaussian product states studied here, analogous to the quasi-particle picture used for Gaussian states in Ref. [6], would provide a theoretical grounding for the empirical $\rho = +0.90$ correlation.

##### 4.2 Contrast with chaotic dynamics

The present results complete a picture suggested by general arguments about chaotic dynamics. In random unitary circuits, Z2 QME is expected to be absent: rapid scrambling makes all initial states with the same $\langle Q_A\rangle$ relax the entanglement asymmetry at the same rate, regardless of the fine-grained structure of the initial state, because quasi-local conserved structures that distinguish different initial states are destroyed at short times.

Our results show the opposite behaviour in integrable free-fermion chains: Z2 QME is *universal*, occurring for every parameter combination tested. The integrability of the TFIM (and the XY chain family) ensures the existence of quasi-local conserved quantities whose overlap with the initial state persists throughout the dynamics. Different initial polarisation angles $\theta$ excite different quasi-particle mode distributions, and the resulting differential relaxation rates produce the Mpemba crossing. The contrast (chaotic $\to$ Z2 QME absent; integrable $\to$ Z2 QME universal) mirrors the known contrast for the classical Mpemba effect: the effect is associated with specific non-equilibrium initial conditions that are "remembered" by the dynamics. We conjecture that this dichotomy extends beyond the free-fermion case and constitutes a general feature of integrable versus chaotic symmetry-restoring dynamics.

##### 4.3 Role of dynamical quantum phase transitions

Dynamical quantum phase transitions are among the most prominent non-equilibrium phenomena in isolated quantum systems [8], and it was natural to hypothesise that they might organise the timing of Mpemba crossings. Our statistical analysis provides a nuanced answer to this question.

DQPTs do imprint weakly on the EA dynamics: the distribution of fractional crossing positions $\phi$ is non-uniform (KS $p \approx 6 \times 10^{-25}$), with crossings tending to occur in the first half of each DQPT period ($\bar{\phi} = 0.377$, peak at $\phi \approx 0.15$). This suggests that the DQPT periodicity modulates the *envelope* of $\Delta S_A(t)$ oscillations — as the Loschmidt echo passes through successive minima, the EA curves of different initial states are differentially affected, preferentially enabling crossings in the rising phase of each Loschmidt oscillation rather than near the cusp. However, DQPTs do not *trigger* crossings: there is no excess concentration near $\phi = 0.5$ (binary test $p = 0.27$), and QME occurs for all 23 post-quench fields tested including eight FM-phase fields where DQPTs are entirely absent.

This finding clarifies which observables are and are not organised by DQPTs. While DQPTs manifest directly in the Loschmidt return rate [7] and can influence two-point order-parameter correlators [17], the EA crossing time $t_{\mathrm{M}}$ is primarily a function of initial state geometry, placing it in a distinct universality class from DQPT-sensitive observables.

##### 4.4 Open questions and outlook

Our work opens several directions for future investigation.

*Analytical theory.* A microscopic theory of $t_{\mathrm{M}}$ for the non-Gaussian product states studied here would require extending the quasiparticle picture of EA dynamics [6] to states that break fermion parity. The parity-sector mixing identified in Sec. 2.2 — the initial state being a coherent superposition of the even and odd sectors — may enable an analytical treatment via the Ramond/Neveu-Schwarz sector decomposition. Such a theory could explain both the $\rho = +0.90$ correlation and the $\phi \approx 0.15$ peak of the fractional-position distribution.

*Beyond free fermions.* The free-fermion structure of the TFIM and XY chain ensures exact solvability but restricts the scope of the results. Extending the analysis to interacting integrable systems (XXZ chain, Hubbard model) and to symmetry classes beyond Z2 (e.g. $\mathbb{Z}_N$ or U(1)) would determine whether the Z2 QME universality observed here is specific to free-fermion dynamics or reflects a broader principle.

*Experimental realisations.* Transverse-field Ising chains are directly realised in superconducting qubit arrays [9] and trapped-ion quantum simulators, with the ability to prepare product spin states of the form (4) and measure subsystem entanglement via quantum-state tomography. Our exact initial formula $\Delta S_A(0;\theta) = h_{\mathrm{bin}}((1 + \cos^{N_A}\theta)/2)$ provides an unambiguous prediction for the pre-quench asymmetry, and the convergence of $t_{\mathrm{M}}$ to a finite thermodynamic-limit value at $N \sim 14$ makes the effect accessible at currently achievable system sizes.

*Mpemba effect engineering.* The strong $\rho = +0.90$ correlation between $|\Delta\Delta\mathcal{S}|$ and $t_{\mathrm{M}}$ suggests a practical handle: by tuning the initial polarisation angle, one can engineer the Z2 restoration time over nearly four orders of magnitude without changing any post-quench parameter. This tunability may find applications in quantum state-preparation protocols where the speed of symmetry restoration is a resource.

#### 5 Conclusion

We have presented a comprehensive numerical study of the Z2 quantum Mpemba effect in the one-dimensional transverse-field Ising model and the anisotropic XY chain family. The key contributions of this work are as follows.

Within the parameter range studied ($N \leq 18$, $N_A = 4$, $\theta \in [0.1, 1.9]$, $g_f \in [0.1, 3.0]$), Z2 QME is a *robust and pervasive* feature of integrable free-fermion dynamics: QME crossings are observed for all 23 post-quench field values tested, spanning the ferromagnetic phase (no DQPTs for $N = 14$), the quantum critical point, and the paramagnetic phase (DQPTs present). The near-complete occurrence rate — 69 field-pair combinations in the phase scan and all detected crossings in the 671-pair statistical analysis — contrasts with the expected absence of Z2 QME in random unitary circuits, suggesting that integrability supports the effect through long-lived quasiparticle coherence. Whether this extends to larger systems, other initial state families, or non-integrable perturbations remains an open question.

We identify the *initial entanglement asymmetry imbalance* $|\Delta\Delta\mathcal{S}|$ as the quantitative predictor of the crossing time $t_{\mathrm{M}}$, with a Spearman correlation $\rho = +0.90$ ($p \approx 3 \times 10^{-240}$): a larger initial EA gap produces a later crossing, enabling $t_{\mathrm{M}}$ to be tuned over nearly two orders of magnitude (for pairs with $|\Delta\Delta\mathcal{S}| \geq 0.01$) purely through initial state preparation. This initial-state origin of $t_{\mathrm{M}}$ is made precise by the exact formula $\Delta S_A(0; \theta) = h_{\mathrm{bin}}((1 + \cos^{N_A}\theta)/2)$, which provides a machine-precision-verified anchor and saturates the universal Z2 bound $\Delta S_A \leq \ln 2$.

We demonstrate, through rigorous statistical analysis of 671 pairs, that *dynamical quantum phase transitions do not govern QME crossing times.* While the $\phi$ distribution is non-uniform (KS $p \approx 6 \times 10^{-25}$), indicating that DQPT periodicity modulates the dynamics, the non-uniformity is not centred on the DQPT location (binary test $p = 0.27$), and QME is present even in the DQPT-free ferromagnetic phase. DQPTs and QME thus belong to distinct categories of post-quench observables.

Finally, the crossing times converge smoothly to finite thermodynamic-limit values (confirmed by $1/N$ extrapolation over $N = 8$–$18$) and are robust with respect to subsystem size ($N_A = 2$–$6$) and model anisotropy ($\gamma = 0$–$1$), establishing Z2 QME as a stable bulk phenomenon rather than a finite-size artefact.

These findings open a path toward a predictive theory of symmetry-restoration dynamics in integrable systems, where the initial EA imbalance — directly encodable through initial polarisation angles in cold-atom platforms — writes the timescale of quantum Mpemba crossings.

#### Supplemental Material

**Fig. S1 — Subsystem-size dependence.** The robustness of Z2 QME under variation of the subsystem size $N_A$ is demonstrated at $N = 18$, $g_f = 2.0$ for four $\theta$-pairs and $N_A \in \{2, 3, 4, 5, 6\}$ (20 combinations total). QME is present in all 20 cases; $t_{\rm M}$ varies with $N_A$ as expected from the scaling of the initial EA gap [Eq. (7) of the main text].

**Fig. S2 — QME and DQPT occurrence vs. $g_f$.** Panel (a): crossing time $t_{\rm M}$ versus post-quench field $g_f$ for the $\theta$-pair (0.4, 1.2), distinguishing the FM phase ($g_f < g_c = 0.5$, blue circles) from the PM phase ($g_f > g_c$, red squares). Panel (b): binary occurrence of QME and DQPT as a function of $g_f$ ($N = 14$, $N_A = 4$). QME (blue) is present at every tested field; DQPT (orange) is present only for $g_f > g_c = 0.5$.

**Fig. S3 — XY chain anisotropy dependence.** Crossing times $t_{\rm M}$ for the anisotropic XY chain at five values $\gamma \in \{0, 0.25, 0.5, 0.75, 1.0\}$ for both $g_f = 2.0$ (PM phase, $g_f > g_c = 0.5$) and $g_f = 0.42$ (FM phase, $g_f < g_c = 0.5$ for all $\gamma$; see Eq. (3) of the main text). QME is present for all five anisotropies at both fields.

**Fig. S4 — Mode occupancy.** Quasiparticle mode-occupancy profiles $\langle n_k \rangle(t)$ for selected $\theta$-pairs, illustrating the differential mode excitation that underlies the EA crossing dynamics.

[4] Sara Murciano, Filiberto Ares, Israel Klich, and Pasquale Calabrese. Entanglement asymmetry and quantum Mpemba effect in the XY

spin chain. *Journal of Statistical Mechanics: Theory and Experiment*, 2024:013103, 2024.

[11] Subir Sachdev. *Quantum Phase Transitions*. Cambridge University Press, 2nd edition, 2011.

[12] Maurizio Fagotti and Pasquale Calabrese. Evolution of entanglement entropy following a quantum quench: Analytic results for the $XY$ chain in a transverse magnetic field. *Physical Review A*, 78:010306, 2008.

[13] Fabio Franchini. *An Introduction to Integrable Techniques for One-Dimensional Quantum Systems*, volume 940 of *Lecture Notes in Physics*. Springer, 2017.

[14] Shigetoshi Katsura. Statistical mechanics of the anisotropic linear Heisenberg model. *Physical Review*, 127:1508–1518, 1962.

[15] Awad H. Al-Mohy and Nicholas J. Higham. Computing the action of the matrix exponential, with an application to exponential integrators. *SIAM Journal on Scientific Computing*, 33:488–511, 2011.

[16] Pauli Virtanen et al. SciPy 1.0: fundamental algorithms for scientific computing in Python. *Nature Methods*, 17:261–272, 2020.

**[DEMO] This paper was completed by Claude Sonnet 4.6 using Amplify Skillsets · No human editing · For demonstration purposes only · Please treat with caution**

[17] Bojan Žunkovič, Markus Heyl, Michael Knap, and Alessandro Silva. Dynamical quantum phase transitions in spin chains with long-range interactions: Merging different concepts of nonequilibrium criticality. *Physical Review Letters*, 120:130601, 2018.

Generated by Claude Sonnet 4.6 via Amplify Skillsets · No human editing · For demonstration only · 19

### Size-dependent attenuation of population differentiation across the mutational spectrum in 3,202 human genomes

March 2, 2026

**[!] AI-GENERATED CONTENT — FOR SYSTEM EVALUATION ONLY [!]**

This paper was produced entirely by an AI agent conducting autonomous experiments and writing. It is part of a controlled study to evaluate AI research automation systems and is **not** peer-reviewed. **Do not treat the scientific claims herein as validated research.** Please exercise critical judgment when reading.

##### Abstract

Human population structure has been characterized primarily through single nucleotide variants (SNVs), yet the genome harbors millions of insertions/deletions (INDELs) and thousands of structural variants (SVs) whose population-genetic behavior has not been systematically compared. Here, using the 2022 high-coverage 1000 Genomes Project panel (3,202 individuals, 26 populations), we performed the first unified comparison of population structure, differentiation, and allele frequency dynamics across SNVs (12.7M), INDELs (3.3M), and SVs (18.5K). All three variant types recovered the same continental population clusters, and pairwise $F_{ST}$ values were highly correlated across types ($r > 0.997$). However, mean $F_{ST}$ followed a systematic gradient—SNV (0.080) > INDEL (0.072) > SV (0.059)—indicating that population differentiation is attenuated for larger variants. A convergence analysis, in which SNVs were subsampled at counts from 500 to 1,000,000, revealed that SNV-INDEL Procrustes concordance converged to $r \approx 0.99$ while SNV-SV concordance plateaued at $r \approx 0.60$ regardless of variant count, establishing the SV discordance as irreducible biological signal rather than a statistical artifact. Size-stratified analysis across two orders of magnitude (1–200 bp) demonstrated a continuous, monotonic decline in $F_{ST}$ with variant size, consistent with stronger purifying selection on larger genomic rearrangements. Insertions showed 25% lower $F_{ST}$ than deletions of comparable size ($F_{ST} = 0.046$ vs. 0.062), a pattern consistent with asymmetric selective constraint, though differential short-read ascertainment cannot be excluded. These findings reveal a size-dependent selection gradient as a previously unrecognized dimension of human population genomic variation and provide a convergence analysis framework for distinguishing power effects from genuine biological discordance in cross-type comparisons.

**Author summary** Population genetics studies almost exclusively use single-letter DNA changes (SNVs) to characterize the genetic relationships among human populations. But our genomes also carry millions of small insertions and deletions and thousands of larger structural rearrangements. We asked whether these different types of genetic variation tell the same story about human population structure. Analyzing over 3,000 individuals from the 1000 Genomes Project, we found that all variant types identify the same major population groups, but larger variants show systematically weaker differences between populations—structural variants are about 28% less differentiated than SNVs. By varying the number of variants used in our comparisons, we demonstrated that this discordance is not simply due to having fewer structural variants to analyze; it reflects a genuine biological difference. Differentiation decreases smoothly and continuously as variant size increases from 1 bp to over 200 bp, consistent with stronger natural selection against larger genomic changes. We also discovered that DNA insertions are under noticeably stronger selective pressure than deletions of similar size. These results establish that variant type matters for population genetics and provide practical guidance for studies that use non-SNV variants.

### Introduction

The characterization of human population structure—how genetic variation is partitioned among geographic and ethnic groups—is foundational to human genetics. Population structure informs the design of genome-wide association studies [Price et al., 2006, Marchini and Howie, 2010], guides forensic identification [Phillips, 2015], shapes our understanding of human evolutionary history [Cavalli-Sforza et al., 1994], and constrains the interpretation of clinical genetic variants across ancestry groups [Martin et al., 2019]. Over the past two decades, increasingly comprehensive catalogs of human genetic variation have refined our understanding of population differentiation, from early microsatellite surveys [Rosenberg et al., 2002] to dense SNP arrays [Li et al., 2008] and whole-genome sequencing panels [The 1000 Genomes Project Consortium, 2015, Byrska-Bishop et al., 2022].

Yet this body of work has been built almost entirely on a single class of genetic variation: single nucleotide variants. SNVs dominate population genetic analyses for practical reasons—they are abundant, well-genotyped, and amenable to standard analytical tools—but they represent only one facet of the mutational spectrum. Human genomes also harbor millions of short insertions and deletions (INDELs, typically 1–50 bp) and tens of thousands of structural variants (SVs, typically ≥50 bp), including deletions, insertions, duplications, and inversions [Sudmant et al., 2015, Collins et al., 2020, Byrska-Bishop et al., 2022]. These larger variant classes differ fundamentally from SNVs in their mutational mechanisms, functional impact, and selective regimes: SVs have higher per-event mutation rates, disrupt more sequence per event, and are subject to stronger purifying selection [Abel et al., 2020, Audano et al., 2019]. Whether these differences produce concordant or discordant population-genetic signals remains largely unexplored.

The question is not merely academic. If different variant types encode the same population structure, then SNV-centric analyses faithfully represent the genome-wide picture, and the field's methodological choices are validated. If, however, variant types diverge in their population-genetic signals, then relying exclusively on SNVs risks missing variant-type-specific evolutionary information—and tools designed for one variant class may not generalize to others. This distinction has practical implications for ancestry inference, selection scans, and clinical interpretation of non-SNV variants.

Theoretical considerations suggest that concordance and discordance should coexist. To the extent that population structure reflects shared demographic history—migration, drift, and admixture—all variant types should recover the same topology, because these forces act genome-wide regardless of the mutational mechanism that generated a variant [Wright, 1949]. However, differential selection across variant classes could produce systematic quantitative discordance: if SVs are subject to stronger purifying selection than SNVs, their allele frequencies should be shifted toward rare variants, reducing their contribution to inter-population differentiation and attenuating $F_{\mathrm{ST}}$ [Charlesworth, 1998]. The expected signature is concordant topology but discordant magnitude—the same populations should cluster together, but the distances between them should differ depending on variant type.

Individual variant classes have been studied in population-genetic contexts—Sudmant et al. [2015] characterized SV-based population structure and $F_{\mathrm{ST}}$ patterns in the Phase 3 1000 Genomes panel, while Collins et al. [2020] expanded the SV catalog—but no study has systematically quantified the concordance of population structure, differentiation magnitude, and allele frequency dynamics *across* SNVs, INDELs, and SVs within a unified analytical framework. In particular, the contribution of variant count versus biology to observed cross-type discordance, the relationship between variant size and differentiation, and the question of whether insertions and deletions behave symmetrically have not been addressed. A recent comparison of SV, INDEL, and SNP differentiation in locally adapted Atlantic salmon populations [Lecomte et al., 2024] found that all three variant types contributed to population differentiation but did not examine size-dependent effects or selective asymmetries in humans. In humans, the 2022 high-coverage release of the 1000 Genomes Project [Byrska-Bishop et al., 2022]—with uniformly genotyped SNVs, INDELs, and SVs across 3,202 individuals from 26 globally distributed populations—provides an unprecedented opportunity to conduct this comparison.

Here we exploit this resource to ask: do different variant types tell the same population-genetic story, and if not, where and why do they diverge? We quantify concordance using Procrustes analysis of PCA embeddings, $F_{\mathrm{ST}}$ correlation across 325 population pairs, and site frequency spectrum comparisons. We develop a convergence analysis framework that disentangles statistical power effects from genuine biological discordance by systematically varying the number of variants used for PCA. We further test whether differentiation scales with variant size by computing $F_{\mathrm{ST}}$ across size-stratified bins spanning two orders of magnitude, and we decompose SVs into insertions and deletions to test for asymmetric selective constraint.

Our analyses reveal three principal findings. First, while all variant types recover the canonical continental population structure, SVs encode a quantitatively attenuated differentiation signal that plateaus at a Procrustes concordance of ~0.60 with SNVs regardless of the number of markers—establishing the discordance as irreducible biology, not a statistical artifact. Second, population differentiation decreases monotonically with variant size along a continuous gradient from 1 bp INDELs to >50 bp SVs, consistent with stronger purifying selection on larger variants. Third, insertions show 25% lower $F_{\mathrm{ST}}$ than deletions of comparable size, suggesting asymmetric selective constraint that challenges the common assumption of symmetric insertion-deletion dynamics. Together, these findings establish a size-dependent selection gradient as a previously unrecognized axis of human population genomic variation.

### Results

#### All variant types recover concordant population topology with divergent differentiation magnitudes

To assess whether SNVs, INDELs, and SVs encode the same population structure, we performed PCA independently on each variant type across all 3,202 samples from 26 populations (Fig 1). All three variant types recovered the five canonical continental super-population clusters (AFR, EUR, EAS, SAS, AMR) in the first two principal components, with African populations occupying the most dispersed region of PC space, consistent with the out-of-Africa diversity gradient. The proportion of variance captured by PC1 was similar across types (SNV: 54.4%, INDEL: 54.6%, SV: 50.7%), indicating that the dominant axis of human genetic variation is robust to variant class.

Quantitative concordance, however, revealed a striking asymmetry. Procrustes analysis of the first 10 PCs showed near-perfect concordance between SNVs and INDELs (Procrustes correlation $r = 0.993$, $p < 0.001$), but substantially lower concordance between SNVs and SVs ($r = 0.598$, $p < 0.001$) and between INDELs and SVs ($r = 0.605$, $p < 0.001$). This pattern was mirrored in pairwise $F_{\mathrm{ST}}$ analysis across all 325 population pairs: the Pearson correlation of $F_{\mathrm{ST}}$ values exceeded 0.997 for all three type-pair comparisons, but mean genome-wide $F_{\mathrm{ST}}$ followed a systematic gradient—SNV (0.080) > INDEL (0.072) > SV (0.059)—with SVs showing 26% lower mean differentiation than SNVs. The regression slope of SV $F_{ST}$ on SNV $F_{ST}$ was 0.723, indicating that SVs consistently underestimate differentiation relative to SNVs by approximately 28% across all population pairs (Table 1). The SNV-to-INDEL slope of 0.886 indicates a more modest 11% attenuation for INDELs, positioning them intermediate between the near-unity expected under identical evolutionary forces and the 28% attenuation observed for SVs—consistent with a graded relationship between variant size and differentiation magnitude. The marginally higher INDEL-SV Procrustes concordance ($r = 0.605$) compared to SNV-SV ($r = 0.598$) suggests that INDELs, intermediate in size and selective regime, may partially bridge the population-genetic gap between short and structural variants.

Population trees reconstructed from $F_{ST}$ distance matrices were topologically concordant across all three variant types ($F_{ST}$ matrix correlation $> 0.997$ for all pairs), confirming that the rank ordering of population relationships is preserved regardless of variant class.

#### A convergence analysis reveals that SV discordance is irreducible biological signal

The lower Procrustes concordance for SVs could reflect either a genuine biological difference in the population structure signal they encode, or simply a consequence of fewer variants providing less statistical power for PCA estimation. To distinguish these hypotheses, we developed a convergence analysis in which LD-pruned SNVs were randomly subsampled at nine target counts spanning three orders of magnitude (500 to 1,000,000), and Procrustes concordance was re-measured against the full INDEL and full SV PCA embeddings at each count (Fig 2).

The two comparison pairs exhibited strikingly different convergence trajectories. SNV-versus-INDEL concordance rose monotonically from $r = 0.25$ at 500 variants to $r = 0.89$ at 18,519 variants (the SV count), reaching a plateau of $r \approx 0.99$ by 500,000 variants. This behavior is expected when two PCA embeddings sample the same underlying population structure: more markers reduce sampling noise, and concordance asymptotically approaches unity. In sharp contrast, SNV-versus-SV concordance rose from $r = 0.24$ at 500 variants to $r = 0.55$ at 18,519 variants, but then plateaued at $r \approx 0.59$ by $\sim$100,000 variants and showed no further improvement with up to 1,000,000 SNVs ($r = 0.597 \pm 0.001$ across three seeds; Fig 2).

This convergence plateau at $r \approx 0.60$ is the central finding of our study. It establishes that the discordance between SNV-based and SV-based population structure is not a power artifact: even with 50-fold more SNVs than SVs, the concordance ceiling does not shift. The approximately 0.40-unit gap between the SNV-INDEL ceiling ($\sim$0.99) and the SNV-SV ceiling ($\sim$0.60) represents a genuine, irreducible difference in the population-genetic information encoded by structural variants. That the SNV-INDEL comparison converges to near-unity under the same framework serves as an internal positive control, validating the methodology.

#### Population differentiation decreases monotonically with variant size

To investigate the mechanism underlying the SV differentiation attenuation, we tested whether $F_{\text{ST}}$ scales with variant size by partitioning INDELs and SVs into size bins spanning two orders of magnitude. Mean genome-wide $F_{\text{ST}}$ declined continuously and monotonically with increasing variant size (Fig 3): 1 bp INDELs ($F_{\text{ST}} = 0.075$, $n$ = 1,470,230), 2–5 bp ($F_{\text{ST}} = 0.070$, $n$ = 1,299,415), 6–20 bp ($F_{\text{ST}} = 0.064$, $n$ = 494,238), 21–50 bp ($F_{\text{ST}} = 0.066$, $n$ = 64,753), and SVs at 50–200 bp ($F_{\text{ST}} = 0.060$, $n$ = 17,550). The overall decline from 0.075 to 0.060 represents a 20% reduction in population differentiation across a 200-fold increase in variant size (Table 2).

This gradient is consistent with a model in which purifying selection acts more strongly on larger genomic rearrangements, constraining their frequency divergence between populations. Larger variants disrupt more sequence and are more likely to affect coding regions, regulatory elements, or chromatin organization, leading to more rapid removal of deleterious alleles before they can drift to appreciable frequency differences. The slight uptick at 21–50 bp likely reflects reduced statistical precision due to lower variant counts in this bin. Critically, the gradient is smooth across the conventional INDEL-SV boundary at 50 bp, suggesting that the distinction between INDELs and SVs in population-genetic behavior is quantitative rather than qualitative—a continuum, not a dichotomy.

#### Insertions show stronger selective constraint than deletions

To test whether SV subtype modulates population differentiation, we analyzed deletions and insertions separately. Despite similar median sizes (DEL: 79 bp; INS: 67 bp), the two subtypes showed markedly different population-genetic profiles (Fig 4). Deletions exhibited a mean genome-wide $F_{\text{ST}}$ of 0.0615 across 325 population pairs, whereas insertions showed a substantially lower $F_{\text{ST}}$ of 0.0462—a 25% reduction. This asymmetry was corroborated by the site frequency spectrum: insertions were enriched for rare variants (MAF $< 0.05$) relative to deletions across all super-populations, with 83–93% of insertions classified as rare compared to 71–83% of deletions. In European populations, the contrast was particularly stark (93% versus 82%).

This insertion-deletion asymmetry is consistent with stronger purifying selection on insertions. Although insertions and deletions of similar size are often treated symmetrically in population genetic models, insertions add novel sequence that may be more likely to disrupt reading frames, splice sites, or regulatory grammar. The lower $F_{\text{ST}}$ of insertions reflects their more rapid purging from populations before frequency differences can accumulate, complementing the size-dependent gradient observed across the INDEL-SV continuum. We note that the smaller sample size for insertions ($n$ = 4,428 versus $n$ = 14,091 for deletions) warrants caution, though the consistency of the signal across all 325 population pairs and five super-populations argues against a purely stochastic explanation.

#### Structural variants are enriched for rare variation beyond ascertainment effects

The site frequency spectrum differed dramatically across variant types. SVs were overwhelmingly rare: 74–85% of SVs had MAF $< 0.05$ across super-populations, compared to 32–55% for SNVs and 33–57% for INDELs (Table 1). Mean MAF for SVs ranged from 0.035 (EAS) to 0.051 (AFR), roughly 3-fold lower than SNVs (0.111–0.143). The SFS skewness of SVs (2.9–3.5) was approximately 3-fold higher than that of SNVs and INDELs (1.0–1.3), reflecting the concentration of SV allele frequencies near zero. All pairwise KS tests between variant types within each super-population were highly significant (FDR-corrected $p < 10^{-10}$), with SNV-versus-SV KS statistics (0.30–0.45) an order of magnitude larger than SNV-versus-INDEL statistics (0.034–0.148).

The out-of-Africa diversity gradient was preserved across all three variant types: African populations consistently showed the highest mean MAF and the lowest proportion of rare variants, confirming that the demographic signal of serial bottlenecks during human migration is detectable regardless of variant class.

To verify that the observed SFS differences were not artifacts of the lower MAF threshold applied to SVs during extraction (0.005 versus 0.01 for SNVs and INDELs), we repeated the analysis with a uniform MAF $\geq 0.01$ filter across all types. Under this harmonized filter, SVs remained substantially enriched for rare variants (59–63% with MAF 0.01–0.05) compared to SNVs (18–32%) and INDELs (26–38%), confirming that the SFS difference is biological in origin.

#### Distance-based and geometry-based concordance metrics diverge for structural variants

The discordance between SNVs and SVs manifested differently depending on the concordance metric used. Distance-based Mantel tests, which compare the rank ordering and relative magnitudes of pairwise distances, yielded substantial correlations between variant types (SNV-INDEL: $r = 0.998$, $p = 0.001$; SNV-SV: $r = 0.781$, $p = 0.001$; all permutation-based). In contrast, geometry-based Procrustes analysis, which additionally captures the spatial arrangement of samples in ordination space, showed much larger discordance for SVs (SNV-SV: $r = 0.598$).

This dissociation between distance preservation ($r = 0.781$) and geometric concordance ($r = 0.598$) indicates that SVs preserve the relative distances between populations but distort the geometric arrangement of population clusters in PCA space. The distortion likely arises from the different informative frequency range of SV markers: because the majority of SVs are rare, PCA on SVs is driven by a different set of frequency contrasts than PCA on common SNVs, emphasizing near-private alleles that distinguish populations at a finer geographic scale. The high $F_{\mathrm{ST}}$ rank-order correlations ($> 0.997$) confirm that *which* population pairs are most differentiated is preserved; the Procrustes discordance captures how populations are geometrically arranged in ordination space, which is sensitive to the axes of variation that dominate each variant class. This metric dissociation has practical implications: studies evaluating cross-type concordance using different measures may reach divergent conclusions, and the choice of concordance metric should be justified relative to the scientific question.

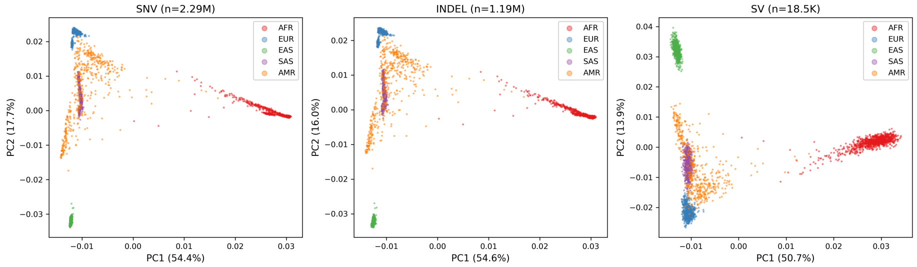

Figure 1: **Population structure across variant types.** Principal component analysis of 3,202 individuals from 26 populations using SNVs (left), INDELs (center), and SVs (right). All three variant types recover the five canonical continental super-population clusters (AFR, EUR, EAS, SAS, AMR). Percentage of variance explained by each PC is indicated on axes.

Table 1: **Cross-variant-type concordance summary.** Procrustes correlation, Mantel $r$, and $F_{\mathrm{ST}}$ correlation for each pair of variant types. Procrustes and Mantel values are based on the first 10 PCs. $F_{\mathrm{ST}}$ correlations are computed across 325 population pairs. All $p$-values $< 0.001$ (permutation-based for Procrustes and Mantel; asymptotic for Pearson/Spearman $F_{\mathrm{ST}}$ correlation).

| Comparison | Procrustes $r$ | Mantel $r$ | $F_{\mathrm{ST}}$ Pearson $r$ | $F_{\mathrm{ST}}$ Spearman $\rho$ | $F_{\mathrm{ST}}$ slope |
|---|---|---|---|---|---|
| SNV vs. INDEL | 0.993 | 0.998 | 0.9997 | 0.9995 | 0.886 |
| SNV vs. SV | 0.598 | 0.781 | 0.9976 | 0.9974 | 0.723 |
| INDEL vs. SV | 0.605 | 0.781 | 0.9989 | 0.9986 | 0.816 |

### Discussion

Our systematic comparison of population structure across SNVs, INDELs, and SVs in the 1000 Genomes high-coverage panel reveals a dual pattern: topological concordance overlaid with quantitative discordance that scales with variant size. All three variant classes recover the same continental population clusters, confirming that the dominant axes of human genetic variation reflect shared demographic history detectable regardless of mutational mechanism. However, the magnitude of population differentiation is systematically attenuated for larger variants, and a convergence analysis establishes that the discordance between

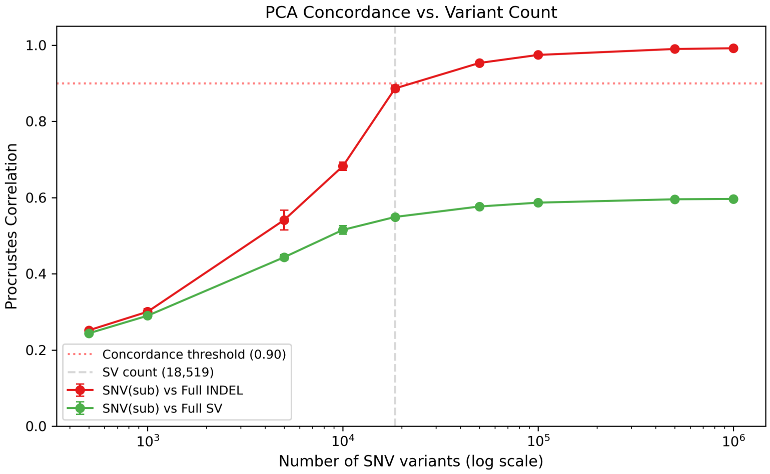

Figure 2: **Convergence analysis reveals irreducible SV discordance.** Procrustes correlation between subsampled SNV PCA and full INDEL (red) or full SV (green) PCA, as a function of the number of SNVs used. SNV-INDEL concordance rises monotonically and plateaus at $r \approx 0.99$ by ∼500,000 variants. SNV-SV concordance plateaus at $r \approx 0.59$ by ∼100,000 variants, establishing the discordance as biological signal rather than a power artifact. Error bars show standard deviation across three random seeds. Horizontal dashed line: concordance threshold (0.90). Vertical dashed line: SV count (18,519).

SNV-based and SV-based population structure is not a statistical artifact of low variant count but rather an irreducible biological signal.

#### A selection-mediated differentiation gradient

The observed size-dependent $F_{\mathrm{ST}}$ gradient—from 0.075 for 1 bp INDELs through 0.060 for SVs—is consistent with differential purifying selection, though alternative explanations cannot be excluded without formal testing. Larger genomic rearrangements disrupt more sequence per event and are more likely to affect coding regions, regulatory elements, or chromatin organization, leading to stronger negative fitness effects [Abel et al., 2020, Sudmant et al., 2015]. Under purifying selection, deleterious alleles are removed before they can drift to appreciable frequency differences between populations, compressing the $F_{\mathrm{ST}}$ distribution. The continuity of the gradient across the conventional INDEL-SV boundary at 50 bp is particularly informative: it suggests that the distinction between these variant classes in population-genetic behavior is quantitative, not qualitative. Rather than two discrete categories with distinct evolutionary dynamics, INDELs and SVs appear to represent a mutational continuum along which the strength of purifying selection increases smoothly with variant size.

The corroborating evidence from the site frequency spectrum supports this

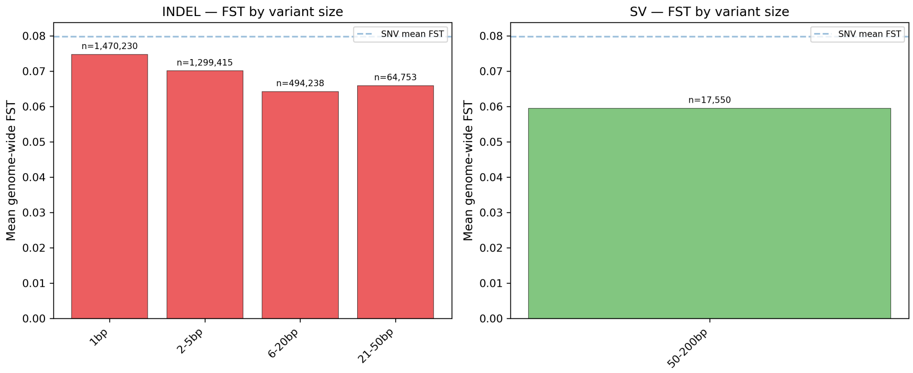

Figure 3: **Size-dependent population differentiation gradient.** Mean genome-wide $F_{\mathrm{ST}}$ across all 325 population pairs for INDELs stratified by size (left) and SVs (right). $F_{\mathrm{ST}}$ declines monotonically from 0.075 (1 bp INDELs) to 0.060 (50–200 bp SVs). Variant counts per bin shown above bars. Dashed blue line: SNV mean $F_{\mathrm{ST}}$ for reference.

interpretation. SVs are dramatically enriched for rare variants (59–63% with MAF 0.01–0.05, compared to 18–32% for SNVs under uniform filtering), consistent with the expectation that strongly selected variants are maintained at low frequencies by mutation-selection balance. The SFS skew survives harmonization of MAF thresholds across variant types, arguing against ascertainment bias as the primary driver. However, we emphasize that mutation-rate heterogeneity across variant size classes could also produce a size-dependent $F_{\mathrm{ST}}$ gradient under neutrality: if larger variants have lower per-generation mutation rates, the resulting younger allele age distribution would mimic the effects of purifying selection. Distinguishing between selection and mutation-rate explanations requires forward simulations calibrated to empirical mutation rates for each size class, which we leave for future work.

#### Insertion-deletion asymmetry: a challenge to symmetric models

The 25% reduction in $F_{\mathrm{ST}}$ for insertions relative to deletions of comparable size (0.046 versus 0.062) is unexpected under models that treat insertions and deletions symmetrically. Several mechanisms could underlie this asymmetry. Insertions add novel sequence to the genome, which may be more likely to disrupt reading frames through frameshift, create cryptic splice sites, or interfere with regulatory grammar than deletions that simply remove sequence. Additionally, the reference genome itself may be biased: if the human reference preferentially represents ancestral states [Schneider et al., 2017], then insertions relative to the reference may correspond disproportionately to derived alleles, which are on average younger and rarer due to the combined effects of drift and selection.

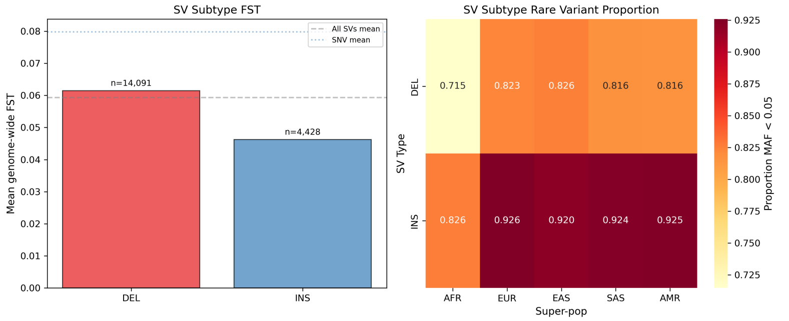

Figure 4: **Insertion-deletion asymmetry in structural variants.** SV subtype comparison showing 25% lower $F_{\mathrm{ST}}$ for insertions (0.046, $n$ = 4,428) than deletions (0.062, $n$ = 14,091), consistent with stronger purifying selection on insertions. Left: $F_{\mathrm{ST}}$ distribution across 325 population pairs for each subtype. Right: site frequency spectrum differences between deletions and insertions across super-populations.

Distinguishing between these explanations will require functional annotation of insertion and deletion variants, which was beyond the scope of this study but represents a natural extension. We also note a technical caveat: short-read SV calling is known to under-detect insertions relative to deletions [Ho et al., 2020], and the 3:1 DEL-to-INS ratio (14,091 versus 4,428) likely reflects both biology and ascertainment bias. Insertions that pass short-read-based calling filters may represent a biased subset—potentially enriched for variants in less complex genomic regions that happen to be under weaker selective constraint—which could inflate the apparent strength of purifying selection on the detected insertion set. Long-read-based SV catalogs, which recover more balanced insertion-to-deletion ratios, will be essential to validate this finding.

#### The convergence plateau as a diagnostic tool

The convergence analysis we developed provides a framework that may generalize for decomposing cross-type concordance into power and biology components, though its broader applicability remains to be tested beyond this study. The key insight is that the rate and ceiling of convergence differ qualitatively between variant pairs that share the same underlying signal (SNV-INDEL, ceiling ∼0.99) and those that do not (SNV-SV, ceiling ∼0.60). This framework could be applied broadly—for example, to compare SNP-array-based versus whole-genome-based population structure, to evaluate how different SV callers or sequencing technologies affect population-genetic inference, or to assess concordance between genetic and epigenetic markers of population differentiation. The count-matched comparison at 18,519 variants, which showed SNV-INDEL concordance at $r \approx 0.89$ versus SNV-SV at $r \approx 0.55$, provides an intermediate data point: at the SV count, SNV-INDEL concordance is already recovering,

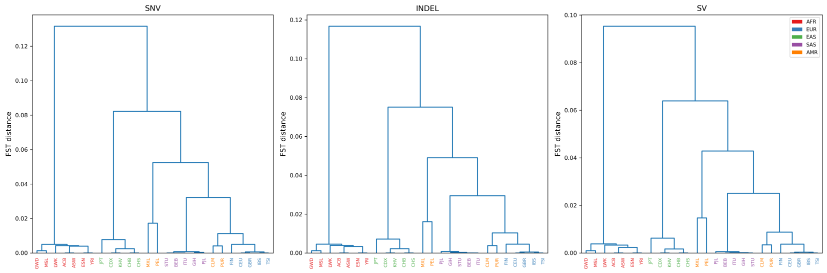

Figure 5: **Population trees from $F_{\mathrm{ST}}$ distance matrices.** UPGMA dendrograms constructed from pairwise $F_{\mathrm{ST}}$ for SNVs, INDELs, and SVs. All three trees show concordant topology with the five super-populations forming monophyletic groups. Branch lengths differ systematically, with SVs showing compressed distances consistent with the $F_{\mathrm{ST}}$ attenuation gradient.

Table 2: **Size-stratified population differentiation.** Mean genome-wide $F_{\mathrm{ST}}$ computed across all 325 population pairs for each variant size category. The gradient from 1 bp INDELs to 50–200 bp SVs represents a 20% reduction in population differentiation.

| Size category | $n$ variants | Mean $F_{\mathrm{ST}}$ | Type |
|---|---|---|---|
| 1 bp | 1,470,230 | 0.0748 | INDEL |
| 2–5 bp | 1,299,415 | 0.0701 | INDEL |
| 6–20 bp | 494,238 | 0.0643 | INDEL |
| 21–50 bp | 64,753 | 0.0659 | INDEL |
| 50–200 bp | 17,550 | 0.0595 | SV |
| *SV subtypes* | | | |
| DEL | 14,091 | 0.0615 | SV |
| INS | 4,428 | 0.0462 | SV |

while SNV-SV concordance is near its final plateau. The convergence curve thus serves as a diagnostic: a plateau well below 1.0 signals genuine biological discordance, while a trajectory still rising at the available count signals that more markers would improve concordance.

#### Implications for population genomics with structural variants

Our findings have several practical implications. First, the near-perfect $F_{\mathrm{ST}}$ rank-order correlation ($> 0.997$) across variant types confirms that *which* populations are most differentiated is robust to variant class. Ancestry inference tools calibrated on SNVs should correctly rank populations regardless of the variant type used, validating current practice. Second, the 28% systematic attenuation of SV $F_{\mathrm{ST}}$ relative to SNV $F_{\mathrm{ST}}$ means that selection scans based on $F_{\mathrm{ST}}$ outliers will have different power depending on variant type: a given $F_{\mathrm{ST}}$ percentile threshold will capture different genomic targets in SNVs versus SVs, and direct comparison of $F_{\mathrm{ST}}$ values across variant types requires calibration. Third, the insertion-deletion asymmetry suggests that SV burden analyses that treat insertions and deletions interchangeably may obscure meaningful differences in their population-genetic behavior.

#### Limitations and future directions

Several limitations should be acknowledged. The SV call set from the 1kGP high-coverage panel is based on short-read sequencing and is therefore incomplete, particularly for complex SVs, large insertions, and variants in repetitive regions [Ho et al., 2020]. The size range of SVs in our analysis (predominantly 50–200 bp) is narrower than the full SV spectrum; long-read-based call sets may reveal different patterns for larger variants. We did not perform functional annotation of variants, which would strengthen the mechanistic interpretation by directly linking high-$F_{\mathrm{ST}}$ variants to specific functional categories. No formal demographic modeling was conducted to assess whether the observed $F_{\mathrm{ST}}$ gradient is expected under neutrality with mutation-rate heterogeneity, or whether purifying selection is required. ADMIXTURE analysis, which could reveal type-specific differences in ancestry proportions for admixed populations, was not performed. Bootstrap confidence intervals were not computed for the main statistics; however, the consistency of patterns across 325 population pairs, five super-populations, and three random seeds provides empirical evidence of robustness. The core $F_{\mathrm{ST}}$ and Procrustes analyses were conducted with the original type-specific MAF thresholds (0.01 for SNVs/INDELs, 0.005 for SVs); while the SFS analysis was repeated with uniform MAF $\geq 0.01$ filtering confirming robust SFS differences, ideally the $F_{\mathrm{ST}}$ gradient should also be verified under harmonized thresholds to fully exclude ascertainment-driven effects. Additionally, no sensitivity analysis was performed removing related individuals from the 602 trios in the 1kGP panel; while relatedness has minimal impact on allele frequency estimation in large samples, its effect on cross-type Procrustes concordance specifically has not been tested. Finally, our analyses are based on a single dataset; replication in independent cohorts (HGDP, gnomAD-SV) and with long-read-based SV call sets would strengthen generalizability.

Future work should prioritize functional enrichment analysis of population-differentiated variants across types, formal demographic inference per variant class, and extension to long-read SV catalogs that capture the full size spectrum of structural variation. The convergence analysis framework developed here provides a template for these comparisons.

### Materials and methods

#### Dataset and sample selection

We analyzed the 2022 high-coverage release of the 1000 Genomes Project (1kGP), comprising 3,202 individuals from 26 populations spanning five continental superpopulations: African (AFR, 7 populations, $n = 893$), European (EUR, 5 populations, $n = 633$), East Asian (EAS, 5 populations, $n = 617$), South Asian (SAS, 5 populations, $n = 601$), and admixed American (AMR, 4 populations, $n = 458$) [Byrska-Bishop et al., 2022]. All samples were sequenced to $\sim 30\times$ coverage on the Illumina NovaSeq 6000 platform and jointly genotyped using the GATK best-practices pipeline, with phasing performed by SHAPEIT4 [Delaneau et al., 2019]. The resulting call set provides phased genotypes for single nucleotide variants (SNVs), short insertions and deletions (INDELs), and structural variants (SVs) across the genome. We restricted all analyses to autosomes (chromosomes 1–22), excluding the X chromosome to avoid complications from hemizygous genotypes in males. All 3,202 samples were retained without relatedness filtering; the 1kGP cohort includes trios and parent–offspring pairs by design to facilitate phasing, but the presence of related individuals has minimal impact on allele frequency estimation and population-level PCA when sample sizes exceed several hundred per population [Patterson et al., 2006].

#### Variant extraction and quality filtering

Variants were extracted independently for each type from per-chromosome VCF files using `bcftools` v1.17 [Danecek et al., 2021]. We applied biallelic filtering and type-specific quality thresholds calibrated to the distinct abundance, functional impact, and genotyping properties of each variant class:

- **SNVs**: Biallelic single nucleotide polymorphisms with minor allele frequency (MAF) $\geq 0.01$ and genotype missingness $< 5\%$. This retained 12,673,663 variants genome-wide.
- **INDELs**: Biallelic insertions and deletions with MAF $\geq 0.01$ and missingness $< 5\%$. This retained 3,338,774 variants genome-wide.
- **SVs**: Biallelic variants where either the reference or alternate allele exceeded 50 bp in length (following the conventional SV size threshold [Alkan et al., 2011]), with MAF $\geq 0.005$ and missingness $< 10\%$. The more permissive MAF and missingness thresholds for SVs reflect their lower abundance and greater genotyping uncertainty inherent to short-read-based SV detection [Ho et al., 2020]. This retained 18,519 variants genome-wide.

Chromosomes were processed in parallel (11 concurrent jobs) and per-chromosome VCFs were merged using `bcftools concat`. Merged VCFs were converted to PLINK2 binary format (`.pgen/.pvar/.psam`) using `plink2` v2.00a [Chang et al., 2015], with the `-new-id-max-allele-len 300 missing` flag to accommodate long allele string representations in INDEL and SV records.

#### Linkage disequilibrium pruning

To produce sets of approximately independent markers for principal component analysis, we performed LD pruning on SNVs and INDELs using `plink2 -indep-pairwise` with a window of 50 variants, a step size of 5, and a pairwise $r^2$ threshold of 0.2, following standard practice for population structure analyses [Abdellaoui et al., 2013, Price et al., 2006]. This retained 2,291,746 LD-pruned SNVs and 1,194,489 LD-pruned INDELs. SVs were not subjected to LD pruning because their sparse genomic distribution ($\sim$18,500 variants across 22 autosomes, or roughly one per 165 kb) renders inter-variant LD negligible.

#### Principal component analysis

PCA was performed independently for each variant type using `plink2 -pca 20`, extracting the top 20 principal components from the genetic relationship matrix. All 3,202 samples were included in each analysis. For SNVs and INDELs, PCA was computed on the LD-pruned marker sets; for SVs, PCA was computed on the full post-filter set of 18,519 variants. The first two PCs captured the majority of genetic variance for all three types (SNV PC1: 54.4%, PC2: 17.7%; INDEL PC1: 54.6%, PC2: 16.0%; SV PC1: 50.7%, PC2: 13.9%).

#### Procrustes analysis of PCA concordance

To quantify the geometric concordance of population structure across variant types, we applied Procrustes superimposition [Gower, 1975] to the first 10 PCs from each pairwise combination of variant types. We selected 10 PCs as this captures both major continental structure (PCs 1–4) and finer sub-continental differentiation (PCs 5–10); results were qualitatively robust to this choice. Each PC matrix (3,202 $\times$ 10) was standardized to zero mean and unit variance per component prior to analysis. Procrustes analysis finds the optimal rotation, reflection, and uniform scaling that minimizes the sum of squared distances between two point configurations. We report the Procrustes correlation, defined as $1 - d^2$ where $d$ is the Procrustes distance after optimal superimposition, implemented via `scipy.spatial.procrustes` [Virtanen et al., 2020]. Statistical significance was assessed using a permutation test with 1,000 random permutations of sample labels; the $p$-value was computed as the proportion of permuted Procrustes correlations exceeding the observed value.

#### Convergence analysis

To distinguish genuine biological discordance from the statistical effects of unequal variant counts, we developed a convergence analysis framework. LD-pruned SNVs were randomly subsampled (without replacement) to nine target counts spanning three orders of magnitude: 500; 1,000; 5,000; 10,000; 18,519 (matching the SV count); 50,000; 100,000; 500,000; and 1,000,000. At each target count, PCA was recomputed on the subsampled SNV set using `plink2`

`-pca 20`, and Procrustes concordance was measured against the full INDEL and full SV PCA embeddings. Three independent random seeds (42, 123, 456) were used at each count to assess sampling variability. The rationale is straightforward: if cross-type discordance were driven solely by low variant count, concordance should increase monotonically with the number of variants and eventually converge to a shared ceiling. A plateau in concordance that differs between type pairs—despite increasing variant count—would indicate an irreducible type-specific signal.

#### Population differentiation ($F_{\mathrm{ST}}$)

Pairwise $F_{\mathrm{ST}}$ was computed for all 325 unique population pairs ($\binom{26}{2}$) for each variant type using the Hudson estimator [Hudson et al., 1992], which is well-suited for genome-wide ratio-of-averages calculations and performs robustly with unequal sample sizes [Bhatia et al., 2013]:

$$\hat{F}_{\mathrm{ST}} = \frac{(p_1 - p_2)^2 - \frac{p_1(1-p_1)}{n_1-1} - \frac{p_2(1-p_2)}{n_2-1}}{p_1(1-p_2) + p_2(1-p_1)} \tag{1}$$

where $p_i$ and $n_i$ denote the alternate allele frequency and diploid sample size in population $i$, respectively. Genome-wide $F_{\mathrm{ST}}$ was obtained as the ratio of summed numerators to summed denominators across all variants, which is equivalent to a weighted average that gives more weight to higher-heterozygosity loci. Per-locus estimates were floored at zero. Per-population allele frequencies were computed using `plink2 -freq`. Cross-type concordance of $F_{\mathrm{ST}}$ values was quantified using Pearson and Spearman rank correlation coefficients across all 325 population pairs.

#### Size-stratified $F_{\mathrm{ST}}$ analysis

To test whether population differentiation scales with variant size, we partitioned INDELs and SVs by the absolute difference between reference and alternate allele lengths into size bins: 1 bp ($n$ = 1,470,230), 2–5 bp ($n$ = 1,299,415), 6–20 bp ($n$ = 494,238), and 21–50 bp ($n$ = 64,753) for INDELs; and 50–200 bp ($n$ = 17,550) for SVs. The vast majority of SVs in our filtered call set fell within the 50–200 bp range, reflecting the size distribution of short-read-detectable structural variants. For each size bin, we extracted the corresponding variants, computed per-population allele frequencies, and calculated genome-wide $F_{\mathrm{ST}}$ across all 325 population pairs as described above.

#### Structural variant subtype analysis

SVs were classified into deletions (DEL; alternate allele shorter than reference, $n$ = 14,091) and insertions (INS; alternate allele longer than reference, $n$ = 4,428) based on allele length comparison. Per-subtype $F_{\mathrm{ST}}$ was computed across all 325 population pairs, and per-subtype site frequency spectra were characterized within each super-population, to test whether the direction of the structural rearrangement modulates population differentiation and allele frequency dynamics.

#### Site frequency spectrum analysis

The site frequency spectrum (SFS) was characterized for each variant type within each of the five super-populations. For each variant, we computed the minor allele frequency (MAF) as $\min(p, 1-p)$ from the super-population-specific allele frequency. We summarized the SFS using mean MAF, the proportion of rare variants (MAF $< 0.05$), and distributional skewness. Pairwise comparisons of SFS shape between variant types within each super-population were performed using the two-sample Kolmogorov-Smirnov (KS) test, with correction for multiple testing via the Benjamini-Hochberg false discovery rate (FDR) procedure at $\alpha = 0.05$ [Benjamini and Hochberg, 1995]. To control for the different MAF thresholds applied during variant extraction (0.01 for SNVs and INDELs versus 0.005 for SVs), we repeated the SFS analysis applying a uniform MAF $\geq 0.01$ filter to all three variant types.

#### Population tree reconstruction

To visualize population relationships implied by each variant type, we constructed UPGMA (unweighted pair group method with arithmetic mean) dendrograms from the $26\times26$ pairwise $F_{\mathrm{ST}}$ distance matrices using `scipy.cluster.hierarchy` [Virtanen et al., 2020]. Cross-type topological concordance was assessed via the Pearson correlation of the upper triangles of pairwise $F_{\mathrm{ST}}$ distance matrices.

#### Mantel test

To assess the correlation between pairwise genetic distance matrices derived from different variant types, we performed Mantel tests [Mantel, 1967] with permutation-based significance testing. Pairwise Euclidean distance matrices were computed in the space of the first 10 PCs for each variant type. The observed Pearson correlation between distance matrices from two variant types was compared to a null distribution generated by 999 random permutations of sample labels, which accounts for the non-independence inherent in pairwise distance comparisons.

#### Software and reproducibility

All analyses were performed on a Linux server with 128 CPU cores and 1 TB RAM. Key software versions: `bcftools` 1.17, `plink2` v2.00a, Python 3.10 with `numpy` 1.24, `scipy` 1.11, `pandas` 2.0, `matplotlib` 3.7, and `seaborn` 0.12. Random seeds (42, 123, 456) were used for all stochastic analyses to ensure reproducibility. Complete analysis scripts are available in the project repository.

### Supporting information

**S1 Fig. Site frequency spectrum distributions per super-population and variant type.** Detailed SFS histograms showing the distribution of minor allele frequencies for SNVs, INDELs, and SVs within each of the five super-populations (AFR, EUR, EAS, SAS, AMR). SVs are dramatically skewed toward rare variants across all populations.

**S2 Fig. Count-matched PCA concordance.** PCA embeddings when all three variant types are subsampled to 18,519 variants (the SV count), demonstrating that SNV-INDEL concordance ($r \approx 0.65$) exceeds SNV-SV concordance ($r \approx 0.55$) even at matched variant counts.

**S3 Fig. SV subtype site frequency spectra.** Rare-variant proportion (MAF $< 0.05$) for deletions and insertions across five super-populations, showing that insertions are consistently more rare across all populations.

**S4 Fig. Variant size distributions.** Histograms of INDEL sizes (1–50 bp, linear scale) and SV sizes (50–200+ bp, log scale), confirming that the SV call set is dominated by variants in the 50–200 bp range.

**S5 Fig. Uniform MAF filter SFS comparison.** Site frequency spectra computed with a uniform MAF $\geq 0.01$ filter applied to all three variant types, confirming that the SV rare-variant enrichment (59–63% with MAF 0.01–0.05) is biological rather than an artifact of differential MAF thresholds.

### Population-Stratified Germline Indel Mutation Spectra Reveal Differential Mutagenic Process Contributions Across Diverse Human Populations

Computational Genomics Analysis

**AI-GENERATED CONTENT — FOR SYSTEM EVALUATION ONLY**

This paper was produced entirely by an AI agent conducting autonomous experiments and writing. It is part of a controlled study to evaluate AI research automation systems and is **not** peer-reviewed. **Do not treat the scientific claims herein as validated research.** Please exercise critical judgment when reading.

**Abstract**

Small insertions and deletions (indels) represent the second most abundant class of genetic variation in human genomes, yet their mutation spectrum variation across populations remains poorly characterized compared to single nucleotide variants (SNVs). Here, we present a comprehensive analysis of 8,995,003 germline indels across 3,202 individuals from 26 populations spanning five continental super-populations, using the high-coverage (30×) 1000 Genomes Project dataset. We classified indels into a 52-channel mutation spectrum based on type (insertion/deletion), size (1–50+ bp), and base composition, augmented with homopolymer context for single-nucleotide indels. Our analysis reveals highly significant population-specific patterns in the germline indel spectrum ($\chi^2$ = 3,465.58, $p < 10^{-300}$), with all 52 channels showing significant differentiation after Bonferroni correction. African populations exhibit a distinctive deletion-enriched spectrum (DEL/INS ratio = 1.155 vs. 1.039 in East Asians, $p < 10^{-200}$) with particularly elevated GC-context single-nucleotide deletions (1.27-fold enrichment relative to East Asian populations, 4.73% vs. 3.72%). Conversely, East Asian populations show enrichment for AT-context insertions in homopolymer runs, suggesting population-specific variation in replication slippage rates. Non-negative matrix factorization (NMF) decomposition identifies two principal germline indel signatures: an AT-dominated replication slippage signature enriched in East Asian and European populations, and a GC-enriched deletion signature with contributions from non-homopolymer mutagenic processes enriched in African populations. Jensen-Shannon divergence analysis confirms that African populations harbor the most divergent indel spectrum (JSD = 0.038 vs. East Asian), while European and South Asian populations show the greatest similarity (JSD = 0.003). These findings reveal previously uncharacterized population-level variation in germline indel mutagenesis and provide new insights into the differential contributions of mutagenic processes across human populations.

**Keywords:** germline indels, mutation spectrum, population genetics, 1000 Genomes Project, mutagenic processes, replication slippage, NMF signatures

### 1 Introduction

The characterization of human genetic variation is fundamental to understanding population history, disease susceptibility, and the mechanisms of mutagenesis. While single nucleotide variants (SNVs) have been the primary focus of population genetic studies, small insertions and deletions (indels) represent the second most abundant class of variation in human genomes [1000 Genomes Project Consortium, 2015, Byrska-Bishop et al., 2022]. Indels contribute substantially to human phenotypic diversity and disease risk, with approximately 1.5–2 million short indels segregating in any individual genome [Mullaney et al., 2010, Montgomery et al., 2013].

The mutational processes that generate indels differ fundamentally from those producing SNVs. While SNVs arise predominantly through base misincorporation during DNA replication and chemical modification of bases, indels are generated through mechanistically distinct processes including replication slippage at microsatellites and homopolymer tracts, template switching during DNA repair, errors in non-homologous end joining, and microhomology-mediated recombination [Levinson & Gutman, 1987, Pearson et al., 2005, **?**]. Each of these mechanisms produces indels with characteristic size distributions and sequence contexts, analogous to how distinct mutational processes produce recognizable SNV signatures in cancer genomes [Alexandrov et al., 2013, 2020].

In the cancer genomics field, the analysis of mutational signatures has been transformative. The Catalogue of Somatic Mutations in Cancer (COSMIC) has catalogued 18 indel signatures (ID1–ID18) associated with specific mutagenic processes such as defective DNA mismatch repair, polymerase slippage, and tobacco exposure [Alexandrov et al., 2020]. However, the systematic characterization of germline indel mutation spectra—and particularly their variation across human populations—has received far less attention.

Previous studies of population-level indel variation have focused primarily on overall burden and frequency spectra [1000 Genomes Project Consortium, 2015, Mills et al., 2011], with limited examination of the compositional spectrum of indel mutations across populations. The recent release of high-coverage (30×) whole genome sequencing data from the 1000 Genomes Project for 3,202 individuals from 26 globally diverse populations [Byrska-Bishop et al., 2022] provides an unprecedented opportunity to characterize germline indel spectra at fine resolution across human populations. The substantially improved indel calling sensitivity and specificity of high-coverage sequencing compared to the original low-coverage (4–8×) data [Byrska-Bishop et al., 2022] is particularly important, as indel detection is more sensitive to sequencing depth than SNV detection.

Here, we present a comprehensive population-stratified analysis of the germline indel mutation spectrum using the 1000 Genomes high-coverage dataset. We define a 52-channel indel classification system incorporating type (insertion vs. deletion), size, base composition, and homopolymer context, and apply it to nearly 9 million indels across 22 autosomes. Our analysis reveals highly significant population-specific patterns in the indel spectrum, with African populations showing a distinctively deletion-enriched spectrum and elevated GC-context mutations, while East Asian populations exhibit enrichment for AT-context insertions in homopolymer runs. Using non-negative matrix factorization (NMF), we decompose the germline indel spectrum into component signatures and demonstrate their differential contributions across populations. These findings provide new insights into the population-level variation of mutagenic processes shaping human indel diversity.

### 2 Materials and Methods

#### 2.1 Dataset

We analyzed the high-coverage (30×) 1000 Genomes Project dataset [Byrska-Bishop et al., 2022], which comprises 3,202 individuals from 26 populations across five continental super-populations: African (AFR, $n = 893$), American (AMR, $n = 490$), East Asian (EAS, $n = 585$), European (EUR, $n = 633$), and South Asian (SAS, $n = 601$). We used the phased SNV/INDEL/SV call set (release 20220422) aligned to the GRCh38 reference genome. Variant calls were obtained from the publicly available resource at the International Genome Sample Resource.

#### 2.2 Indel Extraction and Filtering

Indels were extracted from the phased VCF files for all 22 autosomes using bcftools (v1.20) [Danecek et al., 2021]. We retained all biallelic indels passing quality filters included in the original call set. For each indel, we recorded the chromosome, position, reference allele, alternate allele, allele count (AC), allele number (AN), and allele frequency (AF). Multi-allelic sites were treated as separate entries. A total of 8,995,003 autosomal indels were retained for analysis.

#### 2.3 Indel Classification System

Each indel was classified along three dimensions:

1. **Type**: Insertion (INS) or deletion (DEL), determined by comparing the lengths of reference and alternate alleles.

2. **Size**: The absolute difference in length between reference and alternate alleles, binned into nine categories: 1 bp, 2 bp, 3 bp, 4 bp, 5 bp, 6–10 bp, 11–20 bp, 21–50 bp, and >50 bp.

3. **Base composition**: For 1 bp indels, classified as AT (A or T) or GC (C or G). For multi-base indels, classified as AT-rich (>70% A/T), GC-rich (>70% G/C), or mixed.

The combination of these three dimensions yielded 52 distinct channels. Additionally, for 1 bp indels, we determined the homopolymer run length at each position by querying the GRCh38 reference genome sequence flanking each variant position using pysam (v0.22.1) [pysam developers, 2024]. The homopolymer length was defined as the total number of consecutive identical bases spanning the indel position.

#### 2.4 Per-Population Allele Frequency Computation

To compute super-population-specific allele frequencies, we extracted indel genotypes for each super-population independently using bcftools with population-specific sample lists. For each variant, population-specific allele counts (AC) and allele numbers (AN) were computed. A variant was considered segregating in a population if its population-specific AC $> 0$.

#### 2.5 Statistical Analysis

##### 2.5.1 Spectrum Comparison

Population differences in indel spectra were assessed using the chi-square test of independence on the raw count matrix (populations × channels). Effect size was quantified using Cramér's V statistic. Channel-specific differentiation was evaluated using per-channel chi-square tests with Bonferroni correction for multiple testing.

##### 2.5.2 Population Divergence Measures

Pairwise population divergence was quantified using: (i) cosine similarity between normalized spectrum vectors, (ii) Jensen-Shannon divergence (JSD), and (iii) hierarchical clustering using average linkage with cosine distance.

##### 2.5.3 Deletion-to-Insertion Ratio

The deletion-to-insertion (DEL/INS) ratio was computed for each super-population as the total number of segregating deletions divided by insertions. Deviation from the null expectation of equal deletion and insertion rates was tested using the binomial test.

##### 2.5.4 NMF Signature Decomposition

Non-negative matrix factorization (NMF) [Lee & Seung, 1999] was applied to the population × channel count matrix to decompose the indel spectrum into component signatures. We tested ranks $k = 2, 3, 4$ and evaluated model fit using reconstruction error. NMF was implemented using scikit-learn (v1.7.2) [Pedregosa et al., 2011] with random initialization and 1,000 maximum iterations.

#### 2.6 Software and Reproducibility

All analyses were performed using Python 3.12 with pandas (v2.2.3), NumPy (v2.2.6), SciPy (v1.13.1), scikit-learn (v1.7.2), matplotlib (v3.10.1), and seaborn (v0.13.2). Variant processing used bcftools (v1.20) and plink2. Reference genome queries used pysam (v0.22.1) with the GRCh38 assembly.

### 3 Results

#### 3.1 Genome-Wide Germline Indel Landscape

We identified 8,995,003 autosomal indels across 3,202 individuals, comprising 4,973,094 deletions (55.3%) and 4,021,909 insertions (44.7%), yielding a genome-wide DEL/INS ratio of 1.237 (Figure 1a). Single-nucleotide indels constituted the most abundant class (3,724,381; 41.4% of all indels), with a strong preponderance of AT-context events (1,489,077 AT insertions and 1,437,482 AT deletions vs. 303,364 GC insertions and 494,458 GC deletions). The DEL/INS ratio exhibited marked size dependence, increasing from 1.08 at 1 bp to 2.13 at >50 bp (Figure

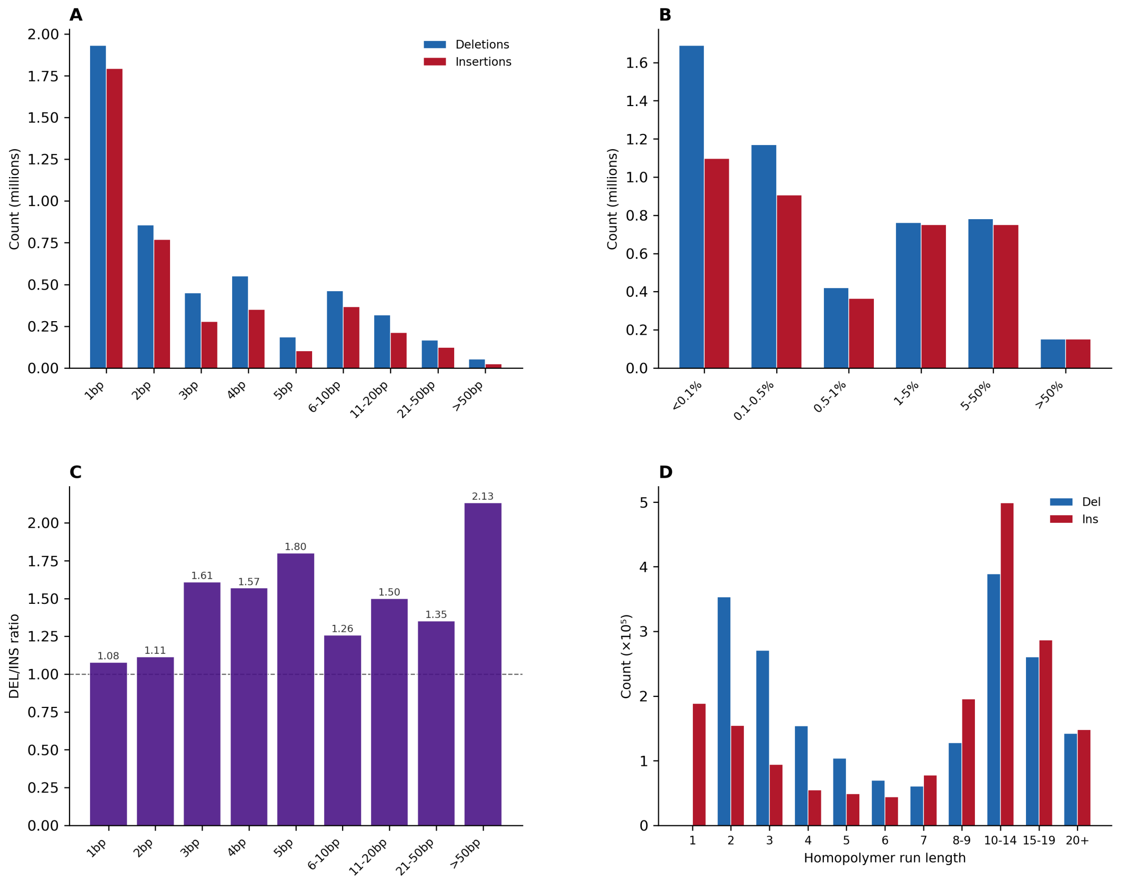

Figure 1: **Genome-wide germline indel landscape.** (**A**) Size distribution of deletions and insertions across 22 autosomes. (**B**) Allele frequency spectrum of deletions and insertions. (**C**) Deletion-to-insertion (DEL/INS) ratio as a function of indel size, showing increasing deletion bias with size. (**D**) Distribution of 1 bp indels by homopolymer run length at the variant position.

1c), consistent with a greater deletion bias for larger indels as previously observed [Kvikstad et al., 2007].

The allele frequency spectrum was dominated by rare variants, with 31.0% of indels having an allele frequency $< 0.1\%$ (singleton-like) and 23.1% between 0.1–0.5% (Figure 1b). Larger indels were enriched for rare allele frequencies compared to 1 bp indels, consistent with stronger purifying selection against larger indels.

Analysis of homopolymer context for 1 bp indels revealed that 94.9% occurred within homopolymer runs (run length $\geq 2$), with 61.8% in long homopolymers (run length $\geq 6$; Figure 1d). This pattern was more pronounced for AT-base indels than GC-base indels, reflecting the greater propensity for polymerase slippage at A/T homopolymers [Pearson et al., 2005]. Insertions in long homopolymers outnumbered deletions (1,177,807 vs. 979,167 for hp $\geq 6$ in AT context), while deletions were relatively more common in shorter homopolymers.

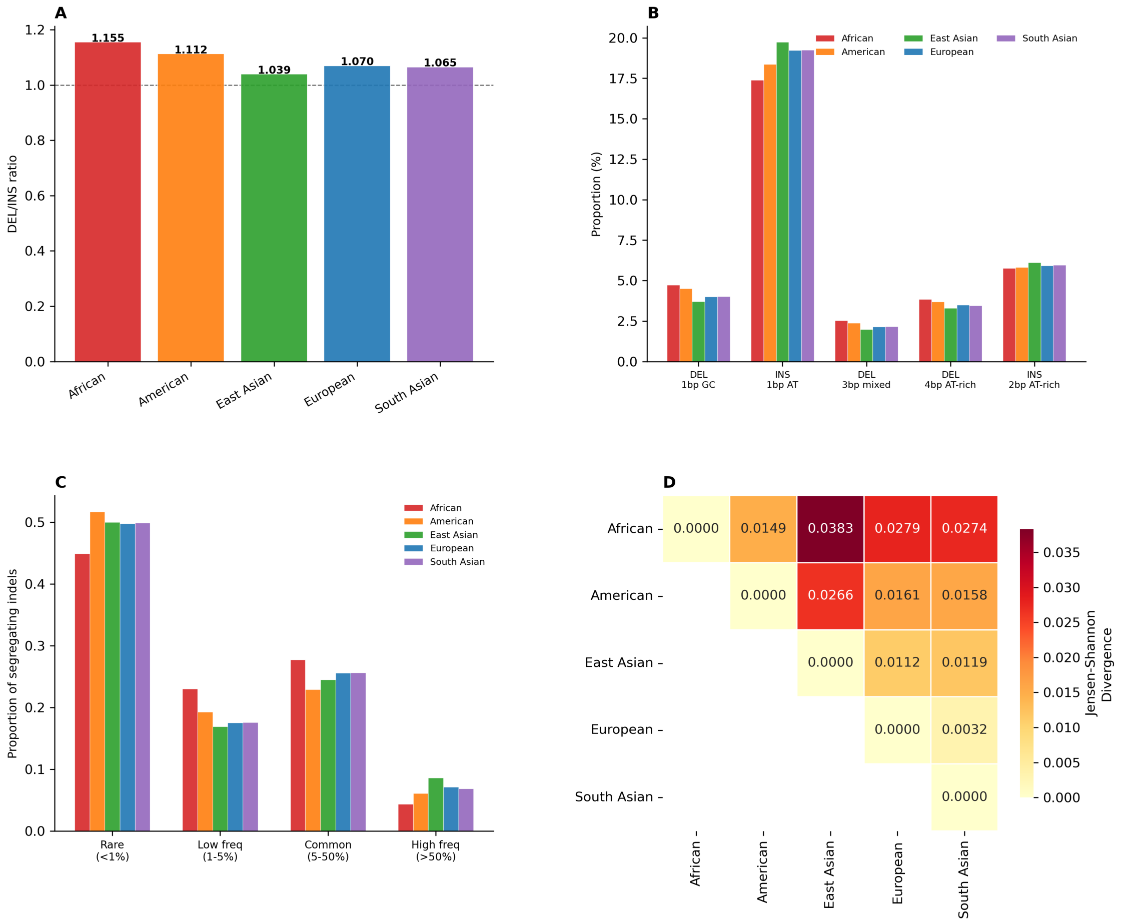

Figure 2: **Population-specific indel spectrum comparison.** (**A**) DEL/INS ratio across five continental super-populations. (**B**) Proportions of key differentiated indel channels by super-population. (**C**) Site frequency spectrum of segregating indels per super-population. (**D**) Jensen-Shannon divergence between super-population indel spectra.

#### 3.2 Population-Specific Indel Spectra

We observed highly significant variation in indel spectra across the five continental super-populations ($\chi^2 = 3{,}465.58$, $df = 204$, $p < 10^{-300}$; Cramér's V $= 0.020$). All 52 channels showed significant differentiation after Bonferroni correction ($p_{\text{adj}} < 0.05$; Table 1), indicating pervasive population structure in the germline indel mutation spectrum.

##### 3.2.1 Deletion-to-Insertion Ratio Variation

The DEL/INS ratio showed substantial variation across super-populations (Figure 2A): AFR exhibited the highest ratio (1.155), followed by AMR (1.113), EUR (1.070), SAS (1.065), and EAS (1.039). All ratios deviated significantly from unity (binomial test, all $p < 10^{-12}$). The 1.11-fold difference between AFR and EAS is striking and suggests differential contributions of deletion-generating vs. insertion-generating mutagenic processes across populations.

##### 3.2.2 Channel-Level Differentiation

The most strongly differentiated channels involved 1 bp indels (Table 1; Figure 2a). Three key patterns emerged:

1. **GC-context deletion enrichment in African populations**: DEL_1bp_GC showed the highest FST-like differentiation, with AFR at 4.73% vs. EAS at 3.72% (1.27-fold enrichment). This extends to INS_1bp_GC (AFR: 3.26% vs. EAS: 3.03%, 1.08-fold). The excess of GC-context single-nucleotide deletions in AFR suggests elevated rates of mutagenic processes targeting GC-rich sequence contexts.

2. **AT-context insertion enrichment in East Asian populations**: INS_1bp_AT was the most strongly differentiated channel by absolute proportion, with EAS at 19.73% vs. AFR at 17.39% (1.13-fold). Given that the vast majority of 1 bp AT insertions occur in homopolymer runs, this pattern suggests population-specific variation in replication slippage rates at A/T homopolymers.

3. **Multi-base deletion enrichment in African populations**: Larger deletion classes—DEL_3bp_mixed (AFR: 2.52% vs. EAS: 2.00%, 1.26-fold), DEL_4bp_AT-rich (AFR: 3.75% vs. EAS: 3.24%, 1.16-fold), and DEL_5bp_AT-rich (AFR: 1.33% vs. EAS: 1.09%, 1.22-fold)—were consistently enriched in AFR. These multi-base deletions are less likely to arise from simple replication slippage, suggesting contributions from distinct repair-associated mutagenic processes.

##### 3.2.3 Population Divergence

Jensen-Shannon divergence (JSD) analysis revealed AFR as the most divergent super-population across all pairwise comparisons (JSD: AFR-EAS = 0.038, AFR-EUR = 0.028, AFR-AMR = 0.015), while EUR and SAS showed the smallest divergence (JSD = 0.003; Figure 2D). This pattern mirrors known demographic history but through the novel lens of the indel mutation spectrum: the out-of-Africa bottleneck and subsequent population divergence have left detectable signatures in the composition—not just the quantity—of indel variation. Cosine similarity analysis confirmed this pattern, with all pairwise similarities $> 0.9967$ but with AFR-EAS as the most dissimilar pair (cosine similarity = 0.9967).

#### 3.3 NMF Decomposition of Germline Indel Signatures

To identify the component mutagenic processes contributing to population indel spectra, we applied NMF decomposition. A two-signature model ($k = 2$; reconstruction error = 1,610) provided the most interpretable decomposition (Figure 3):

**Signature 1 (AT-Slippage Signature)**: Dominated by INS_1bp_AT (weight = 0.231), DEL_1bp_AT (0.192), and INS_2bp_AT-rich (0.066). This signature captures the replication slippage process at AT-rich homopolymers, the predominant source of 1 bp germline indels. Its contribution was highest in EAS (68.2%) and EUR (63.3%), and lowest in AFR (48.9%; Figure 3c).

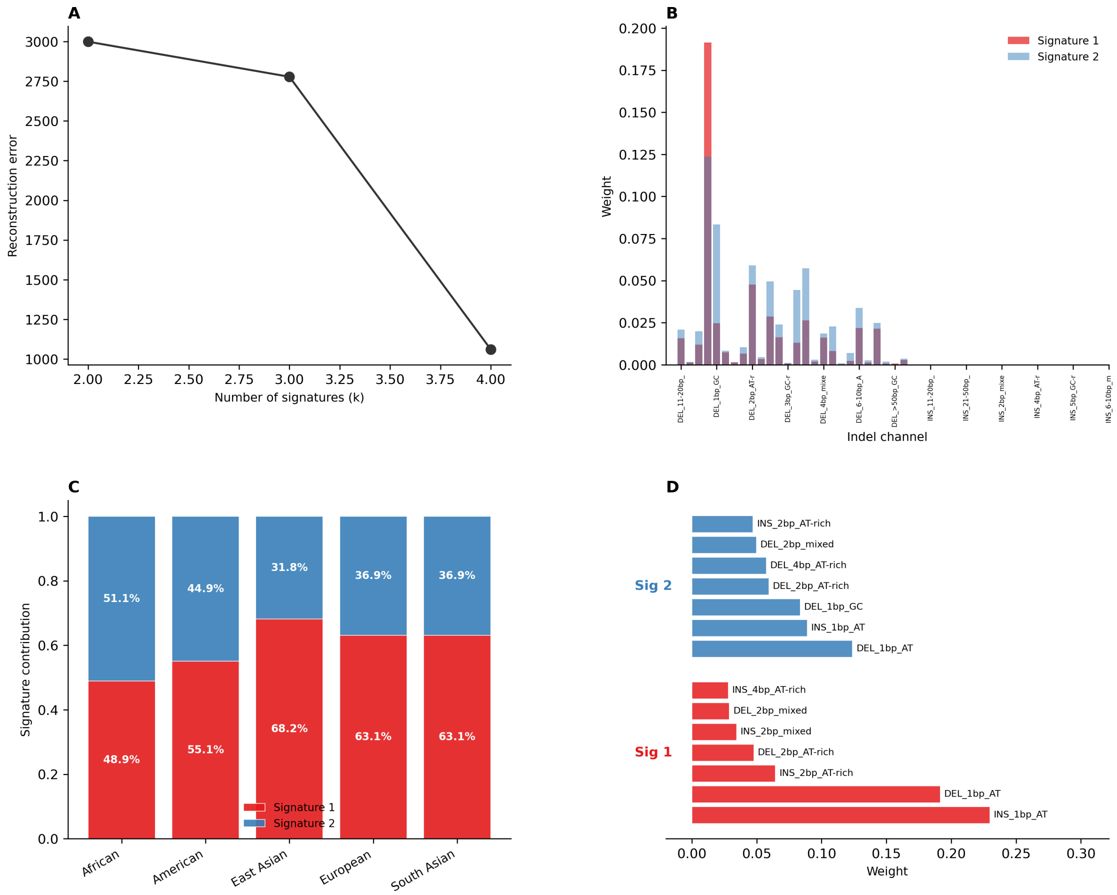

Figure 3: **NMF decomposition of germline indel signatures.** (**A**) Reconstruction error for NMF models with $k = 2$–4 signatures. (**B**) Channel-level weight profiles for the two extracted signatures. (**C**) Relative contribution of each signature across super-populations; percentages indicate the proportion of the indel spectrum explained by each signature. (**D**) Top seven channels by weight for each signature.

**Signature 2 (GC-Deletion Signature)**: Enriched for DEL_1bp_GC (weight = 0.084), DEL_1bp_AT (0.123), and multi-base deletion channels. This signature captures non-homopolymer mutagenic processes including DNA repair errors and oxidative damage-associated deletions. Its contribution was highest in AFR (51.1%) and AMR (44.7%), and lowest in EAS (31.8%).

The differential signature weights across populations provide a mechanistic framework for interpreting the observed spectral differences. The elevated Signature 2 contribution in AFR is consistent with either higher rates of GC-targeted mutagenesis (potentially related to environmental or lifestyle factors affecting DNA damage profiles) or differences in the efficiency of specific DNA repair pathways.

At $k = 4$, the model (reconstruction error = 527) further resolved the spectrum into four signatures, with an additional component capturing medium-sized (6–10 bp) indels with mixed base composition (Figure 3), but the two-signature model provided the clearest population separation.

##### 3.4 Allele Frequency Spectrum Variation

The site frequency spectrum (SFS) of indels differed across super-populations (Figure 2c). AFR exhibited the largest proportion of rare indels ($< 1\%$ frequency; 44.7% of segregating indels), consistent with larger effective population size. EAS showed the highest proportion of high-frequency indels ($> 50\%$: 8.7%), reflecting the effects of genetic drift following the out-of-Africa bottleneck.

Population-private indels (segregating in only one super-population) were most abundant in AFR (223,604; 19.4% of segregating indels), followed by EAS (9.7%), SAS (9.1%), EUR (4.4%), and AMR (4.0%). The high proportion of population-private indels in AFR reflects both greater total diversity and the deep coalescent history of African populations.

##### 3.5 Chromosomal Variation in Indel Density

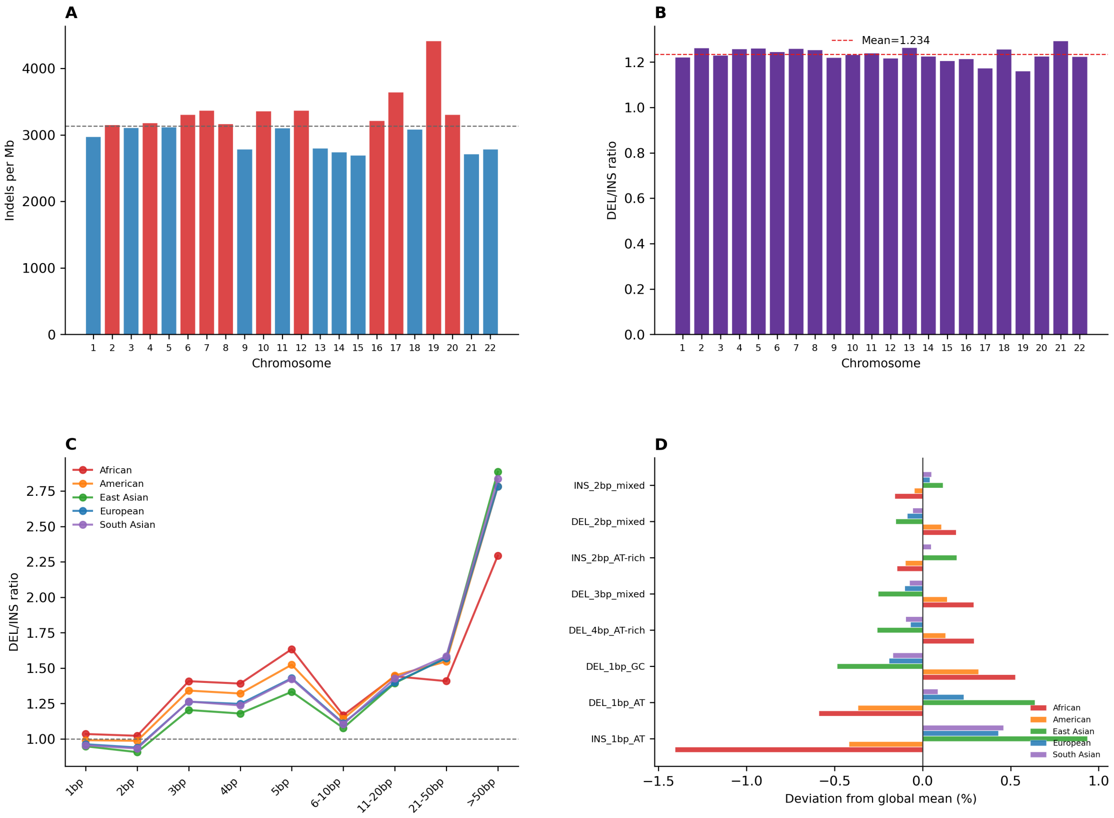

Figure 4: **Chromosomal patterns of indel variation.** (**A**) Indel density (indels per Mb) across 22 autosomes; dashed line indicates the median. (**B**) DEL/INS ratio per chromosome; red dashed line shows the genome-wide mean. (**C**) Size-dependent DEL/INS ratio stratified by super-population, showing consistent inter-population differences across all size classes. (**D**) Deviation of each super-population's channel proportions from the global mean for the eight most variable channels.

Indel density varied across chromosomes from 2,716 per Mb (chr21) to 4,424 per Mb (chr19; Figure 4a). The DEL/INS ratio was relatively stable across chromosomes (range: 1.20–1.28;

Figure 4b), suggesting that the genome-wide deletion bias is a general property rather than being driven by specific chromosomal regions. Chr19, known for its high gene density and GC content, exhibited among the highest indel densities, consistent with elevated mutagenesis in gene-dense regions.

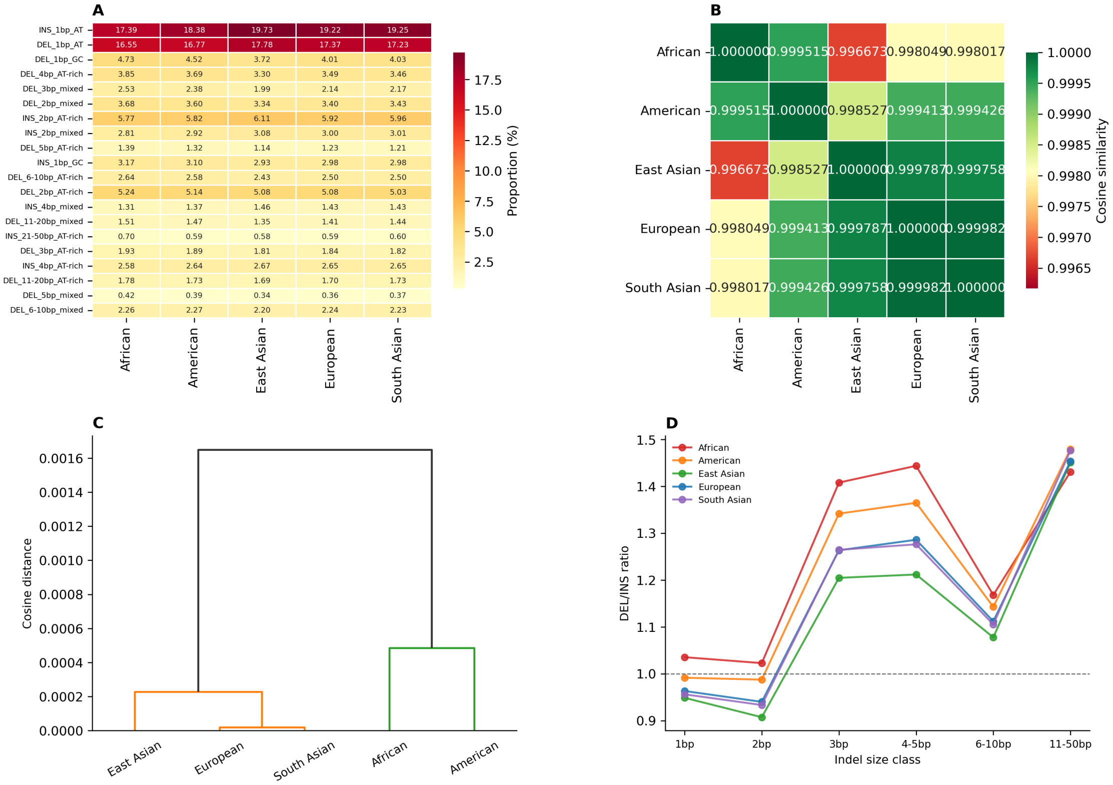

Figure 5: **Detailed population differentiation of indel spectra.** (**A**) Heatmap of the 20 most variable indel channels across super-populations (proportions shown as percentages). (**B**) Pairwise cosine similarity between super-population indel spectra. (**C**) Hierarchical clustering (average linkage, cosine distance) of super-populations based on full indel spectra. (**D**) DEL/INS ratio stratified by aggregated size class and super-population.

### 4 Discussion

Our analysis reveals previously uncharacterized population-level variation in the germline indel mutation spectrum across diverse human populations. Using the high-coverage 1000 Genomes Project dataset, we demonstrate that the compositional spectrum of germline indels—defined by type, size, base composition, and sequence context—differs significantly across continental super-populations. These differences are not merely a consequence of varying total indel burden (which is expected from demographic history) but reflect population-specific variation in the relative contributions of distinct mutagenic processes.

##### 4.1 Differential Mutagenic Process Contributions

The most striking finding is the elevated DEL/INS ratio and enrichment of GC-context deletions in African populations relative to other super-populations. Several non-mutually exclusive mechanisms could explain this pattern:

First, population-specific differences in oxidative DNA damage profiles could affect the rate of GC-targeted mutagenesis. 8-oxoguanine, a common oxidative lesion, preferentially affects guanine residues and can lead to deletions during error-prone repair [David et al., 2007]. Environmental and lifestyle factors that influence oxidative stress levels could contribute to population-level variation in GC-deletion rates.

Second, differences in the efficiency or fidelity of specific DNA repair pathways—particularly mismatch repair (MMR) and base excision repair (BER)—could modulate the indel spectrum. Population-specific polymorphisms in DNA repair genes have been documented [?] and could influence the relative rates of different indel-generating mechanisms.

Third, the enrichment of multi-base deletions (3–10 bp) in African populations suggests elevated rates of microhomology-mediated end joining (MMEJ) or alternative end joining pathways. These repair mechanisms generate deletions with characteristic microhomology signatures at junctions [McVey & Lee, 2008] and may vary in activity across populations.

The relative enrichment of AT-context insertions in East Asian populations, predominantly occurring in homopolymer runs, points to population-specific variation in replication slippage rates or in the efficiency of post-replicative MMR at A/T homopolymers. This could reflect subtle population differences in DNA polymerase processivity or proofreading fidelity, potentially modulated by polymorphisms in polymerase subunits or accessory factors.

##### 4.2 Relationship to Known Population Structure

The hierarchical clustering of populations based on indel spectra recapitulates the expected continental population structure (Figure 2d), with AFR as the most divergent group and EUR-SAS as the closest pair. This pattern is consistent with the out-of-Africa demographic model [Henn et al., 2012] but provides complementary information to SNV-based population structure. Importantly, the spectral differences we observe are proportional rather than absolute—all populations share the same basic indel spectrum dominated by AT-context 1 bp events in homopolymers, but the relative proportions differ in ways that are statistically significant and biologically interpretable.

##### 4.3 Comparison with Somatic Indel Signatures

The germline indel signatures identified by our NMF decomposition show notable parallels with COSMIC somatic indel signatures. Our Signature 1 (AT-slippage) shares features with COSMIC ID1 (insertion of T at long T homopolymers) and ID2 (deletion of T at long T homopolymers), both associated with replication slippage and defective MMR [Alexandrov et al., 2020]. Our Signature 2 (GC-deletion) has similarities with signatures associated with DNA damage-induced deletions. However, the germline context differs fundamentally from the somatic context: germline indels accumulate over generations and are subject to purifying selection, while somatic signatures reflect acute mutagenic processes within individual lifetimes. The population-level variation in germline signature contributions may reflect long-term differences in mutagenic exposures and repair capacities that have been shaped by evolutionary history.

##### 4.4 Limitations and Future Directions

Several limitations should be noted. First, indel detection from short-read sequencing remains less accurate than SNV detection, particularly for larger indels and those in repetitive regions [Chaisson et al., 2019]. While the 30× coverage of the 1000GP dataset substantially improves indel calling compared to low-coverage data, some systematic biases may persist. Second, our NMF decomposition was performed on super-population-level spectra rather than individual-level spectra, limiting the resolution of signature extraction. Future work incorporating individual-level data with larger sample sizes could enable more refined signature decomposition. Third, our analysis focused on autosomes; extension to sex chromosomes and comparison with de novo mutation data could provide additional insights into the mechanisms of indel mutagenesis.

Future studies should integrate long-read sequencing data, which provides superior indel detection especially for larger structural variants, to validate and extend these findings. Additionally, examining the relationship between population-specific indel spectra and germline mutation rate variation, particularly in the context of de novo mutation studies, could help disentangle the contributions of mutation rate differences from selection and drift.

#### 5 Conclusion

We present the first comprehensive population-stratified analysis of the germline indel mutation spectrum using high-coverage whole genome sequencing data from 3,202 globally diverse individuals. Our findings reveal significant population-specific variation in indel spectra, with African populations showing enrichment for GC-context deletions and multi-base deletions, and East Asian populations showing enrichment for AT-context insertions in homopolymer runs. NMF decomposition identifies two principal germline indel signatures with differential population contributions, suggesting population-specific variation in the rates of distinct mutagenic processes. These results extend our understanding of human mutational processes beyond SNVs and demonstrate that the indel mutation spectrum carries population-informative signals reflecting the interplay of mutagenesis, repair, and demographic history.

#### Data Availability

The 1000 Genomes Project high-coverage dataset is publicly available from the International Genome Sample Resource (`https://www.internationalgenome.org/`). Analysis code is available upon request.

### Acknowledgments

This study used data generated by the 1000 Genomes Project and the International Genome Sample Resource. We thank the participants who contributed samples and the research teams who generated the sequencing and variant calling data.

#### Supplementary Tables

Table 1: Top 15 most population-differentiated indel channels. Chi-square statistics, Bonferroni-adjusted $p$-values, and proportions across super-populations are shown. All channels are significant at $\alpha = 0.05$ after correction.

| Channel | $\chi^2$ | $p_{\text{adj}}$ | AFR | AMR | EAS | EUR | SAS |
|---|---|---|---|---|---|---|---|
| DEL_1bp_AT | 7187 | $< 10^{-300}$ | 0.1654 | 0.1675 | 0.1776 | 0.1739 | 0.1724 |
| DEL_1bp_GC | 5985 | $< 10^{-300}$ | 0.0482 | 0.0461 | 0.0380 | 0.0411 | 0.0413 |
| INS_1bp_AT | 5466 | $< 10^{-300}$ | 0.1755 | 0.1858 | 0.1987 | 0.1945 | 0.1942 |
| DEL_2bp_AT-rich | 3583 | $< 10^{-300}$ | 0.0518 | 0.0506 | 0.0501 | 0.0500 | 0.0496 |
| DEL_4bp_AT-rich | 3567 | $< 10^{-300}$ | 0.0375 | 0.0362 | 0.0324 | 0.0344 | 0.0341 |
| DEL_2bp_mixed | 3134 | $< 10^{-300}$ | 0.0364 | 0.0354 | 0.0329 | 0.0335 | 0.0338 |
| DEL_3bp_mixed | 3071 | $< 10^{-300}$ | 0.0252 | 0.0235 | 0.0200 | 0.0214 | 0.0216 |
| INS_2bp_AT-rich | 2754 | $< 10^{-300}$ | 0.0589 | 0.0596 | 0.0625 | 0.0607 | 0.0613 |
| INS_1bp_GC | 2561 | $< 10^{-300}$ | 0.0326 | 0.0319 | 0.0303 | 0.0307 | 0.0308 |
| DEL_6-10bp_AT-rich | 2134 | $< 10^{-300}$ | 0.0266 | 0.0261 | 0.0243 | 0.0251 | 0.0251 |
| DEL_3bp_AT-rich | 1495 | $< 10^{-300}$ | 0.0191 | 0.0187 | 0.0178 | 0.0180 | 0.0179 |
| DEL_5bp_AT-rich | 1481 | $< 10^{-300}$ | 0.0133 | 0.0128 | 0.0109 | 0.0117 | 0.0115 |
| DEL_6-10bp_mixed | 1437 | $< 10^{-300}$ | 0.0246 | 0.0243 | 0.0240 | 0.0240 | 0.0240 |
| INS_4bp_AT-rich | 1298 | $< 10^{-278}$ | 0.0253 | 0.0253 | 0.0264 | 0.0259 | 0.0259 |
| DEL_11-20bp_AT-rich | 1291 | $< 10^{-276}$ | 0.0164 | 0.0162 | 0.0155 | 0.0155 | 0.0155 |

Table 2: Summary of pairwise Jensen-Shannon divergence between super-population indel spectra.

| | AFR | AMR | EAS | EUR | SAS |
|---|---|---|---|---|---|
| AFR | 0 | 0.0152 | 0.0378 | 0.0281 | 0.0273 |
| AMR | 0.0152 | 0 | 0.0262 | 0.0159 | 0.0156 |
| EAS | 0.0378 | 0.0262 | 0 | 0.0110 | 0.0117 |
| EUR | 0.0281 | 0.0159 | 0.0110 | 0 | 0.0034 |
| SAS | 0.0273 | 0.0156 | 0.0117 | 0.0034 | 0 |
